## Supplemental file 1 for "Automated hippocampal unfolding for morphometry and subfield segmentation with HippUnfold"

### HippUnfold Documentation

**HippUnfold Development Team**

**Sep 08, 2022**

### GETTING STARTED

**1 Installation** 3
- 1.1 Requirements 3
- 1.2 Comparison of methods for running HippUnfold 3

**2 Running HippUnfold with Docker** 7

**3 Running HippUnfold with Singularity** 11

**4 Command-line interface** 13
- 4.1 HippUnfold Command-line interface 13
- 4.2 Snakemake command-line interface 17

**5 Running HippUnfold on your data** 33
- 5.1 Selecting the modality to use 33
- 5.2 Selecting and excluding subjects to process 33
- 5.3 Known limitations for BIDS parsing 34
- 5.4 Parsing Non-BIDS datasets with custom paths 35

**6 Specialized scans** 37
- 6.1 Case 1: super high resolution 37
- 6.2 Case 2: one ex-vivo hemisphere 38

**7 Pipeline Details** 39
- 7.1 Overall workflow 39
- 7.2 Pre-processing 40
- 7.3 U-net segmentation 45
- 7.4 Template-based shape injection 45
- 7.5 Laplace & equivolume coordinates 46
- 7.6 Subfields processing 47
- 7.7 Generating warp files 47
- 7.8 Surface processing 48
- 7.9 Additional steps 48

**8 Output Files** 49
- 8.1 anat 49
- 8.2 surf 50
- 8.3 coords 54
- 8.4 warps 54
- 8.5 Additional Files 55

**9 Visualization** 57

This tool aims to automatically model the topological folding structure of the human hippocampus, and computationally unfold the hippocampus to segment subfields and generate hippocampal and dentate gyrus surfaces.

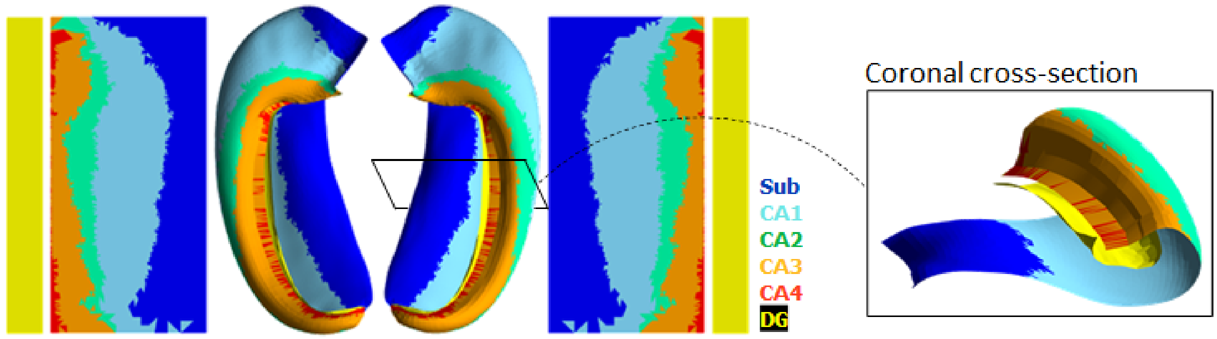

This is especially useful for:

- Visualization
- Topologically-constrained intersubject registration
- Parcellation (ie. registration to an unfolded atlas)
- Morphometry (eg. Thickness, Surface Area, Curvature, and Gyrification measures)

The overall workflow can be summarized in the following steps:

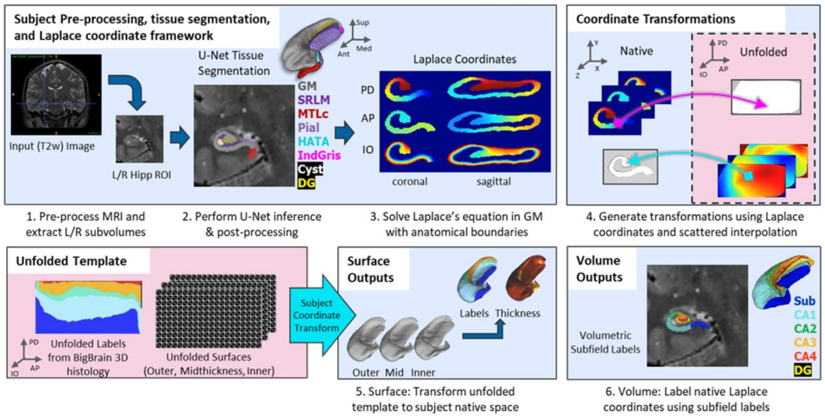

1. Pre-processing MRI images including non-uniformity correction, resampling to 0.3mm isotropic subvolumes, registration and cropping to coronal-oblique subvolumes around each hippocampus
2. Automatic segmentation of hippocampal tissues and surrounding structures via deep convolutional neural network U-net (nnU-net), models available for T1w, T2w, hippocampal b500 dwi, and neonatal T1w, and post-processing with fluid label-label registration to a high-resolution average template
3. Imposing of coordinates across the anterior-posterior and proximal-distal dimensions of hippocampal grey matter via solving Laplace's equation, and using an equivolume solution for laminar coordinates
4. Generating transformations between native and unfolded spaces using scattered interpolation on the hippocampus and dentate gyrus coordinates separately
5. Applying these transformations to generate surfaces meshes of the hippocampus and dentate gyrus, and extraction of morphological surface-based features including thickness, curvature, and gyrification index, sampled at the midthickness surface, and mapping subfield labels from the histological BigBrain atlas of the hippocampus
6. Generating high-resolution volumetric segmentations of the subfields using the same transformations and volumetric representations of the coordinates.

**Full Documentation:** here

**Additional toolbox** for plotting, mapping fMRI, DWI or other data, and manipulating surfaces here

**Relevant papers:**

- DeKraker J, Haast RAM, Yousif MD, Karat B, Köhler S, Khan AR. HippUnfold: Automated hippocampal unfolding, morphometry, and subfield segmentation. bioRxiv 2021.12.03.471134; doi: 10.1101/2021.12.03.471134 link
- DeKraker J, Ferko KM, Lau JC, Köhler S, Khan AR. Unfolding the hippocampus: An intrinsic coordinate system for subfield segmentations and quantitative mapping. Neuroimage. 2018 Feb 15;167:408-418. doi: 10.1016/j.neuroimage.2017.11.054. Epub 2017 Nov 23. PMID: 29175494. link
- DeKraker J, Lau JC, Ferko KM, Khan AR, Köhler S. Hippocampal subfields revealed through unfolding and unsupervised clustering of laminar and morphological features in 3D BigBrain. Neuroimage. 2020 Feb 1;206:116328. doi: 10.1016/j.neuroimage.2019.116328. Epub 2019 Nov 1. PMID: 31682982. link
- DeKraker J, Köhler S, Khan AR. Surface-based hippocampal subfield segmentation. Trends Neurosci. 2021 Nov;44(11):856-863. doi: 10.1016/j.tins.2021.06.005. Epub 2021 Jul 22. PMID: 34304910. link

CHAPTER

ONE

### INSTALLATION

BIDS App for Hippocampal AutoTop (automated hippocampal unfolding and subfield segmentation)

#### 1.1 Requirements

- Docker (Intel Mac/Windows/Linux) or Singularity (Linux)
- For those wishing to contribute or modify the code, `pip install` or `poetry install` are also available (Linux), but will still require singularity to handle some dependencies. See Contributing to HippUnfold.
- GPU not required
- Note: Apple M1 is currently **not supported**. We don't have a Docker arm64 container yet, and hippunfold is unusably slow with the emulated amd64 container.

##### 1.1.1 Notes:

- Inputs to Hippunfold should typically be a BIDS dataset including T1w images or T2w images. Higher-resolution data are preferred (<= 0.8mm) but the pipeline will still work with 1mm T1w images. See Tutorials.
- Other 3D imaging modalities (eg. ex-vivo MRI, 3D histology, etc.) can be used, but may require manual tissue segmentation as the current workflow relies on U-net segmentation trained only on common MRI modalities.

#### 1.2 Comparison of methods for running HippUnfold

There are several different ways of running HippUnfold. In order of increasing complexity/flexibility, we have:

1. CBRAIN Web-based Platform
2. Singularity Container on Linux
3. Docker Container on Windows/Mac (Intel)/Linux
4. Python Environment with Singularity Dependencies

##### 1.2.1 CBRAIN Web-based Platform

HippUnfold is available on the CBRAIN platform, a web-based platform for batch high-performance computing that is free for researchers.

###### Pros:

- No software installation required
- Fully point and click interface (no CLI)
- Can perform batch-processing

###### Cons:

- Must upload data for processing
- Limited command-line options exposed
- Cannot edit code

##### 1.2.2 Docker on Windows/Mac (Intel)/Linux

The HippUnfold BIDS App is available on a DockerHub as versioned releases and development branches.

###### Pros:

- Compatible with non-Linux systems
- All dependencies+models in a single container

###### Cons:

- Typically not possible on shared machines
- Cannot use Snakemake cluster execution profiles
- Cannot edit code

##### 1.2.3 Singularity Container

The same docker container can also be used with Singularity (now Apptainer). Instructions can be found below.

###### Pros:

- All dependencies+models in a single container
- Container stored as a single file (.sif)

###### Cons:

- Compatible on shared systems with Singularity installed
- Cannot use Snakemake cluster execution profiles
- Cannot edit code

##### 1.2.4 Python Environment with Singularity Dependencies

Instructions for this can be found in the **Contributing** documentation page.

###### Pros:

- Complete flexibility to modify code
- External (Non-Python) dependencies as Singularity containers

###### Cons:

- Must use Python virtual environment
- Only compatible on Linux systems with Singularity for external dependencies

CHAPTER

TWO

### RUNNING HIPPUNFOLD WITH DOCKER

Note: These instructions assume you have Docker installed already on your system.

Download and extract a single-subject BIDS dataset for this test:

```
wget https://www.dropbox.com/s/mdbmpmmq6fi8sk0/hippunfold_test_data.tar
tar -xvf hippunfold_test_data.tar
```

This will create a `ds002168/` folder with a single subject, that has a both T1w and T2w images.

```
ds002168/
├── dataset_description.json
├── README.md
└── sub-1425
    └── anat
        ├── sub-1425_T1w.json
        ├── sub-1425_T1w.nii.gz
        ├── sub-1425_T2w.json
        └── sub-1425_T2w.nii.gz

2 directories, 6 files
```

Pull the container:

```
docker pull khanlab/hippunfold:latest
```

Run HippUnfold without any arguments to print the short help:

```
docker run -it --rm \
khanlab/hippunfold:latest
```

Use the `-h` option to get a detailed help listing:

```
docker run -it --rm \
khanlab/hippunfold:latest \
-h
```

Note that all the Snakemake command-line options are also available in HippUnfold, and can be listed with `--help-snakemake`:

```
docker run -it --rm \
khanlab/hippunfold:latest \
--help-snakemake
```

Now let's run it on the test dataset. The `--modality` flag is a required argument, and describes what image we use for segmentation. Here we will use the T1w image. We will also use the `--dry-run/-n` option to just print out what would run, without actually running anything.

```
docker run -it --rm \
-v ds002168:/bids:ro \
-v ds002168_hippunfold:/output \
khanlab/hippunfold:latest \
/bids /output participant --modality T1w -n
```

For those not familiar with Docker, the first three lines of this example are generic Docker arguments to ensure it is run with the safest options and has permission to access your input and output directories The fourth line specifies the HippUnfold Docker container, and the fifth line contains the required arguments for HippUnfold, after which you can additionally specify optional arguments. You may want to familiarize yourself with Docker options, and an overview of HippUnfold arguments is provided in the Command line interface documentation section.

The first three arguments to HippUnfold (as with any BIDS App) are the input folder, the output folder, and then the analysis level. The `participant` analysis level is used in HippUnfold for performing the segmentation, unfolding, and any participant-level processing. The `group` analysis is used to combine subfield volumes across subjects into a single tsv file.

When you run the above command, a long listing will print out, describing all the rules that will be run. This is a long listing, and you can better appreciate it with the `less` tool. We can also have the shell command used for each rule printed to screen using the `-p` Snakemake option:

```
docker run -it --rm \
-v ds002168:/bids:ro \
-v ds002168_hippunfold:/output \
khanlab/hippunfold:latest \
/bids /output participant --modality T1w -np | less
```

Now, to actually run the workflow, we need to specify how many cores to use and leave out the dry-run option. The Snakemake `--cores` option tells HippUnfold how many cores to use. Using `--cores 8` means that HippUnfold will only make use of 8 cores at most. Generally speaking you should use `--cores all`, so it can make maximal use of all the CPU cores it has access to on your system. This is especially useful if you are running multiple subjects.

Running the following command (hippunfold on a single subject) may take ~30 minutes if you have 8 cores, shorter if you have more cores, but could be much longer (several hours) if you only have a single core.

```
docker run -it --rm \
-v ds002168:/bids:ro \
-v ds002168_hippunfold:/output \
khanlab/hippunfold:latest \
/bids /output participant --modality T1w -p --cores all
```

After this completes, you should have a `ds002168_hippunfold` folder with outputs for the one subject.

If you alternatively want to run HippUnfold using a different modality, e.g. the high-resolution T2w image in the BIDS test dataset, you can use the `--modality T2w` option. In this case, since the T2w image in the test dataset has a limited FOV, we should also make use of the `--t1-reg-template` command-line option, which will make use of the T1w image for template registration, since a limited FOV T2w template does not exist.

```
docker run -it --rm \
-v ds002168:/bids:ro \
-v ds002168_hippunfold_t2w:/output \
```

(continues on next page)

(continued from previous page)

```
khanlab/hippunfold:latest \
/bids /output participant --modality T2w --t1-reg-template -p --cores all
```

Note that if you run with a different modality, you should use a separate output folder, since some of the files would be overwritten if not.

CHAPTER

THREE

### RUNNING HIPPUNFOLD WITH SINGULARITY

Note: These instructions assume you have Singularity installed on your system.

Download and extract a single-subject BIDS dataset for this test:

```
wget https://www.dropbox.com/s/mdbmpmmq6fi8sk0/hippunfold_test_data.tar
tar -xvf hippunfold_test_data.tar
```

This will create a `ds002168/` folder with a single subject, that has a both T1w and T2w images:

```
ds002168/
├── dataset_description.json
├── README.md
└── sub-1425
    └── anat
        ├── sub-1425_T1w.json
        ├── sub-1425_T1w.nii.gz
        ├── sub-1425_T2w.json
        └── sub-1425_T2w.nii.gz

2 directories, 6 files
```

Next we will pull the container from dockerhub:

```
singularity pull khanlab_hippunfold_latest.sif docker://khanlab/hippunfold:latest
```

If you encounter any errors pulling the container, it may be because you are running out of disk space in your cache folders. Note, you can change these locations by setting environment variables, e.g.:

```
export SINGULARITY_CACHEDIR=/YOURDIR/.cache/singularity
```

Run HippUnfold any arguments to print the short help:

```
singularity run -e khanlab_hippunfold_latest.sif
```

Use the `-h` option to get a detailed help listing:

```
singularity run -e khanlab_hippunfold_latest.sif -h
```

Note that all the Snakemake command-line options are also available in HippUnfold, and can be listed with `--help-snakemake`:

```
singularity run -e khanlab_hippunfold_latest.sif --help-snakemake
```

Now let's run it on the test dataset. The --modality flag is a required argument, and describes what image we use for segmentation. Here we will use the T1w image. We will also use the --dry-run/-n option to just print out what would run, without actually running anything.

```
singularity run -e khanlab_hippunfold_latest.sif \
ds002168 ds002168_hippunfold participant -n --modality T1w
```

The first three arguments to HippUnfold (as with any BIDS App) are the input folder, the output folder, and then the analysis level. The participant analysis level is used in HippUnfold for performing the segmentation, unfolding, and any participant-level processing. The group analysis is used to combine subfield volumes across subjects into a single tsv file.

When you run the above command, a long listing will print out, describing all the rules that will be run. This is a long listing, and you can better appreciate it with the less tool. We can also have the shell command used for each rule printed to screen using the -p Snakemake option:

```
singularity run -e khanlab_hippunfold_latest.sif \
ds002168 ds002168_hippunfold participant -np --modality T1w | less
```

Now, to actually run the workflow, we need to specify how many cores to use and leave out the dry-run option. The Snakemake --cores option tells HippUnfold how many cores to use. Using --cores 8 means that HippUnfold will only make use of 8 cores at most. Generally speaking you should use --cores all, so it can make maximal use of all the CPU cores it has access to on your system. This is especially useful if you are running multiple subjects.

Running the following command (hippunfold on a single subject) may take ~30 minutes if you have 8 cores, shorter if you have more cores, but could be much longer (several hours) if you only have a single core.

```
singularity run -e khanlab_hippunfold_latest.sif \
ds002168 ds002168_hippunfold participant -p --cores all --modality T1w
```

Note that you may need to adjust your Singularity options to ensure the container can read and write to yout input and output directories, respectively. You can bind paths easily by setting an environment variable, e.g. if you have a /project folder that contains your data, you can add it to the SINGULARITY_BINDPATH so it is available when you are running a container:

```
export SINGULARITY_BINDPATH=/data:/data
```

After this completes, you should have a ds002168_hippunfold folder with outputs for the one subject.

If you alternatively want to run HippUnfold using a different modality, e.g. the high-resolution T2w image in the BIDS test dataset, you can use the --modality T2w option. In this case, since the T2w image in the test dataset has a limited FOV, we should also make use of the --t1-reg-template command-line option, which will make use of the T1w image for template registration, since a limited FOV T2w template does not exist.

```
singularity run -e khanlab_hippunfold_latest.sif \
ds002168 ds002168_hippunfold_t2w participant --modality T2w --t1-reg-template -p --cores␣
↪all
```

Note that if you run with a different modality, you should use a separate output folder, since some of the files would be overwritten if not.

CHAPTER

FOUR

### COMMAND-LINE INTERFACE

#### 4.1 HippUnfold Command-line interface

The following can also be seen by entering `hippunfold -h` into your terminal.

These are all the required and optional arguments HippUnfold accepts in order to run flexibly on many different input data types and with many options, but in most cases only the required arguments are needed.

Snakebids helps build BIDS Apps with Snakemake

```
usage: hippunfold [-h] [--pybidsdb-dir PYBIDSDB_DIR] [--reset-db] [--force-output] [--
↪help-snakemake] [--participant-label PARTICIPANT_LABEL [PARTICIPANT_LABEL ...]]
                  [--exclude_participant_label EXCLUDE_PARTICIPANT_LABEL [EXCLUDE_
↪PARTICIPANT_LABEL ...]] --modality {T1w,T2w,hippb500,segT1w,segT2w,cropseg} [--
↪derivatives DERIVATIVES] [--skip-preproc]
                  [--skip-coreg] [--skip_inject_template_labels] [--inject_template_
↪smoothing_factor INJECT_TEMPLATE_SMOOTHING_FACTOR] [--rigid-reg-template] [--no-reg-
↪template] [--template {CITI168,dHCP}]
                  [--t1_reg_template] [--crop-native-box CROP_NATIVE_BOX] [--atlas
↪{bigbrain,magdeburg,freesurfer} [{bigbrain,magdeburg,freesurfer} ...]] [--generate-
↪myelin-map] [--use-gpu] [--nnunet_enable_tta]
                  [--output_spaces {native,T1w} [{native,T1w} ...]] [--output_density
↪{0p5mm,1mm,2mm,unfoldiso} [{0p5mm,1mm,2mm,unfoldiso} ...]] [--hemi {L,R} [{L,R} ...]]
                  [--laminar-coords-method {laplace,equivolume}] [--keep_work] [--force-
↪nnunet-model {T1w,T2w,T1T2w,b1000,trimodal,hippb500,neonateT1w}] [--filter-T2w FILTER_
↪T2W [FILTER_T2W ...]]
                  [--filter-hippb500 FILTER_HIPPB500 [FILTER_HIPPB500 ...]] [--filter-
↪T1w FILTER_T1W [FILTER_T1W ...]] [--filter-seg FILTER_SEG [FILTER_SEG ...]] [--filter-
↪cropseg FILTER_CROPSEG [FILTER_CROPSEG ...]]
                  [--wildcards-T2w WILDCARDS_T2W [WILDCARDS_T2W ...]] [--wildcards-
↪hippb500 WILDCARDS_HIPPB500 [WILDCARDS_HIPPB500 ...]] [--wildcards-T1w WILDCARDS_T1W␣
↪[WILDCARDS_T1W ...]]
                  [--wildcards-seg WILDCARDS_SEG [WILDCARDS_SEG ...]] [--wildcards-
↪cropseg WILDCARDS_CROPSEG [WILDCARDS_CROPSEG ...]] [--path-T2w PATH_T2W] [--path-
↪hippb500 PATH_HIPPB500] [--path-T1w PATH_T1W]
                  [--path-seg PATH_SEG] [--path-cropseg PATH_CROPSEG]
                  bids_dir output_dir {participant,group}
```

##### 4.1.1 STANDARD

Standard options for all snakebids apps

**--pybidsdb-dir, --pybidsdb_dir** Optional path to directory of SQLite databasefile for PyBIDS. If directory is passed and folder exists, indexing is skipped. If reset_db is called, indexing will persist

**--reset-db, --reset_db** Reindex existing PyBIDS SQLite database

Default: False

**--force-output, --force_output** Force output in a new directory that already has contents

Default: False

**--help-snakemake, --help_snakemake** Options to Snakemake can also be passed directly at the command-line, use this to print Snakemake usage

##### 4.1.2 SNAKEBIDS

Options for snakebids app

**bids_dir** The directory with the input dataset formatted according to the BIDS standard.

**output_dir** The directory where the output files should be stored. If you are running group level analysis this folder should be prepopulated with the results of the participant level analysis.

**analysis_level** Possible choices: participant, group

Level of the analysis that will be performed.

**--participant-label, --participant_label** The label(s) of the participant(s) that should be analyzed. The label corresponds to sub-<participant_label> from the BIDS spec (so it does not include "sub-"). If this parameter is not provided all subjects should be analyzed. Multiple participants can be specified with a space separated list.

**--exclude_participant_label, --exclude-participant-label** The label(s) of the participant(s) that should be excluded. The label corresponds to sub-<participant_label> from the BIDS spec (so it does not include "sub-"). If this parameter is not provided all subjects should be analyzed. Multiple participants can be specified with a space separated list.

**--modality** Possible choices: T1w, T2w, hippb500, segT1w, segT2w, cropseg

Type of image to run hippunfold on. Modality prefixed with seg will import an existing (manual) hippocampal tissue segmentation from that space, instead of running neural network (default: None)

**--derivatives** Path to the derivatives folder (e.g. for finding manual segs) (default: False)

Default: False

**--skip-preproc, --skip_preproc** Set this flag if your inputs (e.g. T2w, dwi) are already pre-processed (default: False)

Default: False

**--skip-coreg, --skip_coreg** Set this flag if your inputs (e.g. T2w, dwi) are already registered to T1w space (default: False)

Default: False

**--skip_inject_template_labels, --skip-inject-template-labels** Set this flag to skip post-processing template injection into CNN segmentation. Note this will disable generation of DG surfaces. (default: False)

Default: False

**--inject_template_smoothing_factor, --inject-template-smoothing-factor** Scales the default smoothing sigma for gradient and warp in template shape injection. Using a value higher than 1 will use result in a smoother warp, and greater capacity to patch larger holes in segmentations. Try setting to 2 if nnunet segmentations have large holes. Note: the better solution is to re-train network on the data you are using (default: 1.0)

Default: 1.0

**--rigid-reg-template, --rigid_reg_template** Use rigid instead of affine for registration to template. Try this if your images are reduced FOV (default: False)

Default: False

**--no-reg-template, --no_reg_template** Use if input data is already in space-CITI168 (default: False)

Default: False

**--template** Possible choices: CITI168, dHCP

Set the template to use for registration to coronal oblique. (default: “CITI168”)

Default: “CITI168”

**--t1_reg_template, --t1-reg-template** Use T1w to register to template space, instead of the segmentation modality. Note: this was the default behavior prior to v1.0.0. (default: False)

Default: False

**--crop-native-box, --crop_native_box** Sets the bounding box size for the crop native (e.g. cropT1w) space). Make this larger if your hippocampi in crop{T1w,T2w} space are getting cut-off (default: “256x256x256vox”)

Default: “256x256x256vox”

**--atlas** Possible choices: bigbrain, magdeburg, freesurfer

Select the atlas (unfolded space) to use for subfield labels. (default: [‘bigbrain’])

Default: [‘bigbrain’]

**--generate-myelin-map, --generate_myelin_map** Generate myelin map using T1w divided by T2w, and map to surface with ribbon approach. Requires both T1w and T2w images to be present. (default: False)

Default: False

**--use-gpu, --use_gpu** Enable gpu for inference by setting resource gpus=1 in run_inference rule (default: False)

Default: False

**--nnunet_enable_tta, --nnunet-enable-tta** Enable test-time augmentation for nnU-net inference, slows down inference by 8x, but potentially increases accuracy (default: False)

Default: False

**--output_spaces, --output-spaces** Possible choices: native, T1w

Sets output spaces for results (default: [‘native’])

Default: [‘native’]

**--output_density, --output-density** Possible choices: 0p5mm, 1mm, 2mm, unfoldiso

Sets the output vertex density for results. Options correspond to approximate vertex spacings of 0.5mm, 1.0mm, and 2.0mm, respectively, with the unfoldiso (32k hipp) vertices legacy option having unequal vertex spacing. (default: [‘0p5mm’])

Default: [‘0p5mm’]

**--hemi** Possible choices: L, R

Hemisphere(s) to process (default: [‘L’, ‘R’])

Default: [‘L’, ‘R’]

**--laminar-coords-method, --laminar_coords_method** Possible choices: laplace, equivolume

Method to use for laminar coordinates. Equivolume uses equivolumetric layering from Waehnert et al 2014 (Nighres implementation). (default: [‘equivolume’])

Default: [‘equivolume’]

**--keep_work, --keep-work** Keep work folder intact instead of archiving it for each subject (default: False)

Default: False

**--force-nnunet-model, --force_nnunet_model** Possible choices: T1w, T2w, T1T2w, b1000, trimodal, hippb500, neonateT1w

Force nnunet model to use (expert option). (default: False)

Default: False

##### 4.1.3 BIDS FILTERS

Filters to customize PyBIDS get() as key=value pairs

**--filter-T2w, --filter_T2w** (default: suffix=T2w extension=.nii.gz datatype=anat invalid_filters=allow space=None)

**--filter-hippb500, --filter_hippb500** (default: suffix=b500 extension=.nii.gz invalid_filters=allow datatype=dwi)

**--filter-T1w, --filter_T1w** (default: suffix=T1w extension=.nii.gz datatype=anat invalid_filters=allow space=None)

**--filter-seg, --filter_seg** (default: suffix=dseg extension=.nii.gz datatype=anat invalid_filters=allow)

**--filter-cropseg, --filter_cropseg** (default: suffix=dseg extension=.nii.gz datatype=anat invalid_filters=allow)

##### 4.1.4 INPUT WILDCARDS

File path entities to use as wildcards in snakemake

**--wildcards-T2w, --wildcards_T2w** (default: subject session acquisition run)

**--wildcards-hippb500, --wildcards_hippb500** (default: subject session)

**--wildcards-T1w, --wildcards_T1w** (default: subject session acquisition run)

**--wildcards-seg, --wildcards_seg** (default: subject session)

**--wildcards-cropseg, --wildcards_cropseg** (default: subject session hemi)

##### 4.1.5 PATH OVERRIDE

Options for overriding BIDS by specifying absolute paths that include wildcards, e.g.: /path/to/my_data/{subject}/t1.nii.gz

**--path-T2w, --path_T2w**

**--path-hippb500, --path_hippb500**

**--path-T1w, --path_T1w**

**--path-seg, --path_seg**

**--path-cropseg, --path_cropseg**

#### 4.2 Snakemake command-line interface

In addition to the above command-line arguments, Snakemake arguments are also be passed at the `hippunfold` command-line.

The most critical of these is the `--cores` or `-c` argument, which is a **required** argument for HippUnfold.

The complete list of Snakemake arguments are below, and mostly act to determine your environment and App behaviours. They will likely only need to be used for running in cloud environments or troubleshooting. These can be listed from the command-line with `hippunfold --help-snakemake`.

Snakemake is a Python based language and execution environment for GNU Make-like workflows.

```
usage: snakemake [-h] [--dry-run] [--profile PROFILE] [--cache [RULE [RULE ...]]] [--
↪snakefile FILE] [--cores [N]] [--jobs [N]] [--local-cores N] [--resources [NAME=INT␣
↪[NAME=INT ...]]]
                 [--set-threads RULE=THREADS [RULE=THREADS ...]] [--max-threads MAX_
↪THREADS] [--set-resources RULE:RESOURCE=VALUE [RULE:RESOURCE=VALUE ...]] [--set-
↪scatter NAME=SCATTERITEMS [NAME=SCATTERITEMS ...]]
                 [--default-resources [NAME=INT [NAME=INT ...]]] [--preemption-default␣
↪PREEMPTION_DEFAULT] [--preemptible-rules PREEMPTIBLE_RULES [PREEMPTIBLE_RULES ...]] [--
↪config [KEY=VALUE [KEY=VALUE ...]]]
                 [--configfile FILE [FILE ...]] [--envvars VARNAME [VARNAME ...]] [--
↪directory DIR] [--touch] [--keep-going]
                 [--rerun-triggers {mtime,params,input,software-env,code} [{mtime,params,
↪input,software-env,code} ...]] [--force] [--forceall] [--forcerun [TARGET [TARGET ...
↪]]] [--prioritize TARGET [TARGET ...]]
```

(continues on next page)

(continued from previous page)

```
                  [--batch RULE=BATCH/BATCHES] [--until TARGET [TARGET ...]] [--omit-from␣
↪TARGET [TARGET ...]] [--rerun-incomplete] [--shadow-prefix DIR] [--scheduler [{ilp,
↪greedy}]] [--wms-monitor [WMS_MONITOR]]
                  [--wms-monitor-arg [NAME=VALUE [NAME=VALUE ...]]] [--scheduler-ilp-
↪solver {PULP_CBC_CMD}] [--scheduler-solver-path SCHEDULER_SOLVER_PATH] [--conda-base-
↪path CONDA_BASE_PATH] [--no-subworkflows]
                  [--groups GROUPS [GROUPS ...]] [--group-components GROUP_COMPONENTS␣
↪[GROUP_COMPONENTS ...]] [--report [FILE]] [--report-stylesheet CSSFILE] [--draft-
↪notebook TARGET] [--edit-notebook TARGET]
                  [--notebook-listen IP:PORT] [--lint [{text,json}]] [--generate-unit-
↪tests [TESTPATH]] [--containerize] [--export-cwl FILE] [--list] [--list-target-rules]␣
↪[--dag] [--rulegraph] [--filegraph] [--d3dag]
                  [--summary] [--detailed-summary] [--archive FILE] [--cleanup-metadata␣
↪FILE [FILE ...]] [--cleanup-shadow] [--skip-script-cleanup] [--unlock] [--list-version-
↪changes] [--list-code-changes]
                  [--list-input-changes] [--list-params-changes] [--list-untracked] [--
↪delete-all-output] [--delete-temp-output] [--bash-completion] [--keep-incomplete] [--
↪drop-metadata] [--version] [--reason]
                  [--gui [PORT]] [--printshellcmds] [--debug-dag] [--stats FILE] [--
↪nocolor] [--quiet [{progress,rules,all} [{progress,rules,all} ...]]] [--print-
↪compilation] [--verbose] [--force-use-threads]
                  [--allow-ambiguity] [--nolock] [--ignore-incomplete] [--max-inventory-
↪time SECONDS] [--latency-wait SECONDS] [--wait-for-files [FILE [FILE ...]]] [--wait-
↪for-files-file FILE] [--notemp] [--all-temp]
                  [--keep-remote] [--keep-target-files] [--allowed-rules ALLOWED_RULES␣
↪[ALLOWED_RULES ...]] [--local-groupid LOCAL_GROUPID] [--max-jobs-per-second MAX_JOBS_
↪PER_SECOND]
                  [--max-status-checks-per-second MAX_STATUS_CHECKS_PER_SECOND] [-T␣
↪RETRIES] [--attempt ATTEMPT] [--wrapper-prefix WRAPPER_PREFIX]
                  [--default-remote-provider {S3,GS,FTP,SFTP,S3Mocked,gfal,gridftp,iRODS,
↪AzBlob,XRootD}] [--default-remote-prefix DEFAULT_REMOTE_PREFIX] [--no-shared-fs] [--
↪greediness GREEDINESS] [--no-hooks]
                  [--overwrite-shellcmd OVERWRITE_SHELLCMD] [--debug] [--runtime-profile␣
↪FILE] [--mode {0,1,2}] [--show-failed-logs] [--log-handler-script FILE] [--log-service
↪{none,slack,wms}]
                  [--cluster CMD | --cluster-sync CMD | --drmaa [ARGS]] [--cluster-config␣
↪FILE] [--immediate-submit] [--jobscript SCRIPT] [--jobname NAME] [--cluster-status␣
↪CLUSTER_STATUS]
                  [--cluster-cancel CLUSTER_CANCEL] [--cluster-cancel-nargs CLUSTER_
↪CANCEL_NARGS] [--cluster-sidecar CLUSTER_SIDECAR] [--drmaa-log-dir DIR] [--kubernetes␣
↪[NAMESPACE]] [--container-image IMAGE]
                  [--tibanna] [--tibanna-sfn TIBANNA_SFN] [--precommand PRECOMMAND] [--
↪tibanna-config TIBANNA_CONFIG [TIBANNA_CONFIG ...]] [--google-lifesciences]
                  [--google-lifesciences-regions GOOGLE_LIFESCIENCES_REGIONS [GOOGLE_
↪LIFESCIENCES_REGIONS ...]] [--google-lifesciences-location GOOGLE_LIFESCIENCES_
↪LOCATION] [--google-lifesciences-keep-cache]
                  [--tes URL] [--use-conda] [--conda-not-block-search-path-envvars] [--
↪list-conda-envs] [--conda-prefix DIR] [--conda-cleanup-envs] [--conda-cleanup-pkgs [
↪{tarballs,cache}]] [--conda-create-envs-only]
                  [--conda-frontend {conda,mamba}] [--use-singularity] [--singularity-
↪prefix DIR] [--singularity-args ARGS] [--use-envmodules]
                  [target [target ...]]
```

##### 4.2.1 EXECUTION

**target** Targets to build. May be rules or files.

**--dry-run, --dryrun, -n** Do not execute anything, and display what would be done. If you have a very large workflow, use –dry-run –quiet to just print a summary of the DAG of jobs.

Default: False

**--profile** Name of profile to use for configuring Snakemake. Snakemake will search for a corresponding folder in /etc/xdg/xdg-ubuntu/snakemake and /home/ROBARTS/alik/.config/snakemake. Alternatively, this can be an absolute or relative path. The profile folder has to contain a file 'config.yaml'. This file can be used to set default values for command line options in YAML format. For example, '–cluster qsub' becomes 'cluster: qsub' in the YAML file. Profiles can be obtained from https://github.com/snakemake-profiles. The profile can also be set via the environment variable $SNAKEMAKE_PROFILE.

**--cache** Store output files of given rules in a central cache given by the environment variable $SNAKEMAKE_OUTPUT_CACHE. Likewise, retrieve output files of the given rules from this cache if they have been created before (by anybody writing to the same cache), instead of actually executing the rules. Output files are identified by hashing all steps, parameters and software stack (conda envs or containers) needed to create them.

**--snakefile, -s** The workflow definition in form of a snakefile.Usually, you should not need to specify this. By default, Snakemake will search for 'Snakefile', 'snakefile', 'workflow/Snakefile', 'workflow/snakefile' beneath the current working directory, in this order. Only if you definitely want a different layout, you need to use this parameter.

**--cores, -c** Use at most N CPU cores/jobs in parallel. If N is omitted or 'all', the limit is set to the number of available CPU cores. In case of cluster/cloud execution, this argument sets the number of total cores used over all jobs (made available to rules via workflow.cores).

**--jobs, -j** Use at most N CPU cluster/cloud jobs in parallel. For local execution this is an alias for –cores.

**--local-cores** In cluster/cloud mode, use at most N cores of the host machine in parallel (default: number of CPU cores of the host). The cores are used to execute local rules. This option is ignored when not in cluster/cloud mode.

Default: 12

**--resources, --res** Define additional resources that shall constrain the scheduling analogously to threads (see above). A resource is defined as a name and an integer value. E.g. –resources mem_mb=1000. Rules can use resources by defining the resource keyword, e.g. resources: mem_mb=600. If now two rules require 600 of the resource 'mem_mb' they won't be run in parallel by the scheduler.

**--set-threads** Overwrite thread usage of rules. This allows to fine-tune workflow parallelization. In particular, this is helpful to target certain cluster nodes by e.g. shifting a rule to use more, or less threads than defined in the workflow. Thereby, THREADS has to be a positive integer, and RULE has to be the name of the rule.

**--max-threads** Define a global maximum number of threads for any job. This can be helpful in a cluster/cloud setting, when you want to restrict the maximum number of requested threads without modifying the workflow definition or overwriting them invidiually with –set-threads.

**--set-resources** Overwrite resource usage of rules. This allows to fine-tune workflow resources. In particular, this is helpful to target certain cluster nodes by e.g. defining a certain partition for a rule, or overriding a temporary directory. Thereby, VALUE has to be a positive integer or a string, RULE has to be the name of the rule, and RESOURCE has to be the name of the resource.

**--set-scatter** Overwrite number of scatter items of scattergather processes. This allows to fine-tune workflow parallelization. Thereby, SCATTERITEMS has to be a positive integer, and NAME has to be the name of the scattergather process defined via a scattergather directive in the workflow.

**--default-resources, --default-res** Define default values of resources for rules that do not define their own values. In addition to plain integers, python expressions over inputsize are allowed (e.g. '2*input.size_mb').When specifying this without any arguments (–default-resources), it defines 'mem_mb=max(2*input.size_mb, 1000)' 'disk_mb=max(2*input.size_mb, 1000)' i.e., default disk and mem usage is twice the input file size but at least 1GB.In addition, the system temporary directory (as given by $TMPDIR, $TEMP, or $TMP) is used for the tmpdir resource. The tmpdir resource is automatically used by shell commands, scripts and wrappers to store temporary data (as it is mirrored into $TMPDIR, $TEMP, and $TMP for the executed subprocesses). If this argument is not specified at all, Snakemake just uses the tmpdir resource as outlined above.

**--preemption-default** A preemptible instance can be requested when using the Google Life Sciences API. If you set a –preemption-default,all rules will be subject to the default. Specifically, this integer is the number of restart attempts that will be made given that the instance is killed unexpectedly. Note that preemptible instances have a maximum running time of 24 hours. If you want to set preemptible instances for only a subset of rules, use –preemptible-rules instead.

**--preemptible-rules** A preemptible instance can be requested when using the Google Life Sciences API. If you want to use these instances for a subset of your rules, you can use –preemptible-rules and then specify a list of rule and integer pairs, where each integer indicates the number of restarts to use for the rule's instance in the case that the instance is terminated unexpectedly. –preemptible-rules can be used in combination with –preemption-default, and will take priority. Note that preemptible instances have a maximum running time of 24. If you want to apply a consistent number of retries across all your rules, use –premption-default instead. Example: snakemake –preemption-default 10 –preemptible-rules map_reads=3 call_variants=0

**--config, -C** Set or overwrite values in the workflow config object. The workflow config object is accessible as variable config inside the workflow. Default values can be set by providing a JSON file (see Documentation).

**--configfile, --configfiles** Specify or overwrite the config file of the workflow (see the docs). Values specified in JSON or YAML format are available in the global config dictionary inside the workflow. Multiple files overwrite each other in the given order. Thereby missing keys in previous config files are extended by following configfiles. Note that this order also includes a config file defined in the workflow definition itself (which will come first).

**--envvars** Environment variables to pass to cloud jobs.

**--directory, -d** Specify working directory (relative paths in the snakefile will use this as their origin).

**--touch, -t** Touch output files (mark them up to date without really changing them) instead of running their commands. This is used to pretend that the rules were executed, in order to fool future invocations of snakemake. Fails if a file does not yet exist. Note that this will only touch files that would otherwise be recreated by Snakemake (e.g. because their input files are newer). For enforcing a touch, combine this with –force, –forceall, or –forcerun. Note however that you loose the provenance information when the files have been created in realitiy. Hence, this should be used only as a last resort.

Default: False

**--keep-going, -k** Go on with independent jobs if a job fails.

Default: False

**--rerun-triggers** Possible choices: mtime, params, input, software-env, code

Define what triggers the rerunning of a job. By default, all triggers are used, which guarantees that results are consistent with the workflow code and configuration. If you rather prefer the traditional way of just considering file modification dates, use '–rerun-trigger mtime'.

Default: ['mtime', 'params', 'input', 'software-env', 'code']

**--force, -f** Force the execution of the selected target or the first rule regardless of already created output.

Default: False

**--forceall, -F** Force the execution of the selected (or the first) rule and all rules it is dependent on regardless of already created output.

Default: False

**--forcerun, -R** Force the re-execution or creation of the given rules or files. Use this option if you changed a rule and want to have all its output in your workflow updated.

**--prioritize, -P** Tell the scheduler to assign creation of given targets (and all their dependencies) highest priority. (EXPERIMENTAL)

**--batch** Only create the given BATCH of the input files of the given RULE. This can be used to iteratively run parts of very large workflows. Only the execution plan of the relevant part of the workflow has to be calculated, thereby speeding up DAG computation. It is recommended to provide the most suitable rule for batching when documenting a workflow. It should be some aggregating rule that would be executed only once, and has a large number of input files. For example, it can be a rule that aggregates over samples.

**--until, -U** Runs the pipeline until it reaches the specified rules or files. Only runs jobs that are dependencies of the specified rule or files, does not run sibling DAGs.

**--omit-from, -O** Prevent the execution or creation of the given rules or files as well as any rules or files that are downstream of these targets in the DAG. Also runs jobs in sibling DAGs that are independent of the rules or files specified here.

**--rerun-incomplete, --ri** Re-run all jobs the output of which is recognized as incomplete.

Default: False

**--shadow-prefix** Specify a directory in which the 'shadow' directory is created. If not supplied, the value is set to the '.snakemake' directory relative to the working directory.

**--scheduler** Possible choices: ilp, greedy

Specifies if jobs are selected by a greedy algorithm or by solving an ilp. The ilp scheduler aims to reduce runtime and hdd usage by best possible use of resources.

Default: "greedy"

**--wms-monitor** IP and port of workflow management system to monitor the execution of snakemake (e.g. http://127.0.0.1:5000) Note that if your service requires an authorization token, you must export WMS_MONITOR_TOKEN in the environment.

**--wms-monitor-arg** If the workflow management service accepts extra arguments, provide. them in key value pairs with –wms-monitor-arg. For example, to run an existing workflow using a wms monitor, you can provide the pair id=12345 and the arguments will be provided to the endpoint to first interact with the workflow

**--scheduler-ilp-solver** Possible choices: PULP_CBC_CMD

Specifies solver to be utilized when selecting ilp-scheduler.

Default: "COIN_CMD"

**--scheduler-solver-path** Set the PATH to search for scheduler solver binaries (internal use only).

**--conda-base-path** Path of conda base installation (home of conda, mamba, activate) (internal use only).

**--no-subworkflows, --nosw** Do not evaluate or execute subworkflows.

Default: False

##### 4.2.2 GROUPING

**--groups** Assign rules to groups (this overwrites any group definitions from the workflow).

**--group-components** Set the number of connected components a group is allowed to span. By default, this is 1, but this flag allows to extend this. This can be used to run e.g. 3 jobs of the same rule in the same group, although they are not connected. It can be helpful for putting together many small jobs or benefitting of shared memory setups.

##### 4.2.3 REPORTS

**--report** Create an HTML report with results and statistics. This can be either a .html file or a .zip file. In the former case, all results are embedded into the .html (this only works for small data). In the latter case, results are stored along with a file report.html in the zip archive. If no filename is given, an embedded report.html is the default.

**--report-stylesheet** Custom stylesheet to use for report. In particular, this can be used for branding the report with e.g. a custom logo, see docs.

##### 4.2.4 NOTEBOOKS

**--draft-notebook** Draft a skeleton notebook for the rule used to generate the given target file. This notebook can then be opened in a jupyter server, exeucted and implemented until ready. After saving, it will automatically be reused in non-interactive mode by Snakemake for subsequent jobs.

**--edit-notebook** Interactively edit the notebook associated with the rule used to generate the given target file. This will start a local jupyter notebook server. Any changes to the notebook should be saved, and the server has to be stopped by closing the notebook and hitting the 'Quit' button on the jupyter dashboard. Afterwards, the updated notebook will be automatically stored in the path defined in the rule. If the notebook is not yet present, this will create an empty draft.

**--notebook-listen** The IP address and PORT the notebook server used for editing the notebook (–edit-notebook) will listen on.

Default: "localhost:8888"

##### 4.2.5 UTILITIES

**--lint** Possible choices: text, json

Perform linting on the given workflow. This will print snakemake specific suggestions to improve code quality (work in progress, more lints to be added in the future). If no argument is provided, plain text output is used.

**--generate-unit-tests** Automatically generate unit tests for each workflow rule. This assumes that all input files of each job are already present. Rules without a job with present input files will be skipped (a warning will be issued). For each rule, one test case will be created in the specified test folder (.tests/unit by default). After successfull execution, tests can be run with 'pytest TESTPATH'.

**--containerize** Print a Dockerfile that provides an execution environment for the workflow, including all conda environments.

Default: False

**--export-cwl** Compile workflow to CWL and store it in given FILE.

**--list, -l** Show available rules in given Snakefile.

Default: False

**--list-target-rules, --lt** Show available target rules in given Snakefile.

Default: False

**--dag** Do not execute anything and print the directed acyclic graph of jobs in the dot language. Recommended use on Unix systems: snakemake –dag | dot | displayNote print statements in your Snakefile may interfere with visualization.

Default: False

**--rulegraph** Do not execute anything and print the dependency graph of rules in the dot language. This will be less crowded than above DAG of jobs, but also show less information. Note that each rule is displayed once, hence the displayed graph will be cyclic if a rule appears in several steps of the workflow. Use this if above option leads to a DAG that is too large. Recommended use on Unix systems: snakemake –rulegraph | dot | displayNote print statements in your Snakefile may interfere with visualization.

Default: False

**--filegraph** Do not execute anything and print the dependency graph of rules with their input and output files in the dot language. This is an intermediate solution between above DAG of jobs and the rule graph. Note that each rule is displayed once, hence the displayed graph will be cyclic if a rule appears in several steps of the workflow. Use this if above option leads to a DAG that is too large. Recommended use on Unix systems: snakemake –filegraph | dot | displayNote print statements in your Snakefile may interfere with visualization.

Default: False

**--d3dag** Print the DAG in D3.js compatible JSON format.

Default: False

**--summary, -S** Print a summary of all files created by the workflow. The has the following columns: filename, modification time, rule version, status, plan. Thereby rule version contains the versionthe file was created with (see the version keyword of rules), and status denotes whether the file is missing, its input files are newer or if version or implementation of the rule changed since file creation. Finally the last column denotes whether the file will be updated or created during the next workflow execution.

Default: False

**--detailed-summary, -D** Print a summary of all files created by the workflow. The has the following columns: filename, modification time, rule version, input file(s), shell command, status, plan. Thereby rule version contains the version the file was created with (see the version keyword of rules), and status denotes whether the file is missing, its input files are newer or if version or implementation of the rule changed since file creation. The input file and shell command columns are self explanatory. Finally the last column denotes whether the file will be updated or created during the next workflow execution.

Default: False

**--archive** Archive the workflow into the given tar archive FILE. The archive will be created such that the workflow can be re-executed on a vanilla system. The function needs conda and git to be installed. It will archive every file that is under git version control. Note that it is best practice to have the Snakefile, config files, and scripts under version control. Hence, they will be included in the archive. Further, it will add input files that are not generated by by the workflow itself and conda environments. Note that symlinks are dereferenced. Supported formats are .tar, .tar.gz, .tar.bz2 and .tar.xz.

**--cleanup-metadata, --cm** Cleanup the metadata of given files. That means that snakemake removes any tracked version info, and any marks that files are incomplete.

**--cleanup-shadow** Cleanup old shadow directories which have not been deleted due to failures or power loss.

Default: False

**--skip-script-cleanup** Don't delete wrapper scripts used for execution

Default: False

**--unlock** Remove a lock on the working directory.

Default: False

**--list-version-changes, --lv** List all output files that have been created with a different version (as determined by the version keyword).

Default: False

**--list-code-changes, --lc** List all output files for which the rule body (run or shell) have changed in the Snakefile.

Default: False

**--list-input-changes, --li** List all output files for which the defined input files have changed in the Snakefile (e.g. new input files were added in the rule definition or files were renamed). For listing input file modification in the filesystem, use –summary.

Default: False

**--list-params-changes, --lp** List all output files for which the defined params have changed in the Snakefile.

Default: False

**--list-untracked, --lu** List all files in the working directory that are not used in the workflow. This can be used e.g. for identifying leftover files. Hidden files and directories are ignored.

Default: False

**--delete-all-output** Remove all files generated by the workflow. Use together with –dry-run to list files without actually deleting anything. Note that this will not recurse into subworkflows. Write-protected files are not removed. Nevertheless, use with care!

Default: False

**--delete-temp-output** Remove all temporary files generated by the workflow. Use together with –dry-run to list files without actually deleting anything. Note that this will not recurse into subworkflows.

Default: False

**--bash-completion** Output code to register bash completion for snakemake. Put the following in your .bashrc (including the accents): *snakemake –bash-completion* or issue it in an open terminal session.

Default: False

**--keep-incomplete** Do not remove incomplete output files by failed jobs.

Default: False

**--drop-metadata** Drop metadata file tracking information after job finishes. Provenance-information based reports (e.g. –report and the –list_x_changes functions) will be empty or incomplete.

Default: False

**--version, -v** show program’s version number and exit

##### 4.2.6 OUTPUT

**--reason, -r** Print the reason for each executed rule (deprecated, always true now).

Default: False

**--gui** Serve an HTML based user interface to the given network and port e.g. 168.129.10.15:8000. By default Snakemake is only available in the local network (default port: 8000). To make Snakemake listen to all ip addresses add the special host address 0.0.0.0 to the url (0.0.0.0:8000). This is important if Snakemake is used in a virtualised environment like Docker. If possible, a browser window is opened.

**--printshellcmds, -p** Print out the shell commands that will be executed.

Default: False

**--debug-dag** Print candidate and selected jobs (including their wildcards) while inferring DAG. This can help to debug unexpected DAG topology or errors.

Default: False

**--stats** Write stats about Snakefile execution in JSON format to the given file.

**--nocolor** Do not use a colored output.

Default: False

**--quiet, -q** Possible choices: progress, rules, all

Do not output certain information. If used without arguments, do not output any progress or rule information. Defining 'all' results in no information being printed at all.

**--print-compilation** Print the python representation of the workflow.

Default: False

**--verbose** Print debugging output.

Default: False

##### 4.2.7 BEHAVIOR

**--force-use-threads** Force threads rather than processes. Helpful if shared memory (/dev/shm) is full or unavailable.

Default: False

**--allow-ambiguity, -a** Don't check for ambiguous rules and simply use the first if several can produce the same file. This allows the user to prioritize rules by their order in the snakefile.

Default: False

**--nolock** Do not lock the working directory

Default: False

**--ignore-incomplete, --ii** Do not check for incomplete output files.

Default: False

**--max-inventory-time** Spend at most SECONDS seconds to create a file inventory for the working directory. The inventory vastly speeds up file modification and existence checks when computing which jobs need to be executed. However, creating the inventory itself can be slow, e.g. on network file systems. Hence, we do not spend more than a given amount of time and fall back to individual checks for the rest.

Default: 20

**--latency-wait, --output-wait, -w** Wait given seconds if an output file of a job is not present after the job finished. This helps if your filesystem suffers from latency (default 5).

Default: 5

**--wait-for-files** Wait –latency-wait seconds for these files to be present before executing the workflow. This option is used internally to handle filesystem latency in cluster environments.

**--wait-for-files-file** Same behaviour as –wait-for-files, but file list is stored in file instead of being passed on the commandline. This is useful when the list of files is too long to be passed on the commandline.

**--notemp, --nt** Ignore temp() declarations. This is useful when running only a part of the workflow, since temp() would lead to deletion of probably needed files by other parts of the workflow.

Default: False

**--all-temp** Mark all output files as temp files. This can be useful for CI testing, in order to save space.

Default: False

**--keep-remote** Keep local copies of remote input files.

Default: False

**--keep-target-files** Do not adjust the paths of given target files relative to the working directory.

Default: False

**--allowed-rules** Only consider given rules. If omitted, all rules in Snakefile are used. Note that this is intended primarily for internal use and may lead to unexpected results otherwise.

**--local-groupid** Name for local groupid, meant for internal use only.

Default: "local"

**--max-jobs-per-second** Maximal number of cluster/drmaa jobs per second, default is 10, fractions allowed.

Default: 10

**--max-status-checks-per-second** Maximal number of job status checks per second, default is 10, fractions allowed.

Default: 10

**-T, --retries, --restart-times** Number of times to restart failing jobs (defaults to 0).

Default: 0

**--attempt** Internal use only: define the initial value of the attempt parameter (default: 1).

Default: 1

**--wrapper-prefix** Prefix for URL created from wrapper directive (default: https://github.com/snakemake/snakemake-wrappers/raw/). Set this to a different URL to use your fork or a local clone of the repository, e.g., use a git URL like 'git+file://path/to/your/local/clone@'.

Default: "https://github.com/snakemake/snakemake-wrappers/raw/"

**--default-remote-provider** Possible choices: S3, GS, FTP, SFTP, S3Mocked, gfal, gridftp, iRODS, AzBlob, XRootD

Specify default remote provider to be used for all input and output files that don't yet specify one.

**--default-remote-prefix** Specify prefix for default remote provider. E.g. a bucket name.

Default: ""

**--no-shared-fs** Do not assume that jobs share a common file system. When this flag is activated, Snakemake will assume that the filesystem on a cluster node is not shared with other nodes. For example, this will lead to downloading remote files on each cluster node separately. Further, it won't take special measures to deal with filesystem latency issues. This option will in most cases only make sense in combination with –default-remote-provider. Further, when using –cluster you will have to also provide –cluster-status. Only activate this if you know what you are doing.

Default: False

**--greediness** Set the greediness of scheduling. This value between 0 and 1 determines how careful jobs are selected for execution. The default value (1.0) provides the best speed and still acceptable scheduling quality.

**--no-hooks** Do not invoke onstart, onsuccess or onerror hooks after execution.

Default: False

**--overwrite-shellcmd** Provide a shell command that shall be executed instead of those given in the workflow. This is for debugging purposes only.

**--debug** Allow to debug rules with e.g. PDB. This flag allows to set breakpoints in run blocks.

Default: False

**--runtime-profile** Profile Snakemake and write the output to FILE. This requires yappi to be installed.

**--mode** Possible choices: 0, 1, 2

Set execution mode of Snakemake (internal use only).

Default: 0

**--show-failed-logs** Automatically display logs of failed jobs.

Default: False

**--log-handler-script** Provide a custom script containing a function 'def log_handler(msg):'. Snakemake will call this function for every logging output (given as a dictionary msg)allowing to e.g. send notifications in the form of e.g. slack messages or emails.

**--log-service** Possible choices: none, slack, wms

Set a specific messaging service for logging output.Snakemake will notify the service on errors and completed execution.Currently slack and workflow management system (wms) are supported.

##### 4.2.8 CLUSTER

**--cluster** Execute snakemake rules with the given submit command, e.g. qsub. Snakemake compiles jobs into scripts that are submitted to the cluster with the given command, once all input files for a particular job are present. The submit command can be decorated to make it aware of certain job properties (name, rulename, input, output, params, wildcards, log, threads and dependencies (see the argument below)), e.g.: $ snakemake –cluster 'qsub -pe threaded {threads}'.

**--cluster-sync** cluster submission command will block, returning the remote exitstatus upon remote termination (for example, this should be usedif the cluster command is 'qsub -sync y' (SGE)

**--drmaa** Execute snakemake on a cluster accessed via DRMAA, Snakemake compiles jobs into scripts that are submitted to the cluster with the given command, once all input files for a particular job are present. ARGS can be used to specify options of the underlying cluster system, thereby using the job properties name, rulename, input, output, params, wildcards, log, threads and dependencies, e.g.: –drmaa ' -pe threaded {threads}'. Note that ARGS must be given in quotes and with a leading whitespace.

**--cluster-config, -u** A JSON or YAML file that defines the wildcards used in 'cluster'for specific rules, instead of having them specified in the Snakefile. For example, for rule 'job' you may define: { 'job' : { 'time' : '24:00:00' } } to specify the time for rule 'job'. You can specify more than one file. The configuration files are merged with later values overriding earlier ones. This option is deprecated in favor of using –profile, see docs.

Default: []

**--immediate-submit, --is** Immediately submit all jobs to the cluster instead of waiting for present input files. This will fail, unless you make the cluster aware of job dependencies, e.g. via: $ snakemake –cluster 'sbatch –dependency {dependencies}. Assuming that your submit script (here sbatch) outputs the generated job id to the first stdout line, {dependencies} will be filled with space separated job ids this job depends on. Does not work for workflows that contain checkpoint rules.

Default: False

**--jobscript, --js** Provide a custom job script for submission to the cluster. The default script resides as 'jobscript.sh' in the installation directory.

**--jobname, --jn** Provide a custom name for the jobscript that is submitted to the cluster (see –cluster). NAME is "snakejob.{name}.{jobid}.sh" per default. The wildcard {jobid} has to be present in the name.

Default: "snakejob.{name}.{jobid}.sh"

**--cluster-status** Status command for cluster execution. This is only considered in combination with the –cluster flag. If provided, Snakemake will use the status command to determine if a job has finished successfully or failed. For this it is necessary that the submit command provided to –cluster returns the cluster job id. Then, the status command will be invoked with the job id. Snakemake expects it to return 'success' if the job was successfull, 'failed' if the job failed and 'running' if the job still runs.

**--cluster-cancel** Specify a command that allows to stop currently running jobs. The command will be passed a single argument, the job id.

**--cluster-cancel-nargs** Specify maximal number of job ids to pass to –cluster-cancel command, defaults to 1000.

Default: 1000

**--cluster-sidecar** Optional command to start a sidecar process during cluster execution. Only active when –cluster is given as well.

**--drmaa-log-dir** Specify a directory in which stdout and stderr files of DRMAA jobs will be written. The value may be given as a relative path, in which case Snakemake will use the current invocation directory as the origin. If given, this will override any given '-o' and/or '-e' native specification. If not given, all DRMAA stdout and stderr files are written to the current working directory.

##### 4.2.9 KUBERNETES

**--kubernetes** Execute workflow in a kubernetes cluster (in the cloud). NAMESPACE is the namespace you want to use for your job (if nothing specified: 'default'). Usually, this requires –default-remote-provider and –default-remote-prefix to be set to a S3 or GS bucket where your . data shall be stored. It is further advisable to activate conda integration via –use-conda.

**--container-image** Docker image to use, e.g., when submitting jobs to kubernetes Defaults to 'https://hub.docker.com/r/snakemake/snakemake', tagged with the same version as the currently running Snakemake instance. Note that overwriting this value is up to your responsibility. Any used image has to contain a working snakemake installation that is compatible with (or ideally the same as) the currently running version.

##### 4.2.10 TIBANNA

**--tibanna** Execute workflow on AWS cloud using Tibanna. This requires –default-remote-prefix to be set to S3 bucket name and prefix (e.g. 'bucketname/subdirectory') where input is already stored and output will be sent to. Using –tibanna implies –default-resources is set as default. Optionally, use –precommand to specify any preparation command to run before snakemake command on the cloud (inside snakemake container on Tibanna VM). Also, –use-conda, –use-singularity, –config, –configfile are supported and will be carried over.

Default: False

**--tibanna-sfn** Name of Tibanna Unicorn step function (e.g. tibanna_unicorn_monty).This works as serverless scheduler/resource allocator and must be deployed first using tibanna cli. (e.g. tibanna deploy_unicorn –usergroup=monty –buckets=bucketname)

**--precommand** Any command to execute before snakemake command on AWS cloud such as wget, git clone, unzip, etc. This is used with –tibanna.Do not include input/output download/upload commands - file transfer between S3 bucket and the run environment (container) is automatically handled by Tibanna.

**--tibanna-config** Additional tibanna config e.g. –tibanna-config spot_instance=true subnet=<subnet_id> security group=<security_group_id>

##### 4.2.11 GOOGLE_LIFE_SCIENCE

**--google-lifesciences** Execute workflow on Google Cloud cloud using the Google Life. Science API. This requires default application credentials (json) to be created and export to the environment to use Google Cloud Storage, Compute Engine, and Life Sciences. The credential file should be exported as GOOGLE_APPLICATION_CREDENTIALS for snakemake to discover. Also, –use-conda, –use-singularity, –config, –configfile are supported and will be carried over.

Default: False

**--google-lifesciences-regions** Specify one or more valid instance regions (defaults to US)

Default: ['us-east1', 'us-west1', 'us-central1']

**--google-lifesciences-location** The Life Sciences API service used to schedule the jobs. E.g., us-centra1 (Iowa) and europe-west2 (London) Watch the terminal output to see all options found to be available. If not specified, defaults to the first found with a matching prefix from regions specified with –google-lifesciences-regions.

**--google-lifesciences-keep-cache** Cache workflows in your Google Cloud Storage Bucket specified by –default-remote-prefix/{source}/{cache}. Each workflow working directory is compressed to a .tar.gz, named by the hash of the contents, and kept in Google Cloud Storage. By default, the caches are deleted at the shutdown step of the workflow.

Default: False

##### 4.2.12 TES

**--tes** Send workflow tasks to GA4GH TES server specified by url.

##### 4.2.13 CONDA

**--use-conda** If defined in the rule, run job in a conda environment. If this flag is not set, the conda directive is ignored.

Default: False

**--conda-not-block-search-path-envvars** Do not block environment variables that modify the search path (R_LIBS, PYTHONPATH, PERL5LIB, PERLLIB) when using conda environments.

Default: False

**--list-conda-envs** List all conda environments and their location on disk.

Default: False

**--conda-prefix** Specify a directory in which the 'conda' and 'conda-archive' directories are created. These are used to store conda environments and their archives, respectively. If not supplied, the value is set to the '.snakemake' directory relative to the invocation directory. If supplied, the *–use-conda* flag must also be set. The value may be given as a relative path, which will be extrapolated to the invocation directory, or as an absolute path. The value can also be provided via the environment variable $SNAKEMAKE_CONDA_PREFIX.

**--conda-cleanup-envs** Cleanup unused conda environments.

Default: False

**--conda-cleanup-pkgs** Possible choices: tarballs, cache

Cleanup conda packages after creating environments. In case of 'tarballs' mode, will clean up all downloaded package tarballs. In case of 'cache' mode, will additionally clean up unused package caches. If mode is omitted, will default to only cleaning up the tarballs.

**--conda-create-envs-only** If specified, only creates the job-specific conda environments then exits. The *–use-conda* flag must also be set.

Default: False

**--conda-frontend** Possible choices: conda, mamba

Choose the conda frontend for installing environments. Mamba is much faster and highly recommended.

Default: "mamba"

##### 4.2.14 SINGULARITY

**--use-singularity** If defined in the rule, run job within a singularity container. If this flag is not set, the singularity directive is ignored.

Default: False

**--singularity-prefix** Specify a directory in which singularity images will be stored.If not supplied, the value is set to the '.snakemake' directory relative to the invocation directory. If supplied, the *–use-singularity* flag must also be set. The value may be given as a relative path, which will be extrapolated to the invocation directory, or as an absolute path.

**--singularity-args** Pass additional args to singularity.

Default: ""

##### 4.2.15 ENVIRONMENT MODULES

**--use-envmodules** If defined in the rule, run job within the given environment modules, loaded in the given order. This can be combined with –use-conda and –use-singularity, which will then be only used as a fallback for rules which don't define environment modules.

Default: False

CHAPTER

FIVE

### RUNNING HIPPUNFOLD ON YOUR DATA

This section goes over the command-line options you will find most useful when running HippUnfold on your dataset, along with describing some of the issues you might face.

Note: Please first refer to the simple example in the Installation section, which goes over running HippUnfold on a test dataset, and the essential required options.

#### 5.1 Selecting the modality to use

The `--modality` option must be chosen when running HippUnfold, and it affects what U-net model will be used, and how the pre-processing will be performed on the images.

If you have sub-millimetric, isotropic, whole-brain T1w data, the `--modality T1w` option is recommended.

If you T2w data, you can use the `--modality T2w` option, however, you may need to also use the T1w data for template registration (`--t1-reg-template`), especially if you have a limited FOV. This is typically most robust as long as a full brain FOV T1w image is available. If this registration is still failing then it may be improved with the `--rigid-reg-template` flag.

For protocols employing high-resolution, b-value 500, hippocampal diffusion-weighted imaging, the `--modality hippb500` option can be used, and does not require registration to a template (providing your acquisition is axial and oblique to the hippocampus).

Specifying a manual segmentation (eg. `--modality segT1w`) expects to additionally find an input file with the suffix `_dseg` which should contain labels following the protocol outlined here. More details are provided on using manual segmentations on the following page.

#### 5.2 Selecting and excluding subjects to process

By default, hippunfold will run on **all** the subjects in a dataset. If you want to run only on a subset of subjects, you can use the `--participant_label` flag, e.g. adding:

```
--participant-label 001
```

would run only on `sub-001`. You can add additional subjects by listing additional arguments to this option, e.g.:

```
--participant-label 001 002
```

runs for `sub-001` and `sub-001`.

Also, if you want to exclude a subject, you can use the `--exclude-participant-label` option.

#### 5.3 Known limitations for BIDS parsing

HippUnfold uses snakebids, which makes use of pybids to parse a BIDS-compliant dataset. However, because of the way Snakebids and Snakemake operate, one limitation is that the input files in your BIDS dataset need to be consistent in terms of what optional BIDS entities exist in them. We can use the acquisition (acq) entity as an example. HippUnfold should have no problem parsing the following dataset:

```
PATH_TO_BIDS_DIR/
└── dataset_description.json
└── sub-001/
    └── anat/
        ├── sub-001_acq-mprage_T1w.nii.gz
└── sub-002/
    └── anat/
        ├── sub-002_acq-spgr_T1w.nii.gz
...
```

as the path (with wildcards) will be interpreted as sub-{subject}_acq-{acq}_T1w.nii.gz.

However, the following dataset will raise an error:

```
PATH_TO_BIDS_DIR/
└── dataset_description.json
└── sub-001/
    └── anat/
        ├── sub-001_acq-mprage_T1w.nii.gz
└── sub-002/
    └── anat/
        ├── sub-002_T1w.nii.gz
...
```

because two distinct paths (with wildcards) would be found for T1w images:

```
sub-{subject}_acq-{acq}_T1w.nii.gz
```

and

```
sub-{subject}_T1w.nii.gz
```

Similarly, you could not have some subjects with the ses identifier, and some subjects without it.

There will soon be added functionality in snakebids to filter out extra files, but for now, if your dataset has these issues you will need to rename or remove extraneous files.

More examples of possible BIDS-compliant datasets can be found in hippunfold/test_data/.

#### 5.4 Parsing Non-BIDS datasets with custom paths

Custom paths can be used to parse input datasets if the data are not in BIDS format, but still are uniquely identified by subject (or subject+session) identifiers. For example:

```
PATH_TO_nonBIDS_DIR/
└── s_001_T1w.nii.gz
└── s_001_T2SPACE.nii.gz
└── s_001_TSE.nii.gz
└── s_002_T1w.nii.gz
└── s_002_T2SPACE.nii.gz
└── s_002_TSE.nii.gz
...
```

This directory doesn't separate subjects into different folders or contain an `anat/` folder for structural images. However, we can still specify what subjects and images to use with subject `wildcards`. This is done by using the `--path-{modality}` options to specify the absolute location of the `nii.gz` files. Note that here, T2SPACE and TSE are both T2-weighted acquisitions, and can be captured by using the `--path-T2w` flag to specify exactly which of these file(s) to use as inputs. For example, the following command:

```
hippunfold - PATH_TO_OUTPUT_DIR participant \
--modality T2w \
--t1-reg-template \
--path_T1w PATH_TO_nonBIDS_DIR/s_{subject}_T1w.nii.gz \
--path_T2w PATH_TO_nonBIDS_DIR/s_{subject}_T2SPACE.nii.gz
```

will search for any files following the naming scheme and fill in `{subject}` IDs for any files it finds, using the T1w and T2SPACE images for `T1w` and `T2w` inputs

##### 5.4.1 Prerequisities for using custom path parsing:

Not all non-BIDS datasets can be parsed, and may still need some reformatting or renaming.

Specifically:

- The subject (or subject/session) wildcard(s) can only contain letters or numbers, e.g. they cannot include underscores, hyphens, or spaces.
- The subject (or subject/session) wildcard(s) must be the only unique identifiers in the filenames.

For example, this datasets would be **ineligible**:

```
PATH_TO_nonBIDS_DIR/
└── s_2019-05-29_001_T1w.nii.gz
└── s_2019-05-29_001_T2SPACE.nii.gz
└── s_2019-05-29_001_TSE.nii.gz
└── s_2018-02-24_002_T1w.nii.gz
└── s_2018-02-24_002_T2SPACE.nii.gz
└── s_2018-02-24_002_TSE.nii.gz
...
```

You would need to rename/symlink your images to remove the additional unique date identifiers, or integrate it into thes subject wildcard, ensuring only letters and numbers appear in the wildcard, e.g.:

```
PATH_TO_nonBIDS_DIR/
└── s_20190529s001_T1w.nii.gz
└── s_20190529s001_T2SPACE.nii.gz
└── s_20190529s001_TSE.nii.gz
└── s_20180224s002_T1w.nii.gz
└── s_20180224s002_T2SPACE.nii.gz
└── s_20180224s002_TSE.nii.gz
...
```

### CHAPTER SIX

### SPECIALIZED SCANS

This tutorial will cover how HippUnfold can be applied to non-standard data including ex-vivo scans, super-high resolution data (eg. <0.3mm isotropic), non-MRI 3D imaging data, or scans where a corresponding whole-brain T1w image is not available.

We will show how the available flags can be adapted for these use-cases with several worked examples.

#### 6.1 Case 1: super high resolution

In this example, we have only a limited field of view covering the hippocampus, and the resolution and contrast do not closely match the training data of HippUnfold (0.3-1.0mm isotropic T1w, T2w, or DWI data). This could be ex-vivo MRI data, or it could even be 3D microscopy data as in our recent 3D BigBrain publication. Thus we don't expect HippUnfold's inbuilt UNet to be successful in segmenting hippocampal tissue before unfolding, and we do not want to downsample our data to accommodate HippUnfold's usual UNet and unfolding workflow in `space-corobl` (which consists of 0.3mm isotropic resampling cropped coronally-oblique to the hippocampus).

This will require manual segmentation of hippocampal grey matter, SRLM, and neighbouring structures, though in the future we hope to include models trained with higher resolution data (and contrasts more common in ex-vivo scanning). This should be done according to the protocol outlined here or, more recently, the video example here. This manual segmentation file should have the `_dseg` suffix.

Here is an example of what the input directory might look like:

```
exvivo/
└── sub-001/
    ├── sub-001_hemi-R_desc-hippo_T2w.nii.gz
    └── sub-001_hemi-R_desc-hippo_dseg.nii.gz
```

This can be unfolded with the command:

```
hippunfold . PATH_TO_OUTPUT_DIR participant --modality cropseg \
--path_cropseg exvivo/sub-{subject}/sub-{subject}_hemi-{hemi}_desc-hippo_dseg.nii.gz \
--hemi R --skip_inject_template_labels
```

Explanation: `--modality cropseg` informs HippUnfold that the input manual segmentation should not be resampled and UNet does not need to be run. Because of a limitation in bids parsing for the `hemi` entity, we need to use the generic path input, `--path_cropseg` in this case, making sure we use the `{subject}` and `{hemi}` wildcards in the filename. Output files will be named with `space-corobl` because HippUnfold is coded to effectively treat all files as already being in this space. We need the `--hemi R` to prevent HippUnfold looking for both hemispheres. Finally, because this segmentation was performed manually on very high resolution data, we can optionally consider skipping the template shape injection step with `--skip_inject_template_labels`. Template shape injection can fix minor errors in segmentation from UNet or from an imperfect manual rater, at the cost of smoothing out some details of the hippocampus due to the fact that it uses deformable registration with inherent smoothness contraints.

Note that because we are not resampling to the CITI168 template or using UNet, the T2w image in this example is effectively not being used at all. Instead, the provided manual segmentation makes up the basis for unfolding.

#### 6.2 Case 2: one ex-vivo hemisphere

In this example, we have a single hemisphere that was scanned ex-vivo at a nearly standard resolution and T2w contrast. Because the resolution and contrast are similar to the HippUnfold training data, we expect UNet will work and so we don't need to perform manual segemntation. However, due to gross deformations and the missing hemisphere, we don't expect this sample to register well to the standard CITI168 template. Thus, we will need to manually resample the image to the CITI168 template prior to running HippUnfold, focusing in particular on aligning the hippocampus. Once done, we may have a directory like this:

```
PATH_TO_EXVIVO_DIR/
└── sub-001/
    ├── sub-001_hemi-R_desc-exvivo_T2w.nii.gz
    ├── sub-001_hemi-R_affine-exvivo-to-CITI168_xfm.txt
    └── sub-001_hemi-R_desc-exvivo_space-CITI168_T2w.nii.gz
```

Note that only the last file is needed for unfolding:

```
hippunfold . PATH_TO_OUTPUT_DIR participant --output_spaces corobl --hemi R --no_reg_
↪template \
--path_T2w PATH_TO_EXVIVO_DIR/sub-001/sub-001_hemi-R_desc-exvivo_space-CITI168_T2w.nii.
↪gz \
--output_spaces corobl
```

Here we need to use `--path_T2w` to specify which input should be used, and `--no_reg_template` to specify that it is already in `space-CITI168`. In this case, we also specified `--output_spaces corobl`. This is not needed, but is useful when we are interested in only the hippocampus as `space-corobl` is higher resolution and cropped more nicely around the hippocampus then the original scan, making it a good space to perform subsequent analyses. Alternatively, outputs can be transformed back to the original space using the inverted transform `sub-001_hemi-R_affine-exvivo-to-CITI168_xfm.txt`.

This same usage could also be applied in a standard MRI case where no T1w image is available.

### PIPELINE DETAILS

This section describes the HippUnfold workflow, that is, the steps taken to produce the intermediate and final files. HippUnfold is a Snakemake workflow, and thus the workflow is a directed acyclic graph (DAG) that is automatically configured based on a set of rules.

#### 7.1 Overall workflow

Below is a example *simplified* visualization of the workflow DAG for the `--modality T1w` workflow. Each rounded rectangle in the DAG represents a *rule*, that is, some code or script that produces output file(s), and the arrows represent file inputs and outputs to these rules. It is *simplified* in that multiple instances of each rule are not shown, e.g. the `run_inference` rule runs on both left and right hemispheres (`hemi=L`, `hemi=R`), but only one `run_inference` box is shown here.

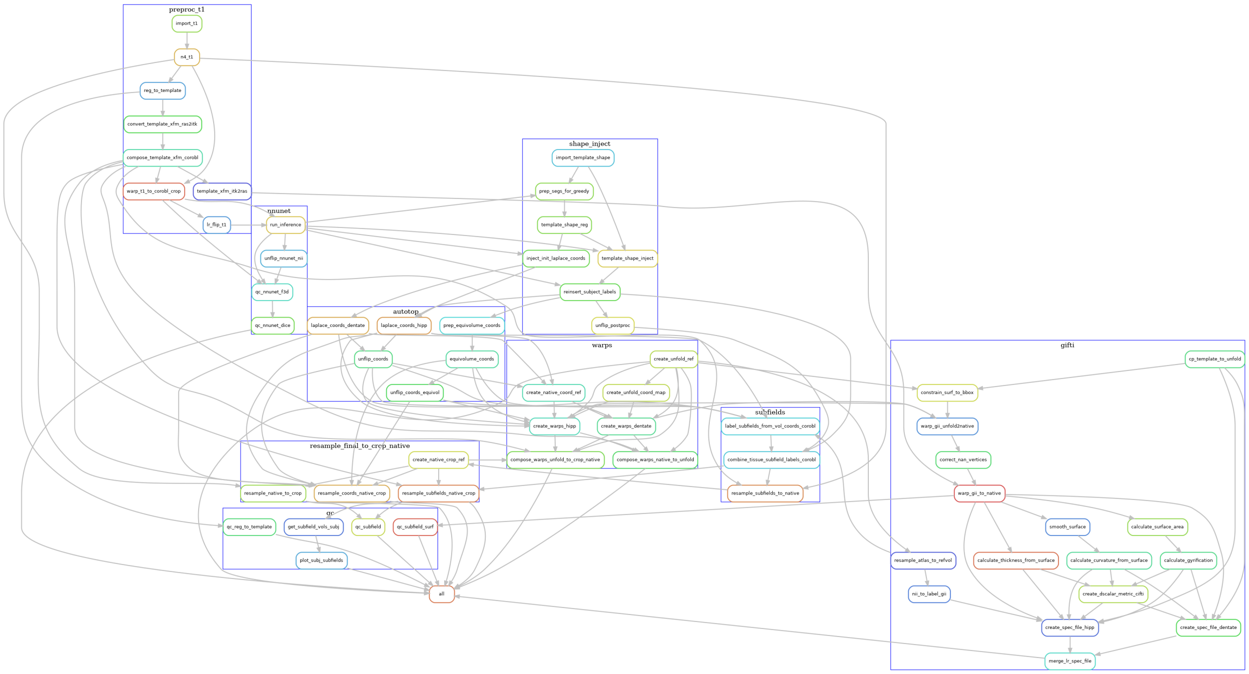

Although it may still look very complex (click on the image to enlarge), it is also organized into groups of rules, each representing the main phases of the workflow. Each grouped set of rules also exist in separate rule files, which can be found in the rules sub-folder
in the workflow source. For example, the preproc_t1 file contains the rules related to pre-processing the T1w images, and these are grouped together in the above diagram by a blue rectangle labelled `preproc_t1`.

The main phases of the workflow are described in the sections below, zooming in on the rules used in each blue rectangle, one at a time.

#### 7.2 Pre-processing

The pre-processing workflow for HippUnfold is generated based on the input data (e.g. whether there are multiple T2w images or a single T2w image), what modality is used (e.g. `--modality T1w` or `--modality T2w`), and what optional arguments are specified (e.g. `--t1-reg-template`).

##### 7.2.1 T1w pre-processing

T1w images are imported, intensity-corrected using N4, and linearly registered to the template image (default: CITI168 - an HCP T1w template). An existing transformation to align the images in a coronal oblique (`space-corobl`) orientation is concatenated, and this space is used to define the left and right hippocampus bounding boxes in 0.3mm isotropic space. The left hippocampus subvolume is left-right flipped at this stage too (subsequent steps in the `corobl` space operate on both the `hemi-R` and `hemi-Lflip` images).

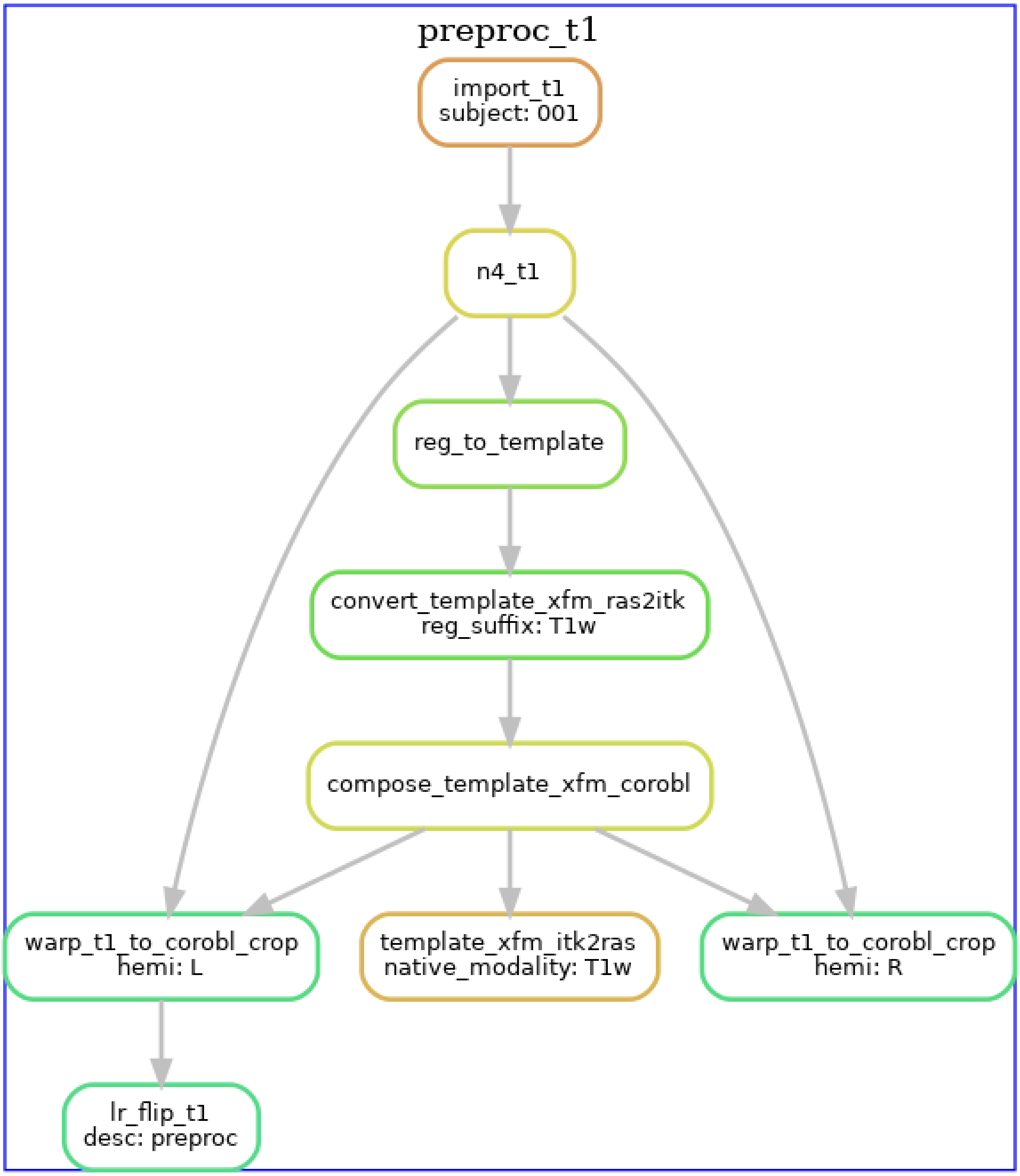
preproc_t1
import_t1
subject: 001
n4_t1
reg_to_template
convert_template_xfm_ras2itk
reg_suffix: T1w
compose_template_xfm_corobl
warp_t1_to_corobl_crop
hemi: L
template_xfm_itk2ras
native_modality: T1w
warp_t1_to_corobl_crop
hemi: R
lr_flip_t1
desc: preproc

##### 7.2.2 T2w pre-processing

T2w images are processed similarly, except the T2w version of the template is used. If multiple T2w images exist, these are motion-corrected and averaged prior to N4 correction. The diagram below shows the T2w pre- processing workflow for a dataset with three T2w runs.

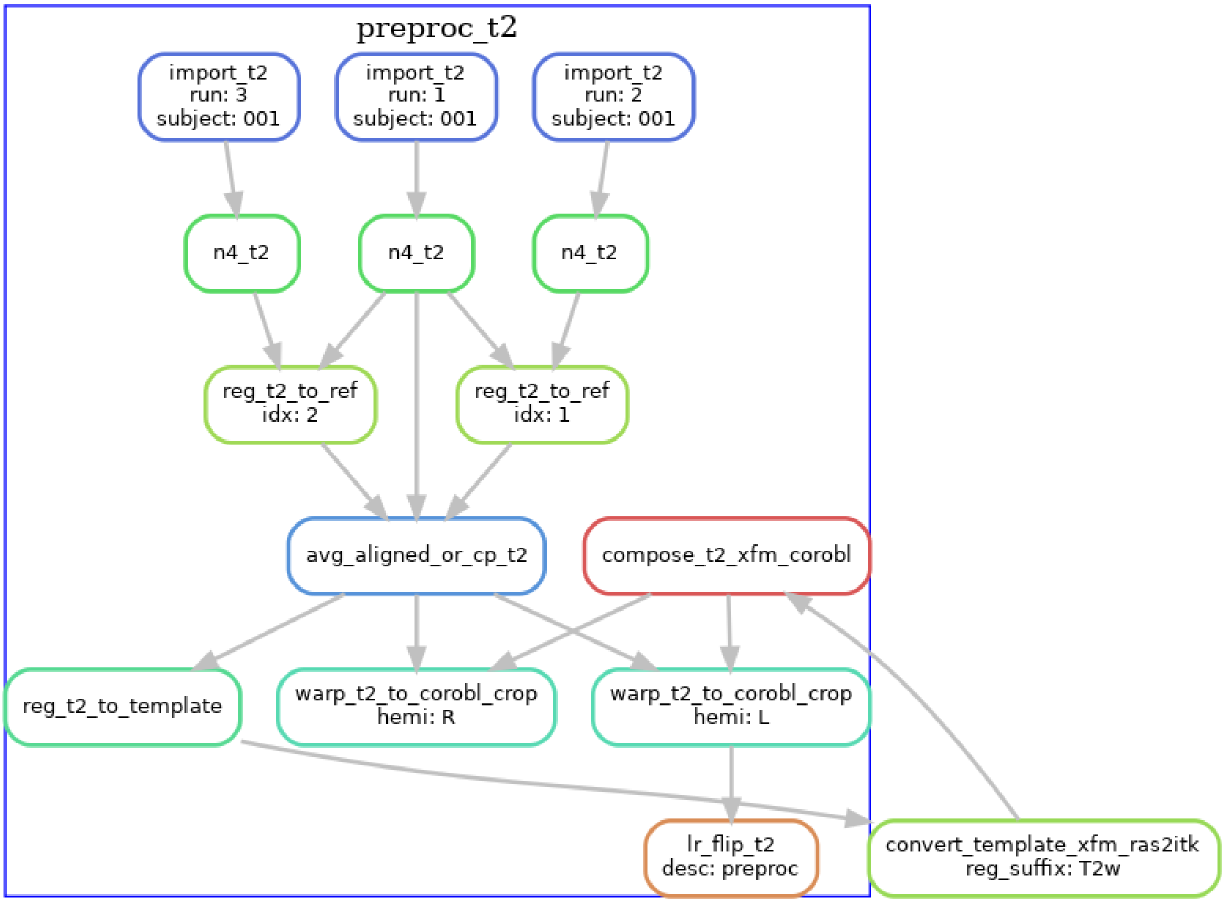

##### 7.2.3 T2w with T1w template registration

For T2w images where template registration is failing (e.g. because the T2w images have a limited FOV), the `--t1-reg-template` option can be used, and will perform template registration with the T1w images, along with a within-subject registration of the T2w to the T1w, concatenating all the transforms. This is shown in the diagrams below (with a single T2w image in this case):

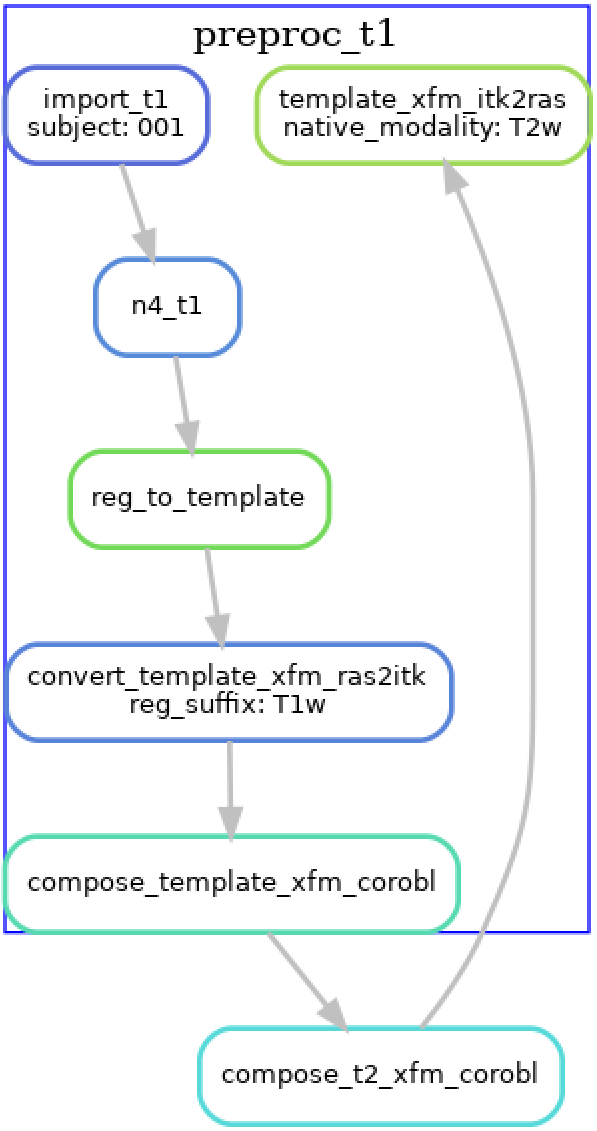
preproc_t1
import_t1
subject: 001
template_xfm_itk2ras
native_modality: T2w
n4_t1
reg_to_template
convert_template_xfm_ras2itk
reg_suffix: T1w
compose_template_xfm_corobl
compose_t2_xfm_corobl

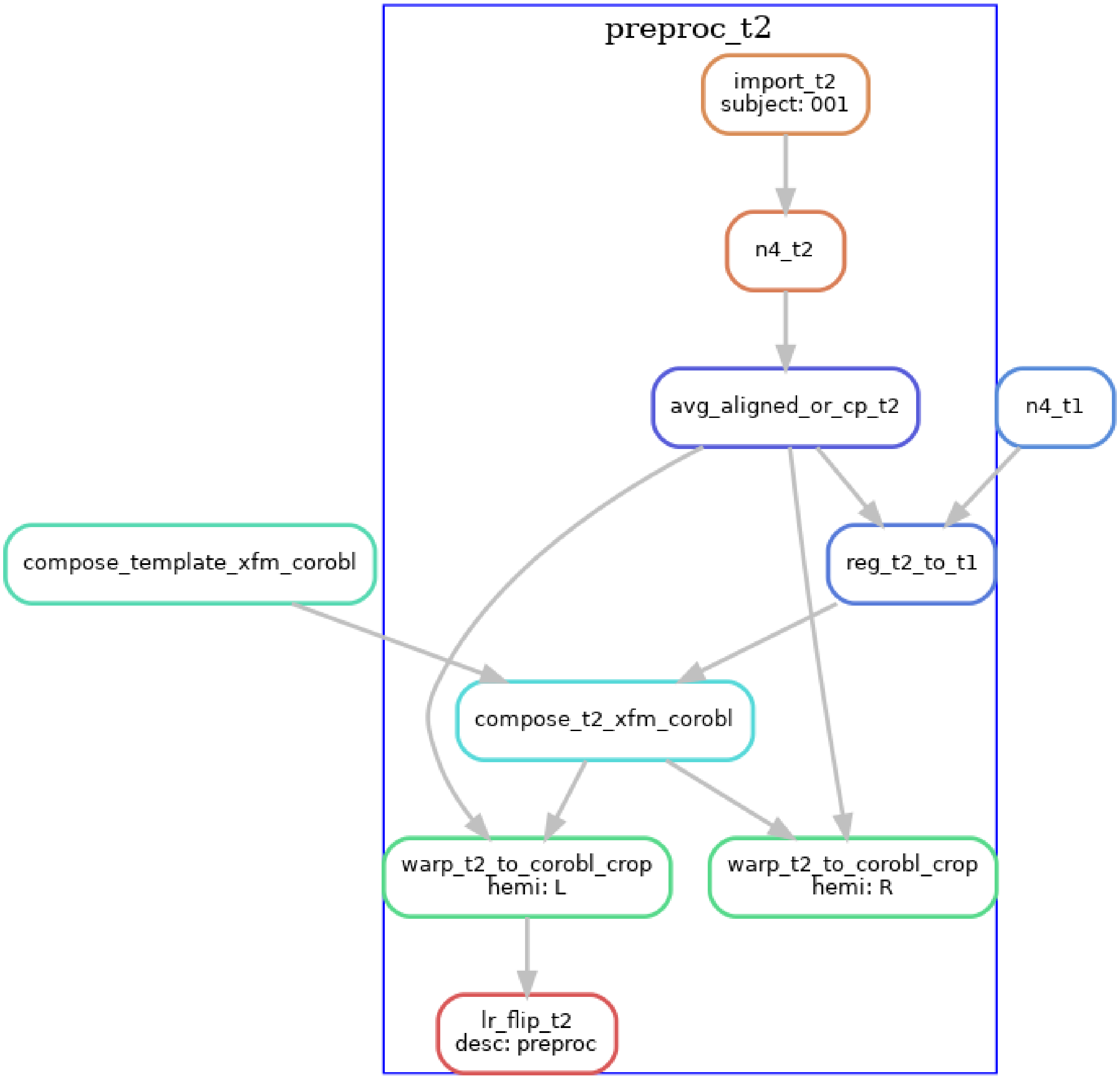

Note that these are not the only workflow configurations possible, several other variants exist by using the command-line flags. For example, if you have T1w and T2w images that are already pre-processed and co-registered (e.g. HCP processed data), then you should use the `--skip-preproc` and `--skip-coreg` options to skip N4 and T1w/T2w co-registration.

#### 7.3 U-net segmentation

The U-net segmentation is performed on the cropped, `space-corobl` images, producing tissue segmentations (gray matter, SRLM, and anatomical landmarks for unfolding). This step is done in a single rule, which runs inference on the image using the corresponding nnU-net model based on the `modality` chosen. This is done on the `R` and `Lflip` hippocampus images, and the `Lflip` is subsequently unflipped.

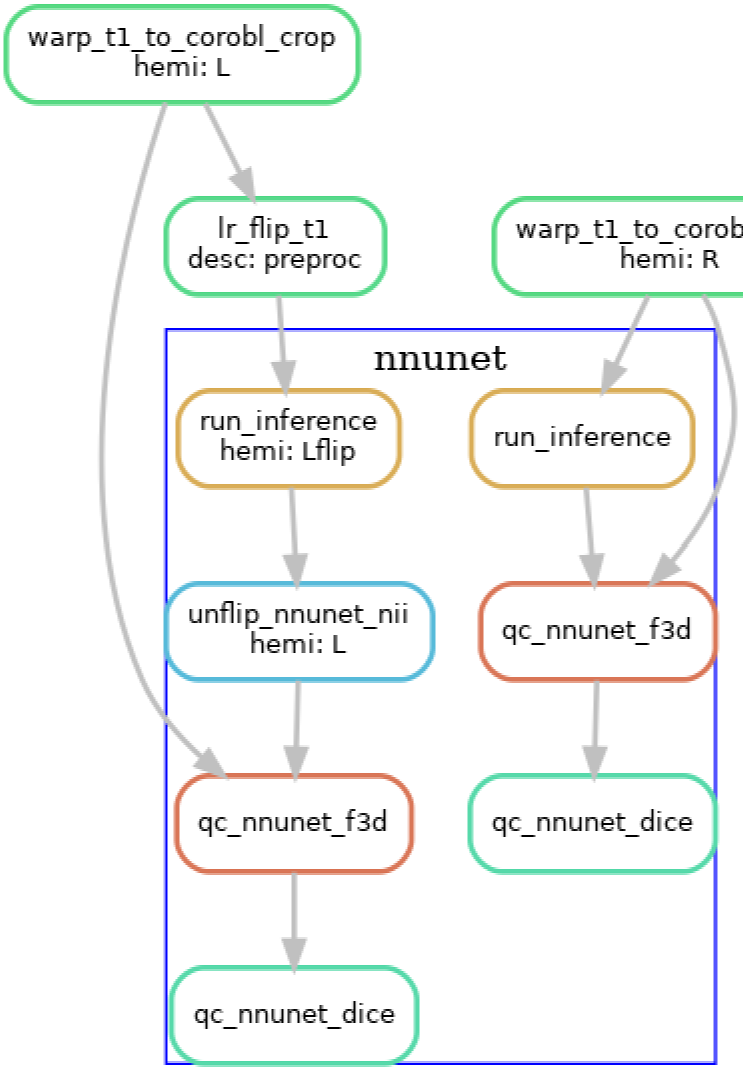

#### 7.4 Template-based shape injection

Since the nnU-net segmentation may possibly contain topological errors that can cause issues when the Laplace-based coordinates, we perform an additional registration-based correction step, a shape injection, where we perform non-linear registration of a template hippocampus segmentation and the U-net segmentation, to warp the template shape. Regularization from the registration ensures topology from the template is preserved while it is warped to match the subject hippocampus. In addition to the segmentation, we also propagate Laplace coordinates to serve as an initialization to the next step.

The following diagram shows the workflow, but simplified to contain one hemisphere (`--hemi R`), and excluding the dentate gyrus.

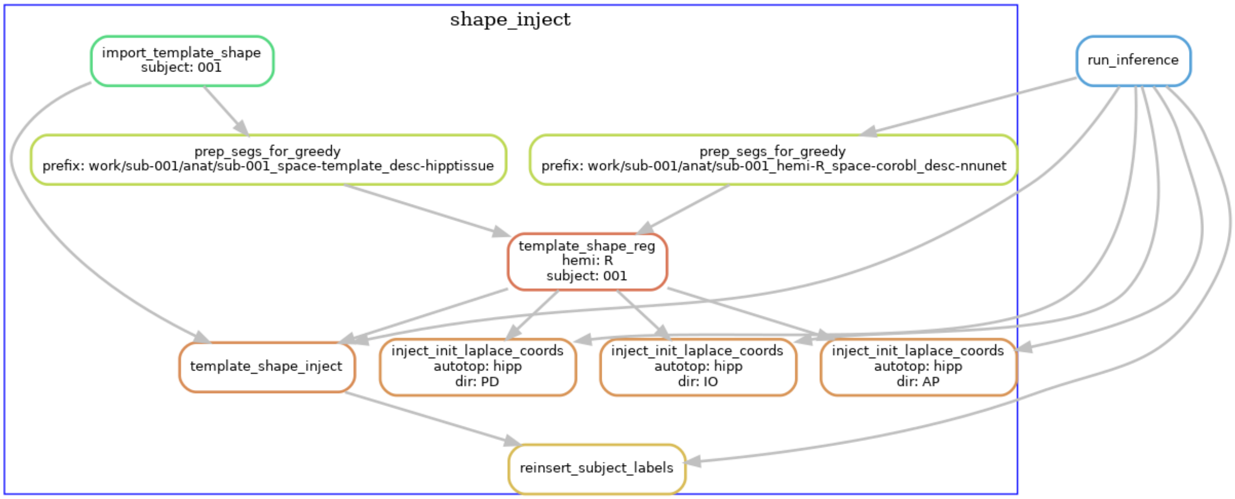

#### 7.5 Laplace & equivolume coordinates

The basis of the hippocampal unfolding is the definition of the Laplace coordinates. Here, Laplace's equation is solved on the domain of the gray matter, using the anatomical landmarks to define boundary conditions. This provides the intrinsic set of anatomical coordinates (AP, PD, IO) for unfolding the hippocampus. For the IO (laminar) coordinates we make use of the equivolume solution instead of Laplace.

The following diagram shows the workflow, but simplified to contain one hemisphere (`--hemi R`), and excluding the dentate gyrus.

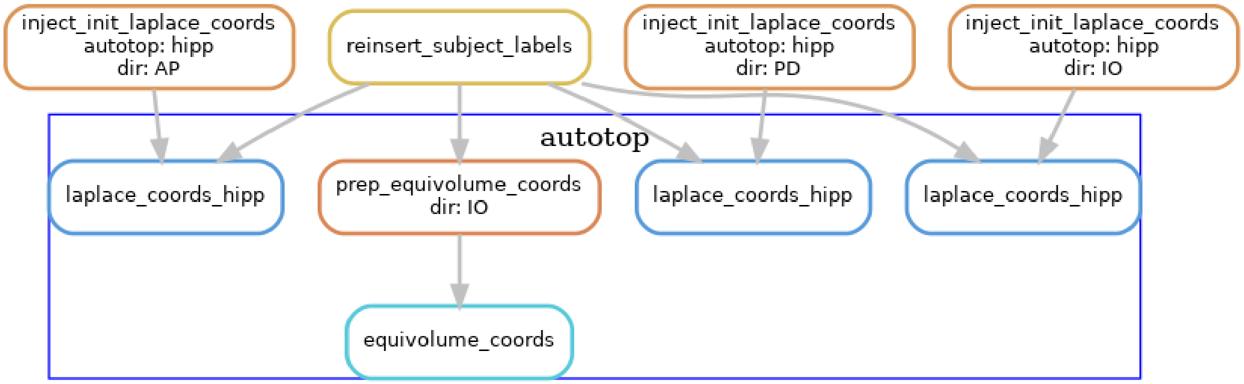

#### 7.6 Subfields processing

The volumetric subfield segmentation is derived from the coord images from the last step, along with the atlas that defines how the coordinates map to subfields.

The following diagram shows the workflow, but simplified to contain one hemisphere (`--hemi R`), and excluding the dentate gyrus.

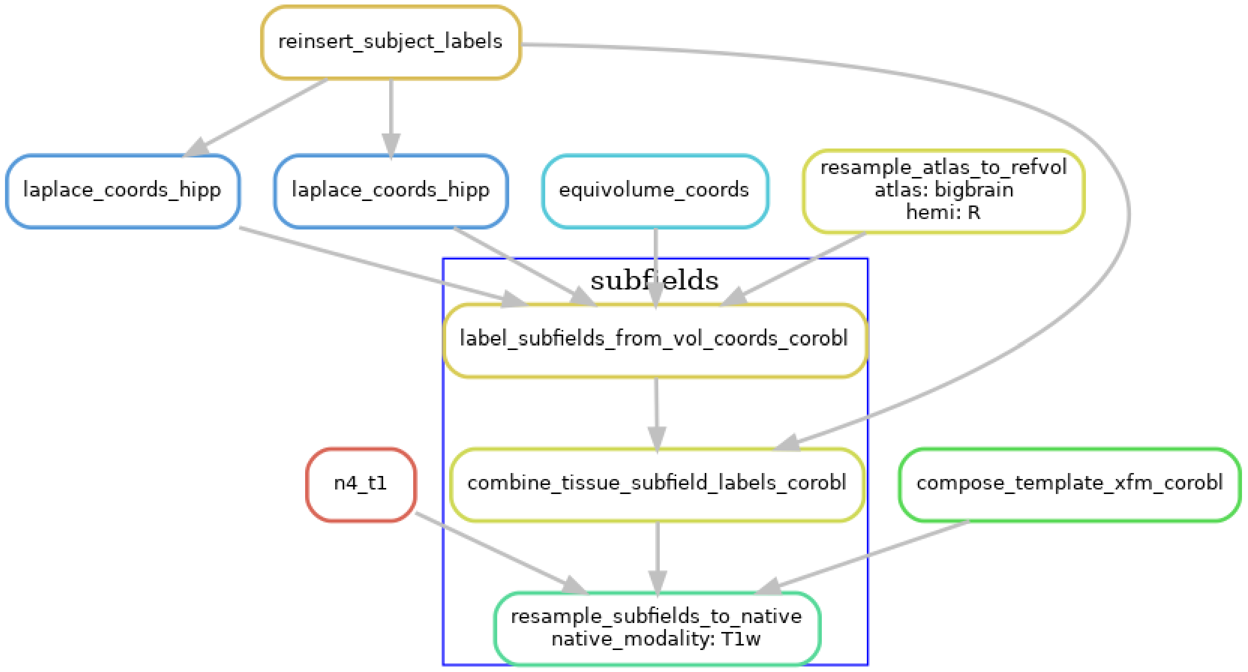

#### 7.7 Generating warp files

To allow users to transform data between the different spaces, we generate warp files that can be applied to transform volumes of surfaces to and from the native and unfolded spaces.

The following diagram shows the workflow, but simplified to contain one hemisphere (`--hemi R`), and excluding the dentate gyrus.

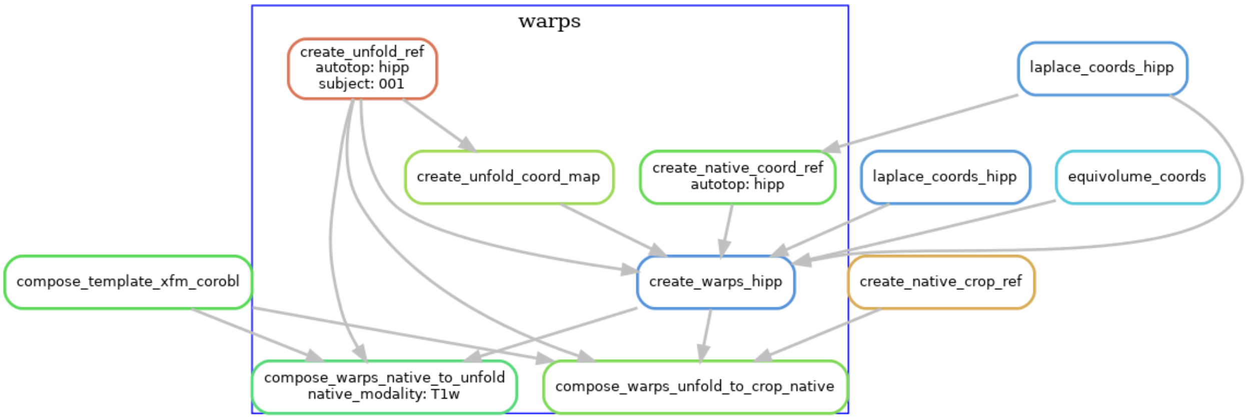

#### 7.8 Surface processing

Using the warps, we transform standard template unfolded meshes to each subject hippocampus, in order to obtain surface meshes in the native space. These are stored in GIFTI format, and we also produce metric files to quantify surface morphometry (thickness, gyrification, curvature).

The following diagram shows the workflow, but simplified to contain one hemisphere (`--hemi R`), and excluding the dentate gyrus.

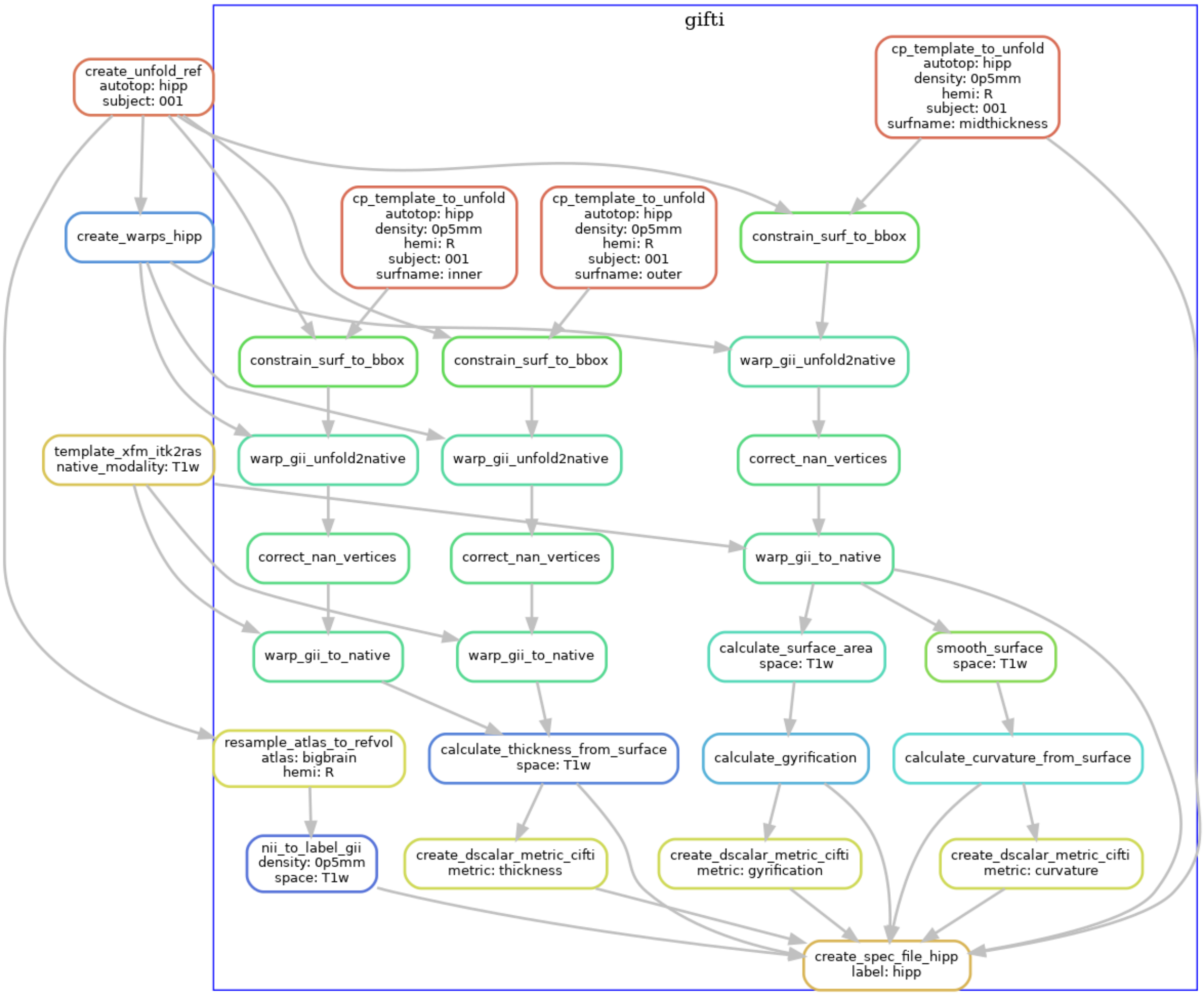

#### 7.9 Additional steps

Resampling to output resolution, quality control snapshot generation, and archiving the work folder are steps that are also carried out by the workflow, but the DAGs are now shown here because of the many inputs/outputs, and generally have straightforward workflow structures.

### CHAPTER EIGHT

### OUTPUT FILES

The PATH_TO_OUTPUT_DIR folder contains a logs and work folder for troubleshooting, but for most purposes all the outputs of interest will be in a subfolder called hippunfold with the following structure:

```
hippunfold/
└── sub-{subject}
    ├── anat
    ├── coords
    ├── qc
    ├── surf
    └── warps
```

Briefly, anat contains preprocessed volumetric input images and output segmentations in nifti format, surf contains surface data in gifti format, coords contain Laplace fields spanning the hippocampus, warps contains transformations between unfolded and native or 'unfolded' space, and qc contains snapshots and useful diagnostic information for quality control.

#### 8.1 anat

This folder contains input anatomical images that have been non-uniformity corrected, motion-corrected, and, where appropriate, averaged and registered. In this example, a T1w image was used as a standard reference image, but a T2w was also registered and used in tissue segemntation:

```
sub-001
 └── anat
     ├── sub-001_desc-preproc_T1w.nii.gz
     ├── sub-001_space-T1w_desc-preproc_T2w.nii.gz
     ├── sub-001_hemi-R_space-T1w_desc-subfields_atlas-bigbrain_dseg.nii.gz
     ├── sub-001_hemi-R_space-cropT1w_desc-preproc_T2w.nii.gz
     └── sub-001_hemi-R_space-cropT1w_desc-subfields_atlas-bigbrain_dseg.nii.gz
```

As per BIDS guidelines, desc-preproc refers to preprocessed input images, space-T1w refers to the volume to which the image is registered, hemi refers to the left or right hemisphere (only shown for the right in this example), and dseg (discrete-segmentation) images with desc-subfields contains subfield labels (coded as integers as described in the included volumes.tsv file). The subfield atlas used will also be included, by default as atlas-bigbrain. Note that HippUnfold does most intermediate processing in an unshown (available in the work/ folder) space-corobl which is cropped, upsampled, and rotated. Downsampling to the original T1w space can thus degrade the results and so they are also provided in a higher resolution space-cropT1w space which is ideal for conducting volumetry or morphometry measures with high precision and detail.

For example, the following image shows a whole-brain `T1w` image, a `space-cropT1w` overlay of the upsampled T2w image (centre square), and a similarly upsampled output subfield segmentation (colour).

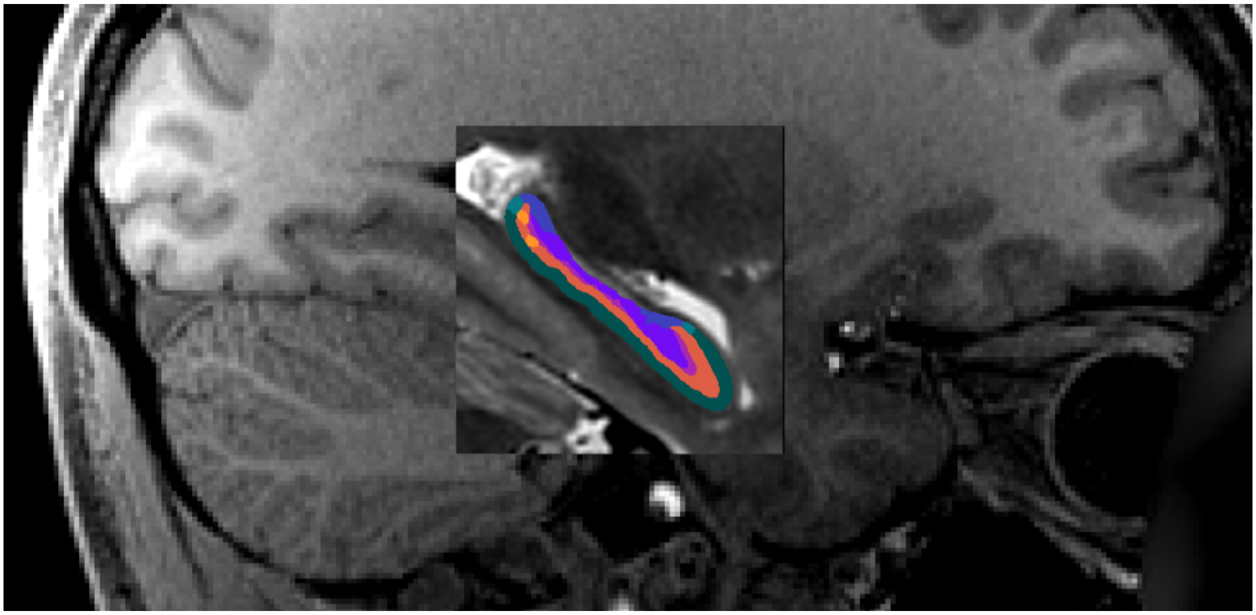

#### 8.2 surf

##### 8.2.1 surface meshes

Surface meshes (geometry files) are in `.surf.gii` format, and are provided in both the native space (`space-T1w`) and the unfolded space (`space-unfolded`). In each space, there are `inner`, `midthickness`, and `outer` surfaces, which correspond to `white`, `midthickness`, and `pial` for cortical surfaces:

```
sub-{subject}
 └── surf
     └── sub-001_hemi-R_space-{T1w,unfolded}_den-0p5mm_{inner,midthickness,outer}.surf.
↪gii
```

The following shows surfaces `inner`, `midthickness`, and `outer` in yellow, orange, and red, respectively.

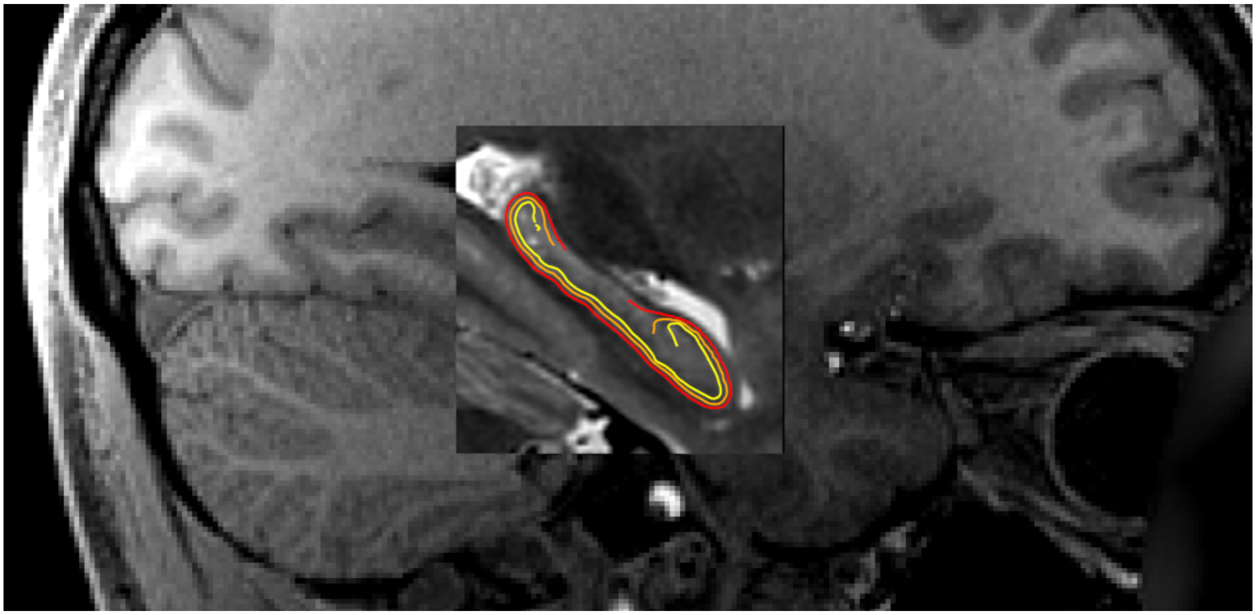

##### 8.2.2 surface densities

Surfaces are provided in different density configurations, and are labelled based on the approximate vertex spacing in each. The default density is `0p5mm`, which has an approximate vertex spacing of 0.5mm. There are also `1mm` and `2mm` surfaces which have 1mm or 2mm spacing, respectively (suitable for lower-resolution BOLD data). Previous versions of hippunfold exclusively used a `unfoldiso` template surface, formed by a 254x126 grid in the unfolded space, however a downside of this template is that it results in very non-uniform vertex spacing when transformed to the native space. The newer `0p5mm`, `1mm` and `2mm` surfaces are designed to have closer to uniform vertex spacing in native space, though vertex spacing will not remain uniform when unfolded. This is illustrated in the the following `den-1mm` mesh in folded and unfolded space.

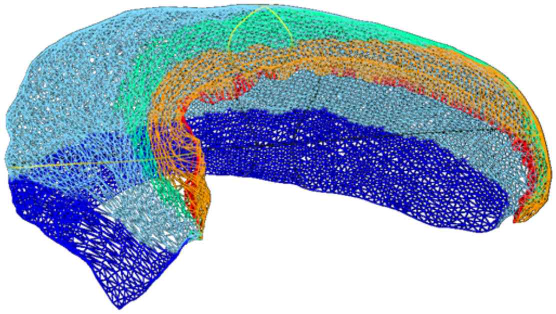

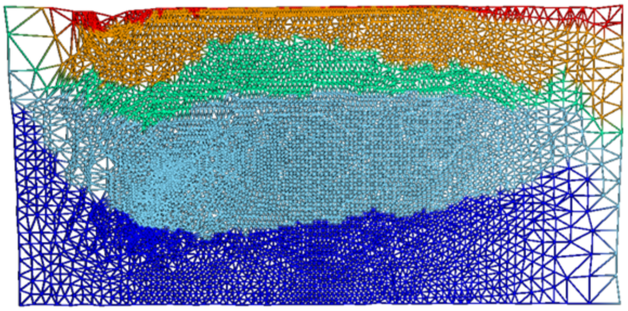

All surfaces of the same density (e.g. `1mm`), in both `space-T1w` and `space-unfolded`, share the same mesh topology and have corresponding vertices with each other. The vertex locations for unfolded surfaces are identical for all subjects as well (note that this of course is not the case for the `space-T1w` surfaces).

##### 8.2.3 surface metrics

In addition to the geometry files, surface-based shape metrics are provided in `.shape.gii` format. The thickness, curvature and surface area are computed using the same methods as cortical surfaces, based on the surface geometry files, and are provided in the `T1w` space. The gyrification metric is the ratio of native to unfolded surface area, or equivalently, the scaling or distortion factor when unfolding:

```
sub-{subject}
 └── surf
     └── sub-001_hemi-{L,R}_space-T1w_den-0p5mm_label-hipp_{thickness,curvature,
↪gyrification}.shape.gii
     └── sub-001_hemi-{L,R}_space-T1w_den-0p5mm_label-dentate_{curvature,gyrification}.
↪shape.gii
```

These metrics are shown in both folded and unfolded space in the images below. Note that these results are from group-averaged data and so individual subject maps may show considerably more variability.

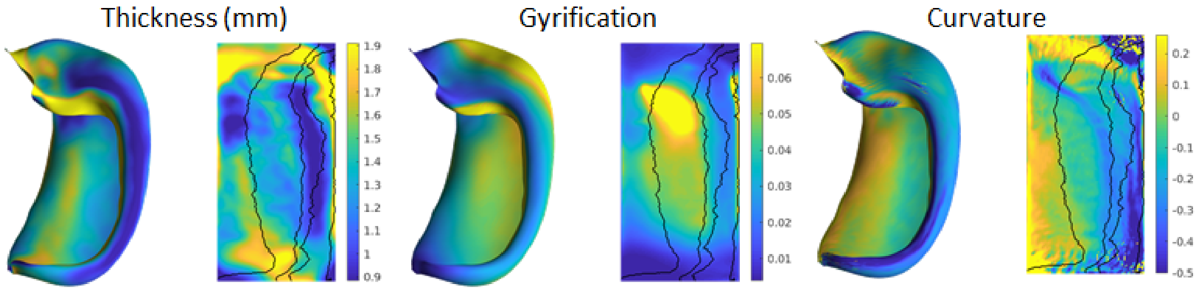

##### 8.2.4 surface labels

The subfield labels from unfolded atlases are also provided for each subject, in `.label.gii` format. Analogous to the volume-based labels, the name of the atlas (default: `bigbrain`) is in the file name.

```
sub-{subject}
 └── surf
     └── sub-001_hemi-{L,R}_space-T1w_den-0p5mm_label-hipp_atlas-bigbrain_subfields.
↪label.gii
```

##### 8.2.5 cifti files

In addition to lateralized `.shape.gii` and `.label.gii` metrics and labels, we also provide data mapped to hippocampi from hemispheres in a single file using the corresponding CIFTI formats, `.dscalar.nii` and `.dlabel.nii`. Note: since CIFTI does not support hippocampus surfaces (yet), we make use of the `CORTEX_LEFT` and `CORTEX_RIGHT` labels for the hippocampal surfaces.

```
sub-{subject}
 └── surf
     ├── sub-001_space-T1w_den-0p5mm_label-{hipp,dentate}_{thickness,curvature,
↪gyrification}.dscalar.nii
     └── sub-001_space-T1w_den-0p5mm_label-hipp_atlas-bigbrain_subfields.dlabel.nii
```

##### 8.2.6 spec files

Finally, these files are packaged together for easy viewing in Connectome Workbench, `wb_view`, in the following `.spec` files, for each hemisphere and structure separately, and combined:

```
sub-{subject}
 └── surf
     ├── sub-001_hemi-{L,R}_space-T1w_den-0p5mm_label-{hipp,dentate}_surfaces.spec
     └── sub-001_space-T1w_den-0p5mm_label-{hipp,dentate}_surfaces.spec
```

##### 8.2.7 New: label-dentate

HippUnfold v1.0.0 introduces `label-dentate` files which represent a distinct surface making up the dentate gyrus (reflecting its distinct topology from the rest of the cortex). The rest of the surfaces are given the name `label-hipp` to differentiate them from these new files.

These are illustrated in the following image (orange represents the usual hippocampal midthickness surface, while violet shows the new `dentate` surface):

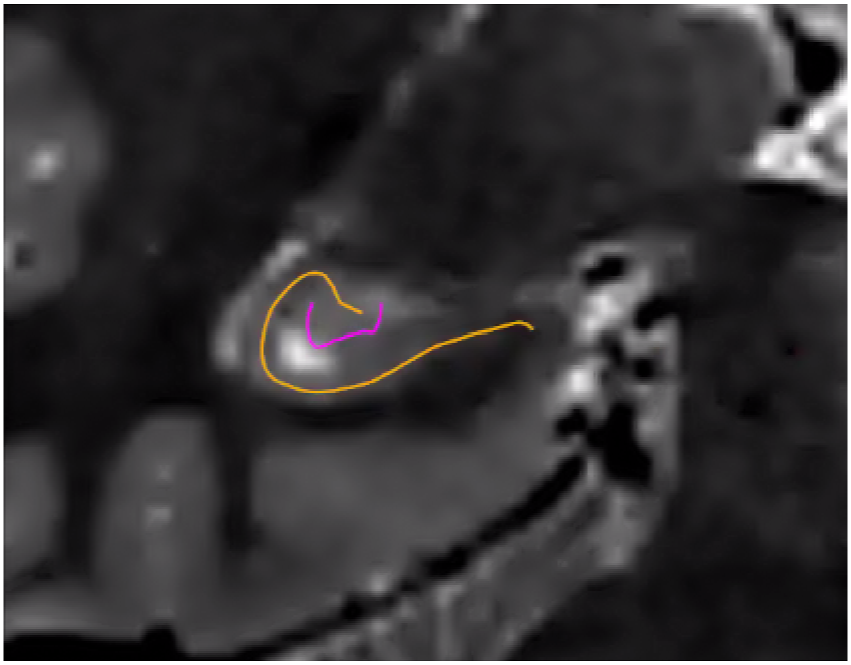

Note that the dentate uses the same unfolding methods as the rest of the hippocampus, but with several caveats. Given its small size, its boundaries are not easily deliminated and so `inner`, `outer`, and `thickness` gifti surfaces are omitted. Furthermore, Laplace coordinates and therefore vertex spacing are not guaranteed to be topologically equivalent as they are obtained through volumetric registration with the template shape injection step of this workflow.

Corresponding `coords` and `warp` files are also generated.

##### 8.2.8 New: myelin maps

If your dataset has T1w and T2w images (and you are using --modality=T1w or --modality=T2w), then you can enable the generation of myelin maps as the ratio of T1w over T2w images. This division is done in the corobl space, and provides myelin.shape.gii surface metrics, and also includes these in the CIFTI and spec files.

This option is enabled with the --generate-myelin-maps command-line option.

#### 8.3 coords

Hippunfold also provides images that represent anatomical gradients along the 3 principal axes of the hippocampus, longitudinal from anterior to posterior, lamellar from proximal (dentate gyrus) to distal (subiculum), and laminar from inner (SRLM) to outer. These are provided in the images suffixed with coords.nii.gz with the direction indicated by dir-{direction} as AP, PD or IO, and intensities from 0 to 1, e.g. 0 representing the Anterior end and 1 the Posterior end.

Here is an example showing coronal slices of the hippocampus with the PD, IO, and AP (sagittal slice) overlaid.

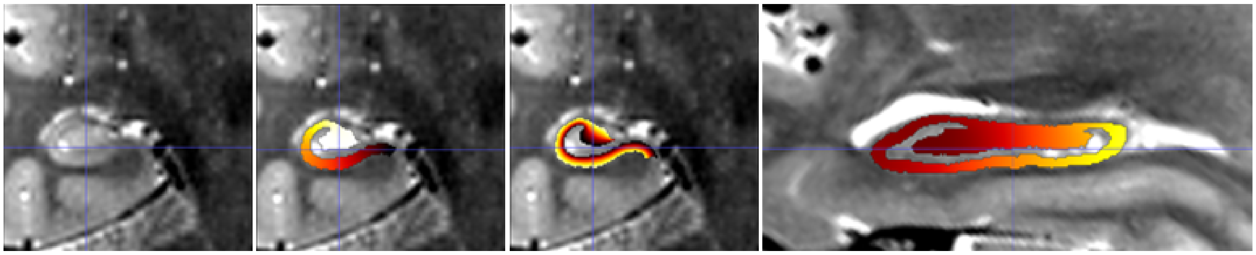

Note that these images have been resampled to space-corobl which is the space in which most processing is done internally. These can be seen in the work/ output directory or specified as a possible output space.

#### 8.4 warps

ITK transforms to warp images between the T1w space to the unfolded space, are provided for each hippocampus:

```
sub-{subject}
 └── seg
     └── sub-001_hemi-R_from-T1w_to-unfold_mode-image_xfm.nii.gz
     └── sub-001_hemi-R_from-unfold_to-T1w_mode-image_xfm.nii.gz
```

These are ITK transforms that can transform any image that is in T1w space (can be any resolution and FOV, as long as aligned to T1w), to the unfolded hippocampal volume space, and vice-versa. You can use the warp itself as a reference image, e.g.:

```
antsApplyTransforms -d 3 \
-i sub-001_space-T1w_FA.nii.gz \
-o sub-001_hemi-L_space-unfolded_FA.nii.gz \
-t sub-001_hemi-L_from-T1w_to-unfold_mode-image_xfm.nii.gz \
-r sub-001_space-unfolded_refvol.nii.gz
```

#### 8.5 Additional Files

The top-level PATH_TO_OUTPUT_DIR contains additional folders:

```
├── hippunfold
├── logs
├── work
└── .snakemake
```

The hidden .snakemake folder contains a record of the code and parameters used, and paths to the inputs.

Workflow steps that write logs to file are stored in the logs subfolder, with file names based on the rule wildcards (e.g. subject, hemi, etc..).

Intermediate files are stored in the work folder. These files and folders, similar to results, are generally named according to BIDS. This folder will have tar.gz files for each subject, unless the --keep_work option is used.

If the app is run in workflow mode (--workflow-mode/-W) which enables direct use of the snakemake CLI to run hippunfold, the hippunfold and work folders will be placed in a results folder.

CHAPTER

NINE

### VISUALIZATION

#### 9.1 Freeview (volumes and surfaces)

Freeview is a powerful viewer that works well with both volume and surface data. It (currently) has several quirks worth noting, such as:

- All surfaces should be loaded before volumes (otherwise their orientation will be incorrect).
- Loading surface metric data can be difficult, see the examples below.
- Crashes can still occur occasionally, especially when trying to load surfaces or surface overlay data with the incorrect buttons.

##### 9.1.1 The example below will act as a guide to avoid common mistakes:

- open Freeview by simple typing `freeview` in the command line.
- Add a hippunfold surface: File > Load Surface > Navigate to a surface (eg. `hippunfold/sub-01/surf/sub-02_hemi-L_space-T1w_den-0p5mm_label-hipp_midthickness.surf.gii`).
- Add the correponding T1w image: File > Load Volume > Navigate to a volume (eg. `hippunfold/sub-01/anat/sub-02_desc-preproc_T1w.nii.gz`). You should now see a hippocampal surface projected onto the coronal, sagittal, and axials views over a T1w image. You should also see a 3D model of the hippocampus. You can toggle visibility of each file by unticking or ticking it in the left panel. You can also adjust the ordering ov overlays here.
- Add shape data as an overlay on the midthickness surface: With the surface file selected, use the left information panel to select Overlay > Load Generic and navigate to a metric file (eg. `hippunfold/sub-01/surf/sub-02_hemi-L_space-T1w_den-0p5mm_label-hipp_gyrification.shape.gii`). You should now have a Configure button which you can use to adjust the windowing and colormap of this data.
- Subfield labels (`.label.gii` files) can be loaded similarly to metric data, but using the Annotation button on the left panel instead of Overlay.
- Add a segmentation image (eg. `hippunfold/sub-01/anat/sub-01_hemi-L_space-cropT1w_desc-subfield_dseg.nii.gz`). On the left panel, set Colormap > Lookup Table and then Select Lookup Table > Load Lookup Table > Navigate to `YOUR_HIPPUNFOLD_INSTALLATION_DIRECTORY/hippunfold/resources/desc-subfields_freeview_desg.tsv`. You should now see standardized subfield colours and names in the bottom panel when you mouse over a given subfield.

##### 9.1.2 Other visualization tips and tricks:

- Consider adding dentate surfaces (`label-dentate`), unfolded surfaces (`space-unfolded`), or other overlay data (eg. `thickness`).
- Ensure surfaces and metric data are sampled with the same number of vertices (eg. `label-hipp` and den-'0p5mm'), and note that folded and unfolded surfaces will appear far apart in the 3D viewer and so you may need to zoom out quite far to navigate to them.
- Any metric data loaded on a folded (eg. `space-T1w`) surface can also be viewed on an unfolded surface that has the same number of vertices.
- Add Laplace coordinates (eg. `hippunfold/sub-01/coords/sub-01_dir-AP_hemi-L_space-cropT1w_label-hipp_desc-nii.gz`) over your T1w image, then set the minimum windowing to 0.001 and tick the box `Clear background` in the left panel to maintain visibility of the T1w underlay.
- Note that `space-T1w` and `space-cropT1w` should appear in equivalent positions in Freeview, despite having a different field of view and resolution. If loading these data into Matlab or Python, you will need to first resample these images to the same reference image in order to index voxels from equivalent points.

#### 9.2 HippUnfold Toolbox

The HippUnfold Toolbox provides examples and functions for mapping data onto hippocampal surfaces, plotting surfaces, and performing comparisons or statistical tests between subjects. Note that this can be done in other programs, like Connectome workbench, but these Python & Matlab tools should give an idea of how this can be done in a fully customizable fashion.

#### 9.3 ITK-SNAP (volumes)

ITK-SNAP is a lightweight tool able to quickly open volumes, and is ideal for manual segmentation or edits. Segmentation images (`_dseg.nii.gz`) can be loaded as overlays and ITK-SNAP will create a 3D rendering of the contours of each label. Use the `space-cropT1w` or `space-cropT2w` images in the `anat/` folder to visualize one hemisphere at a time.

#### 9.4 Connectome Workbench (surfaces)

Connectome Workbench view tool allows for advanced visualization of surfaces and surface data. Loading one of the `.spec` files produced by HippUnfold into `wb_view` will allow you to visualize the hippocampus in native and unfolded configurations, and also overlay metric and label data (e.g. subfields and thickness).

CHAPTER

TEN

### CONTRIBUTING TO HIPPUNFOLD

Hippunfold dependencies are managed with Poetry, which you'll need installed on your machine. You can find instructions on the poetry website.

#### 10.1 Set-up your development environment:

Clone the repository and install dependencies and dev dependencies with poetry:

```
git clone http://github.com/khanlab/hippunfold
cd hippunfold
poetry install
```

Poetry will automatically create a virtual environment. To customize where these virtual environments are stored see poetry docs here

Then, you can run hippunfold with:

```
poetry run hippunfold
```

or you can activate a virtualenv shell and then run hippunfold directly:

```
poetry shell
hippunfold
```

You can exit the poetry shell with `exit`.

#### 10.2 Running code format quality checking and fixing:

Hippunfold uses poethepoet as a task runner. You can see what commands are available by running:

```
poetry run poe
```

We use `black` and `snakefmt` to ensure formatting and style of python and Snakefiles is consistent. There are two task runners you can use to check and fix your code, and can be invoked with:

```
poetry run poe quality_check
poetry run poe quality_fix
```

Note that if you are in a poetry shell, you do not need to prepend `poetry run` to the command.

#### 10.3 Dry-run testing your workflow:

Using Snakemake's dry-run option (`--dry-run`/`-n`) is an easy way to verify any changes to the workflow are working correctly. The `test_data` folder contains a number of *fake* bids datasets (i.e. datasets with zero-sized files) that are useful for verifying different aspects of the workflow. These dry-run tests are part of the automated github actions that run for every commit.

You can use the hippunfold CLI to perform a dry-run of the workflow, e.g. here printing out every command as well:

```
hippunfold test_data/bids_singleT2w test_out participant --modality T2w -np
```

As a shortcut, you can also use `snakemake` instead of the hippunfold CLI, as the `snakebids.yml` config file is set-up by default to use this same test dataset, as long as you run snakemake from the `hippunfold` folder that contains the `workflow` folder:

```
cd hippunfold
snakemake -np
```

#### 10.4 Instructions for Compute Canada

This section provides an example of how to set up a `pip installed` copy of HippUnfold on CompateCanada's `graham` cluster.

##### 10.4.1 Setting up a dev environment on graham:

Here are some instructions to get your python environment set-up on graham to run HippUnfold:

1. Create a virtualenv and activate it:

   ```
   mkdir $SCRATCH/hippdev
   cd $SCRATCH/hippdev
   module load python/3.8
   virtualenv venv
   source venv/bin/activate
   ```

2. Install HippUnfold

   ```
   git clone https://github.com/khanlab/hippunfold.git
   pip install hippunfold/
   ```

3. Run hippunfold:

   ```
    hippunfold ...
   ```

Note if you want to run hippunfold with modifications to your cloned repository, you either need to pip install again, or run hippunfold the following, since an `editable` pip install is not allowed with pyproject:

```
    python <YOUR_HIPPUNFOLD_DIR>/hippunfold/run.py
```

##### 10.4.2 Running hippunfold jobs on graham:

Note that this requires neuroglia-helpers for regularSubmit or regularInteractive wrappers, and the cc-slurm snakemake profile for cluster execution with slurm.

In an interactive job (for testing):

```
regularInteractive -n 8
hippunfold PATH_TO_BIDS_DIR PATH_TO_OUTPUT_DIR participant \
--participant_label 001 -j 8
```

Here, the last line is used to specify only one subject from a BIDS directory presumeably containing many subjects.

Submitting a job (for larger cores, more subjects), still single job, but snakemake will parallelize over the 32 cores:

```
regularSubmit -j Fat \
hippunfold PATH_TO_BIDS_DIR PATH_TO_OUTPUT_DIR participant  -j 32
```

Scaling up to ~hundred subjects (needs cc-slurm snakemake profile installed), submits 1 16core job per subject:

```
hippunfold PATH_TO_BIDS_DIR PATH_TO_OUTPUT_DIR participant \
--profile cc-slurm
```

Scaling up to even more subjects (uses group-components to bundle multiple subjects in each job), 1 32core job for N subjects (e.g. 10):

```
hippunfold PATH_TO_BIDS_DIR PATH_TO_OUTPUT_DIR participant \
--profile cc-slurm --group-components subj=10
```

#### 10.5 Deep learning nnU-net model files

The trained model files we use for hippunfold are large and thus are not included directly in this github repository, and instead are downloaded from Zenodo releases. If you are using the docker/singularity container, `docker://khanlab/hippunfold`, they are pre-downloaded there, in `/opt/hippunfold_cache`.

If you are not using this container, you will need to download the models before running hippunfold, by running:

```
hippunfold_download_models
```

This console script (installed when you install hippunfold) downloads all the models to a cache dir on your system, which on Linux is typically `~/.cache/hippunfold`. To override this, you can set the `HIPPUNFOLD_CACHE_DIR` environment variable before running `hippunfold_download_models` and `hippunfold`.

#### 10.6 Overriding Singularity cache directories

By default, singularity stores image caches in your home directory when you run `singularity pull` or `singularity run`. As described above, hippunfold also stores deep learning models in your home directory. If your home directory is full or otherwise inaccessible, you may want to change this with the following commands:

```
export SINGULARITY_CACHEDIR=/YOURDIR/.cache/singularity
export SINGULARITY_BINDPATH=/YOURDIR:/YOURDIR
export HIPPUNFOLD_CACHE_DIR=/YOURDIR/.cache/hippunfold/
```

If you are running `hippunfold` with the `--use-singularity` option, hippunfold will download the required singularity containers for rules that require it. These containers are placed in the `.snakemake` folder in your hippunfold output directory, but this can be overriden with the Snakemake option: `--singularity-prefix DIRECTORY`
