## Supplemental file 3 for "Automated hippocampal unfolding for morphometry and subfield segmentation with HippUnfold"

hemi=R,subject=6086470

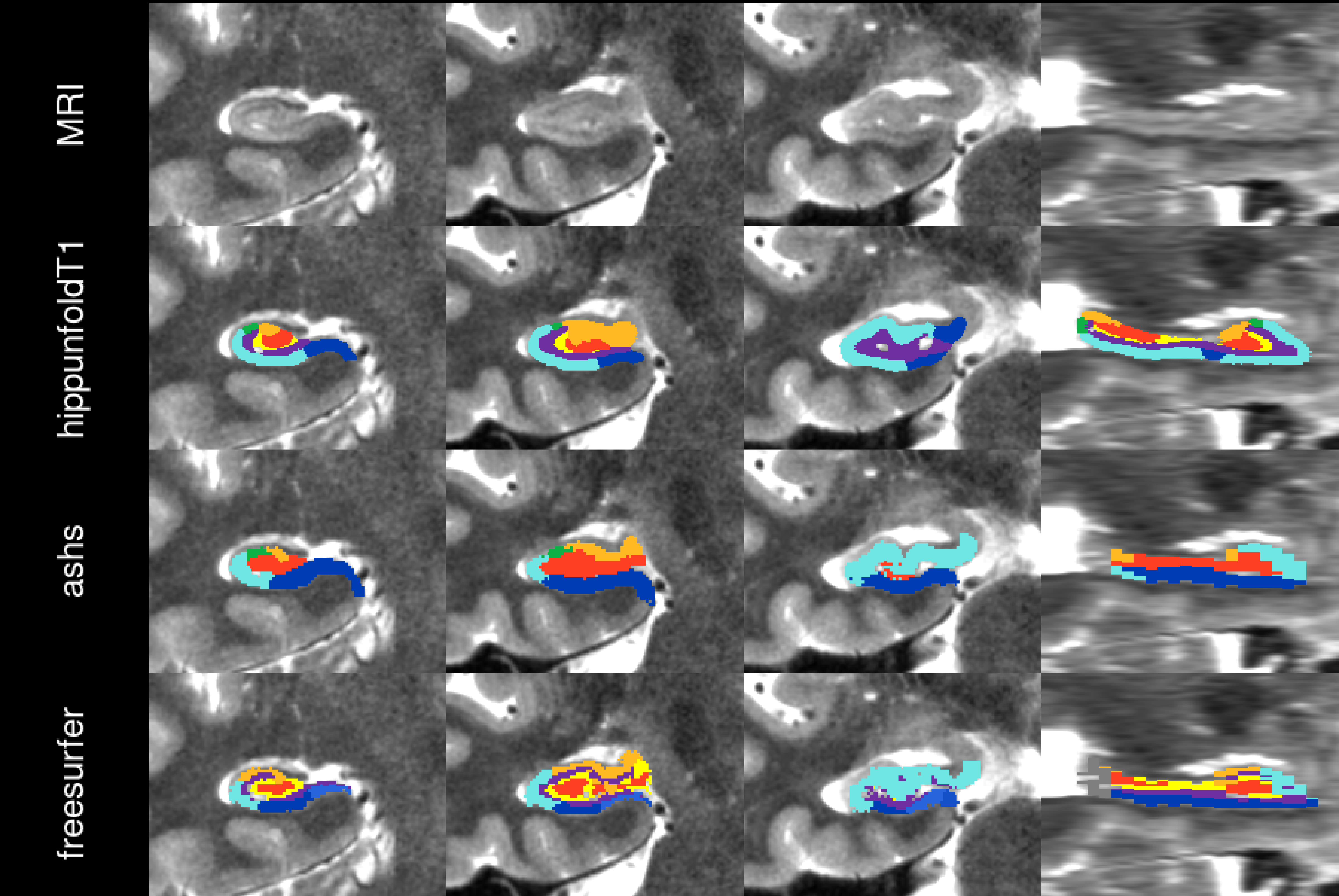

hemi=R,subject=6117051

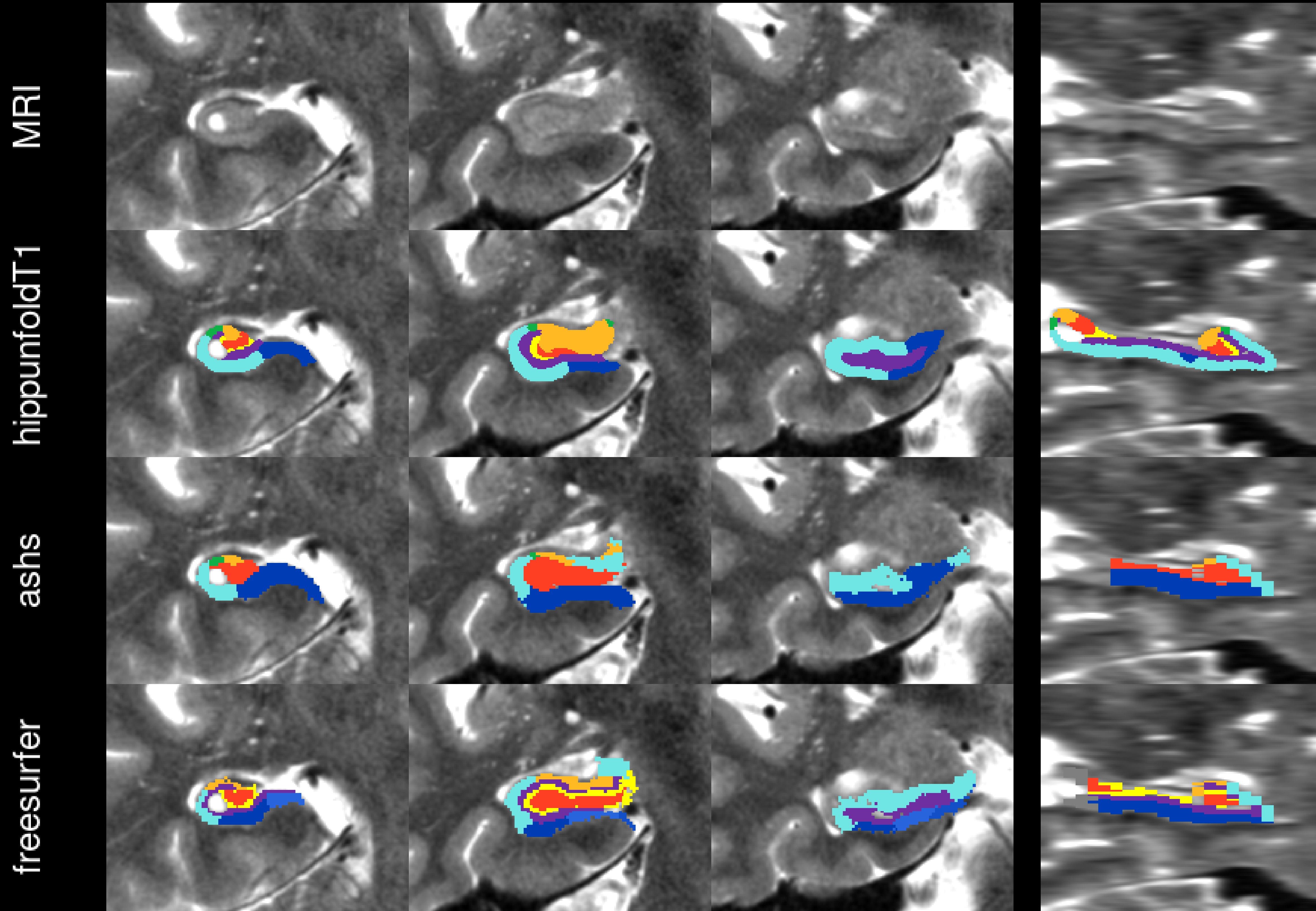

hemi=R,subject=6166973

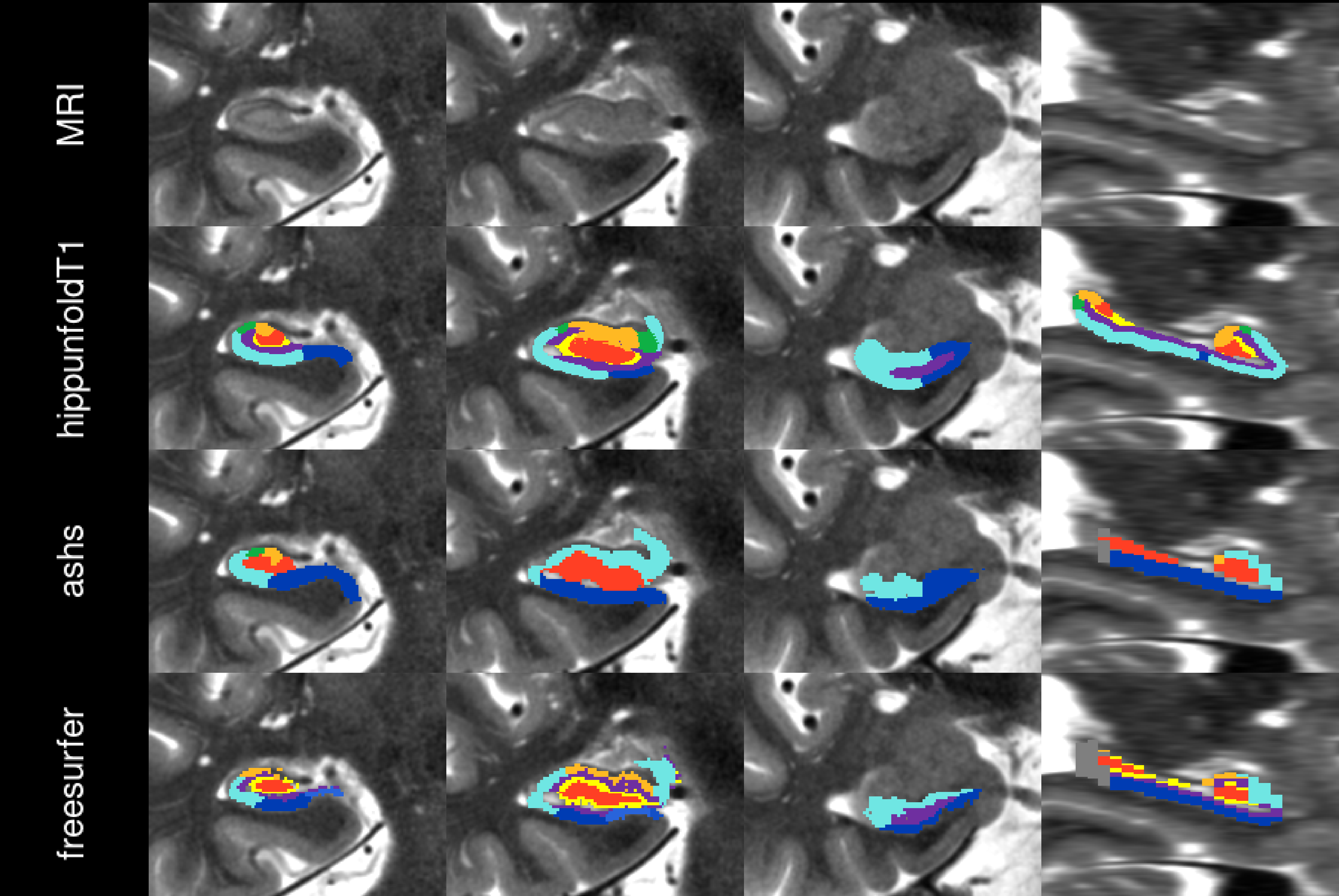

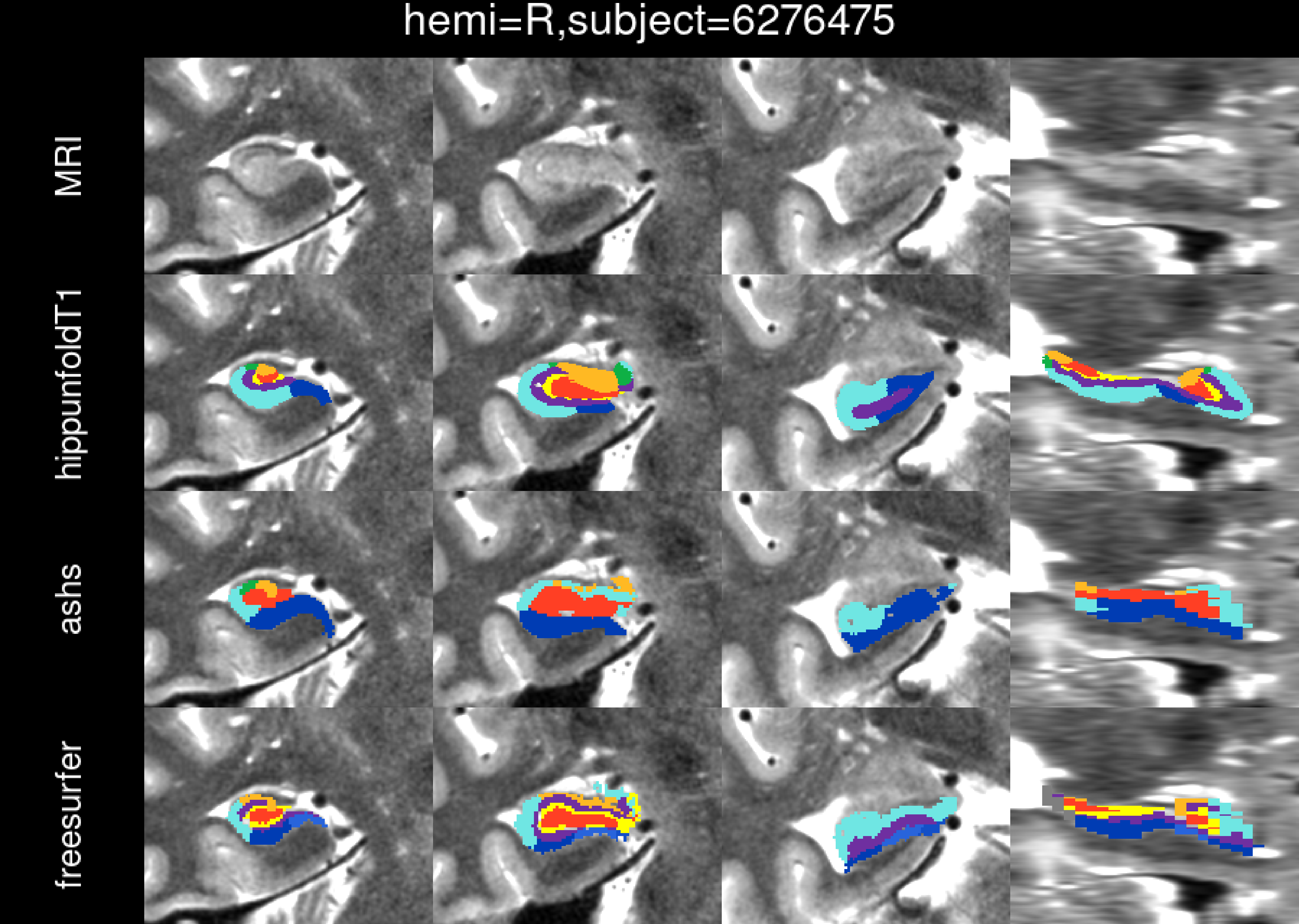
hemi=R,subject=6276475
MRI
hippunfoldT1
ashs
freesurfer

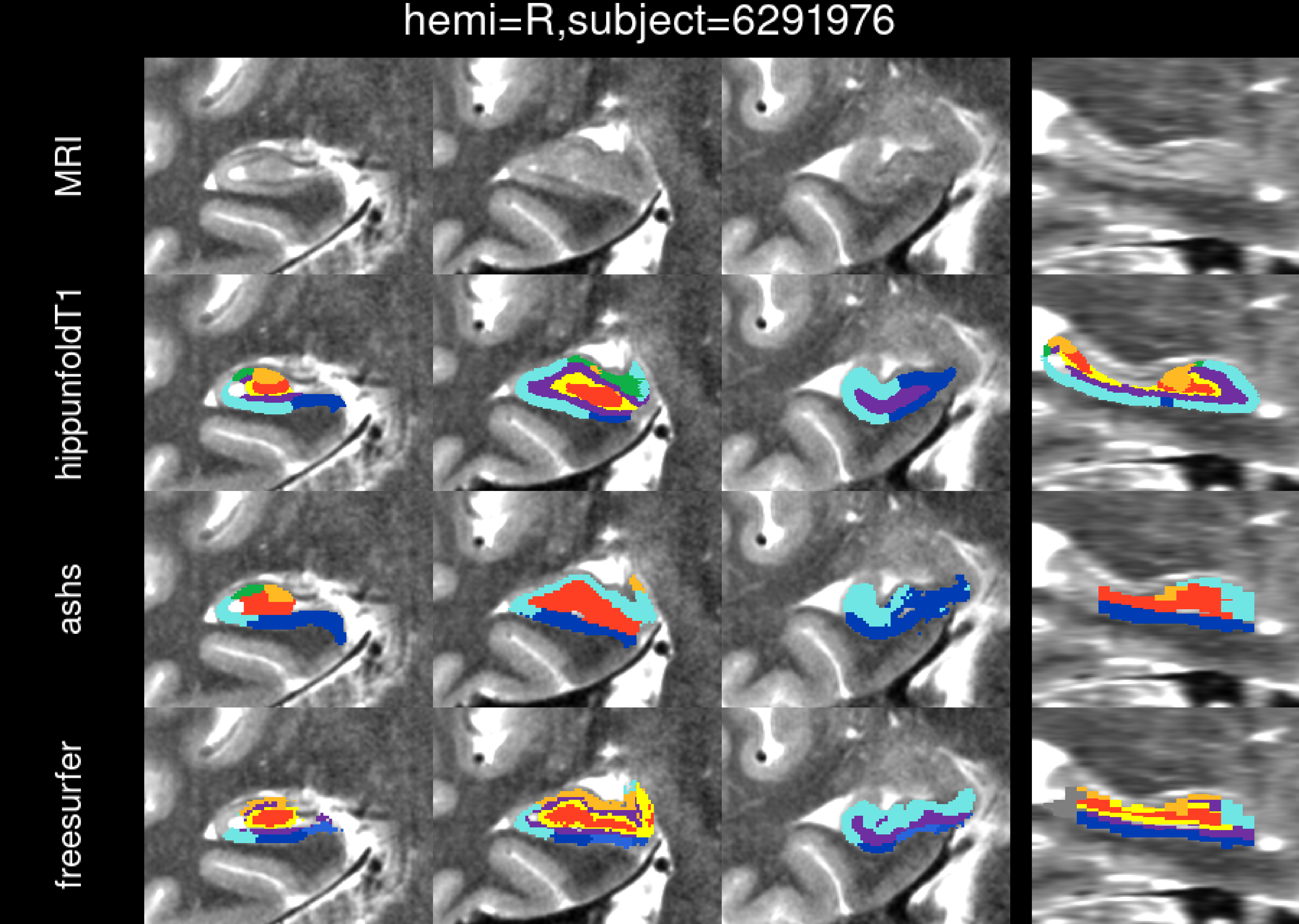
hemi=R,subject=6291976
MRI
hippunfoldT1
ashs
freesurfer

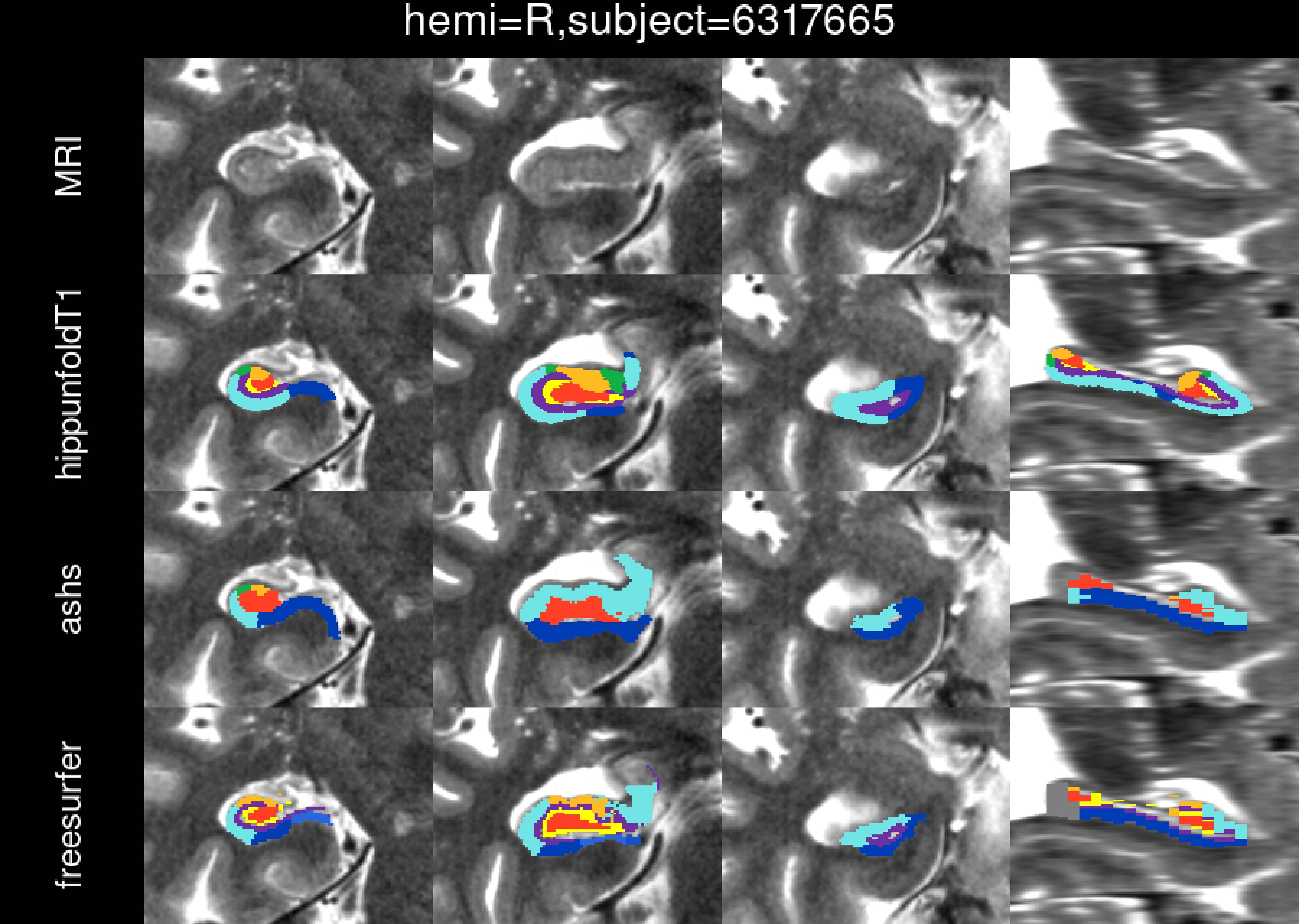
hemi=R,subject=6317665
MRI
hippunfoldT1
ashs
freesurfer

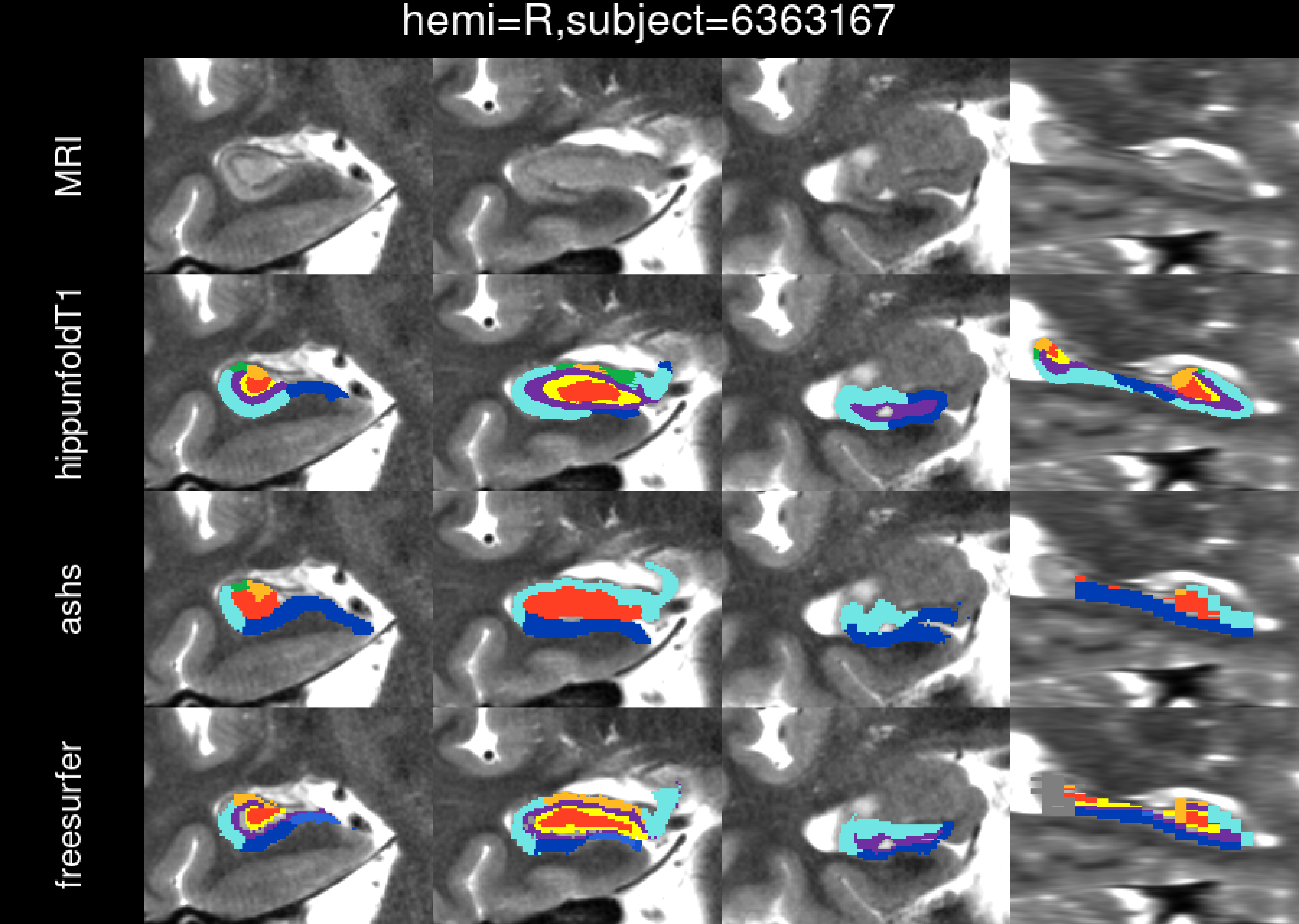
hemi=R,subject=6363167
MRI
hippunfoldT1
ashs
freesurfer

hemi=R,subject=6405157

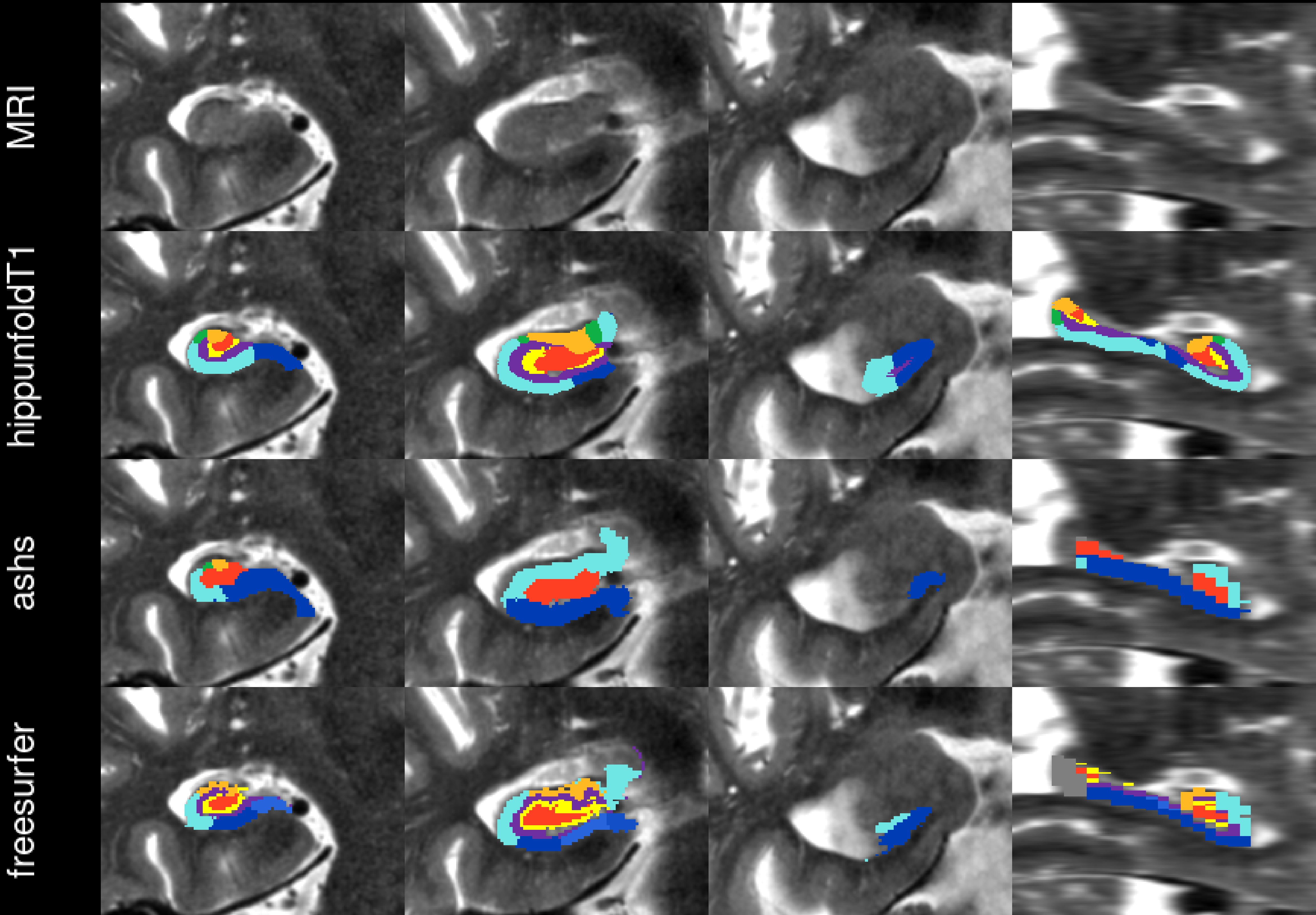

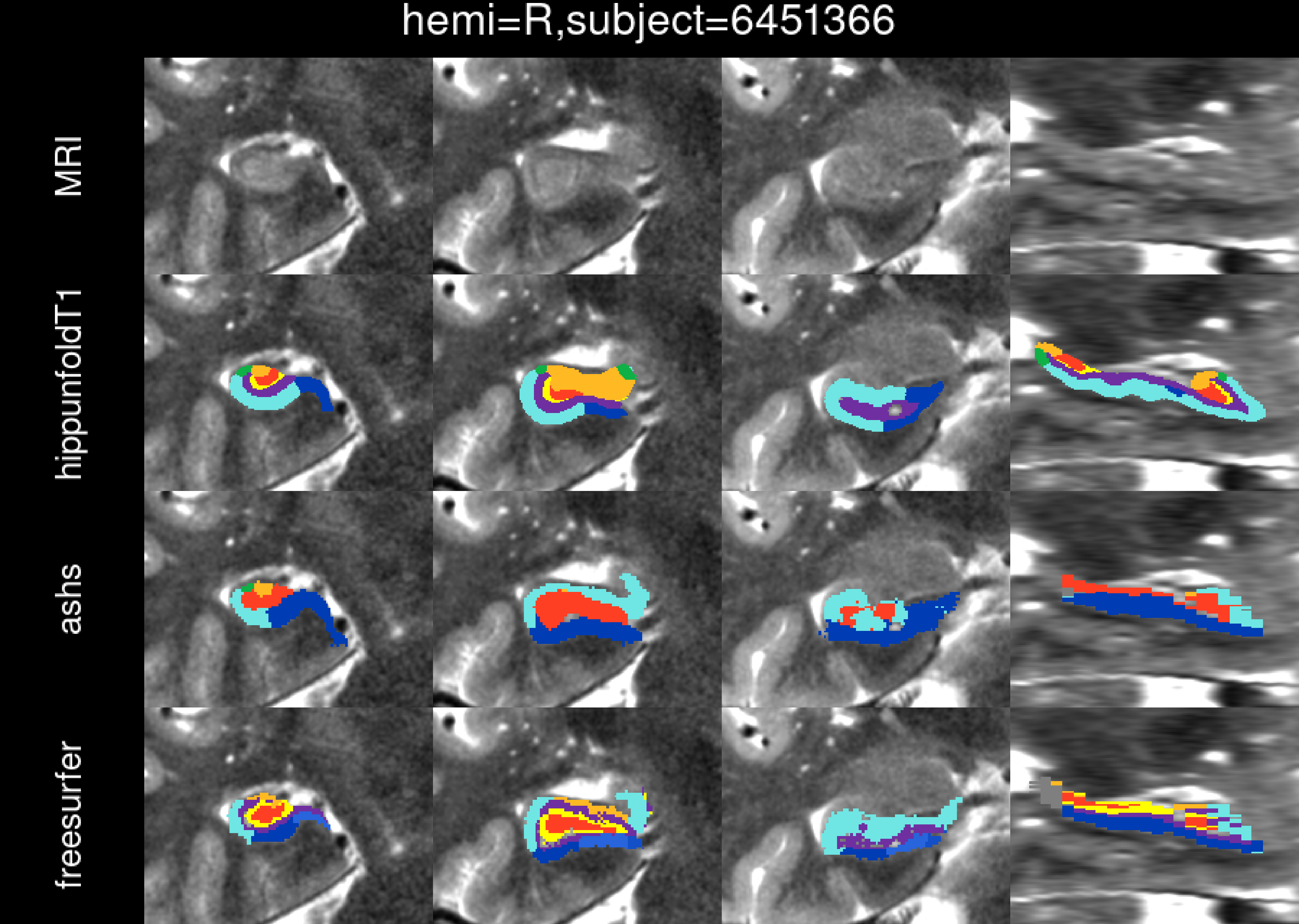
hemi=R,subject=6451366
MRI
hippunfoldT1
ashs
freesurfer

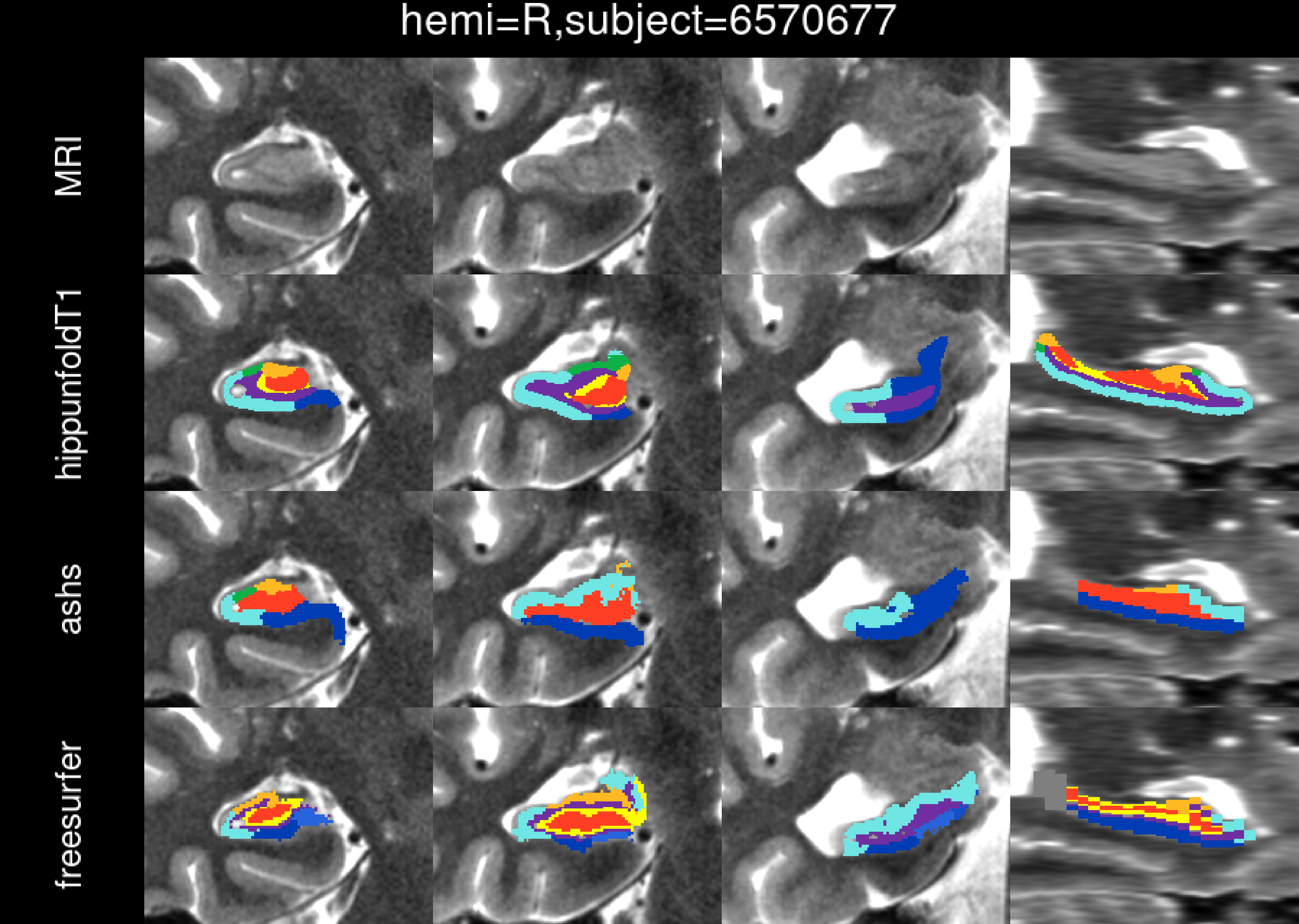
hemi=R,subject=6570677
MRI
hippunfoldT1
ashs
freesurfer

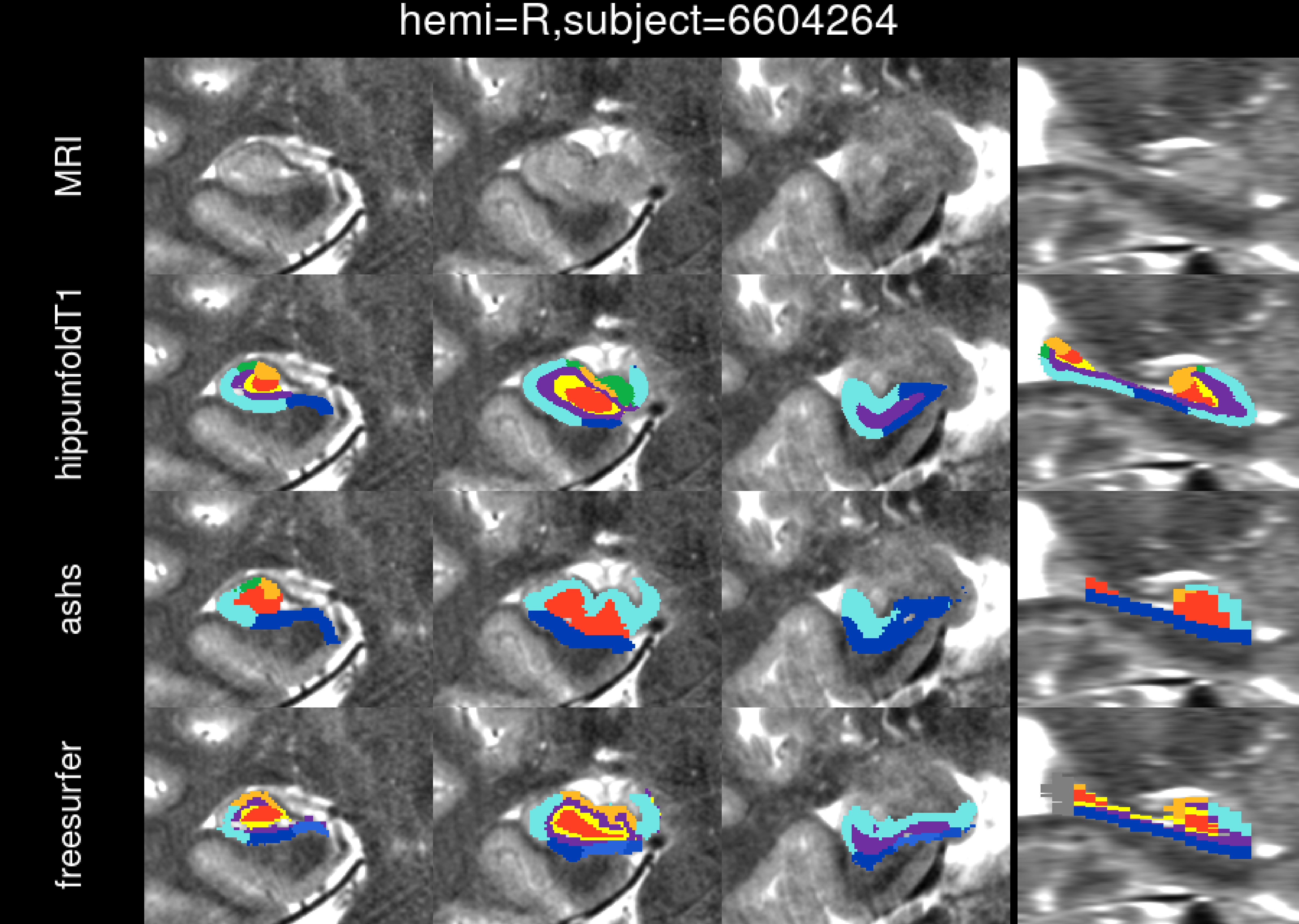
hemi=R,subject=6604264
MRI
hippunfoldT1
ashs
freesurfer

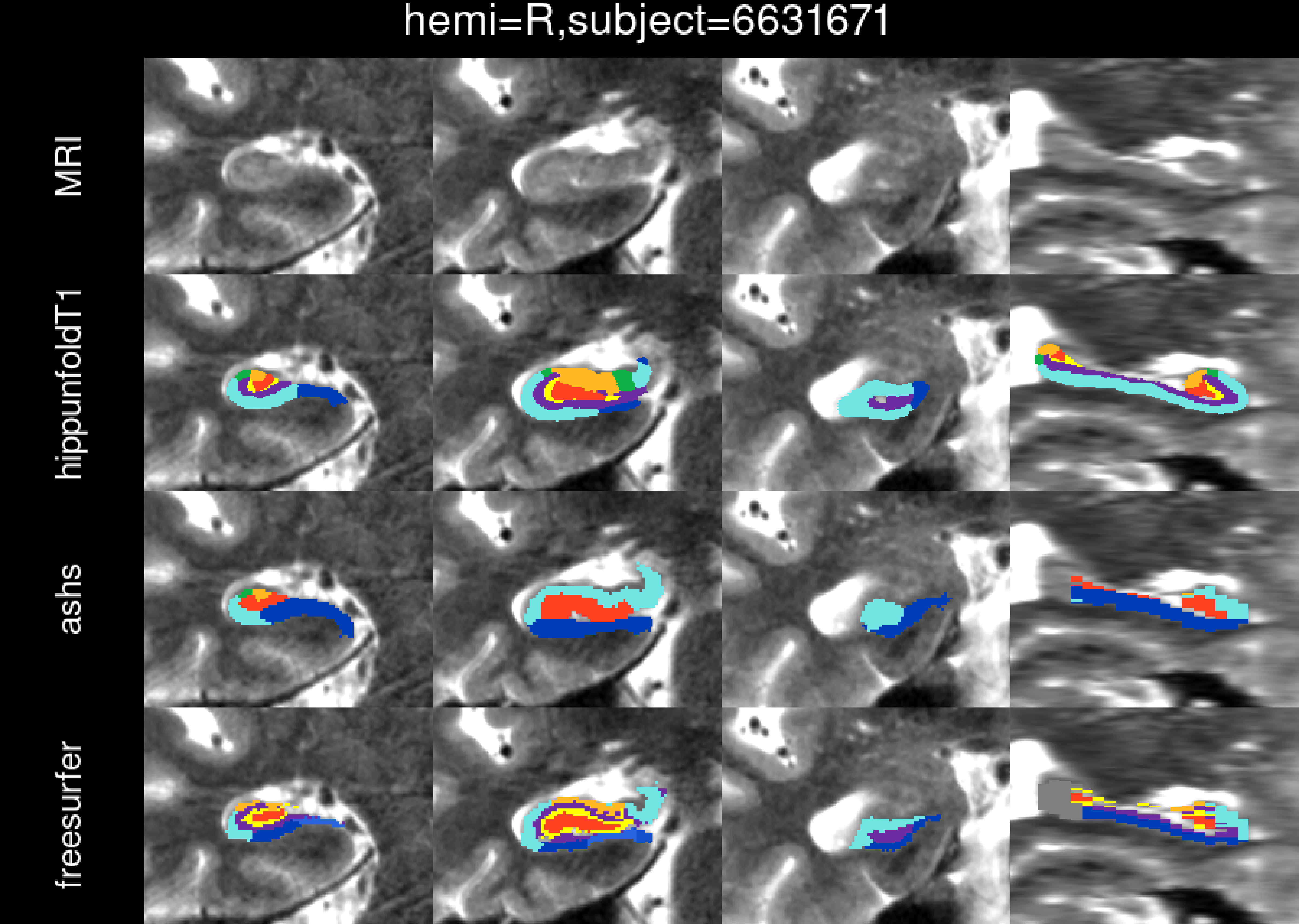
hemi=R,subject=6631671
MRI
hippunfoldT1
ashs
freesurfer

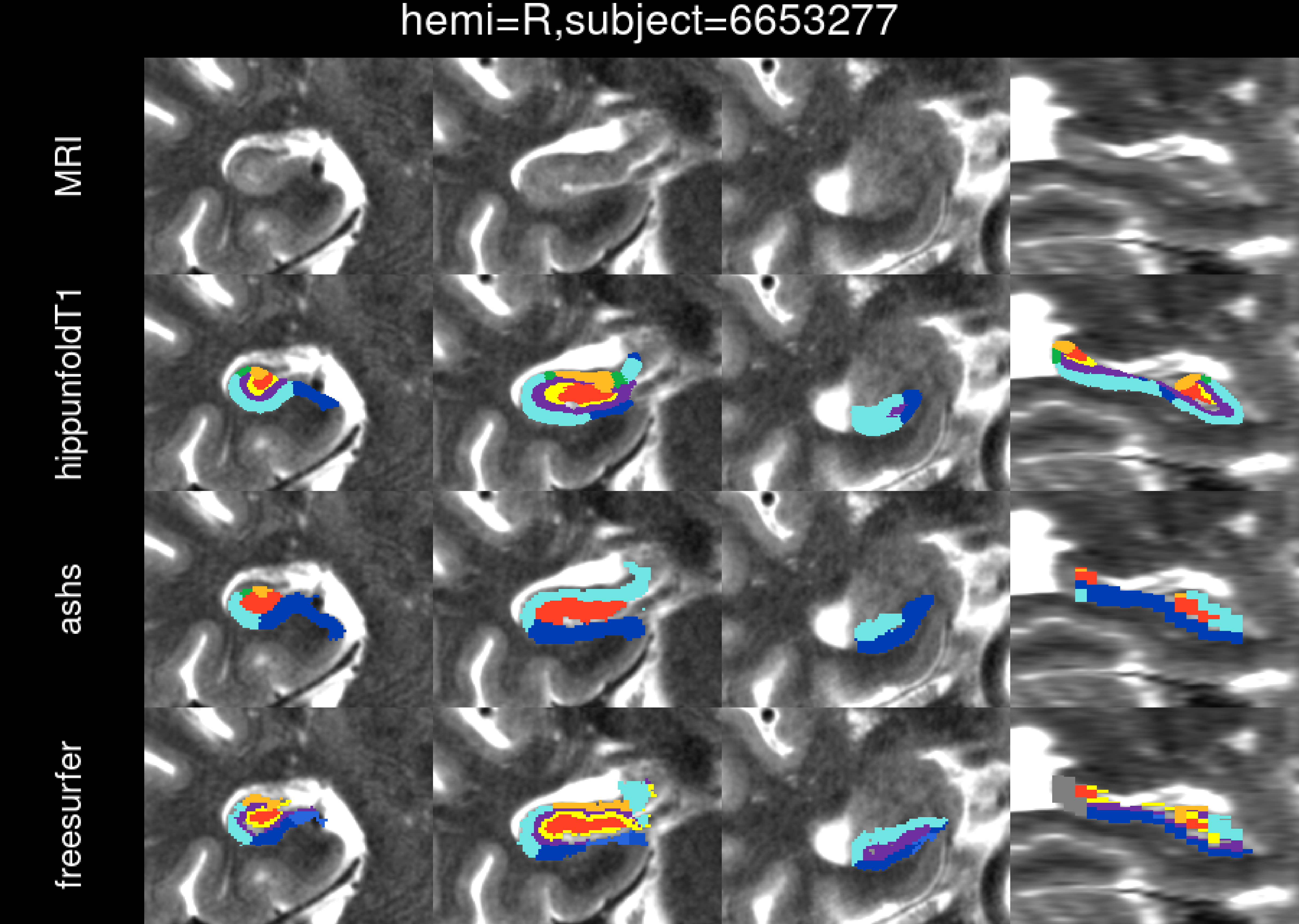
hemi=R,subject=6653277
MRI
hippunfoldT1
ashs
freesurfer

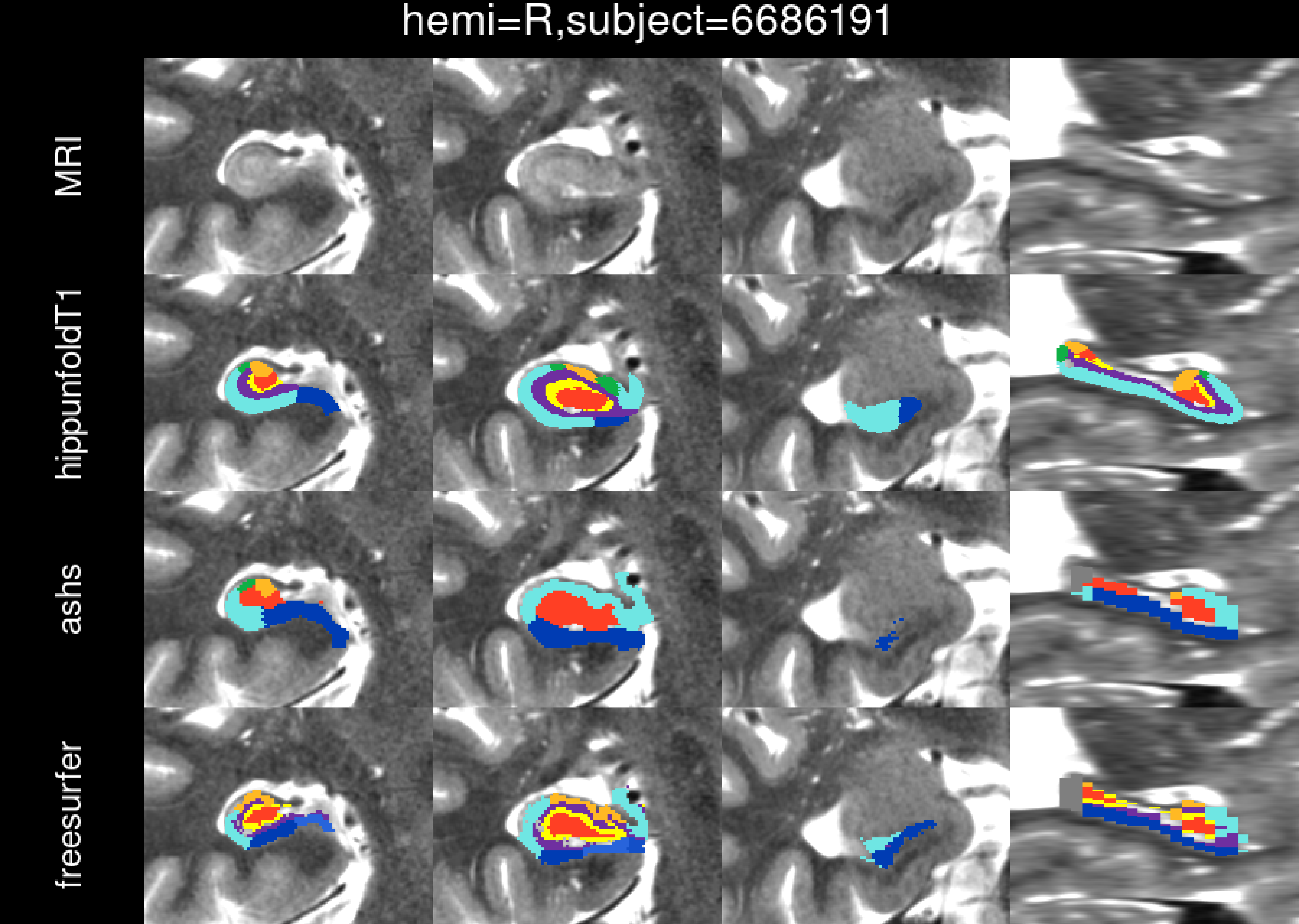

hemi=R,subject=6686191
MRI
hippunfoldT1
ashs
freesurfer

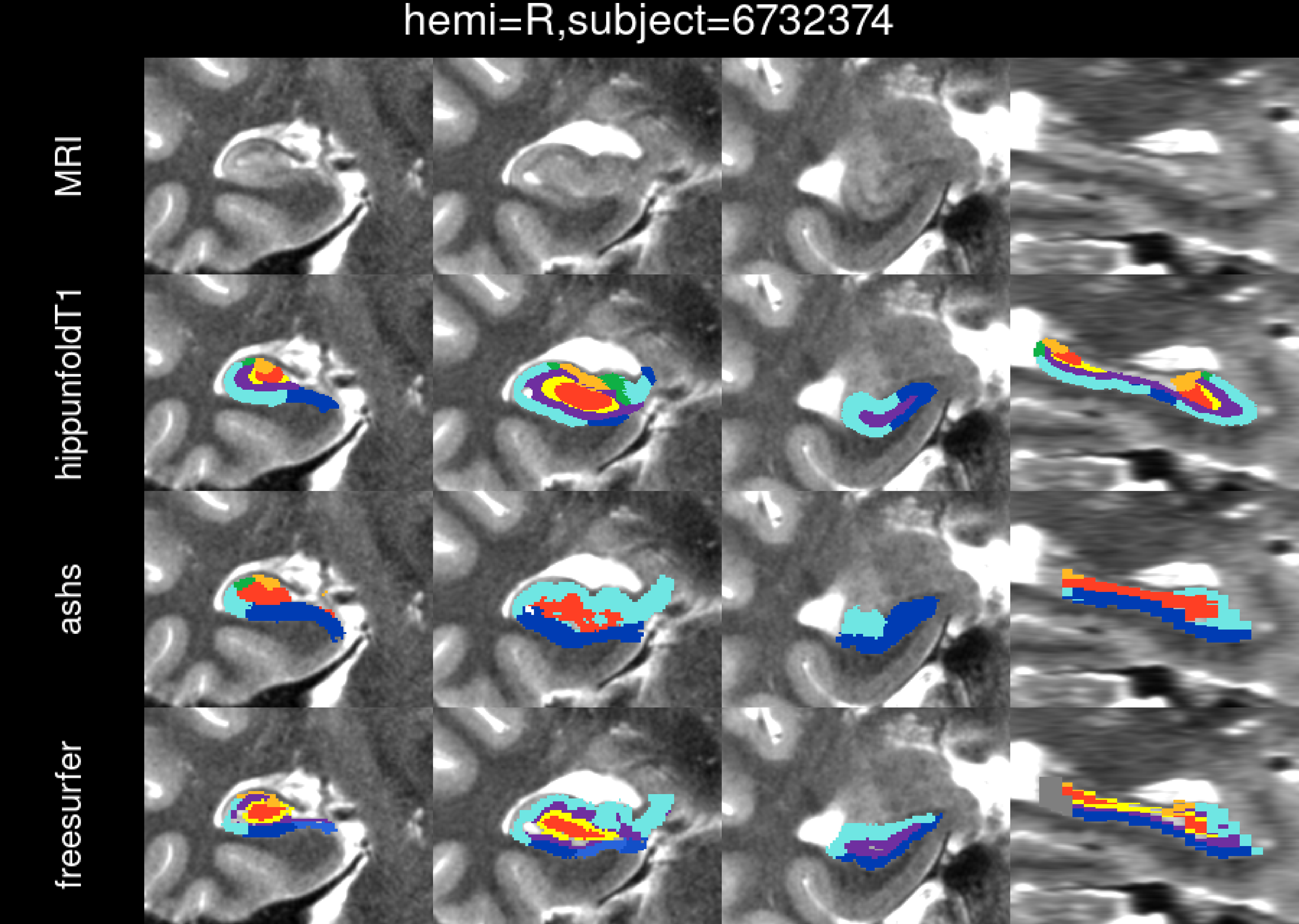
hemi=R,subject=6732374
MRI
hippunfoldT1
ashs
freesurfer

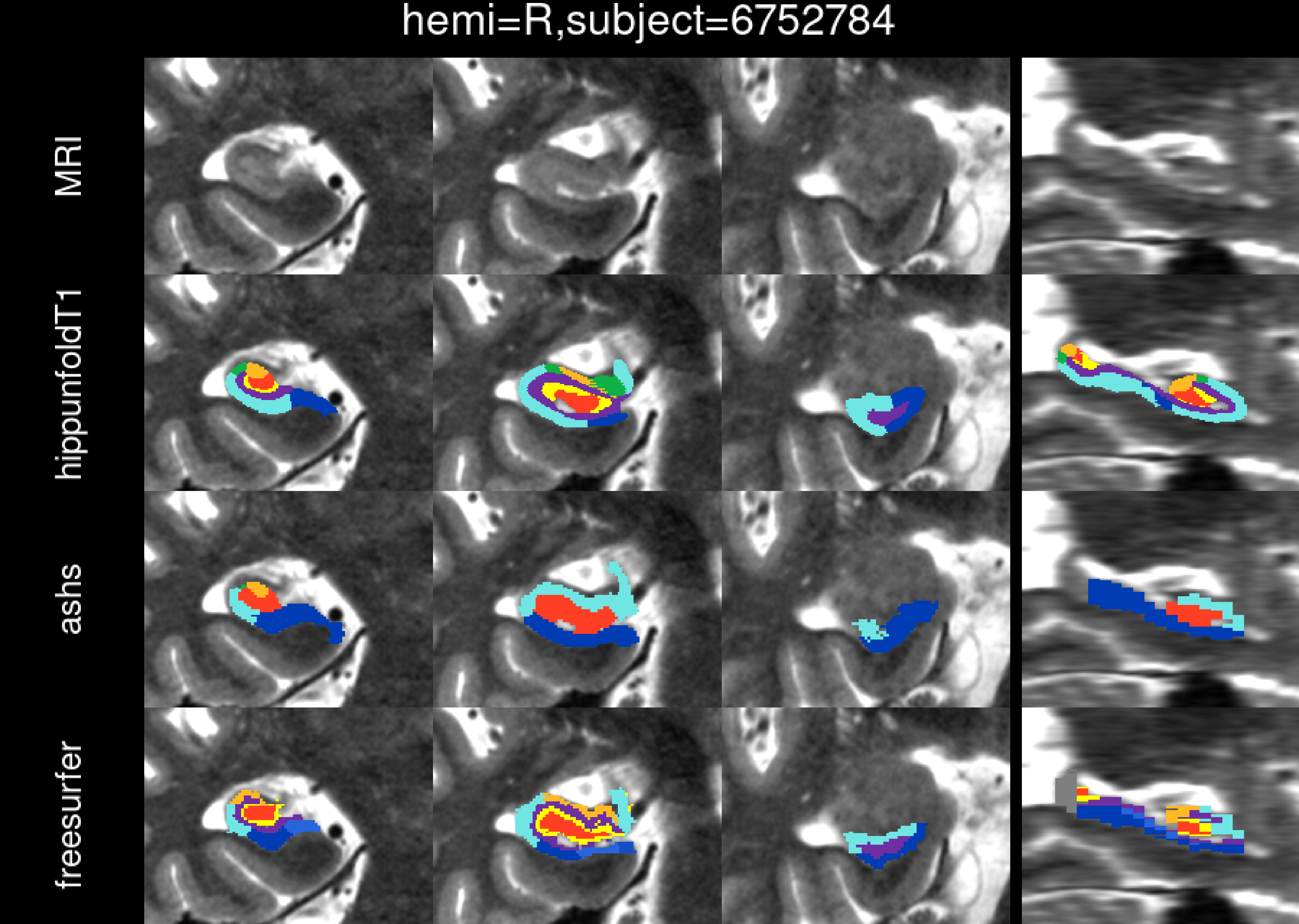

hemi=R,subject=6752784
MRI
hippunfoldT1
ashs
freesurfer

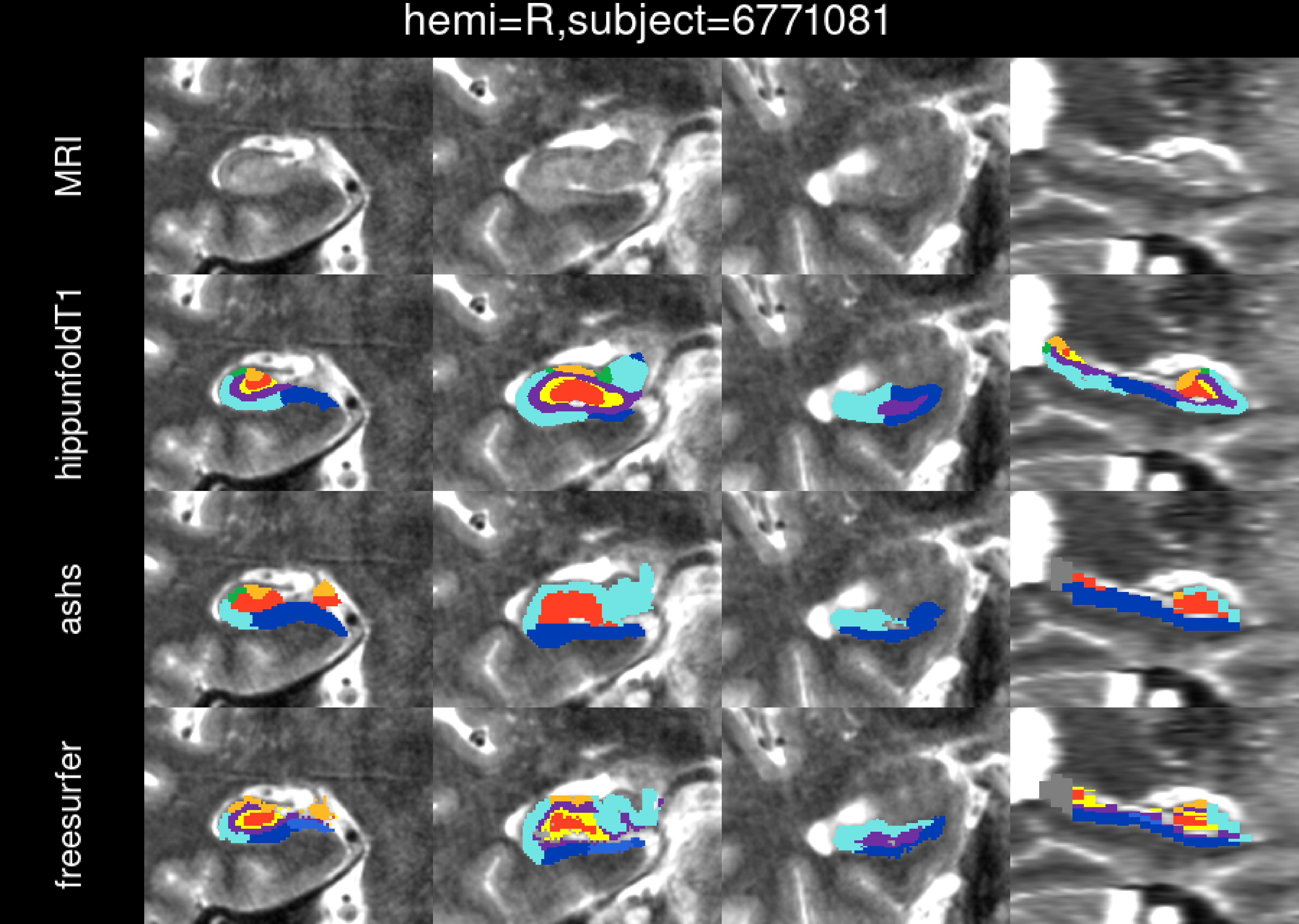
hemi=R,subject=6771081
MRI
hippunfoldT1
ashs
freesurfer

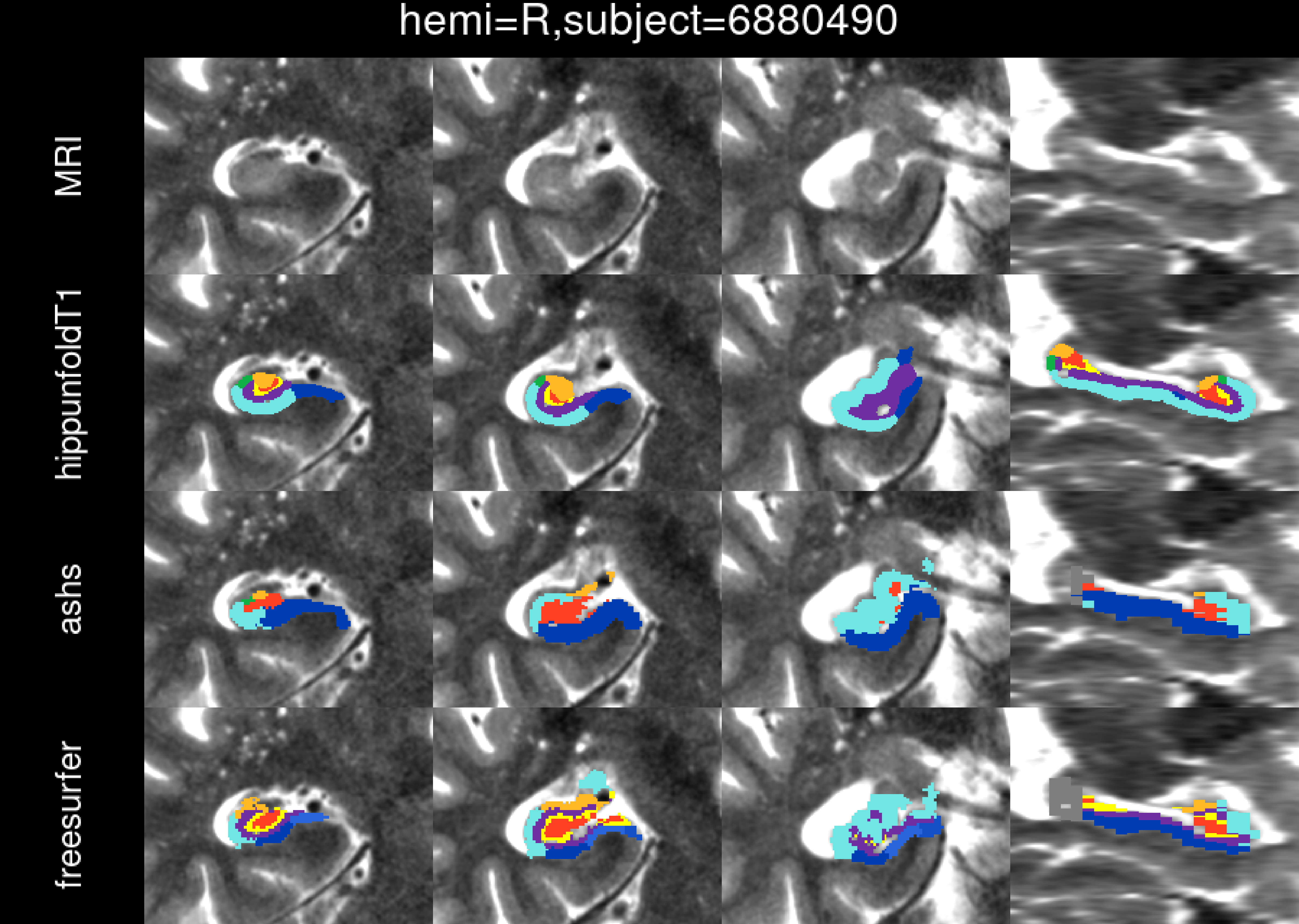

hemi=R,subject=6880490
MRI
hippunfoldT1
ashs
freesurfer

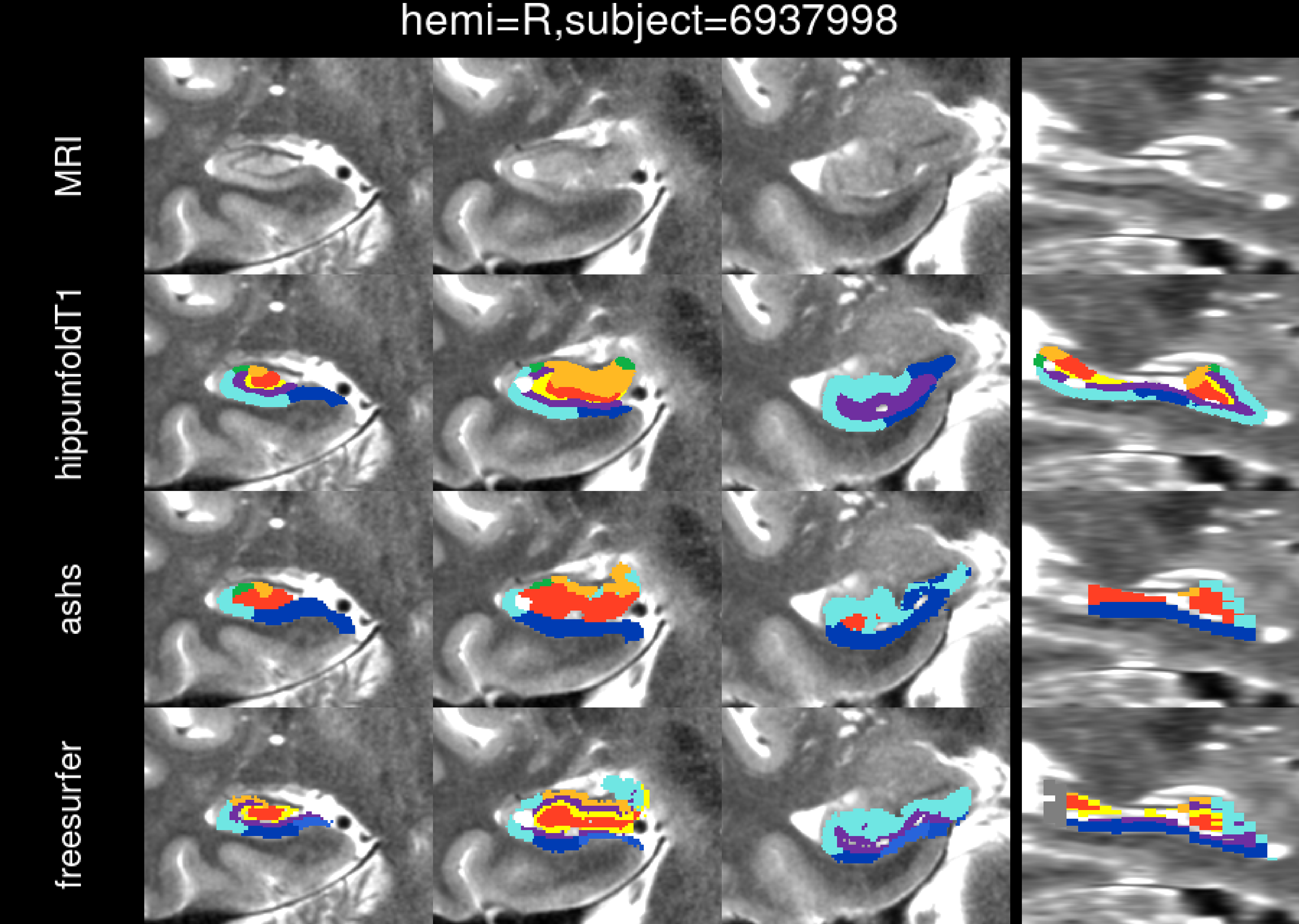
hemi=R,subject=6937998
MRI
hippunfoldT1
ashs
freesurfer

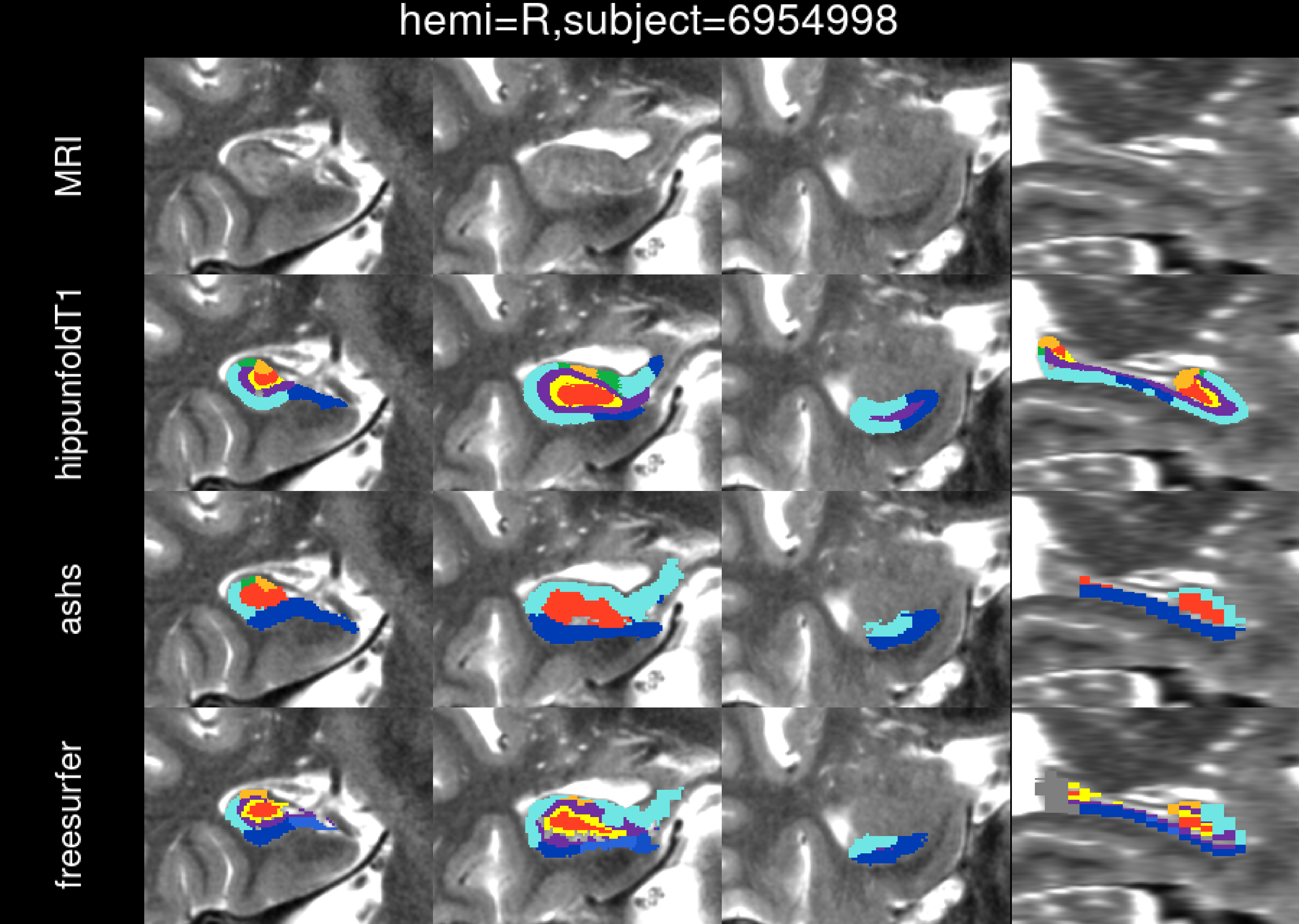

hemi=R,subject=6954998
MRI
hippunfoldT1
ashs
freesurfer

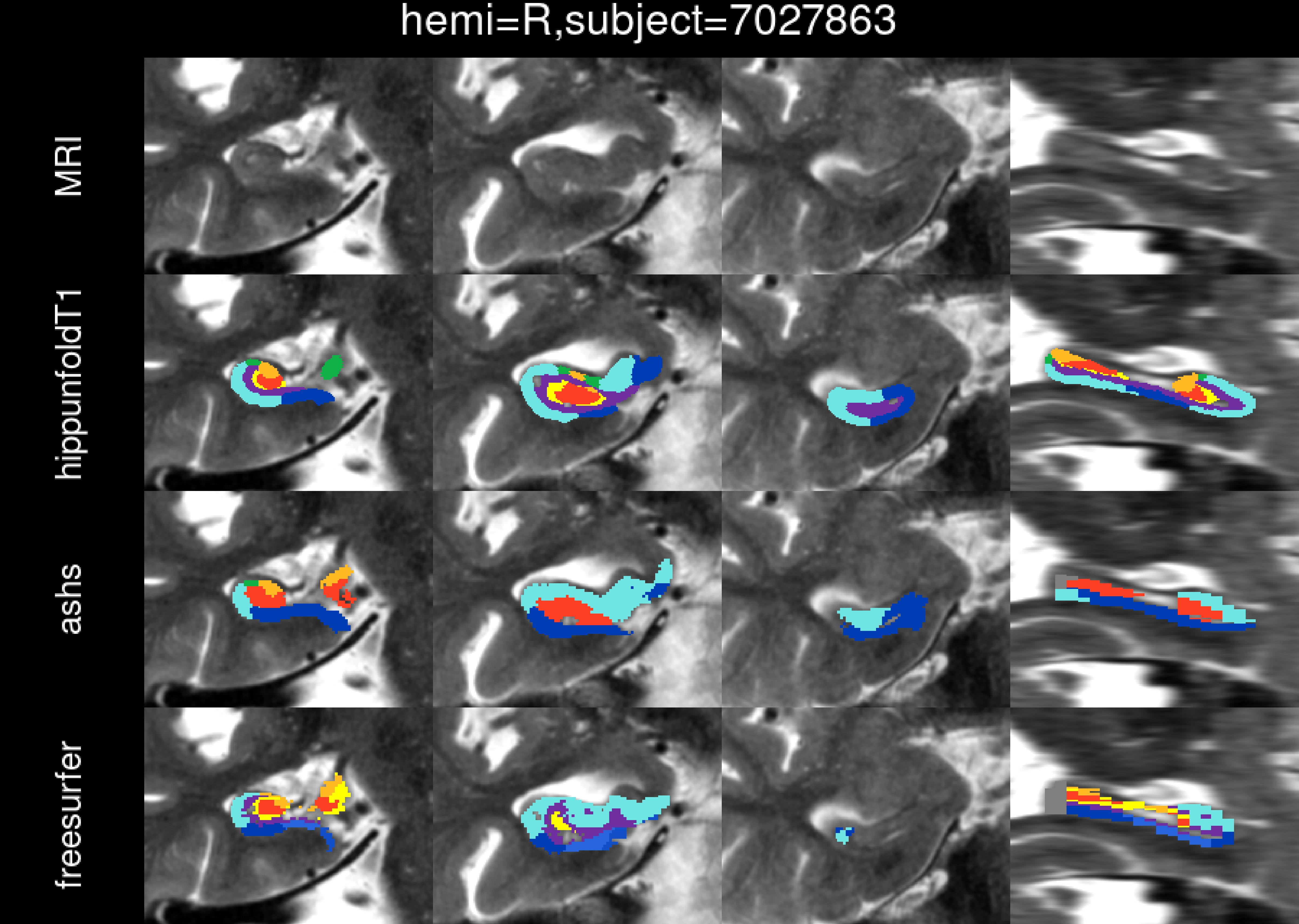
hemi=R,subject=7027863
MRI
hippunfoldT1
ashs
freesurfer

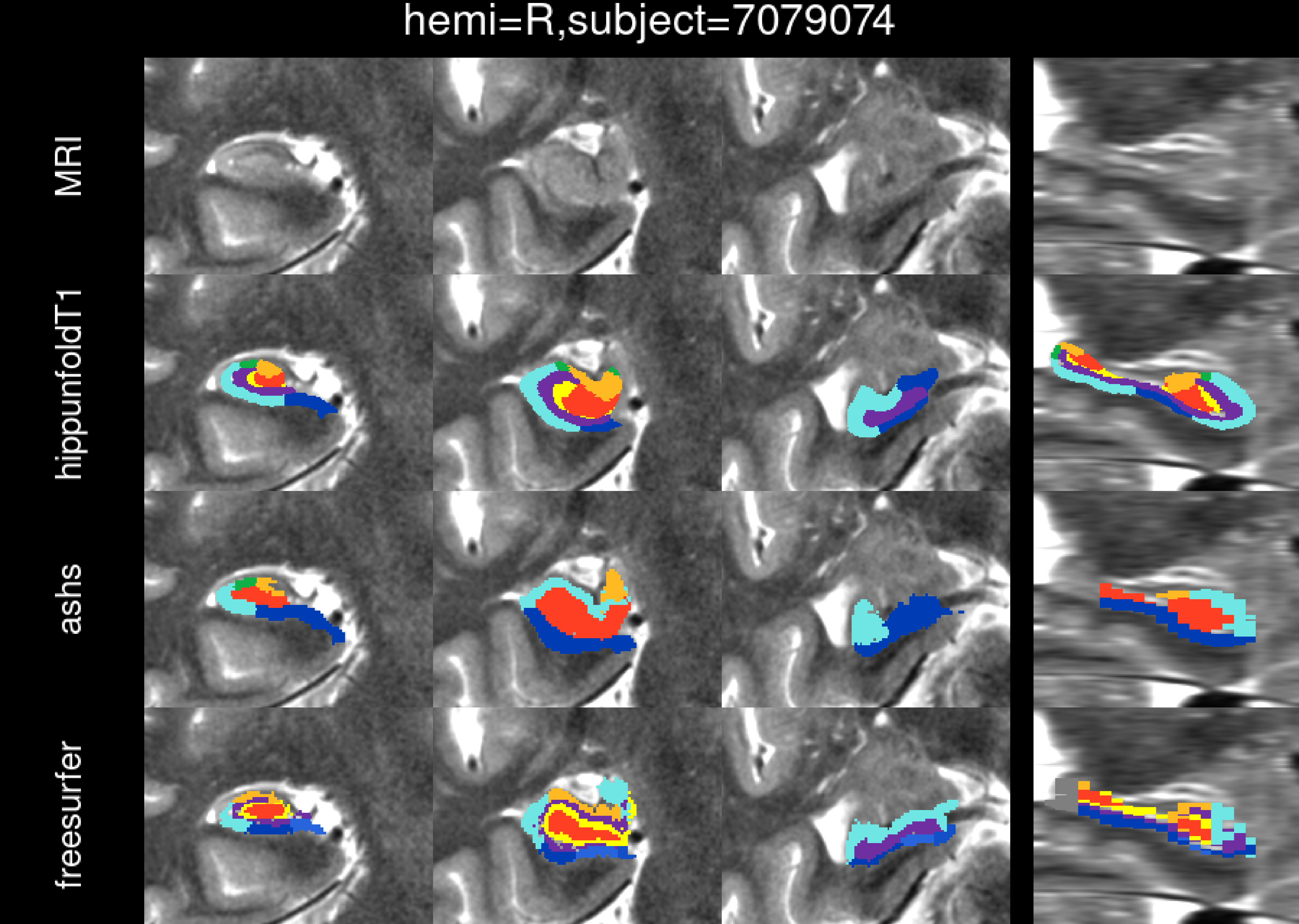
hemi=R,subject=7079074
MRI
hippunfoldT1
ashs
freesurfer

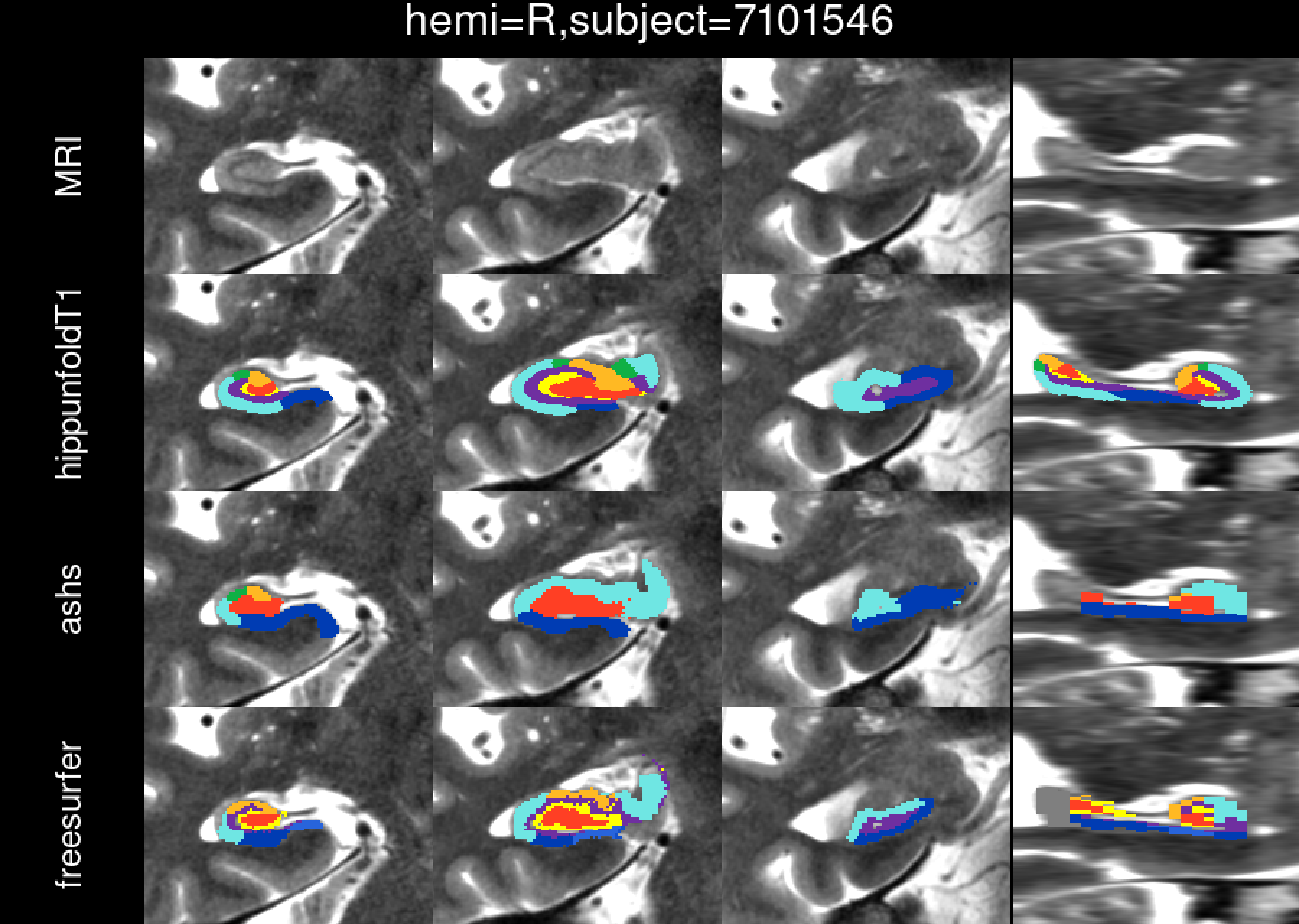
hemi=R,subject=7101546
MRI
hippunfoldT1
ashs
freesurfer

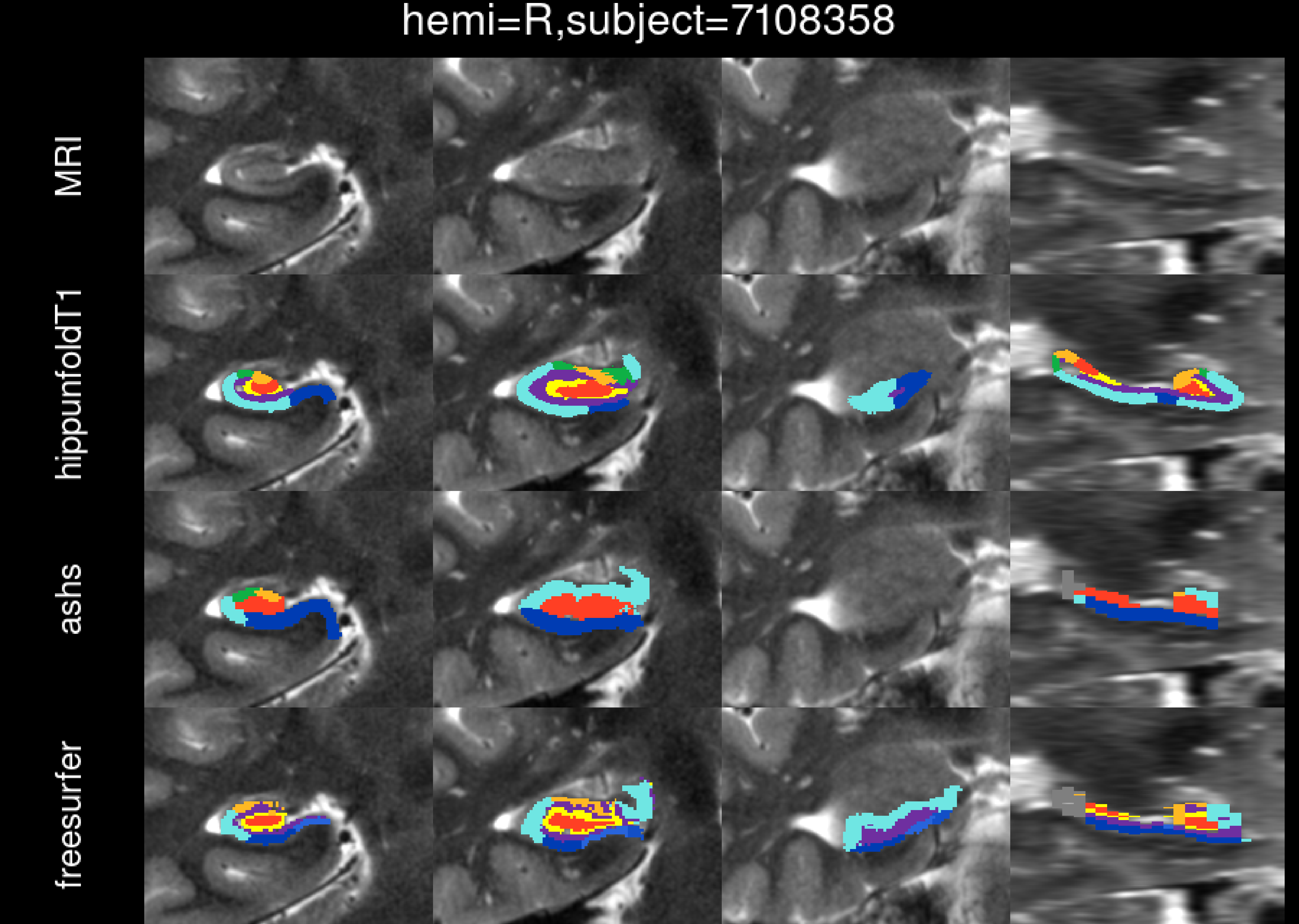
hemi=R,subject=7108358
MRI
hippunfoldT1
ashs
freesurfer

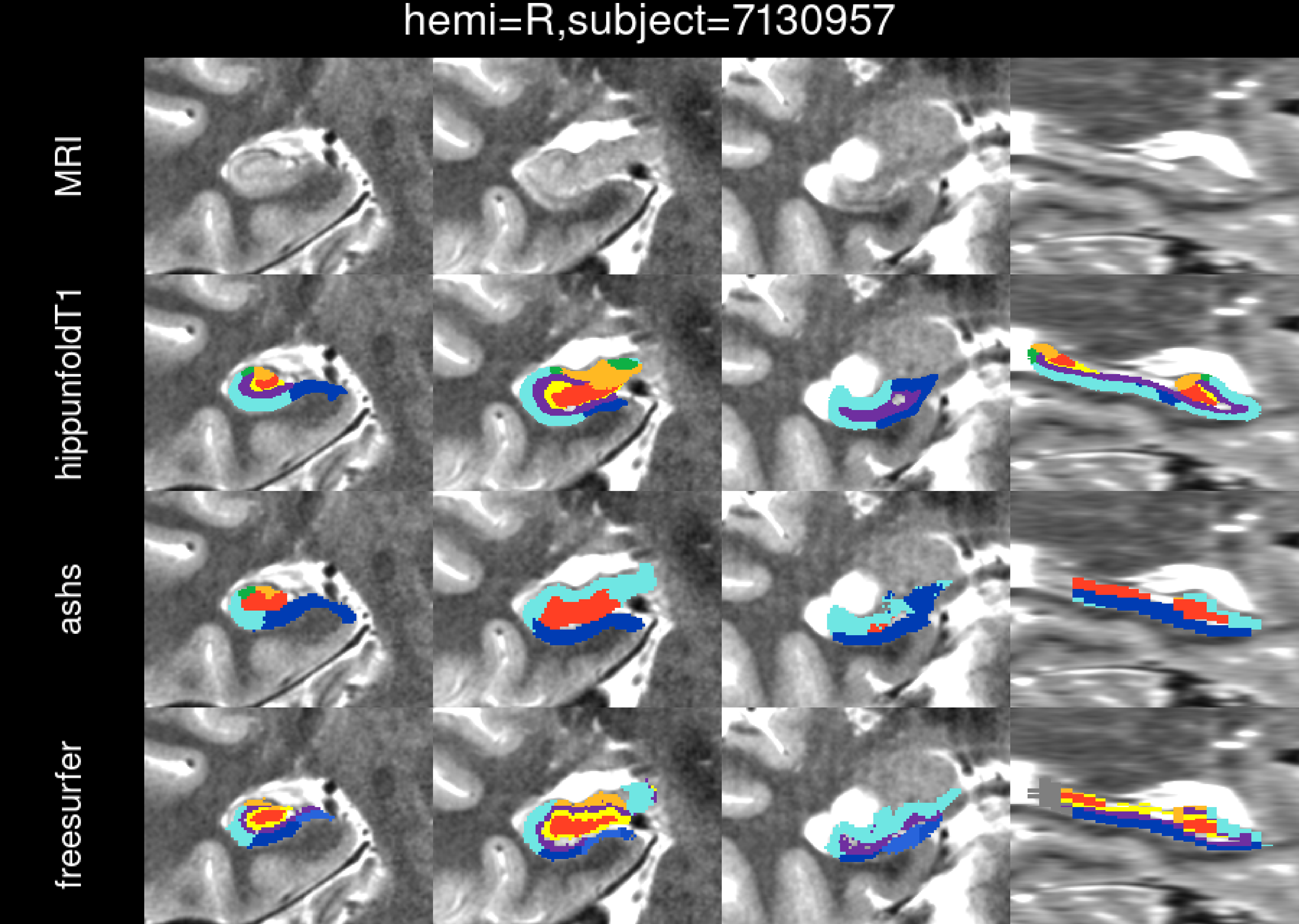
hemi=R,subject=7130957
MRI
hippunfoldT1
ashs
freesurfer

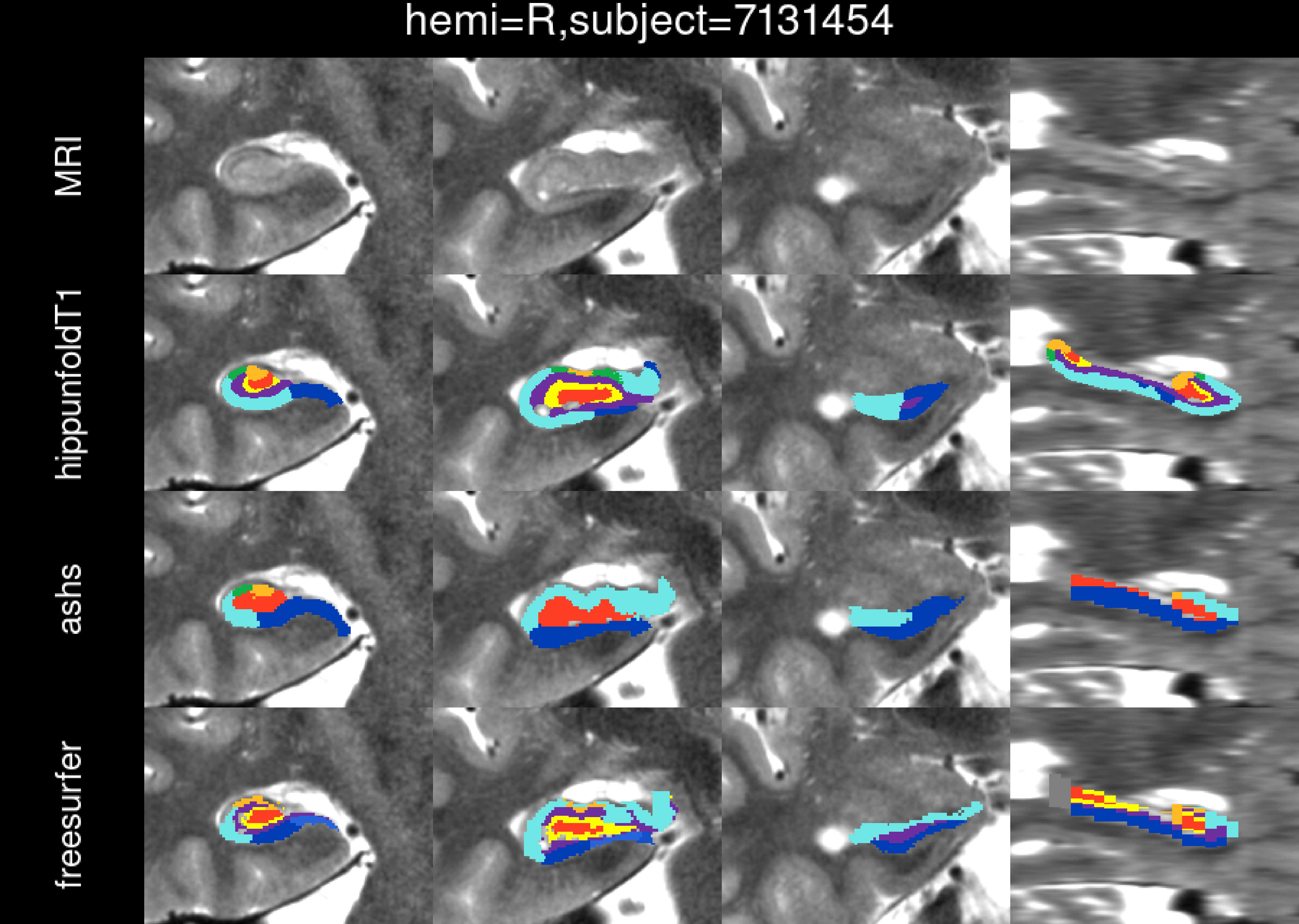
hemi=R,subject=7131454
MRI
hippunfoldT1
ashs
freesurfer

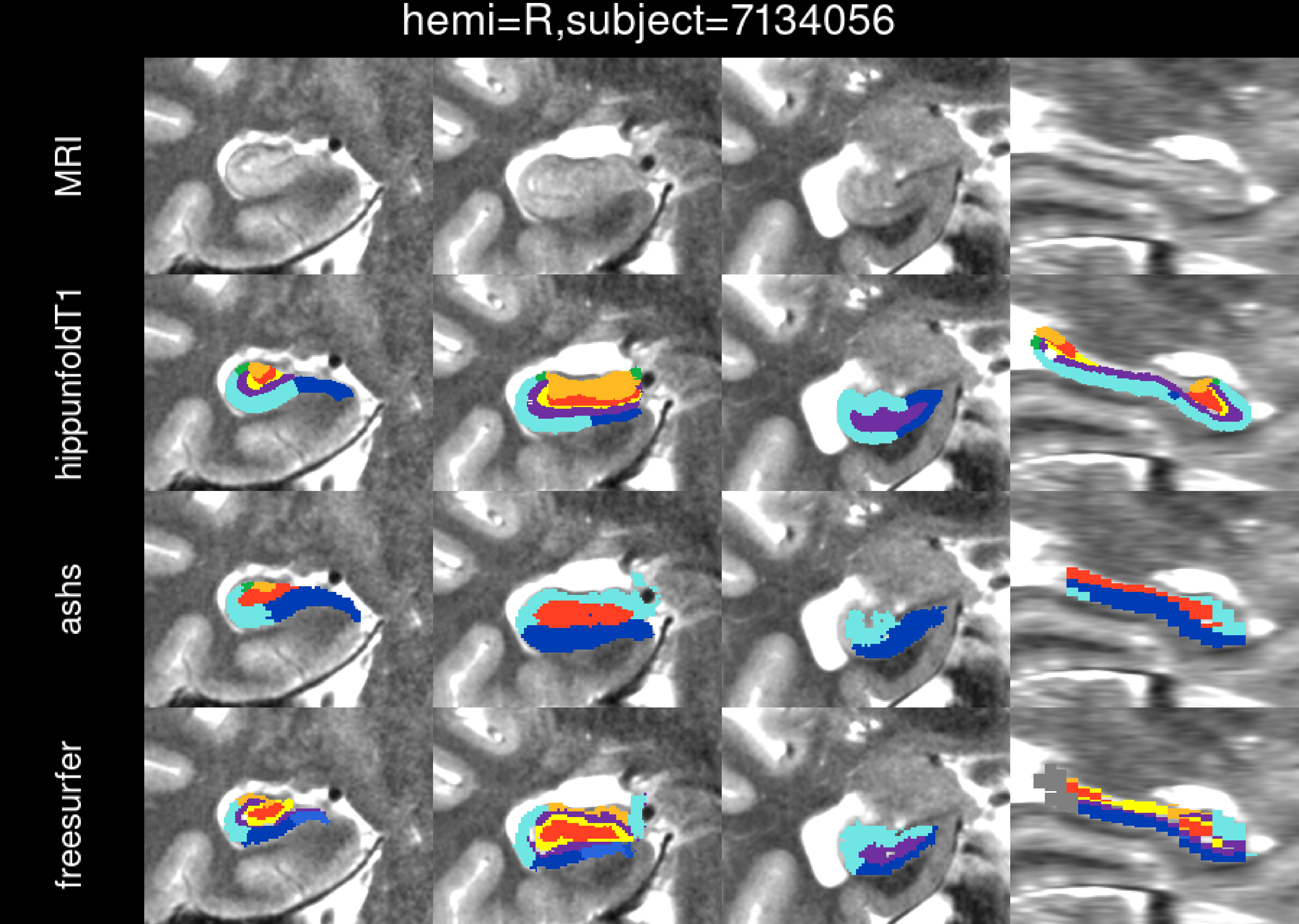
hemi=R,subject=7134056
MRI
hippunfoldT1
ashs
freesurfer

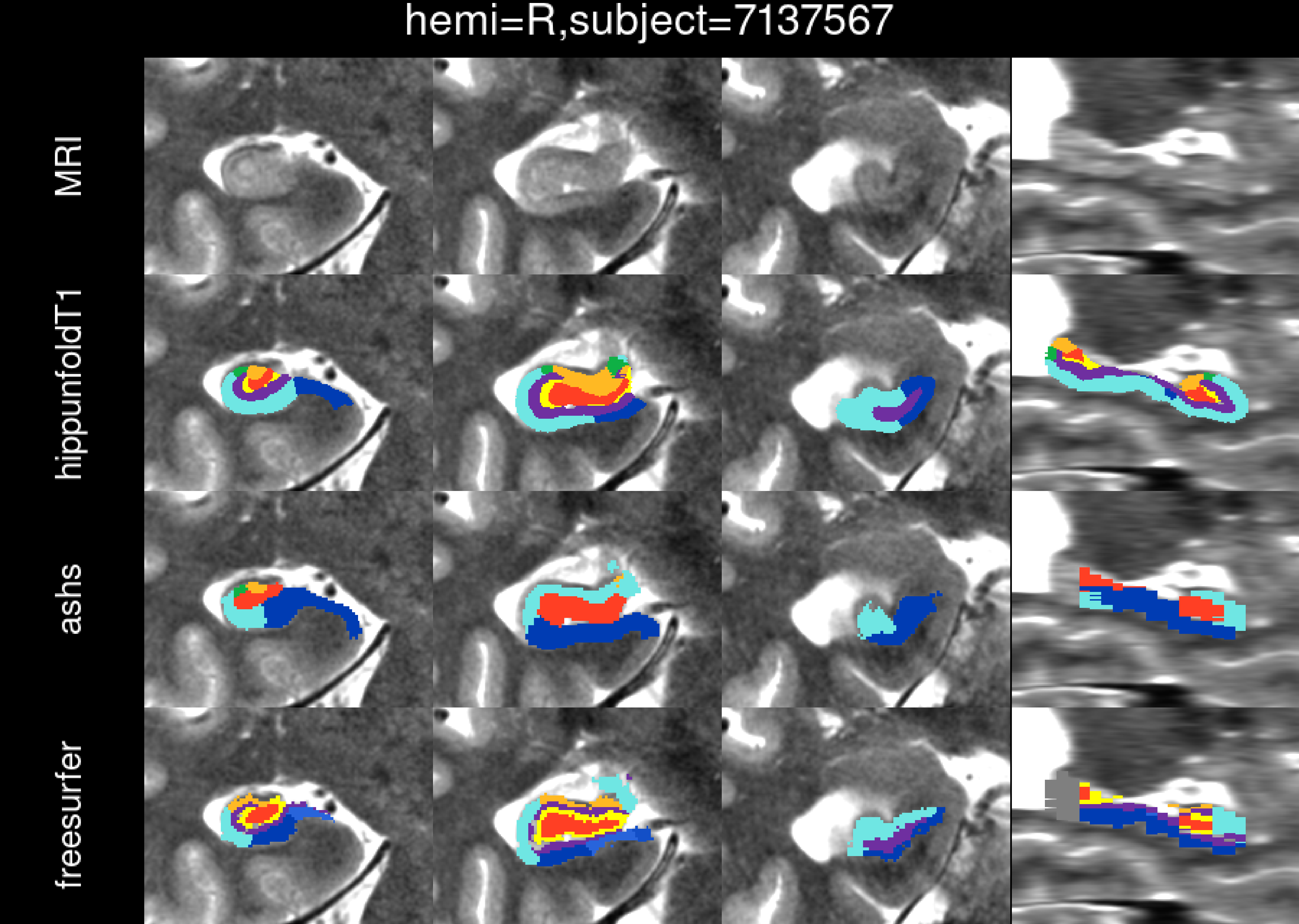
hemi=R,subject=7137567
MRI
hippunfoldT1
ashs
freesurfer

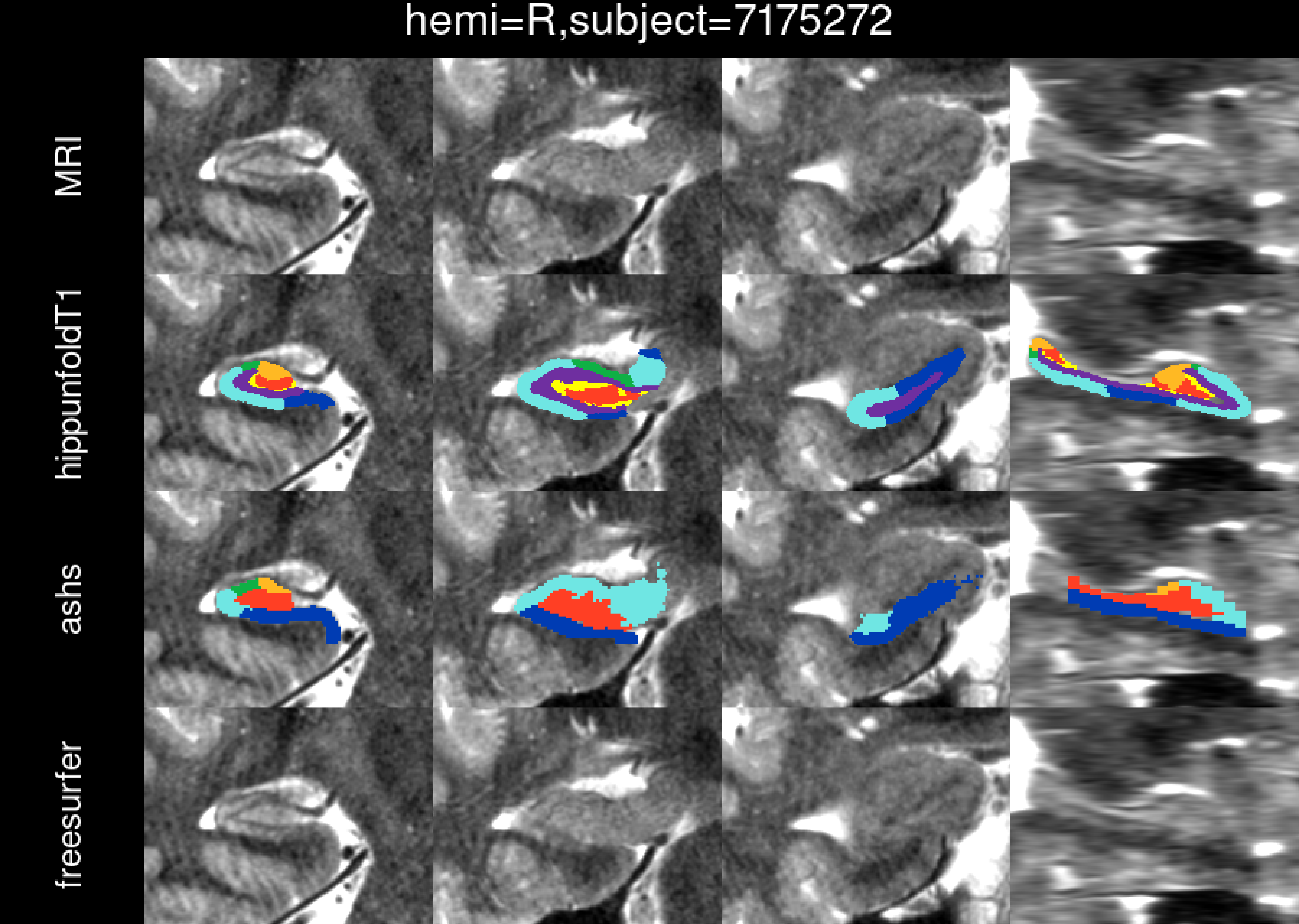
hemi=R,subject=7175272
MRI
hippunfoldT1
ashs
freesurfer

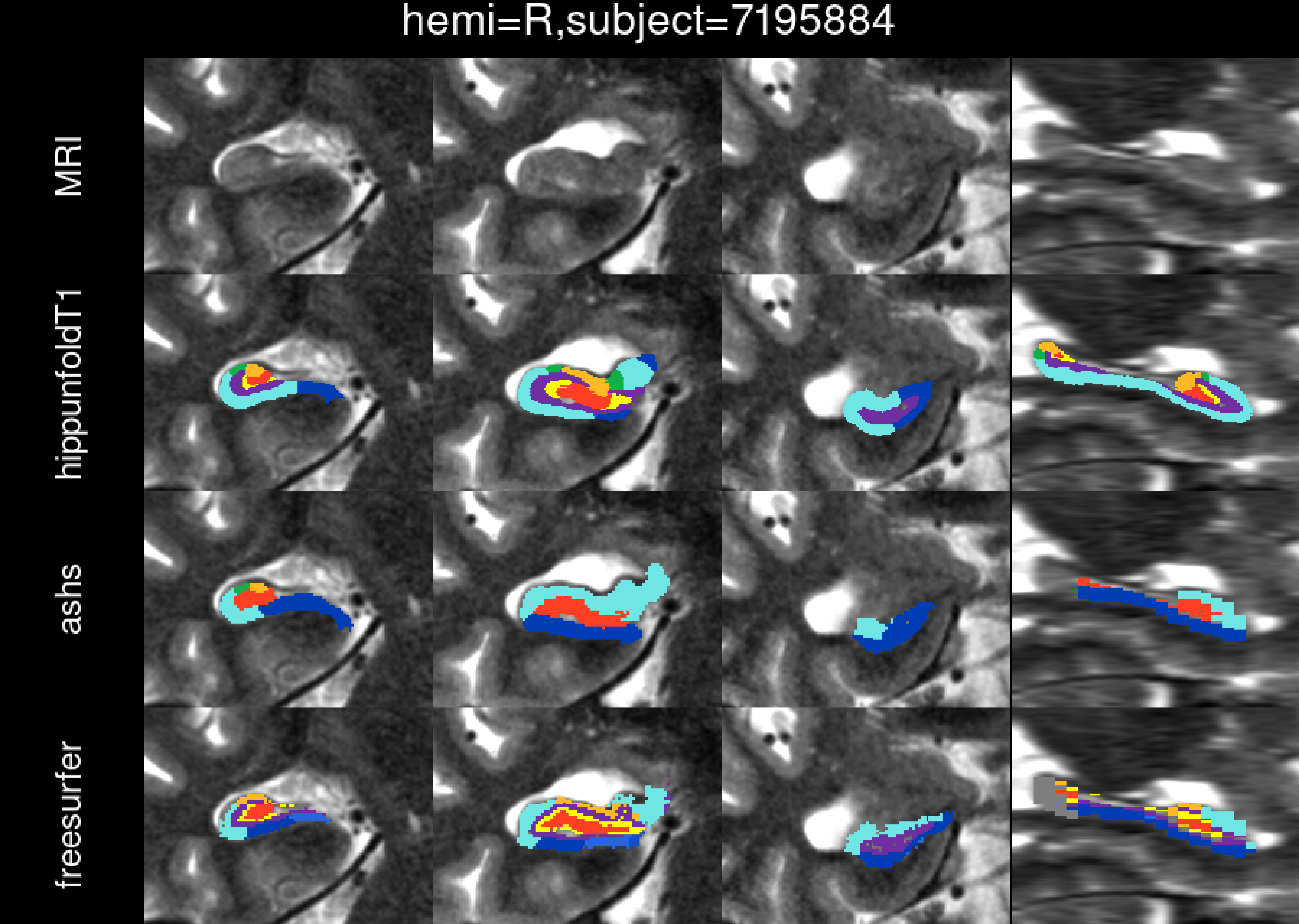

hemi=R,subject=7195884
MRI
hippunfoldT1
ashs
freesurfer

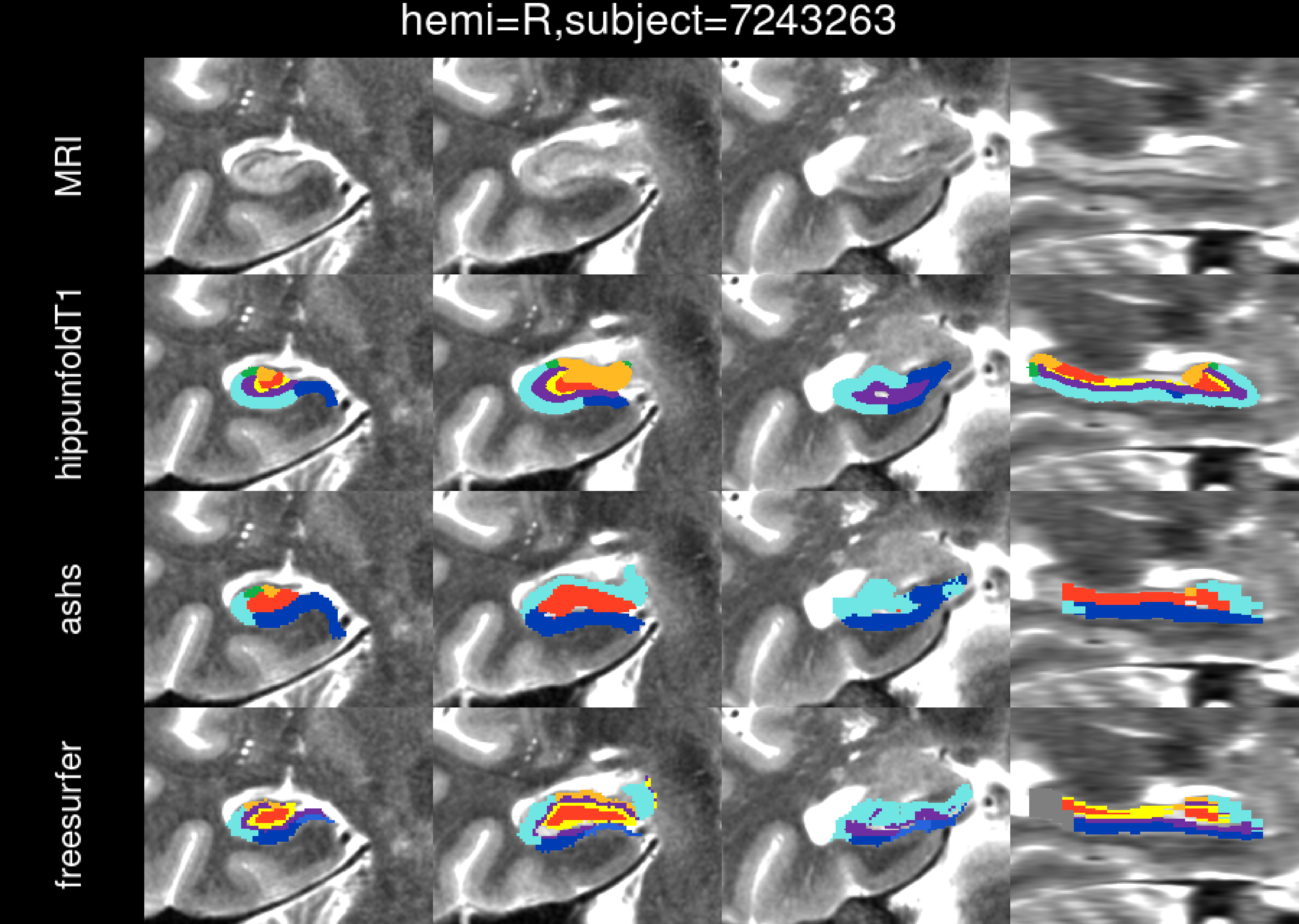

hemi=R,subject=7243263
MRI
hippunfoldT1
ashs
freesurfer

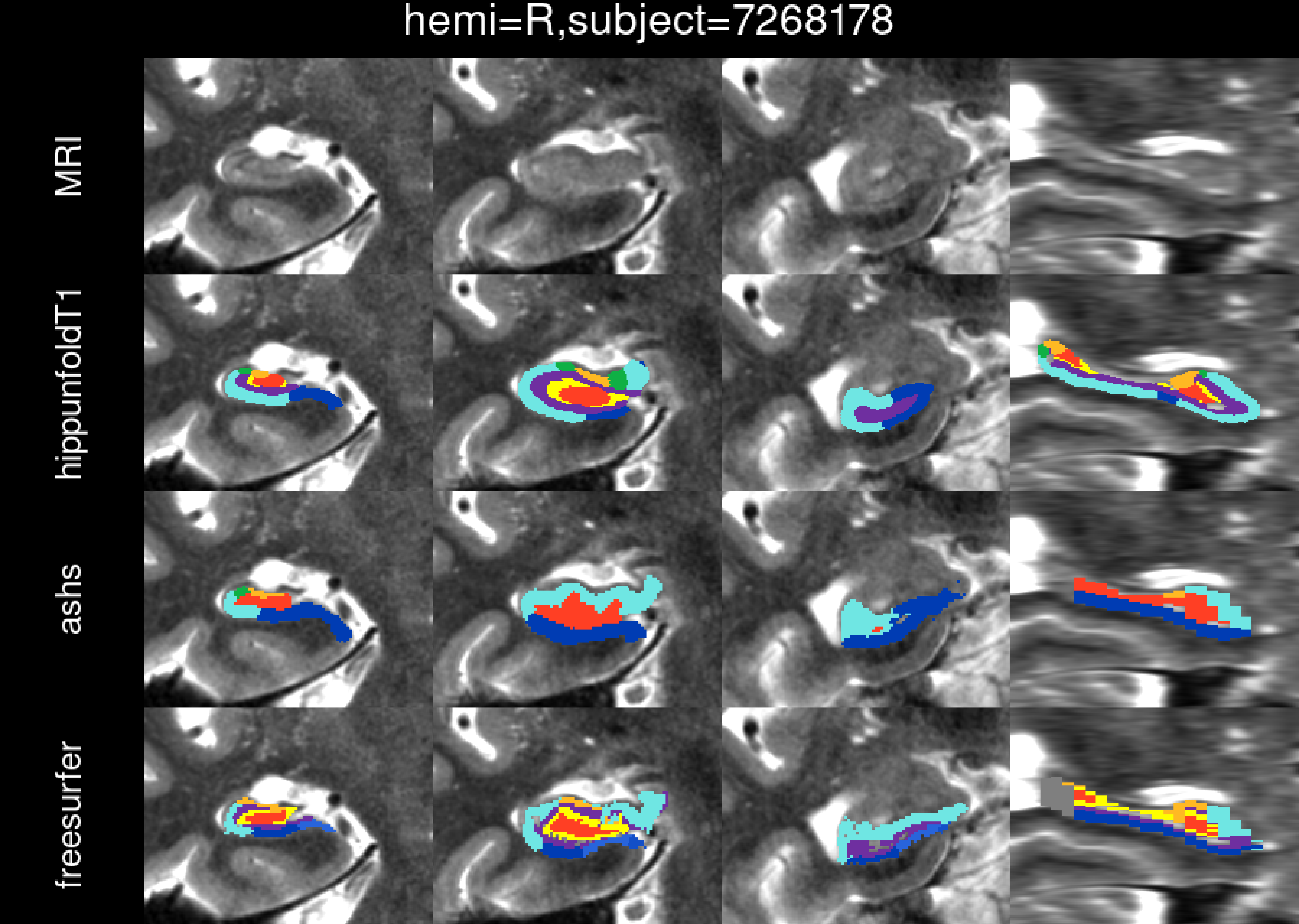
hemi=R,subject=7268178
MRI
hippunfoldT1
ashs
freesurfer

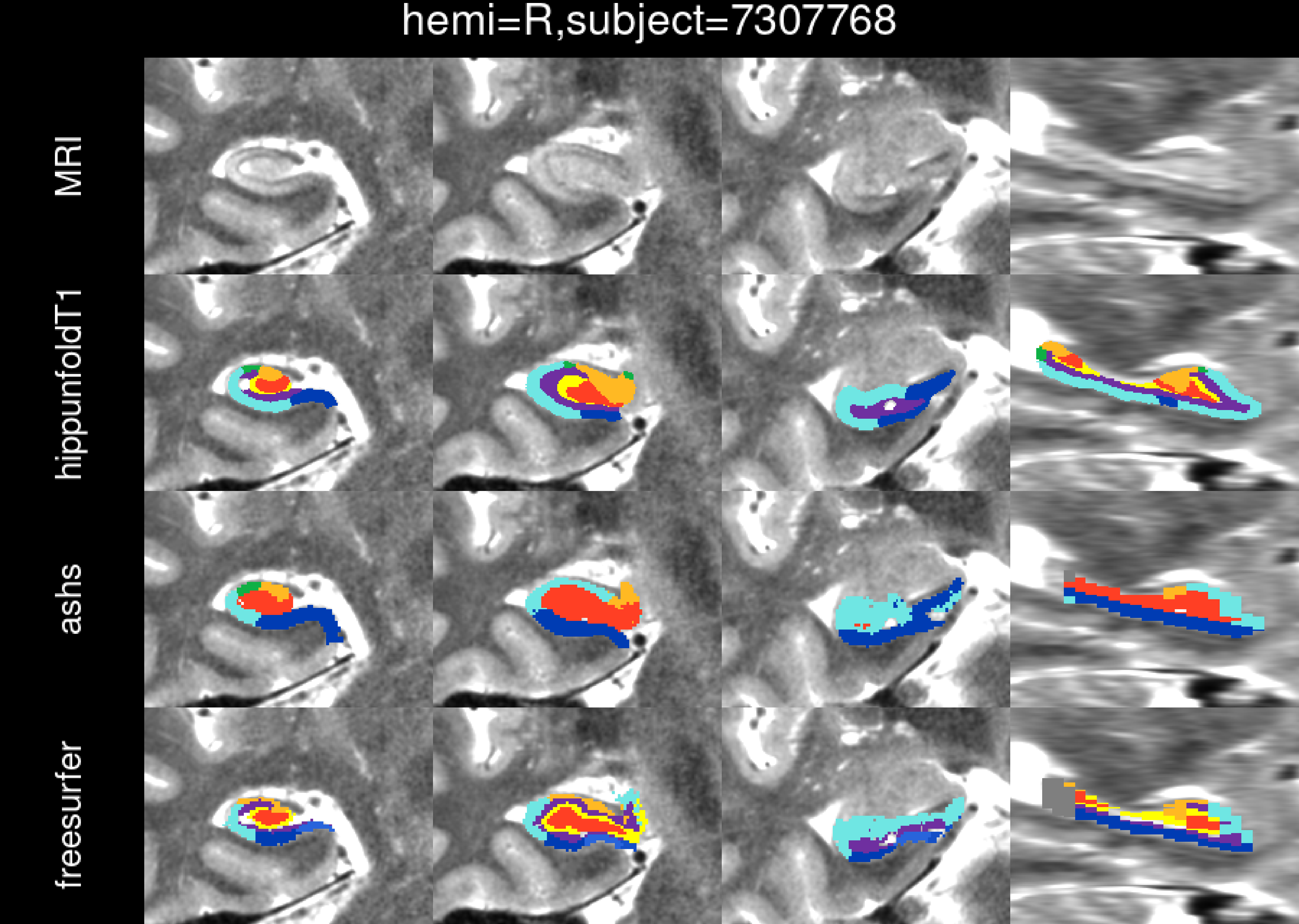

hemi=R,subject=7307768
MRI
hippunfoldT1
ashs
freesurfer

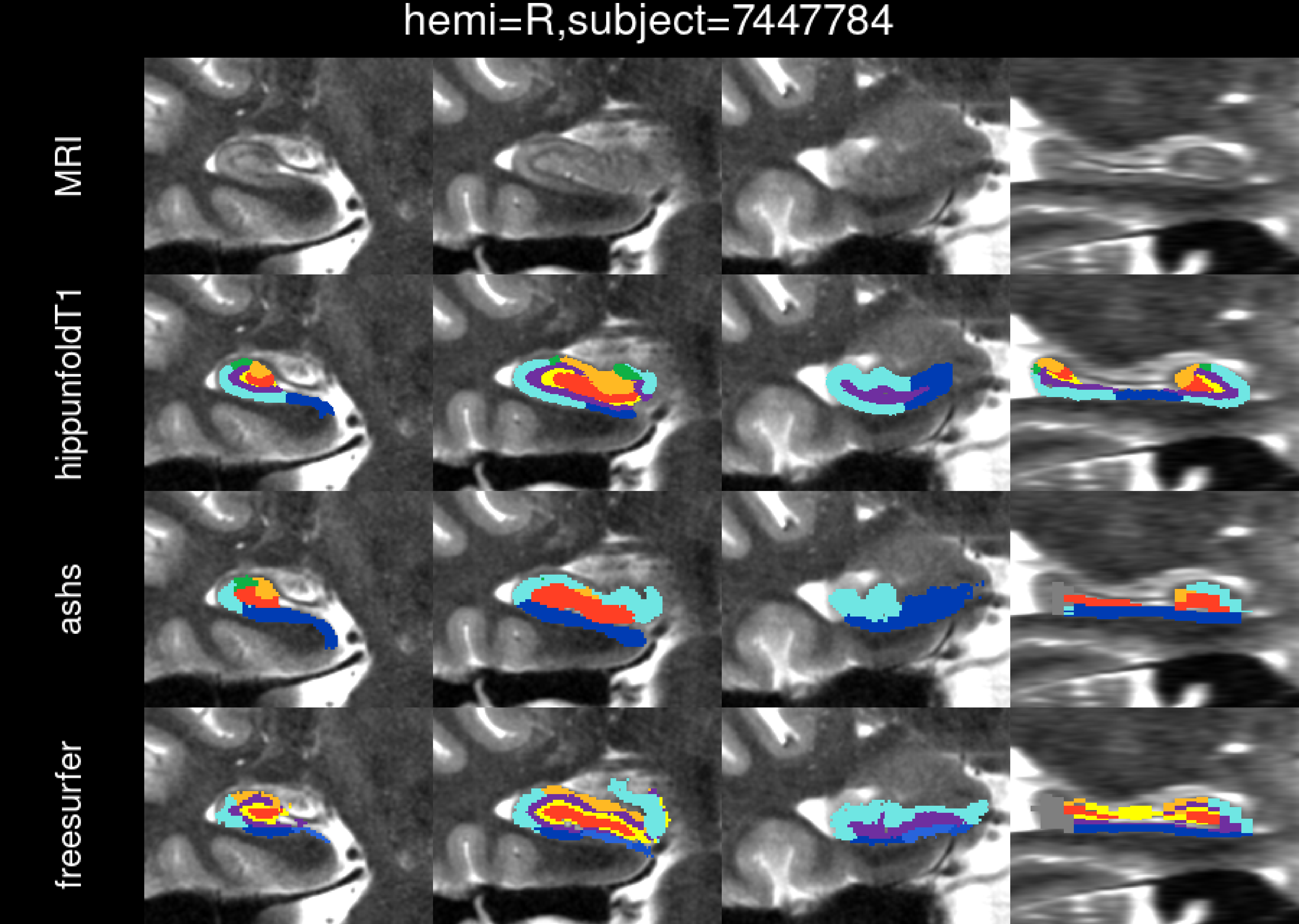
hemi=R,subject=7447784
MRI
hippunfoldT1
ashs
freesurfer

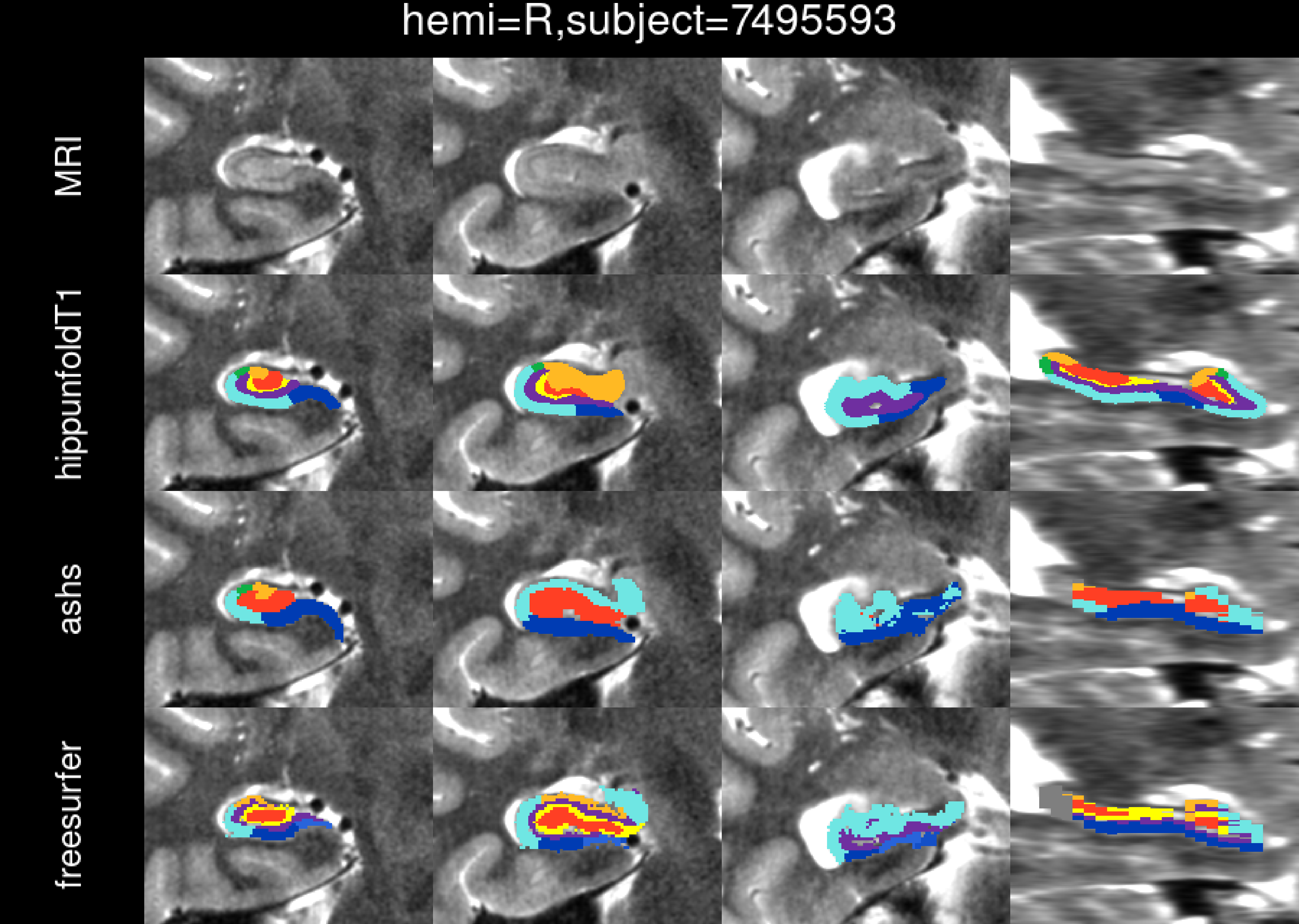
hemi=R,subject=7495593
MRI
hippunfoldT1
ashs
freesurfer

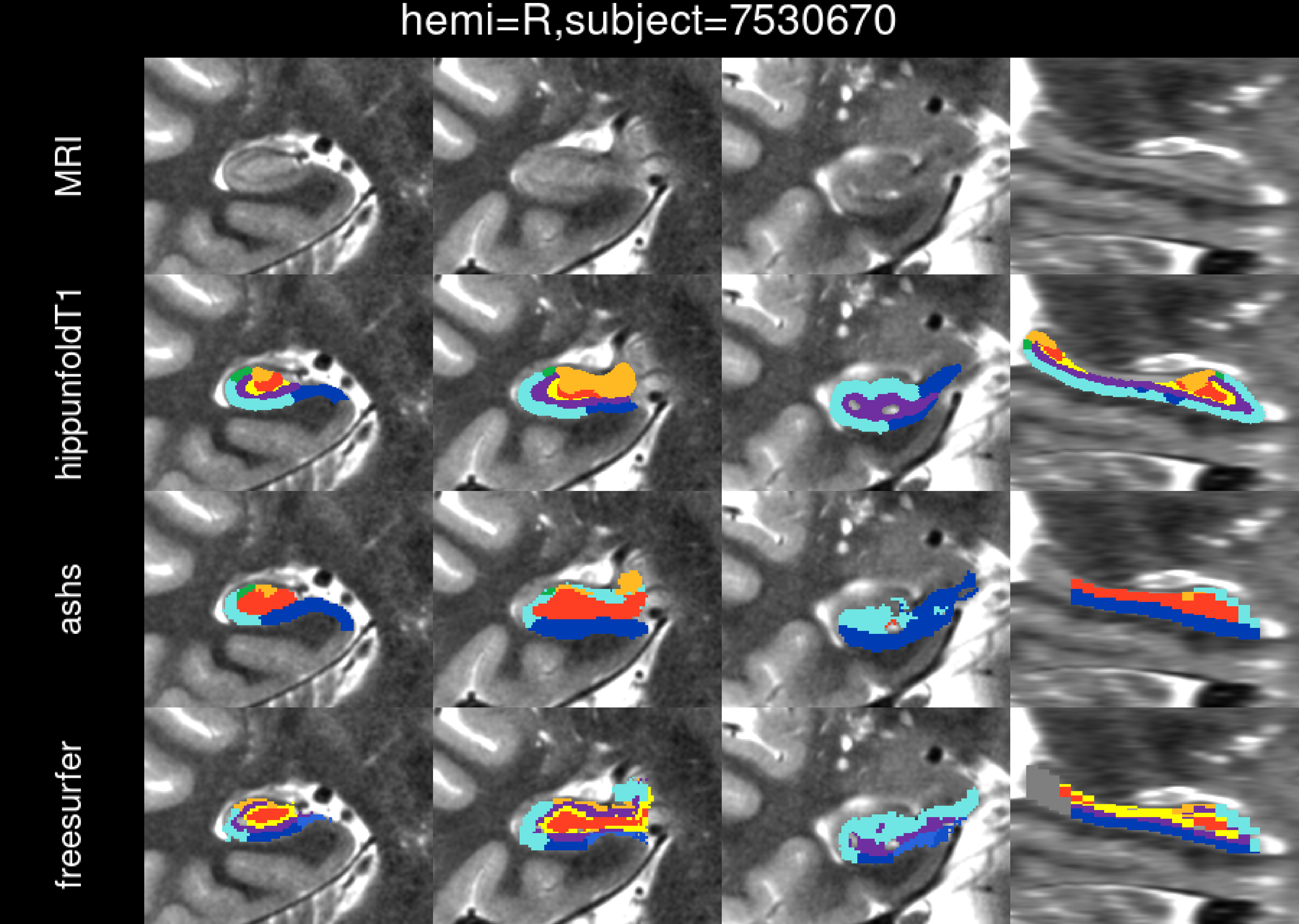
hemi=R,subject=7530670
MRI
hippunfoldT1
ashs
freesurfer

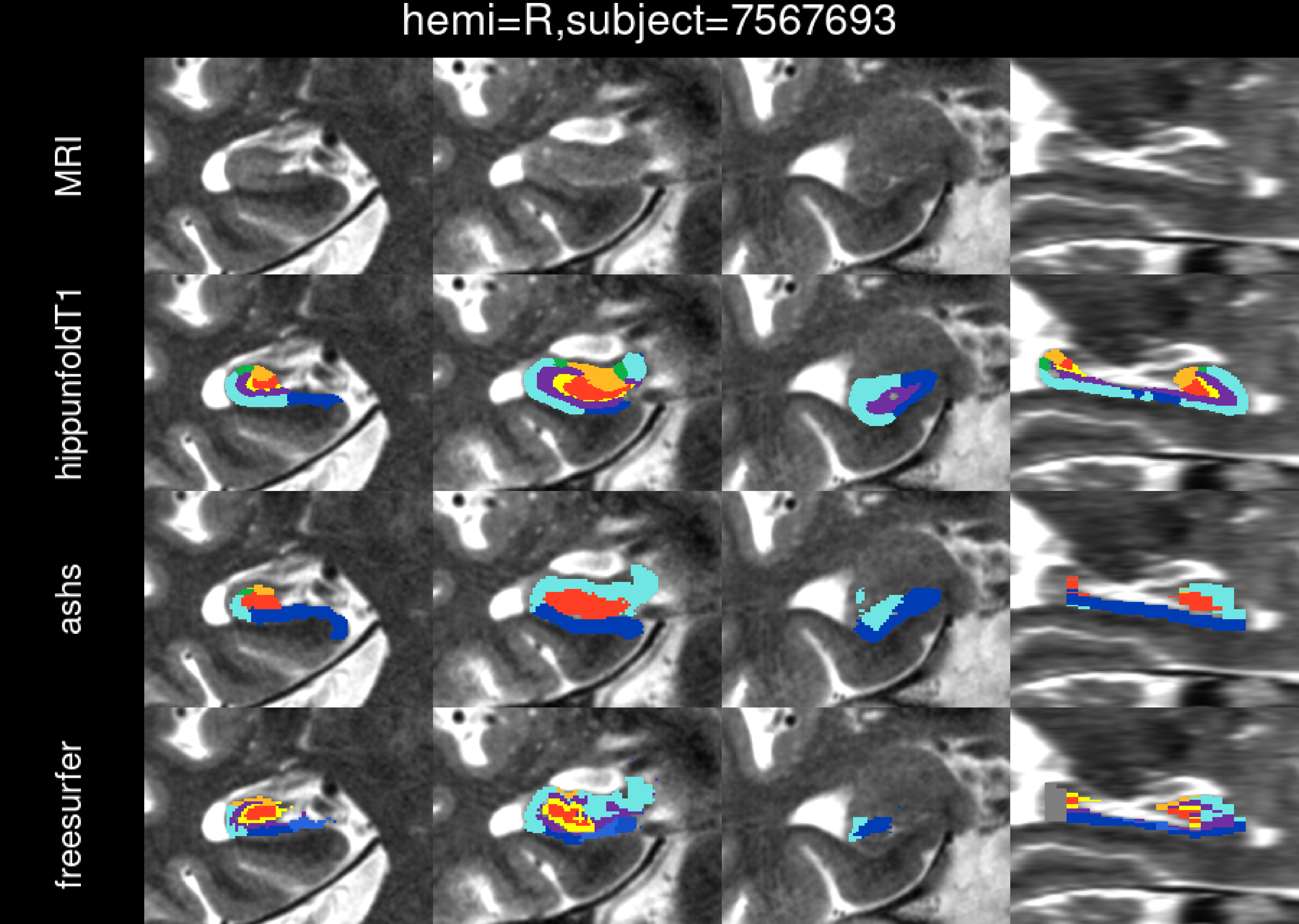
hemi=R,subject=7567693
MRI
hippunfoldT1
ashs
freesurfer

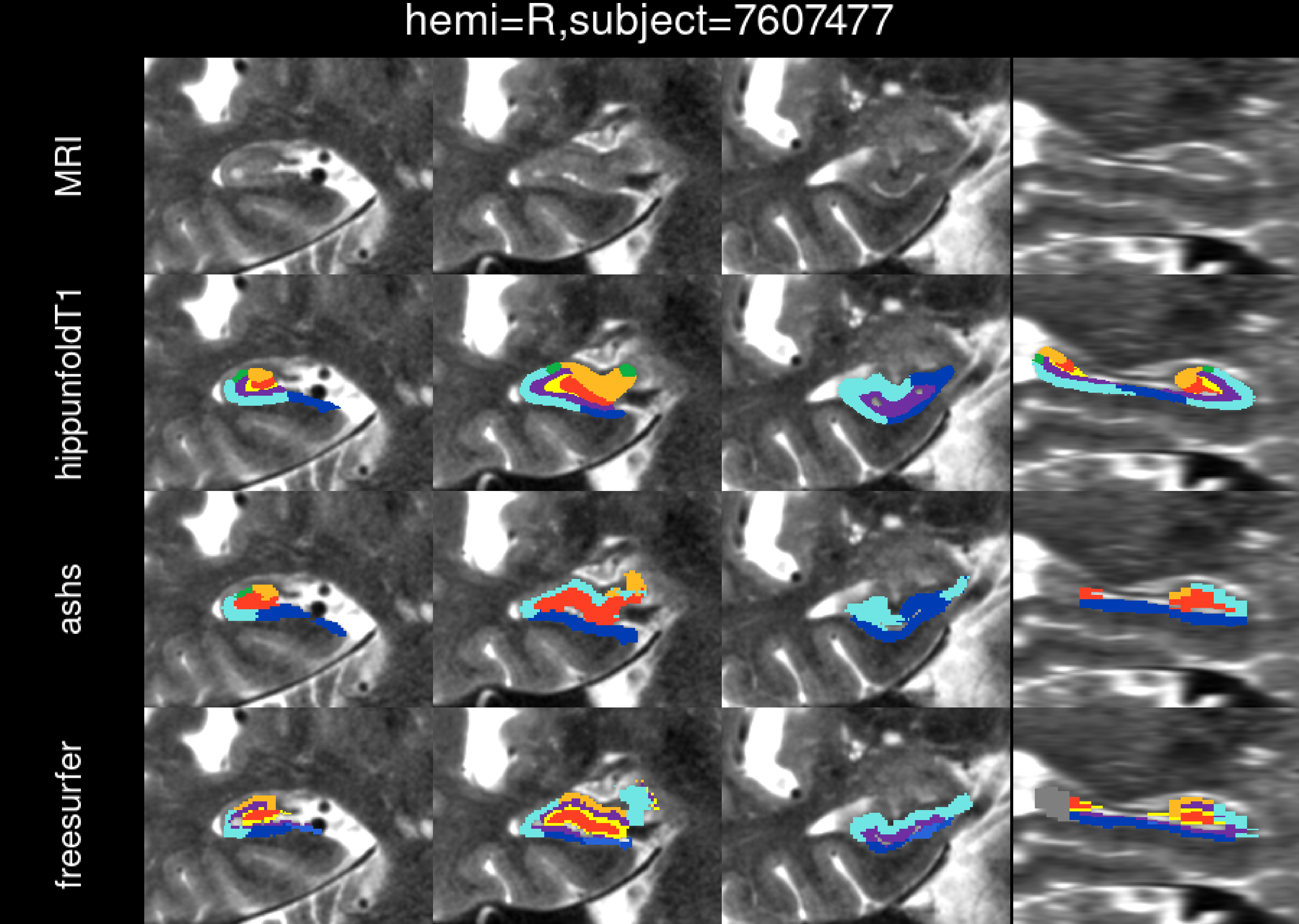
hemi=R,subject=7607477
MRI
hippunfoldT1
ashs
freesurfer

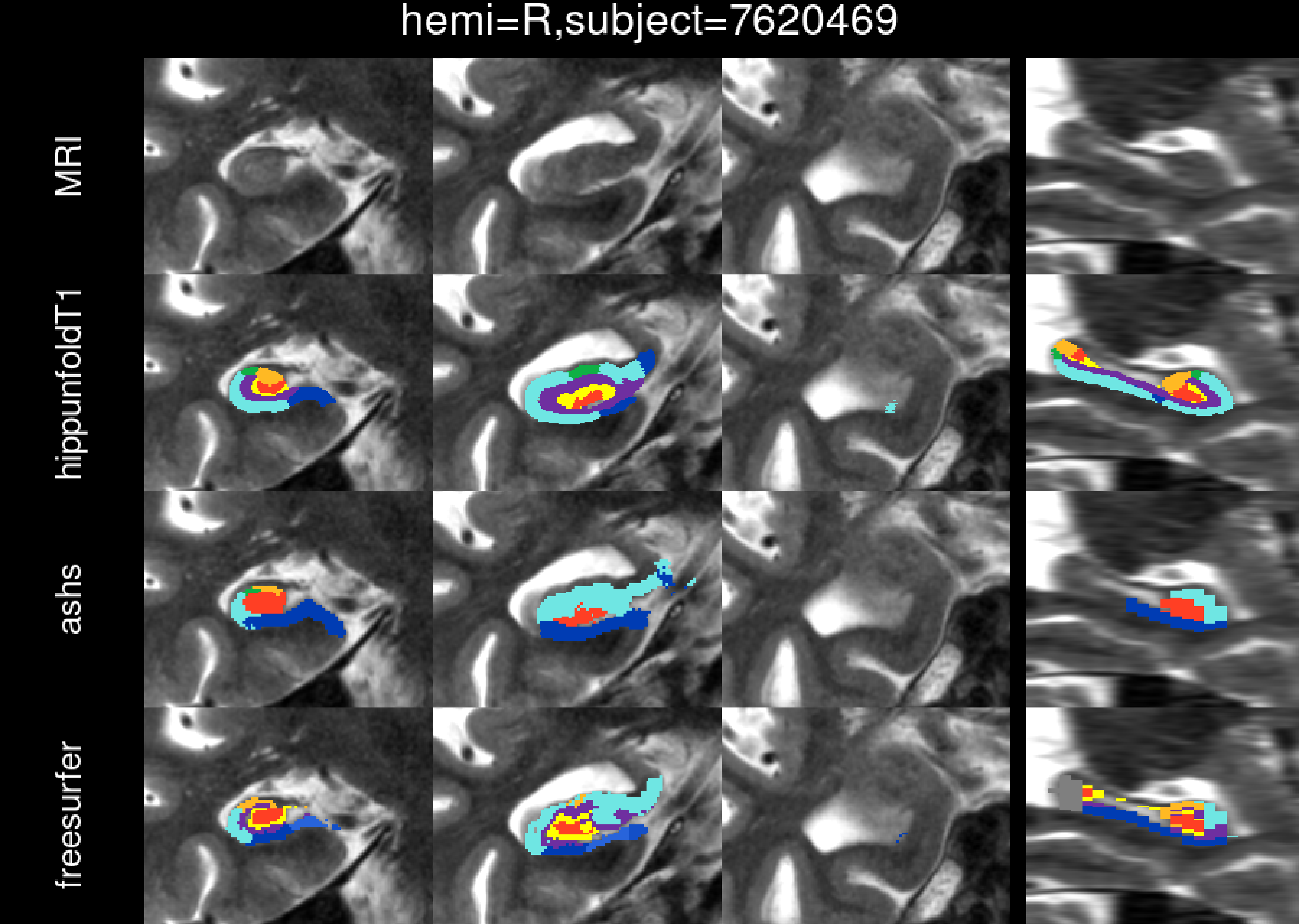
hemi=R,subject=7620469
MRI
hippunfoldT1
ashs
freesurfer

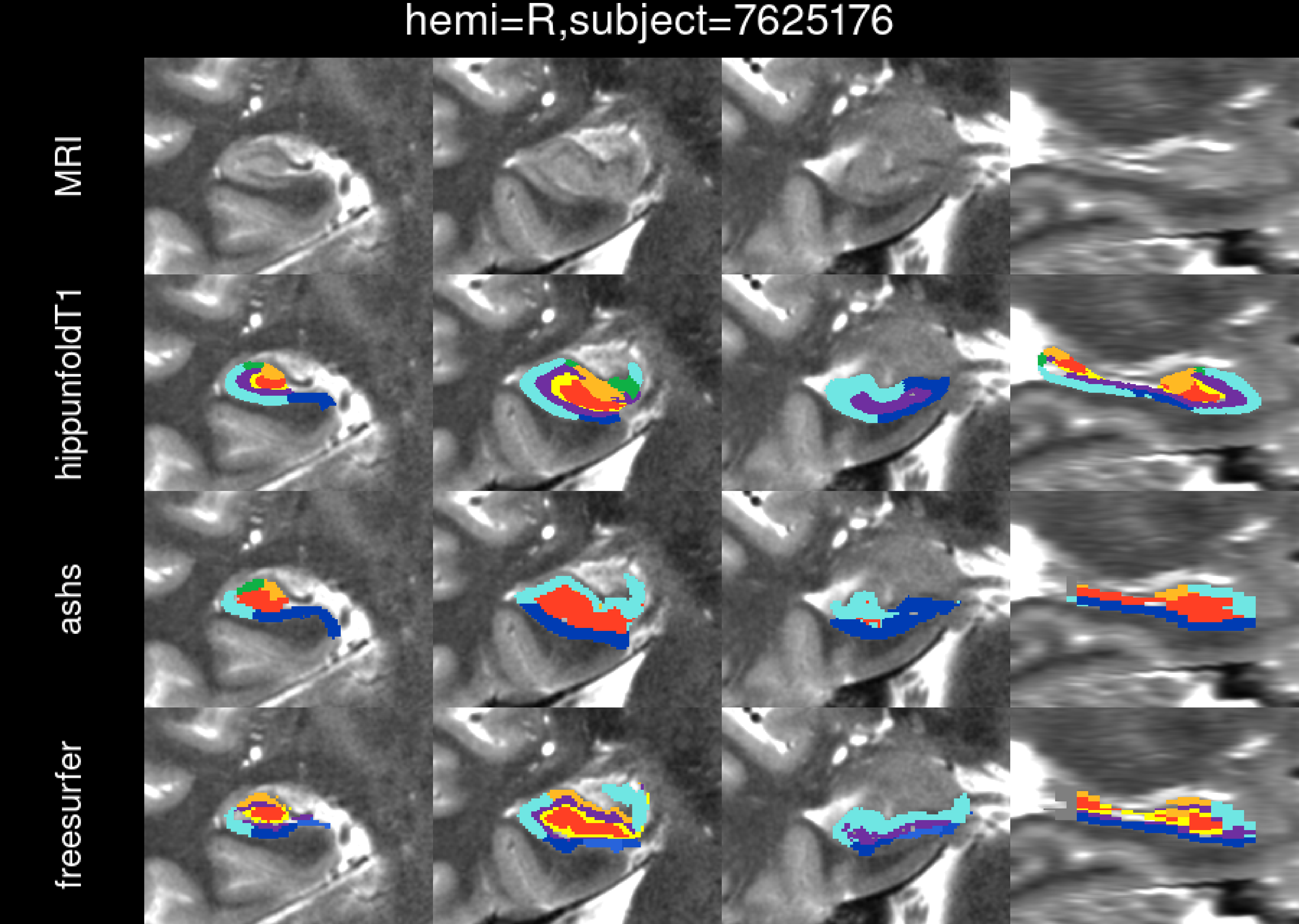
hemi=R,subject=7625176
MRI
hippunfoldT1
ashs
freesurfer

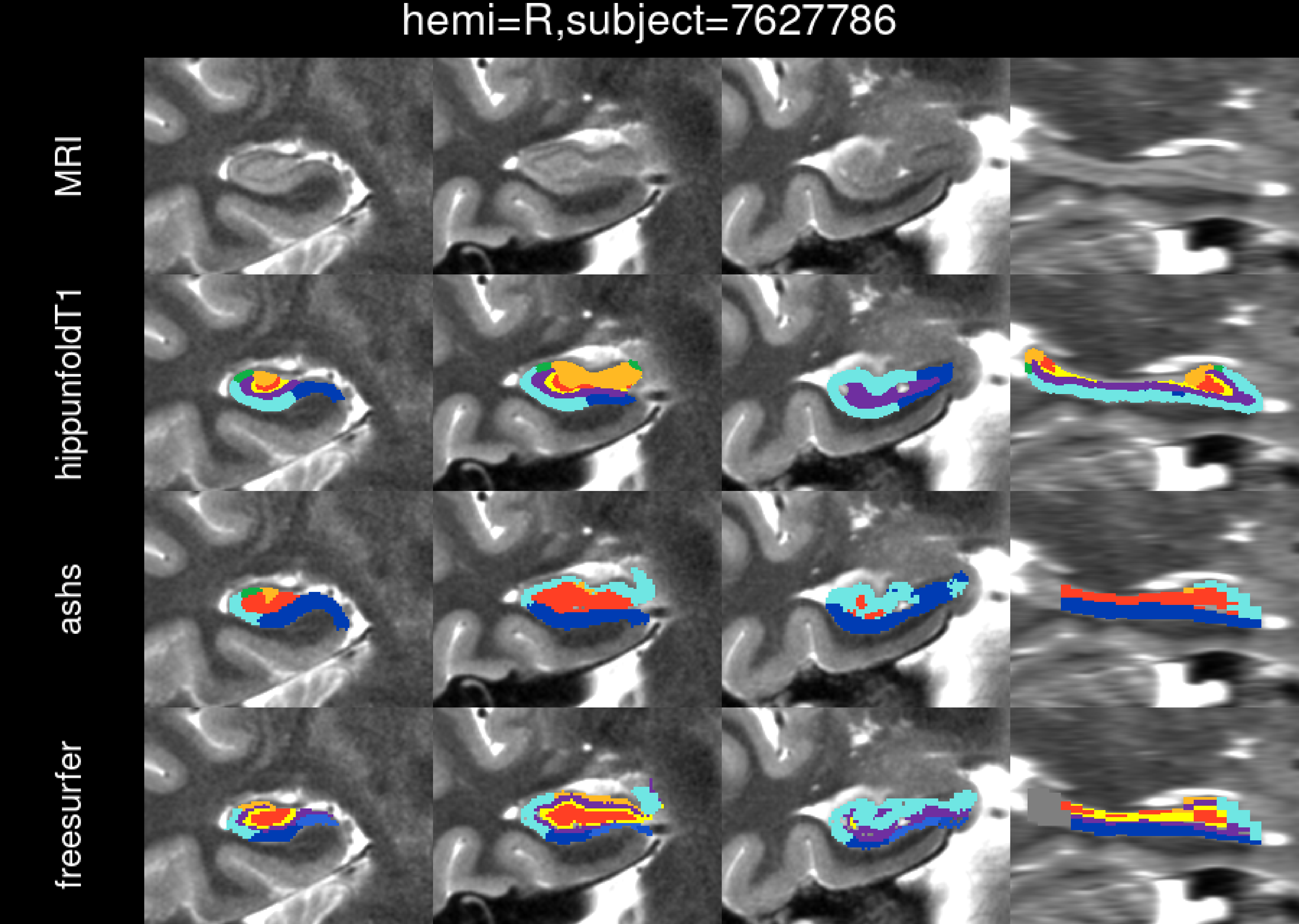
hemi=R,subject=7627786
MRI
hippunfoldT1
ashs
freesurfer

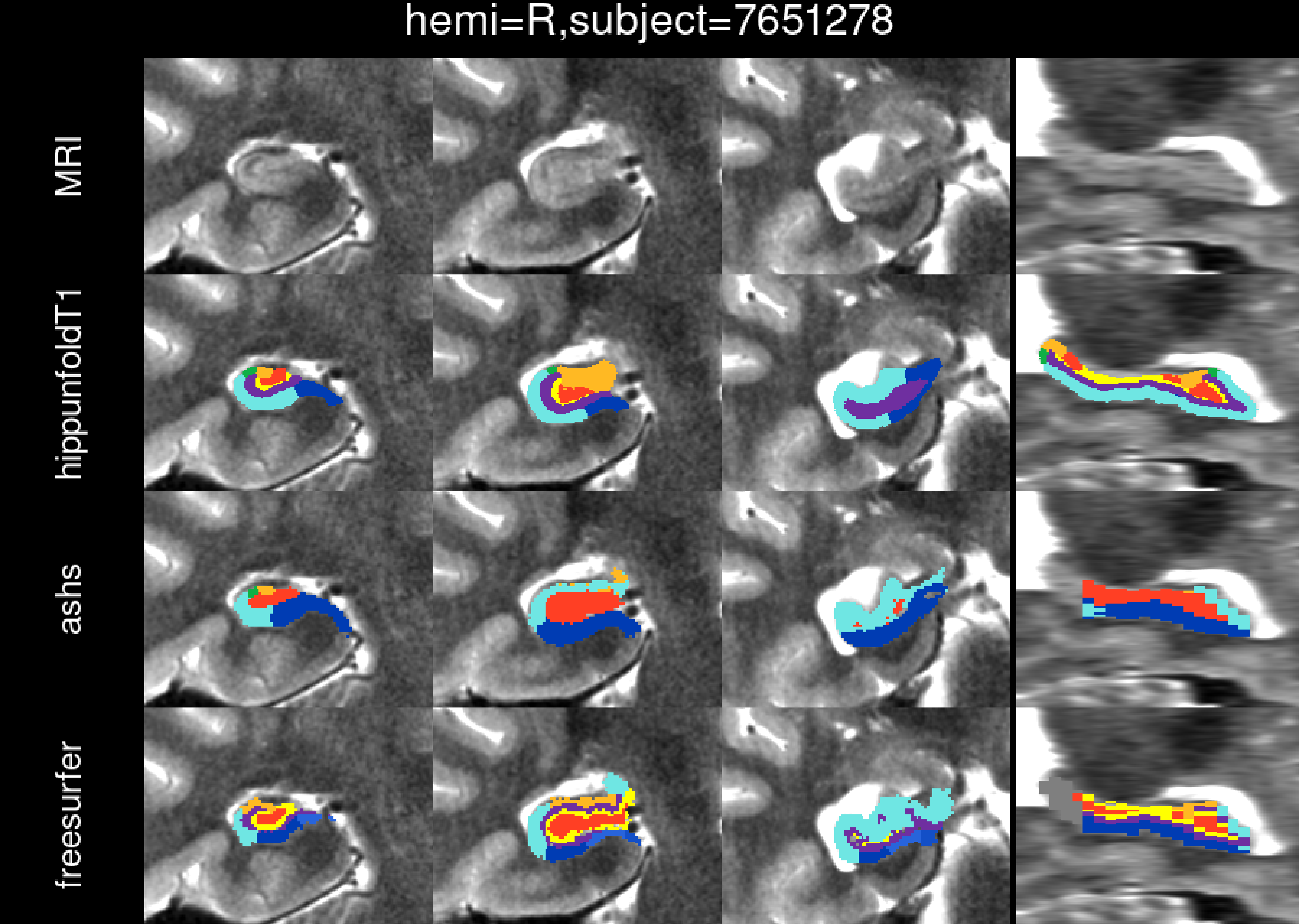
hemi=R,subject=7651278
MRI
hippunfoldT1
ashs
freesurfer

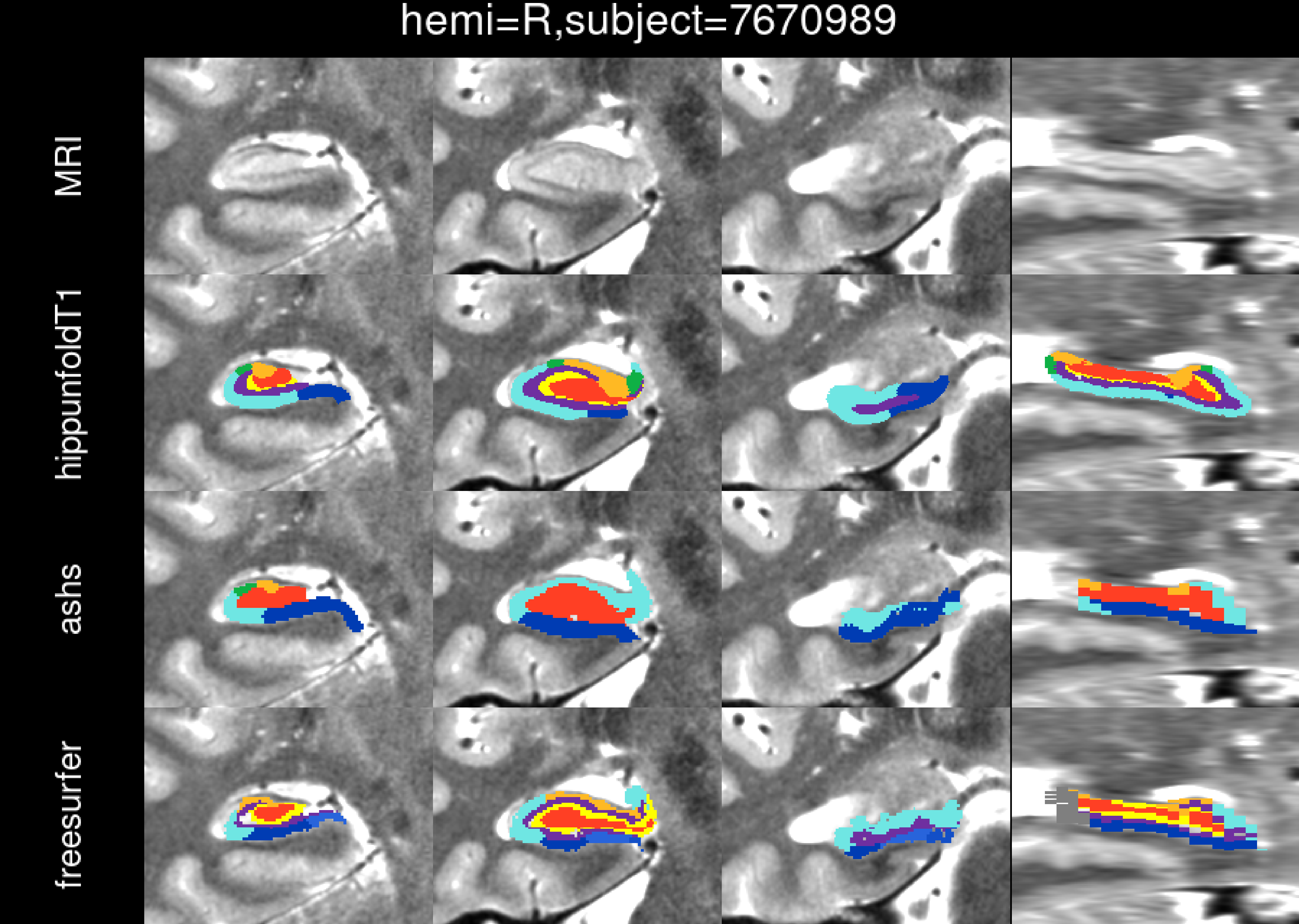
hemi=R,subject=7670989
MRI
hippunfoldT1
ashs
freesurfer

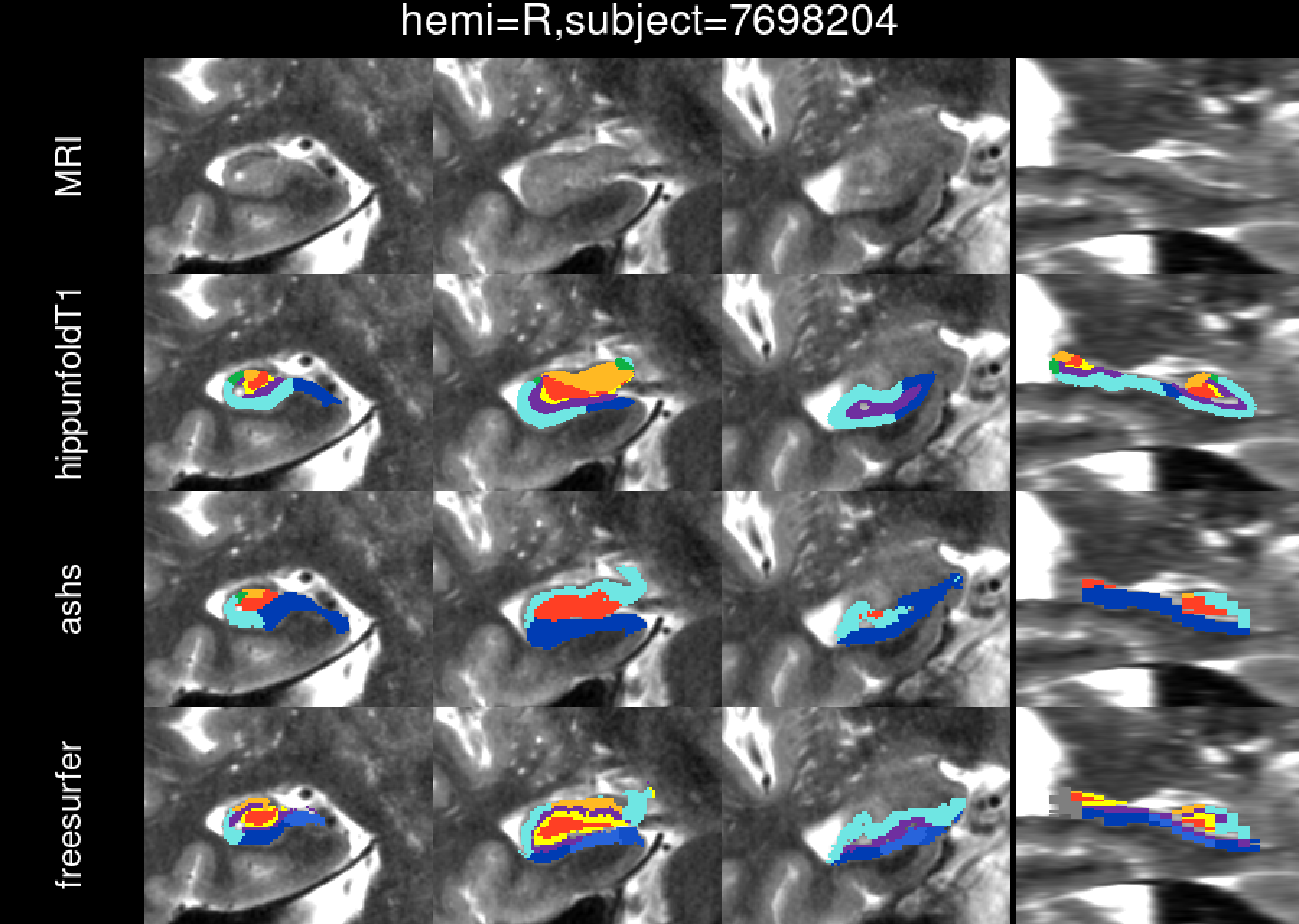
hemi=R,subject=7698204
MRI
hippunfoldT1
ashs
freesurfer

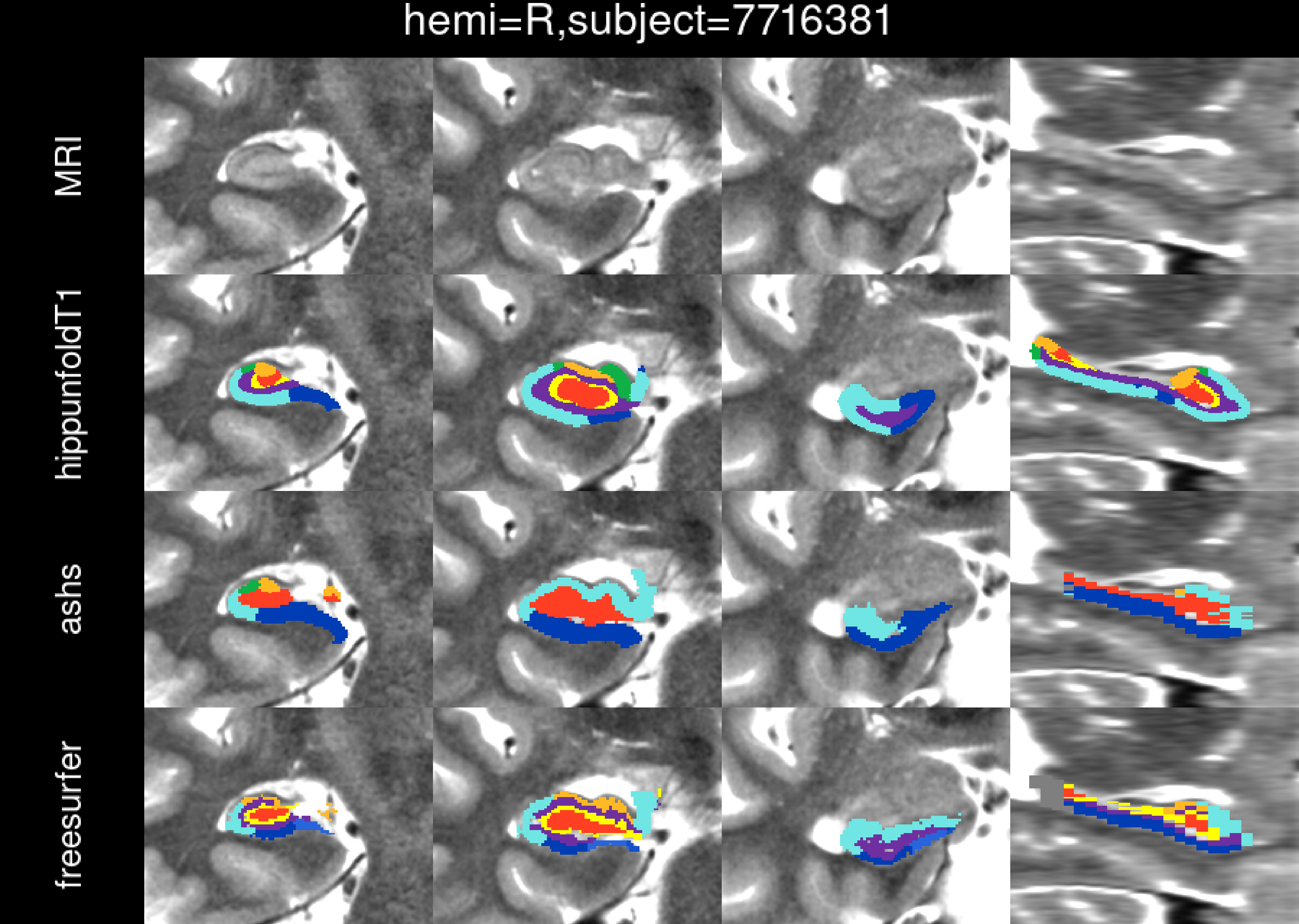
hemi=R,subject=7716381
MRI
hippunfoldT1
ashs
freesurfer

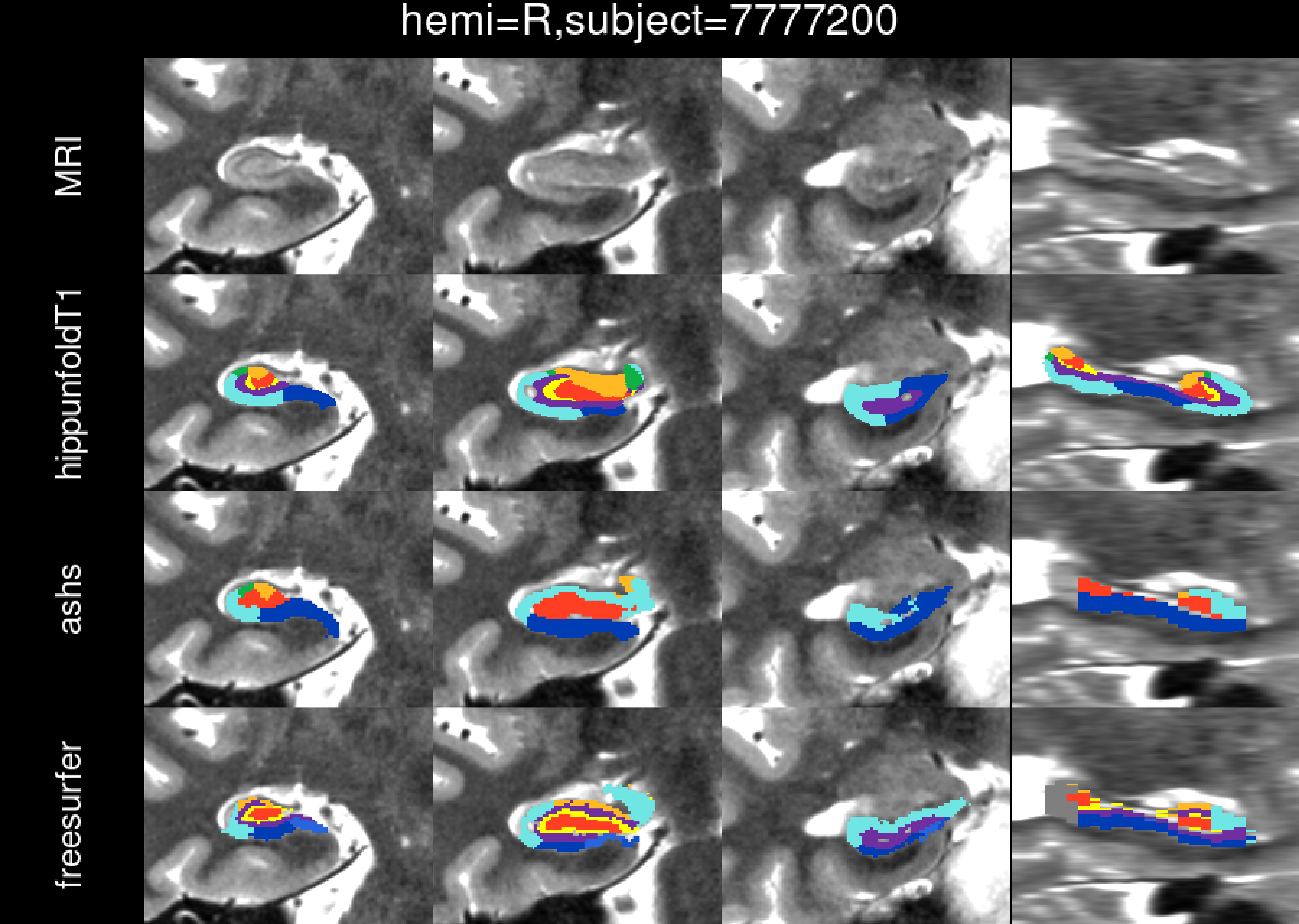
hemi=R,subject=7777200
MRI
hippunfoldT1
ashs
freesurfer

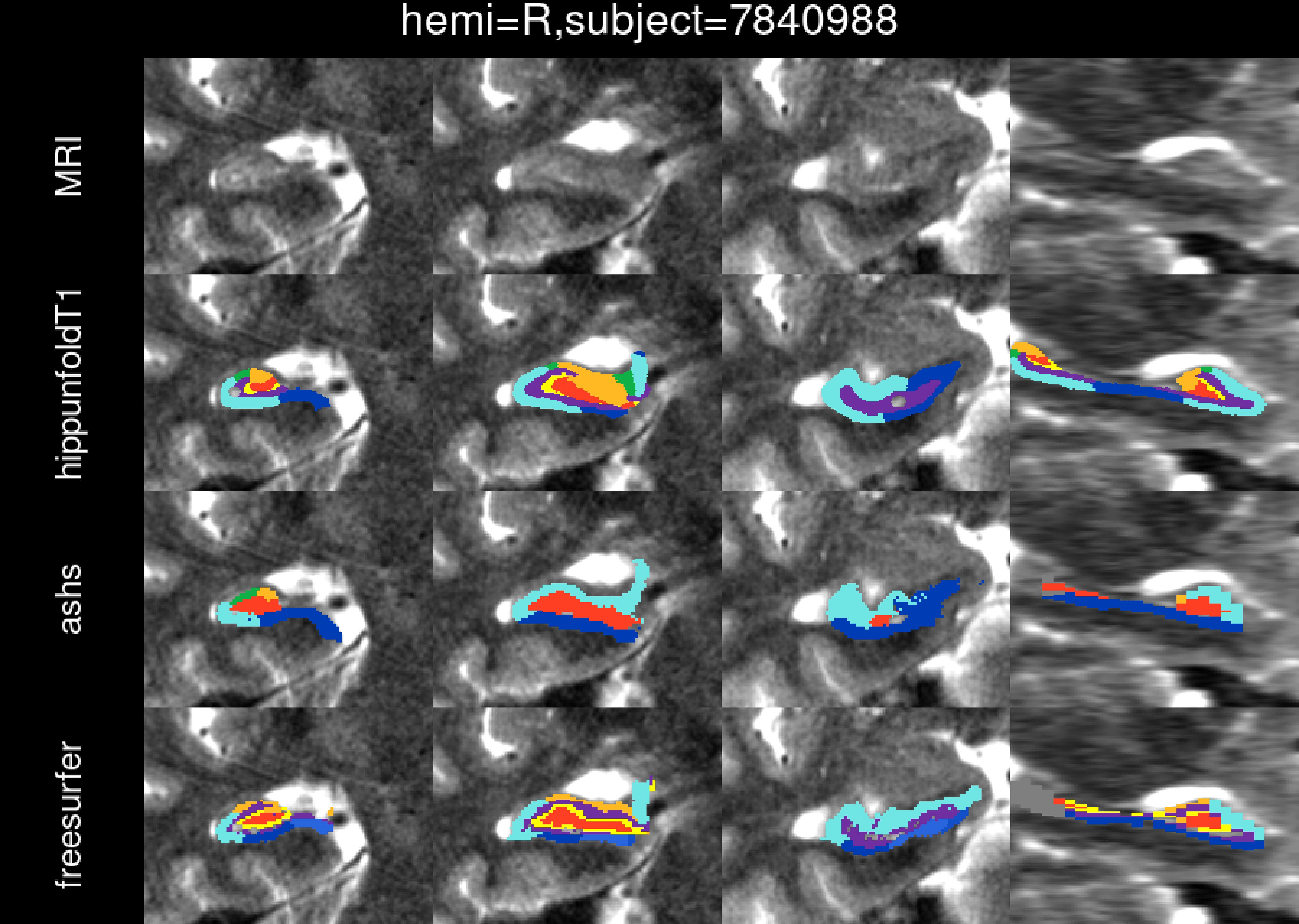

hemi=R,subject=7840988
MRI
hippunfoldT1
ashs
freesurfer

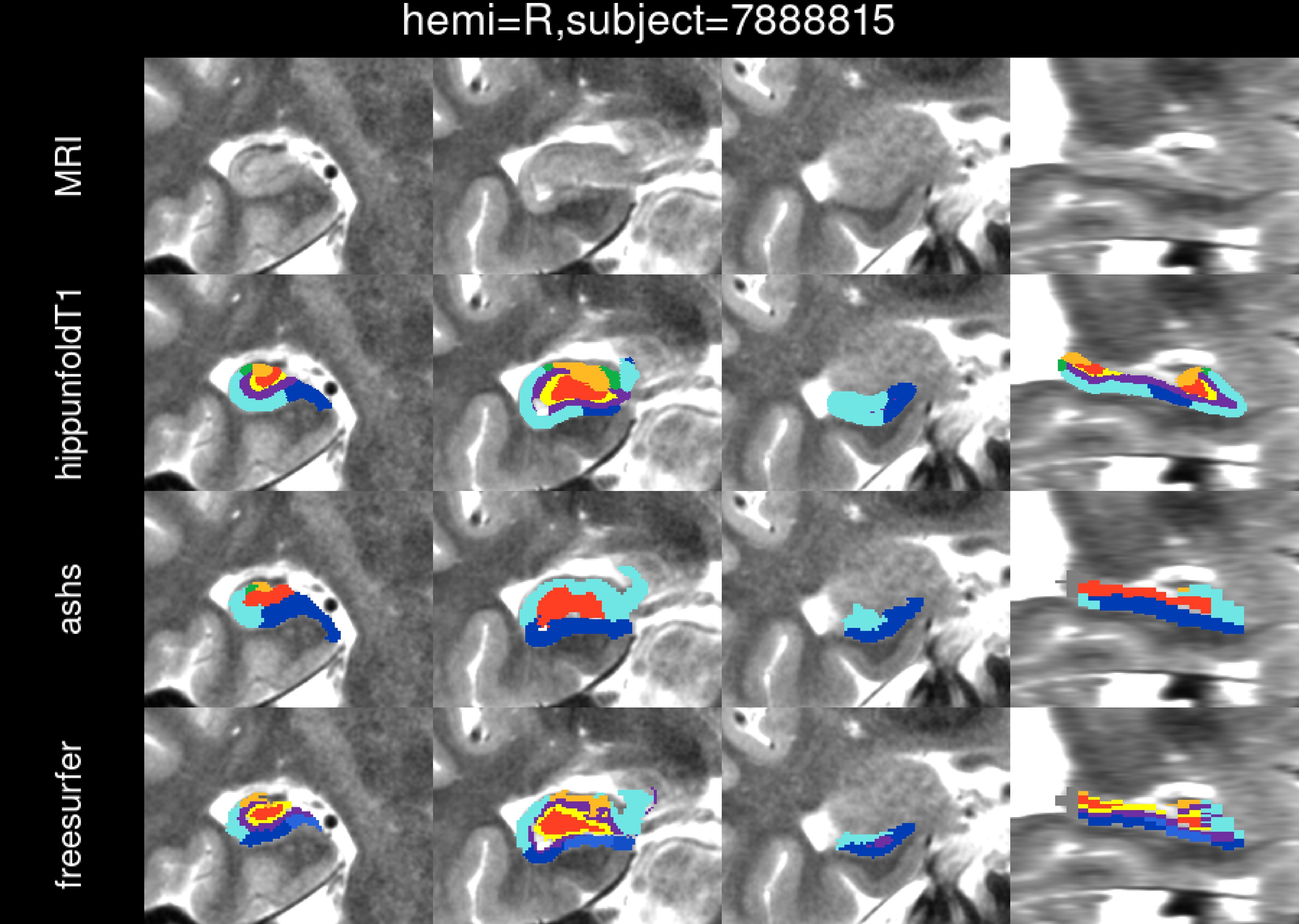

hemi=R,subject=7888815
MRI
hippunfoldT1
ashs
freesurfer

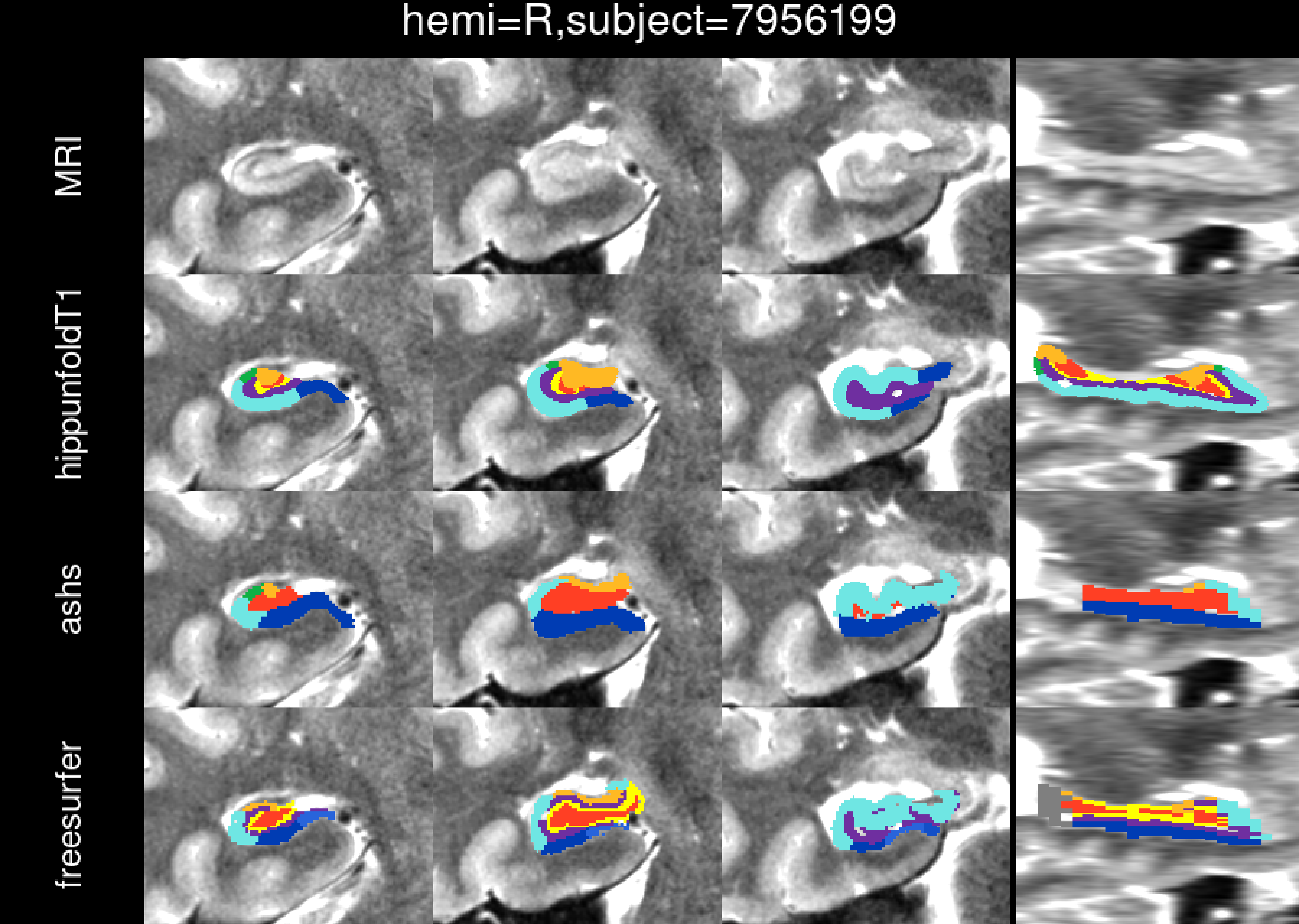
hemi=R,subject=7956199
MRI
hippunfoldT1
ashs
freesurfer

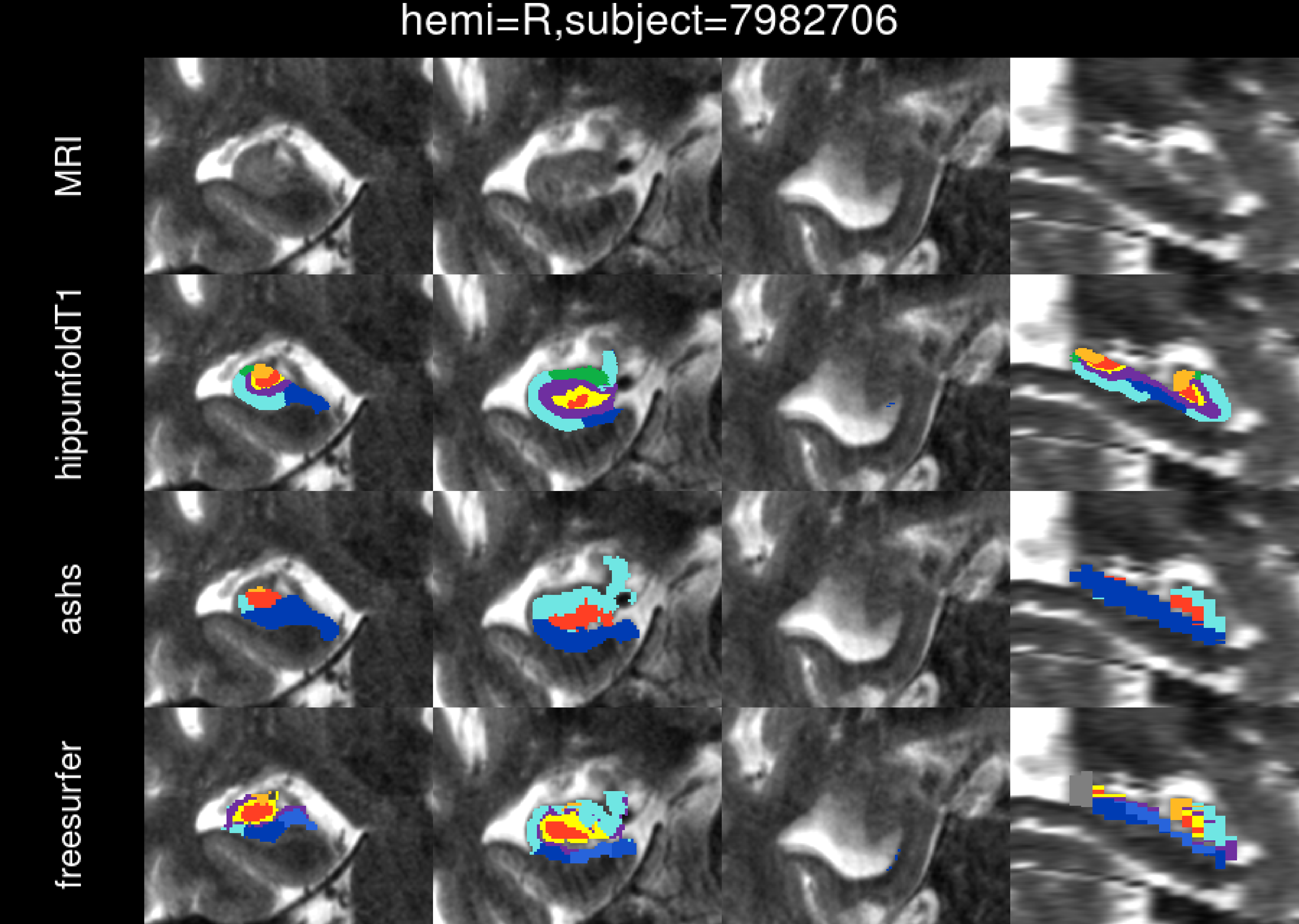

hemi=R,subject=7982706
MRI
hippunfoldT1
ashs
freesurfer

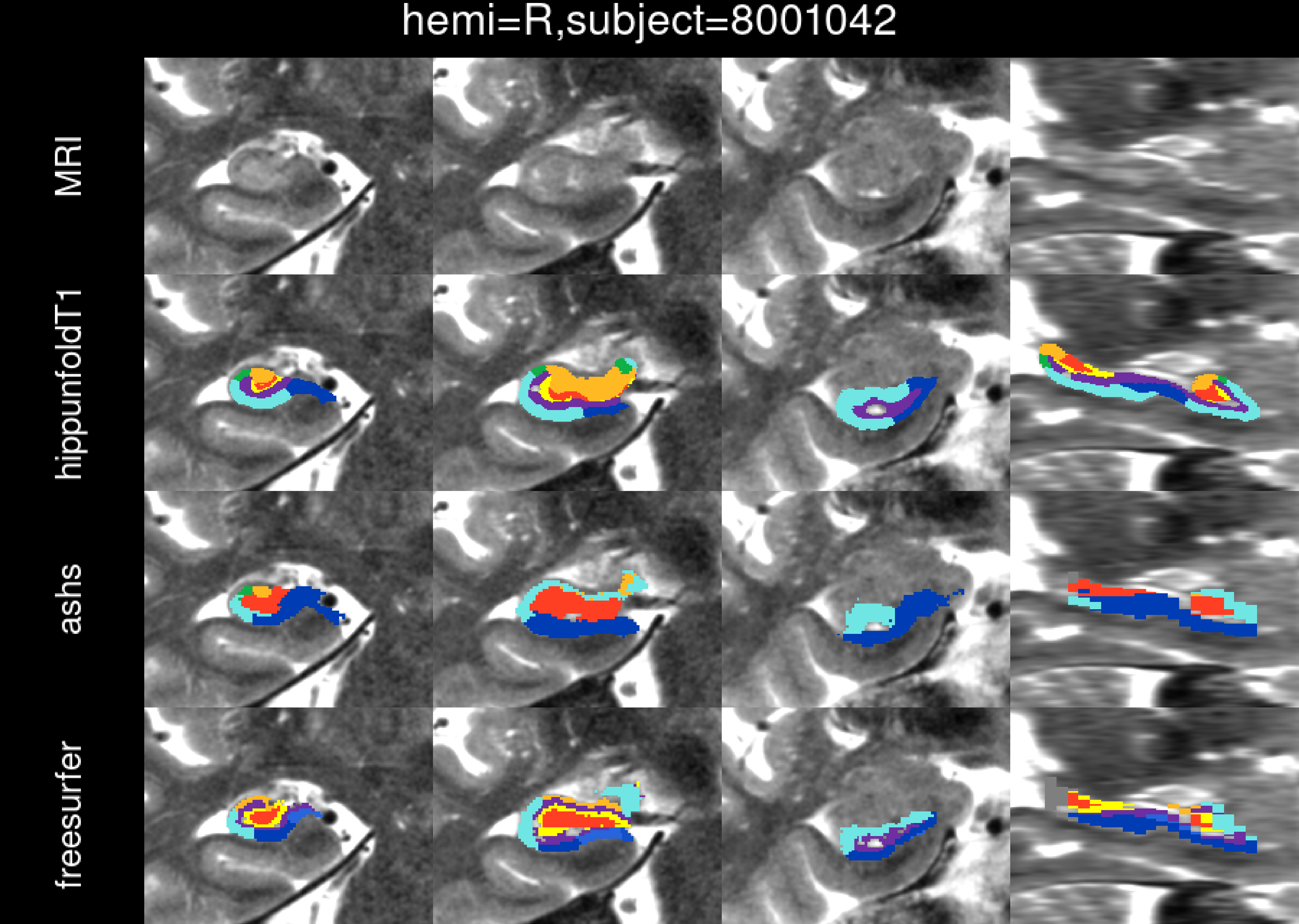
hemi=R,subject=8001042
MRI
hippunfoldT1
ashs
freesurfer

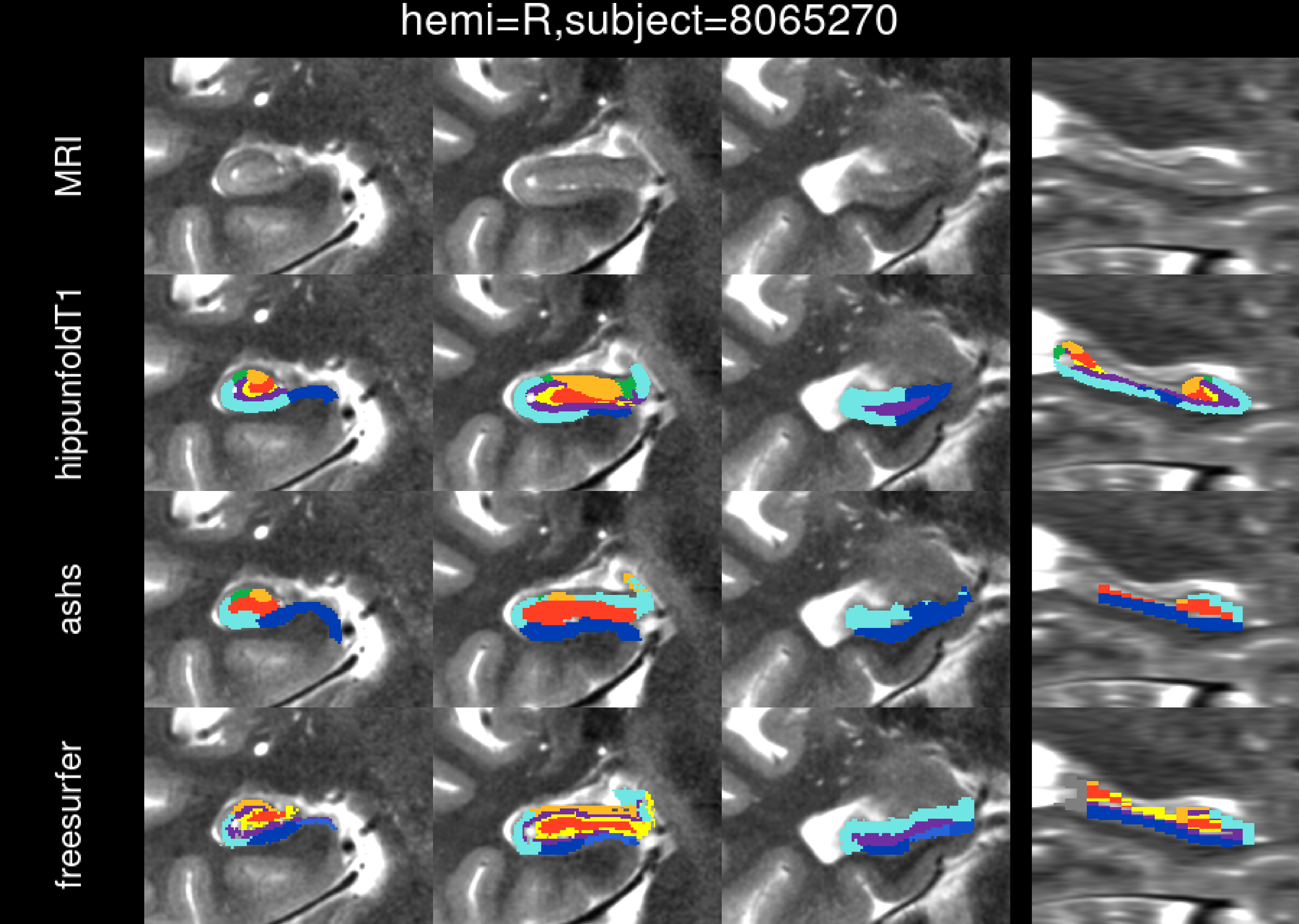
hemi=R,subject=8065270
MRI
hippunfoldT1
ashs
freesurfer

hemi=R,subject=8127165

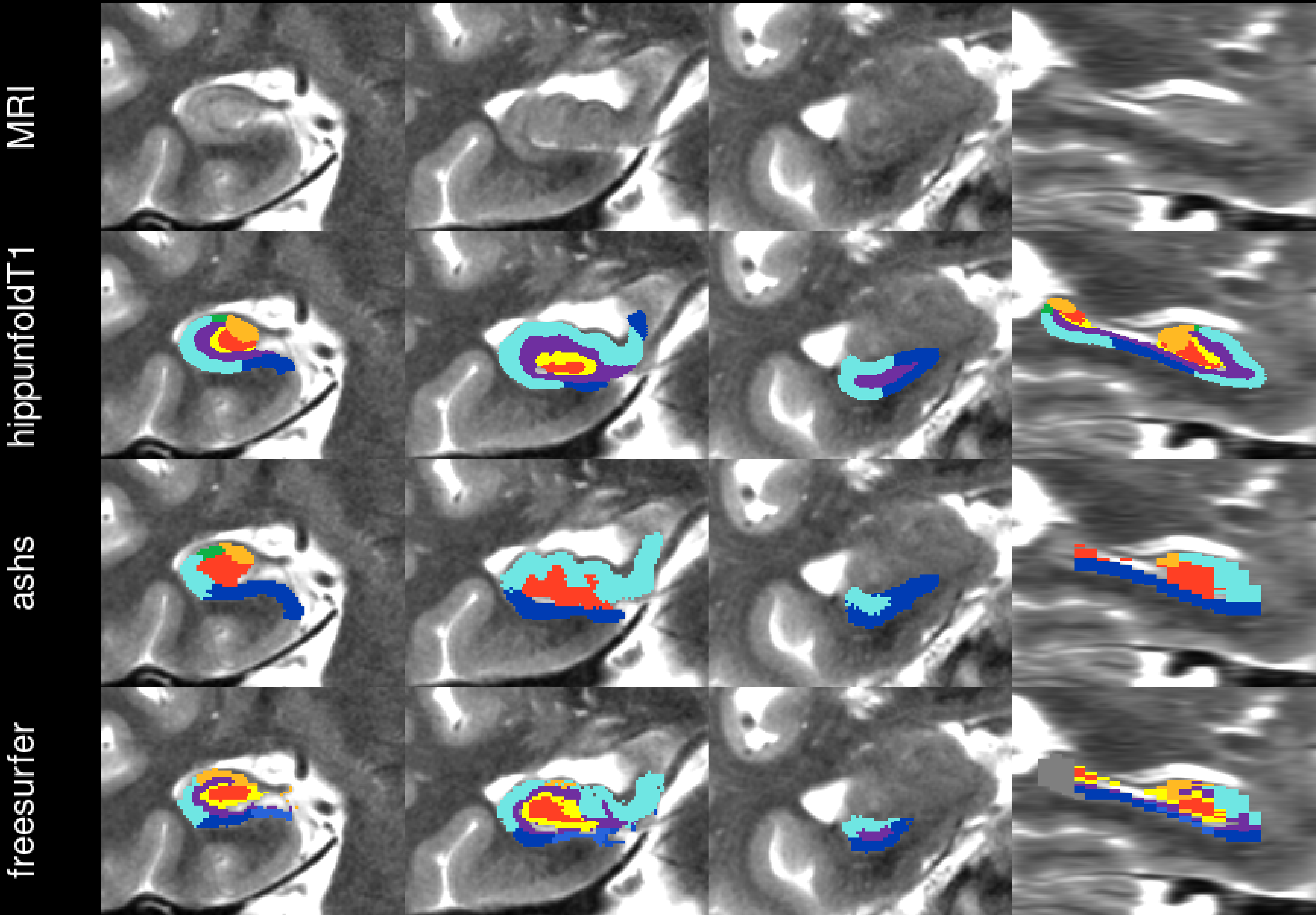

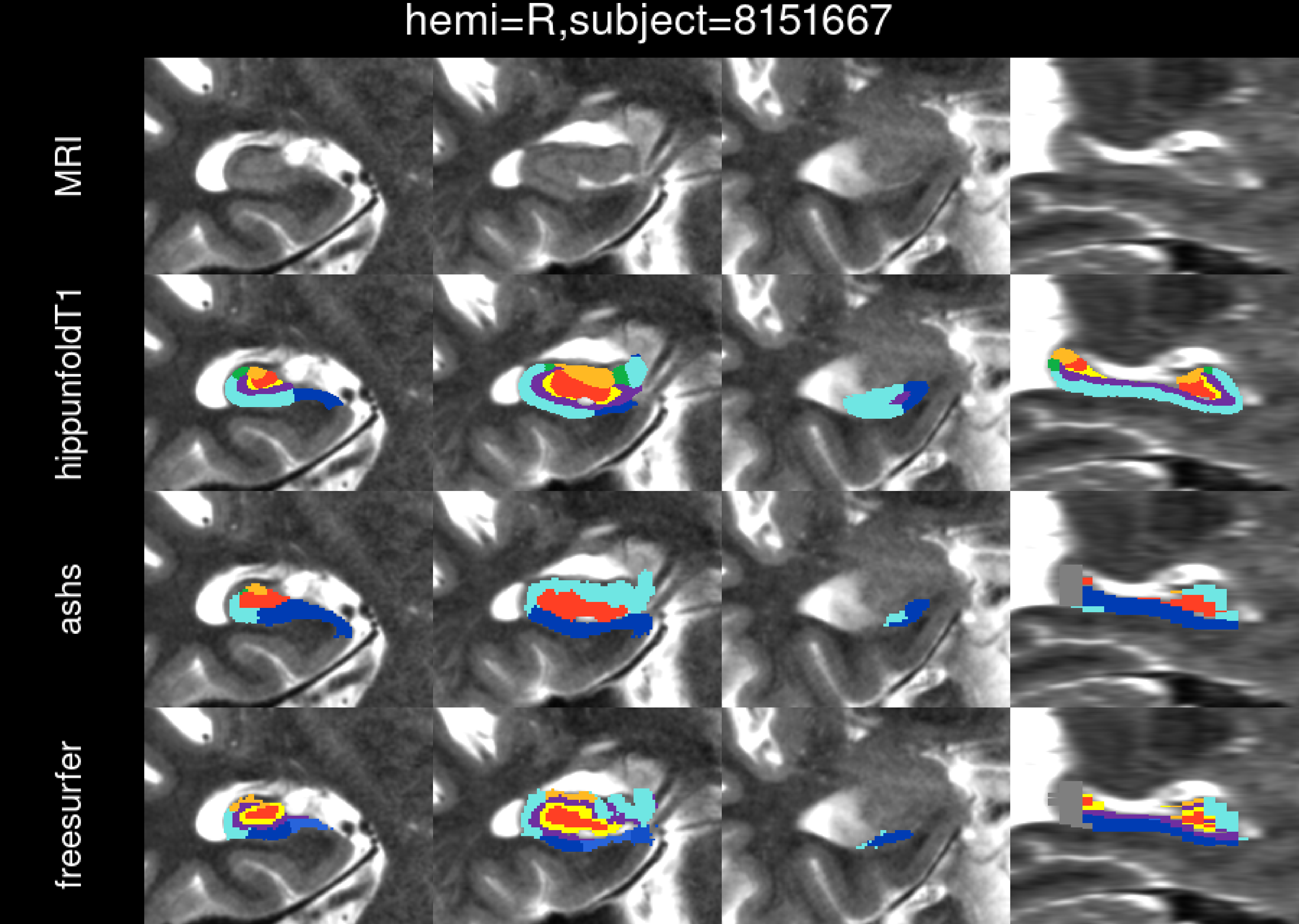
hemi=R,subject=8151667
MRI
hippunfoldT1
ashs
freesurfer

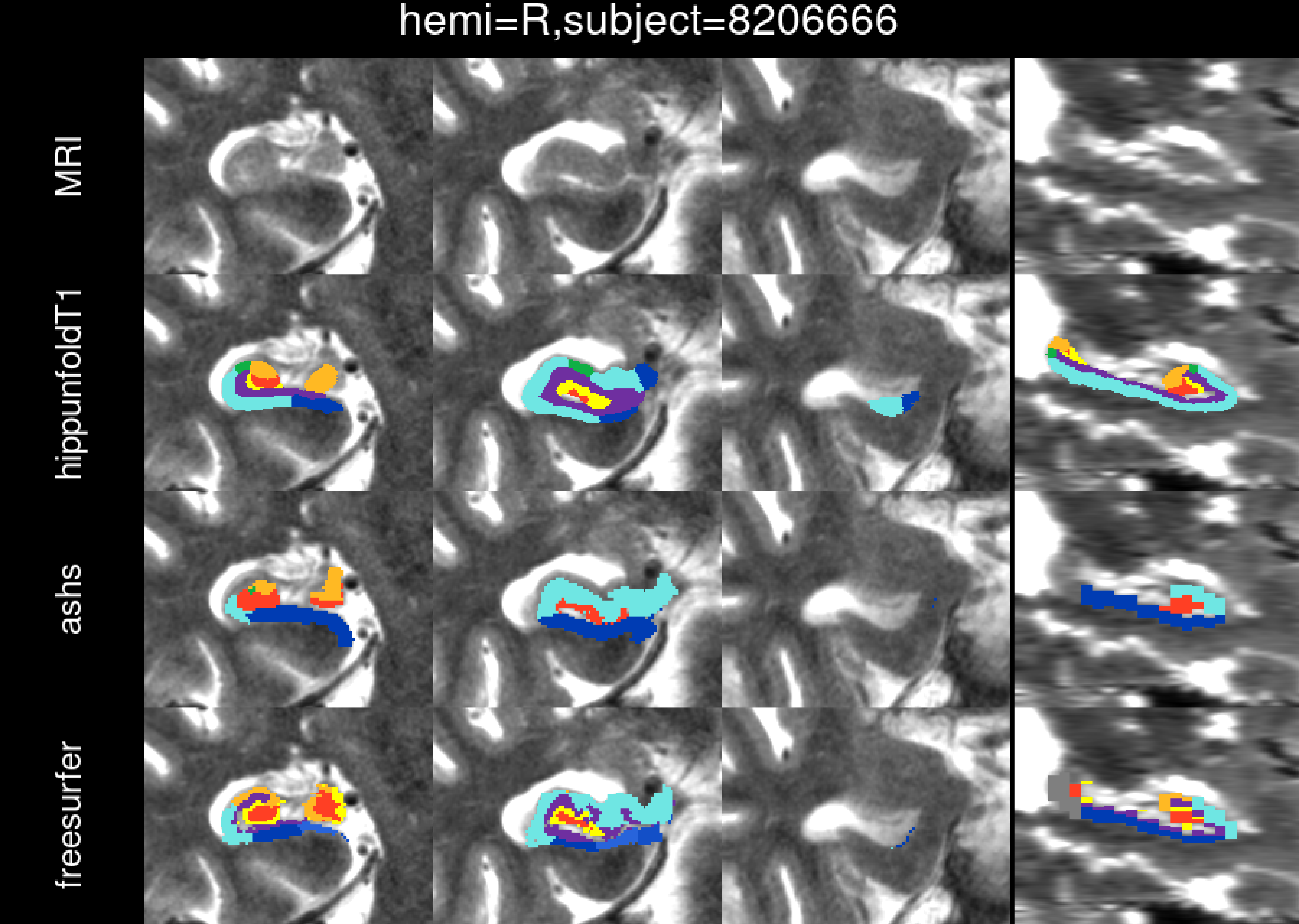
hemi=R,subject=8206666
MRI
hippunfoldT1
ashs
freesurfer

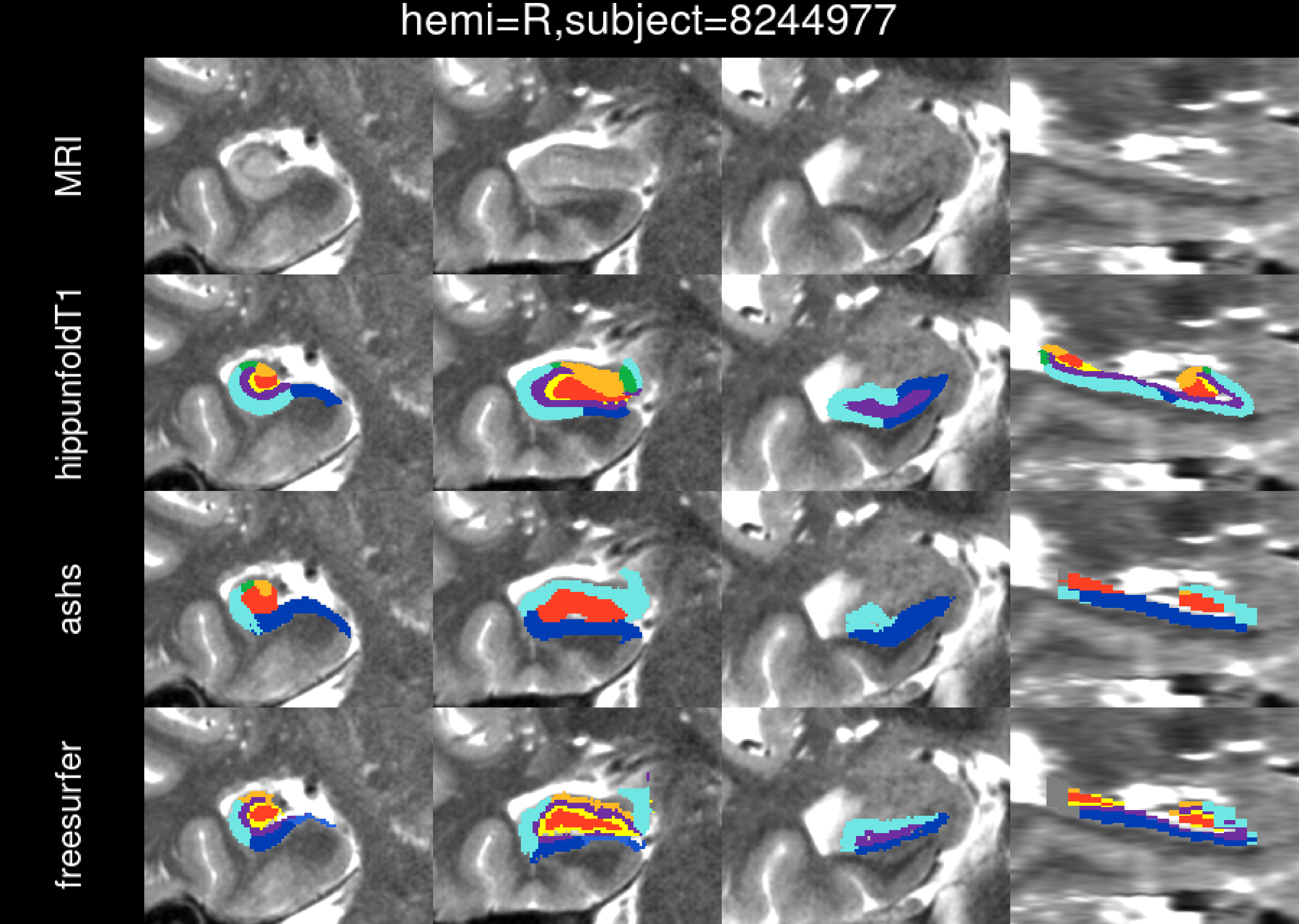
hemi=R,subject=8244977
MRI
hippunfoldT1
ashs
freesurfer

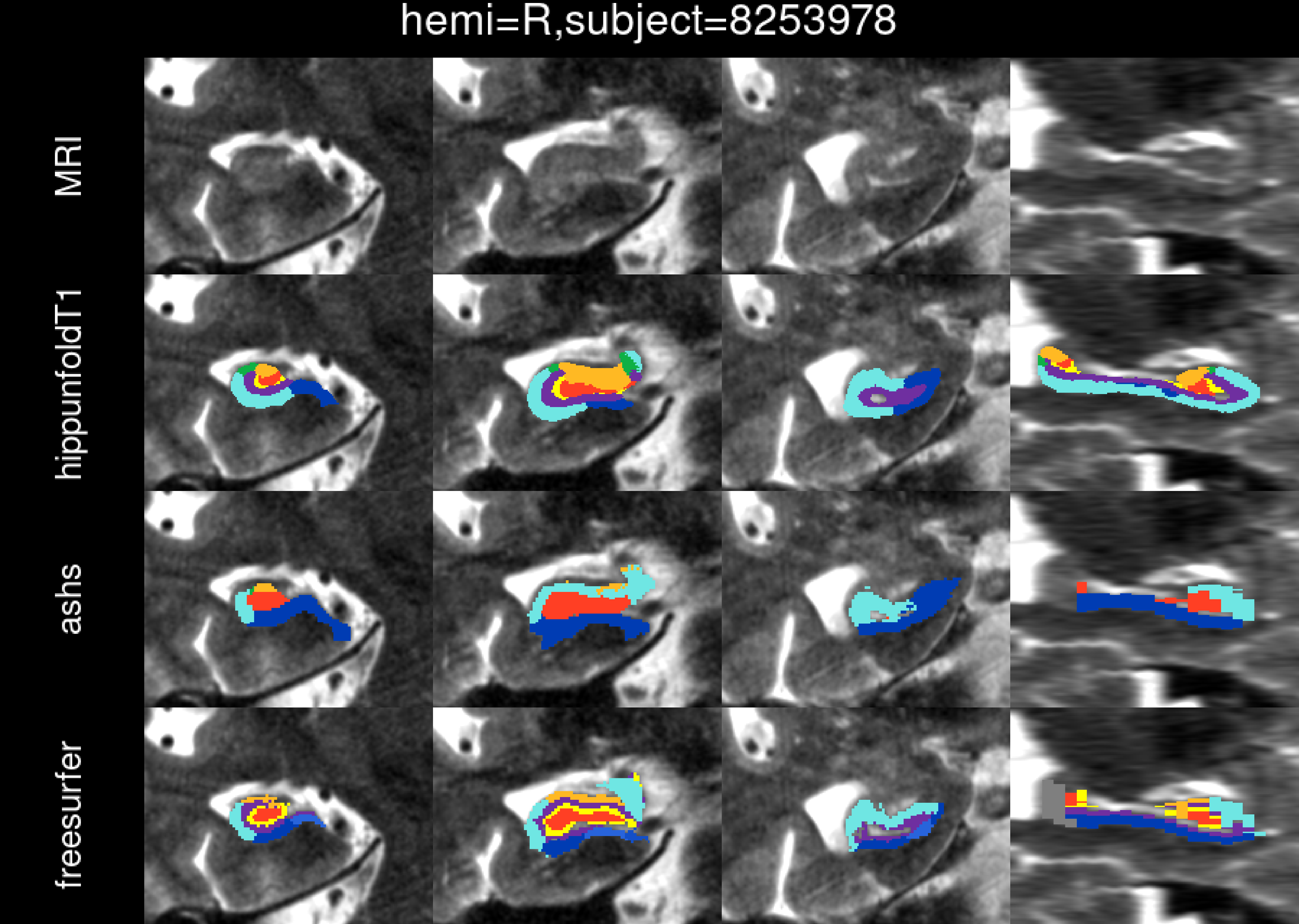
hemi=R,subject=8253978
MRI
hippunfoldT1
ashs
freesurfer

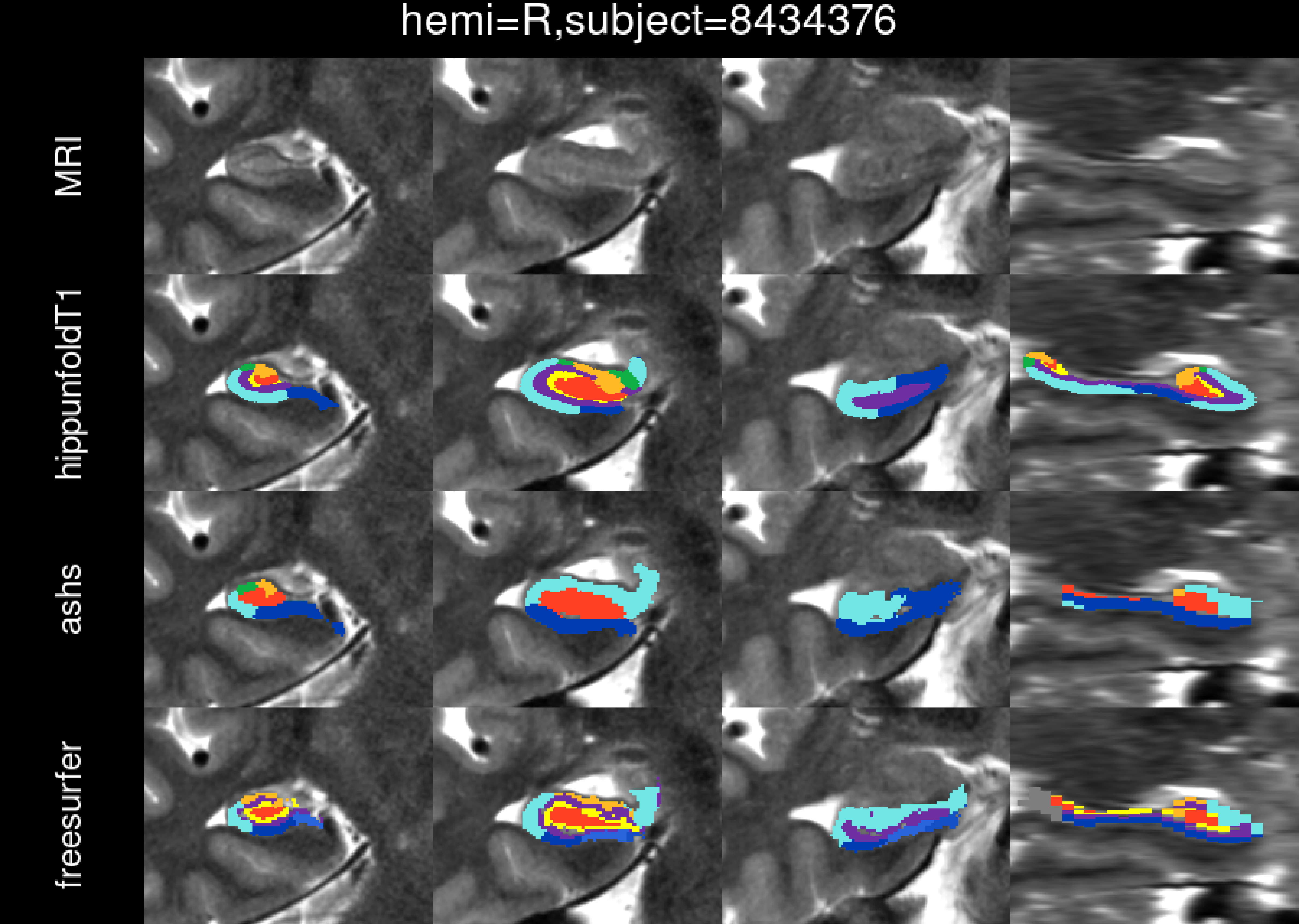
hemi=R,subject=8434376
MRI
hippunfoldT1
ashs
freesurfer

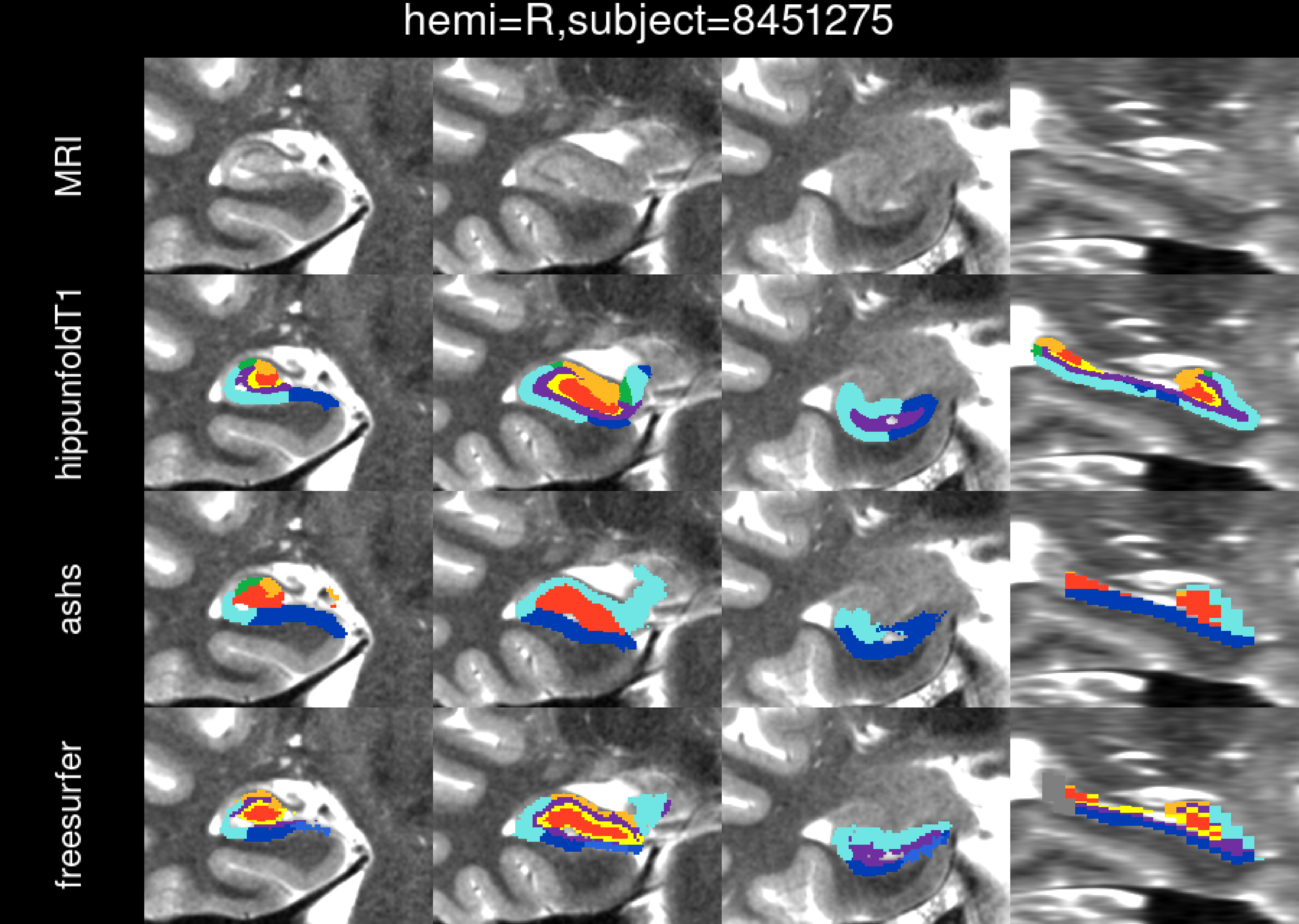
hemi=R,subject=8451275
MRI
hippunfoldT1
ashs
freesurfer

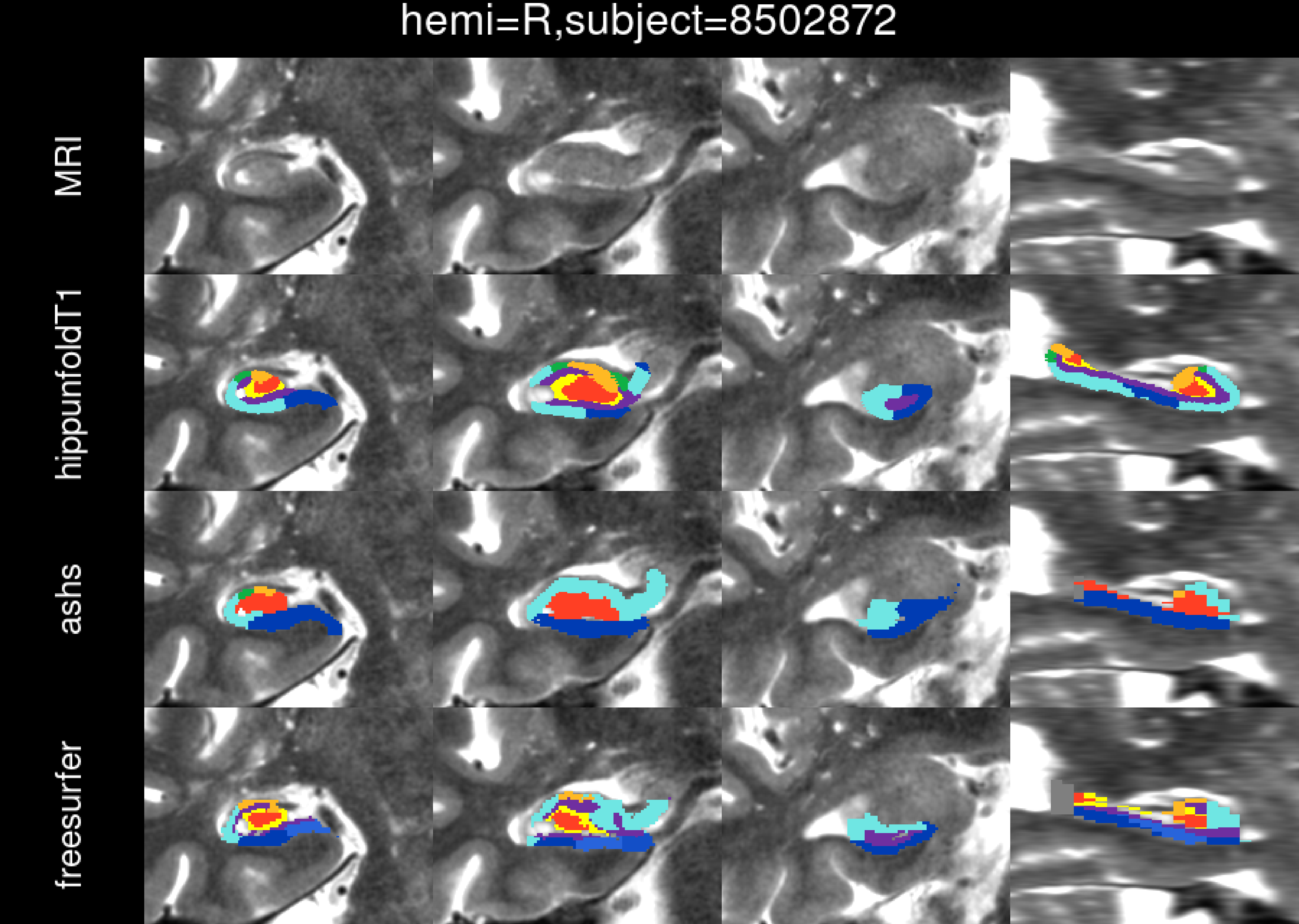
hemi=R,subject=8502872
MRI
hippunfoldT1
ashs
freesurfer

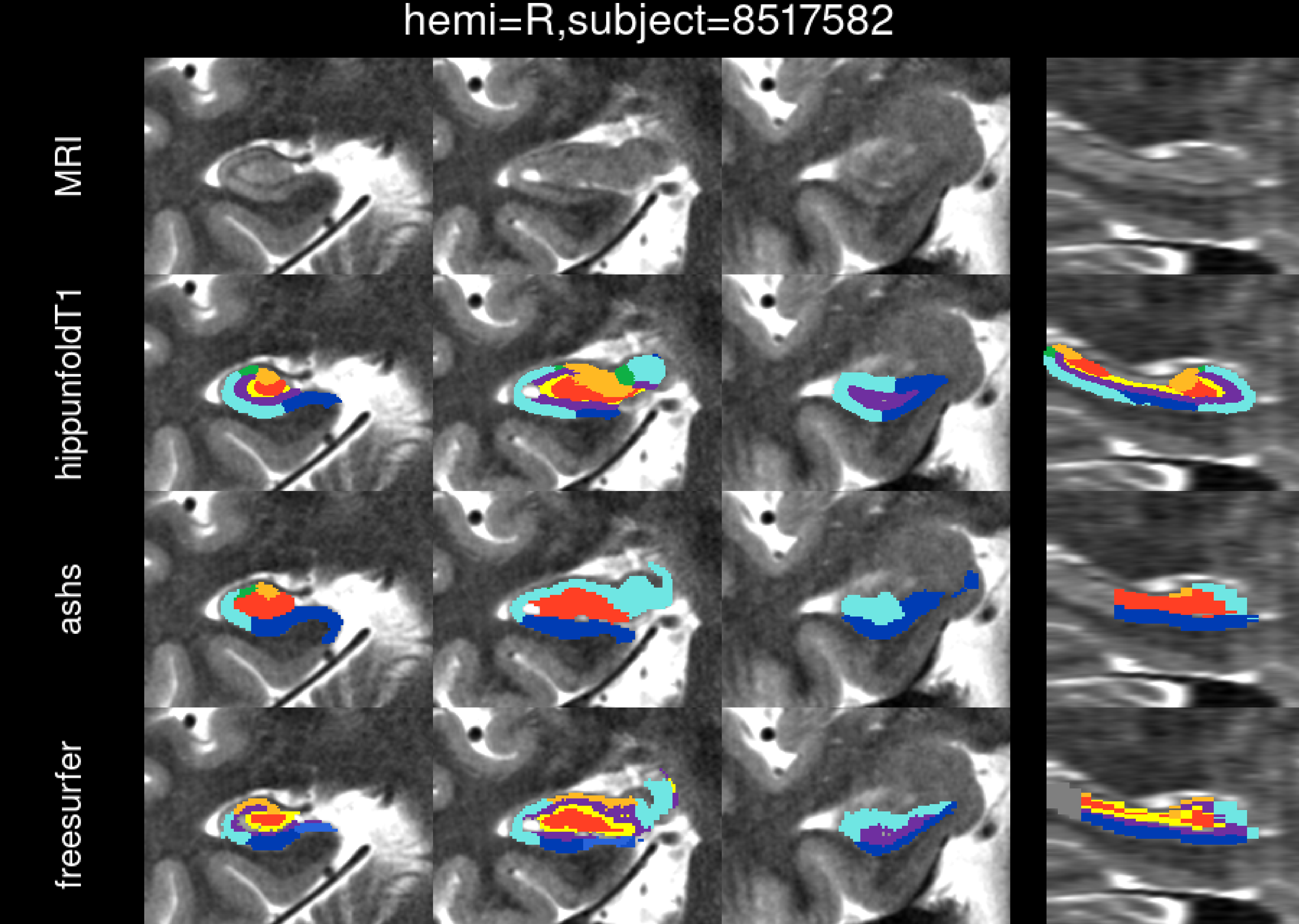

hemi=R,subject=8517582
MRI
hippunfoldT1
ashs
freesurfer

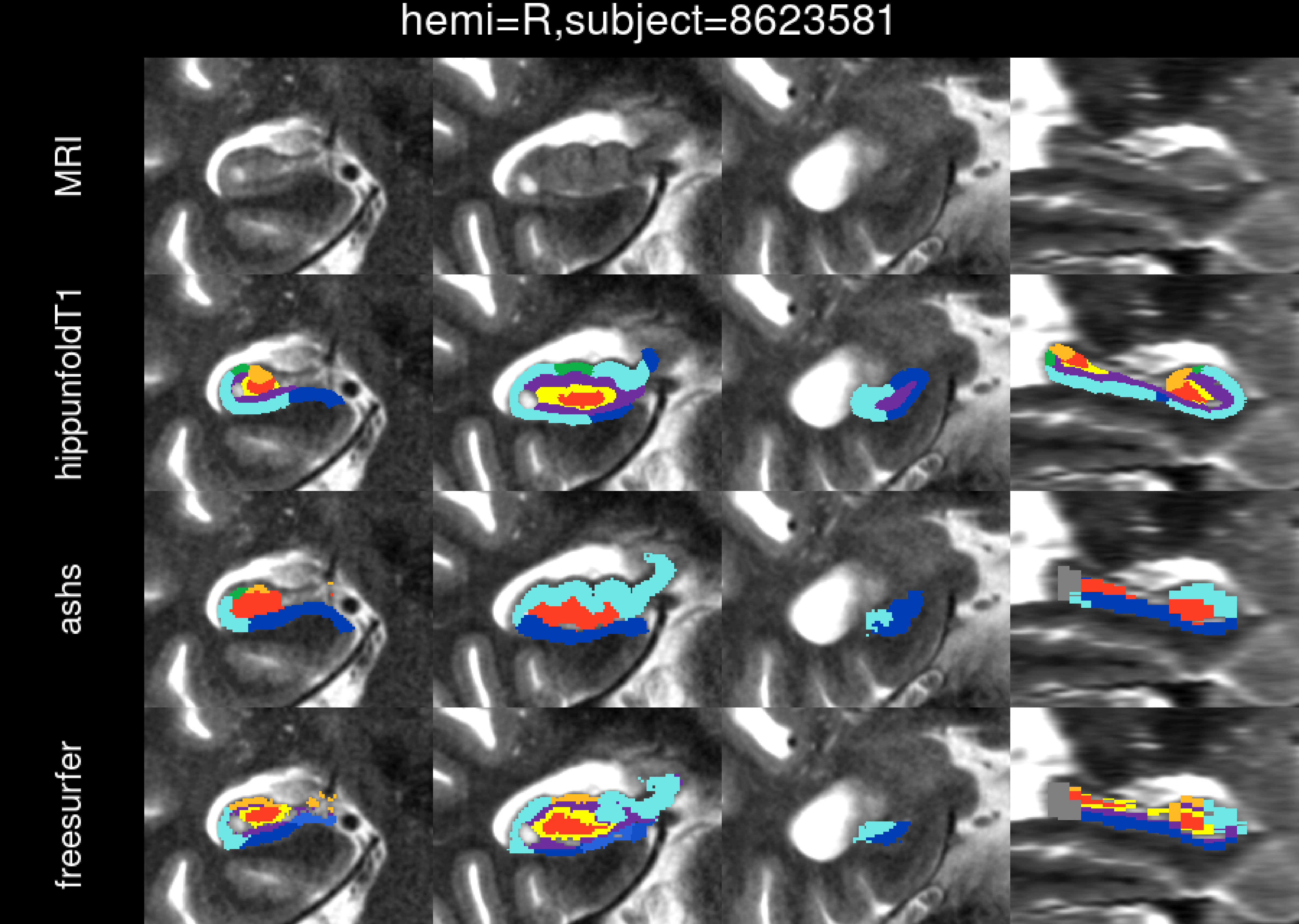

hemi=R,subject=8623581
MRI
hippunfoldT1
ashs
freesurfer

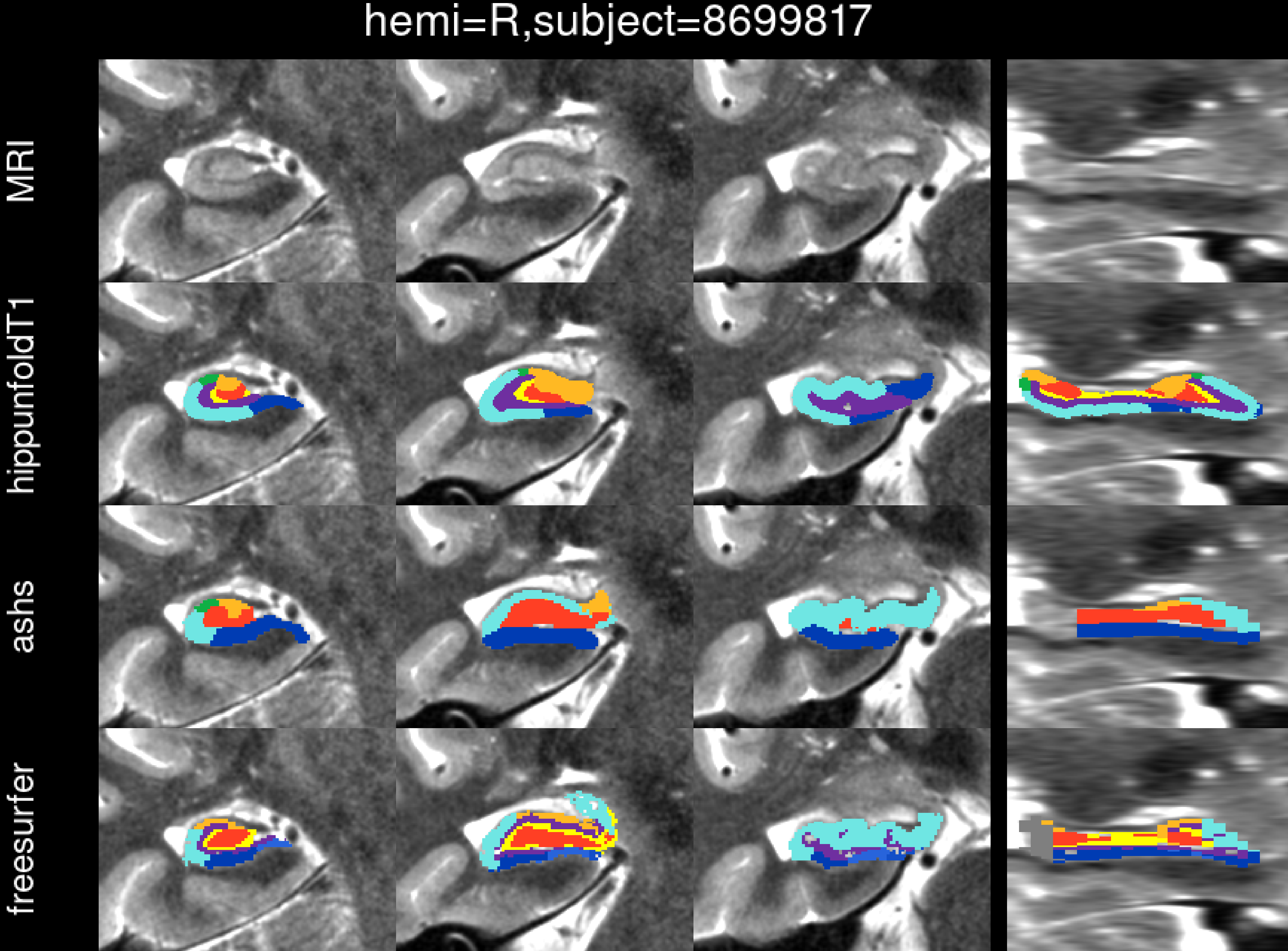
hemi=R,subject=8699817
MRI
hippunfoldT1
ashs
freesurfer

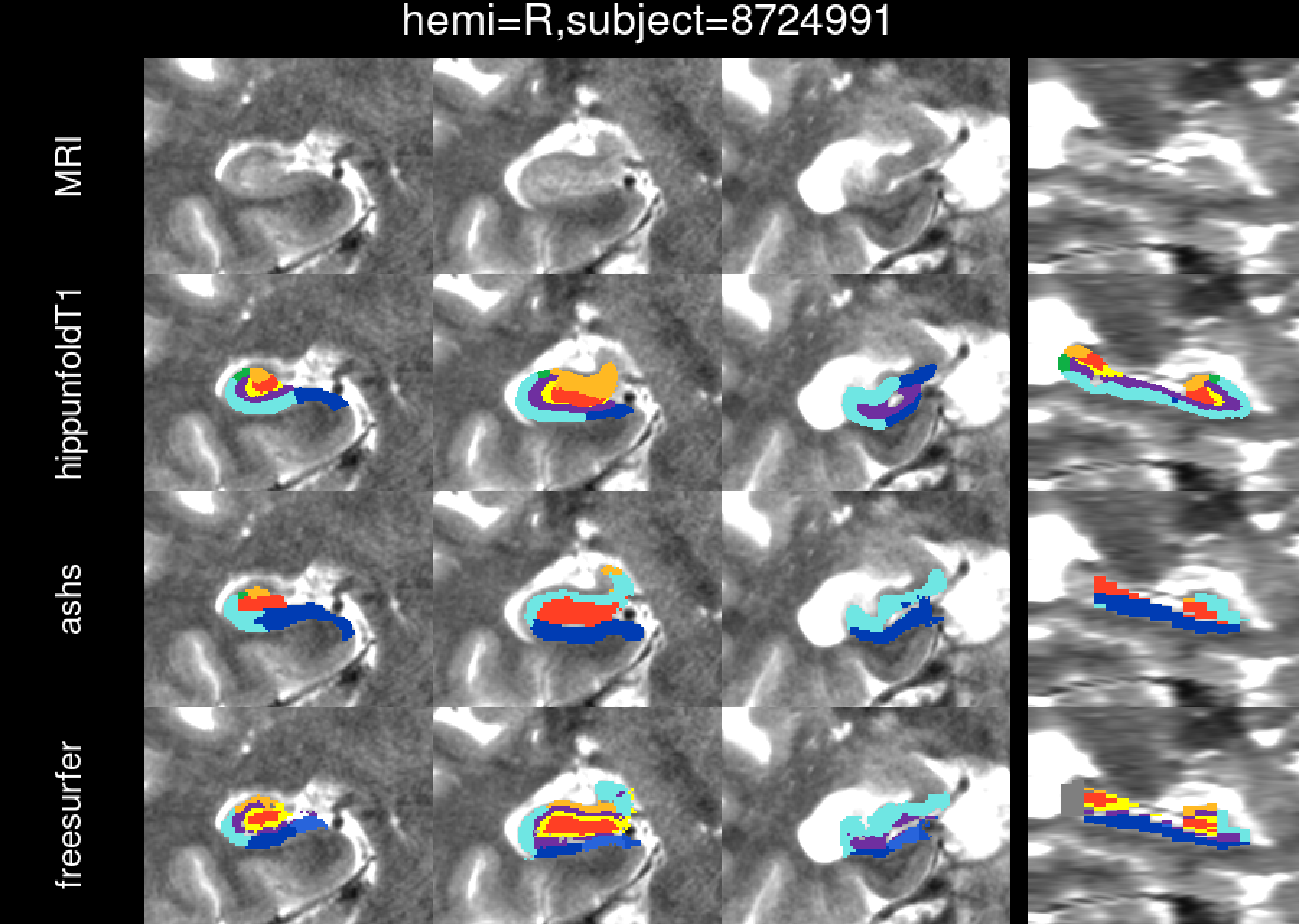
hemi=R,subject=8724991
MRI
hippunfoldT1
ashs
freesurfer

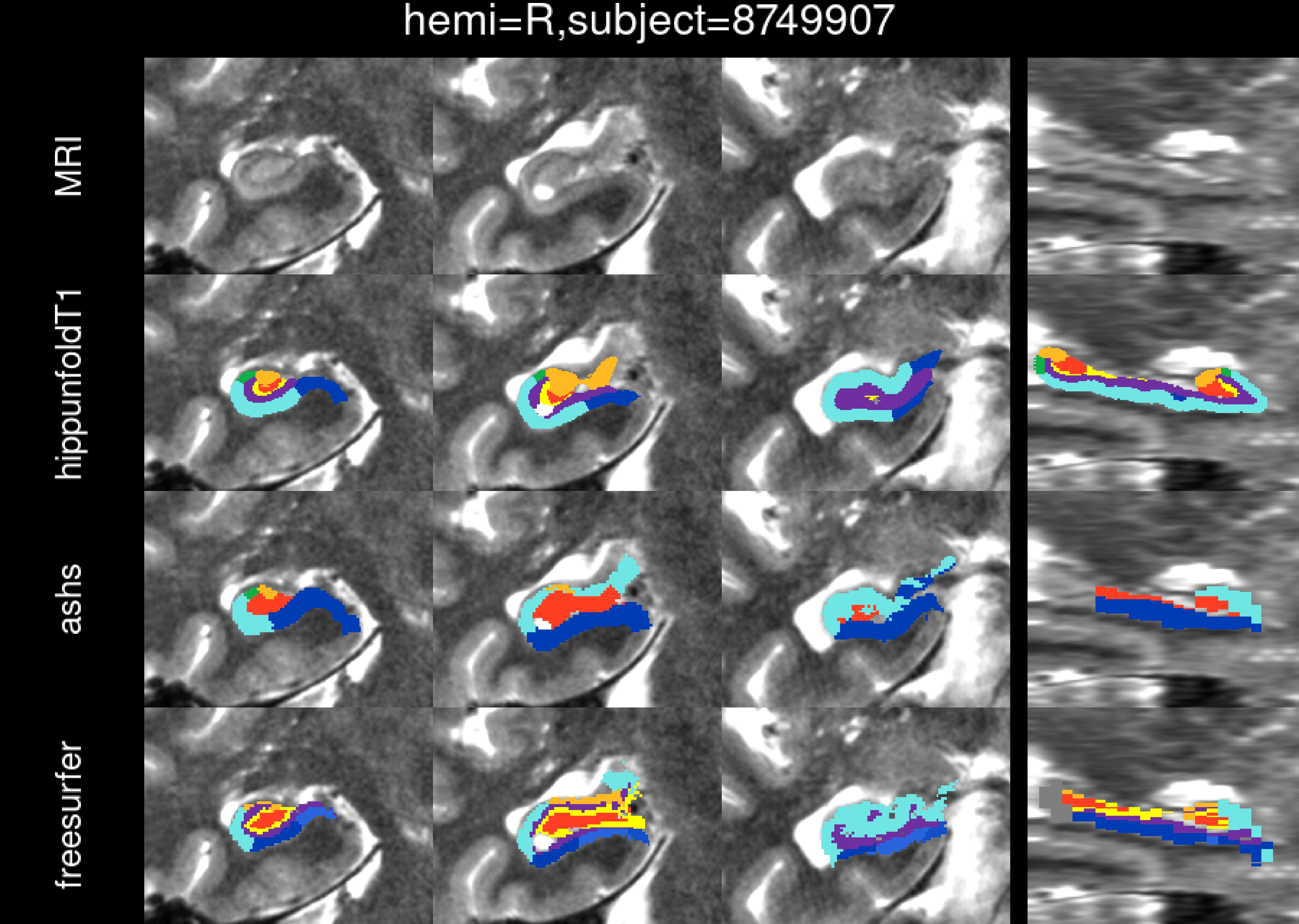
hemi=R,subject=8749907
MRI
hippunfoldT1
ashs
freesurfer

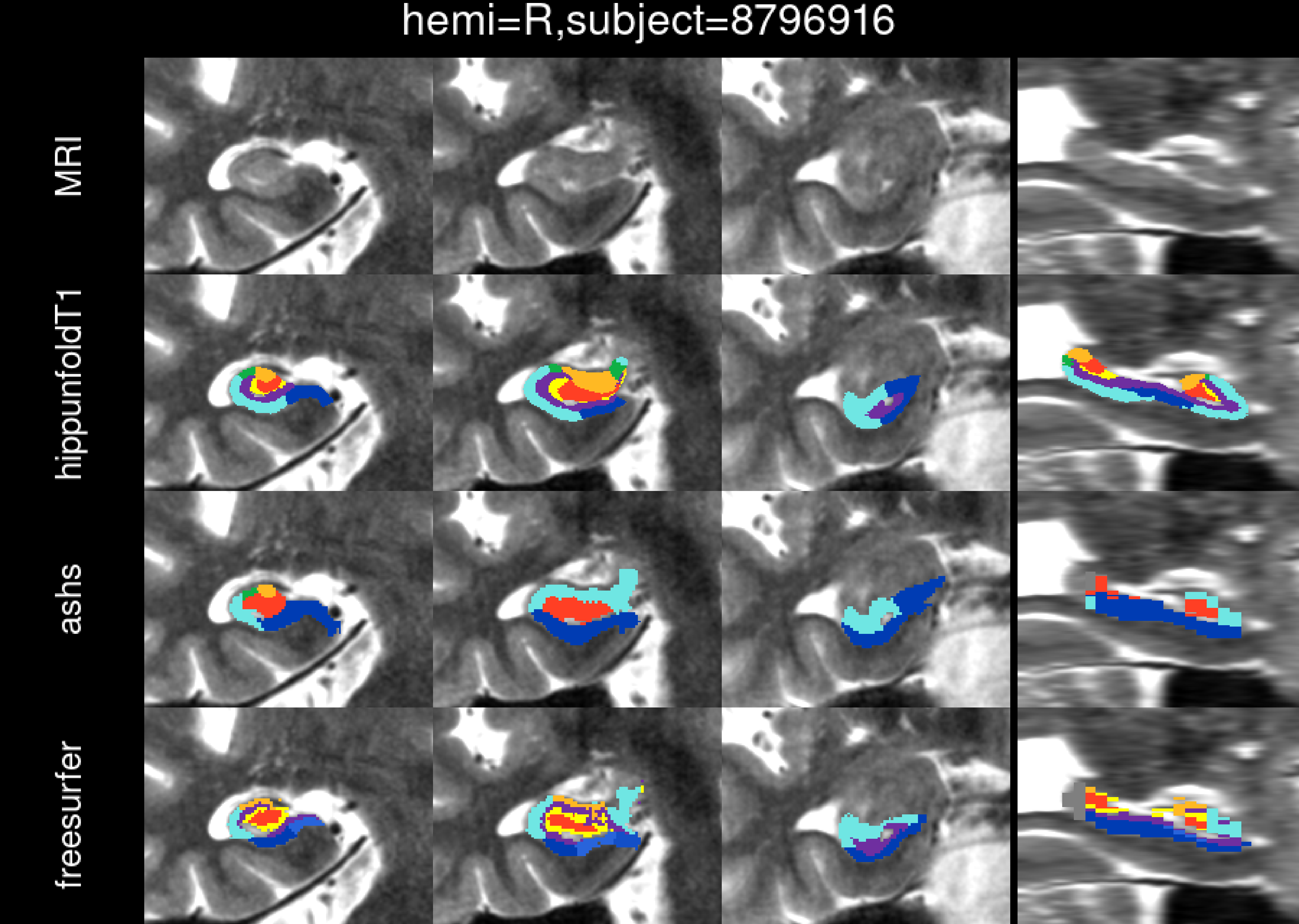
hemi=R,subject=8796916
MRI
hippunfoldT1
ashs
freesurfer

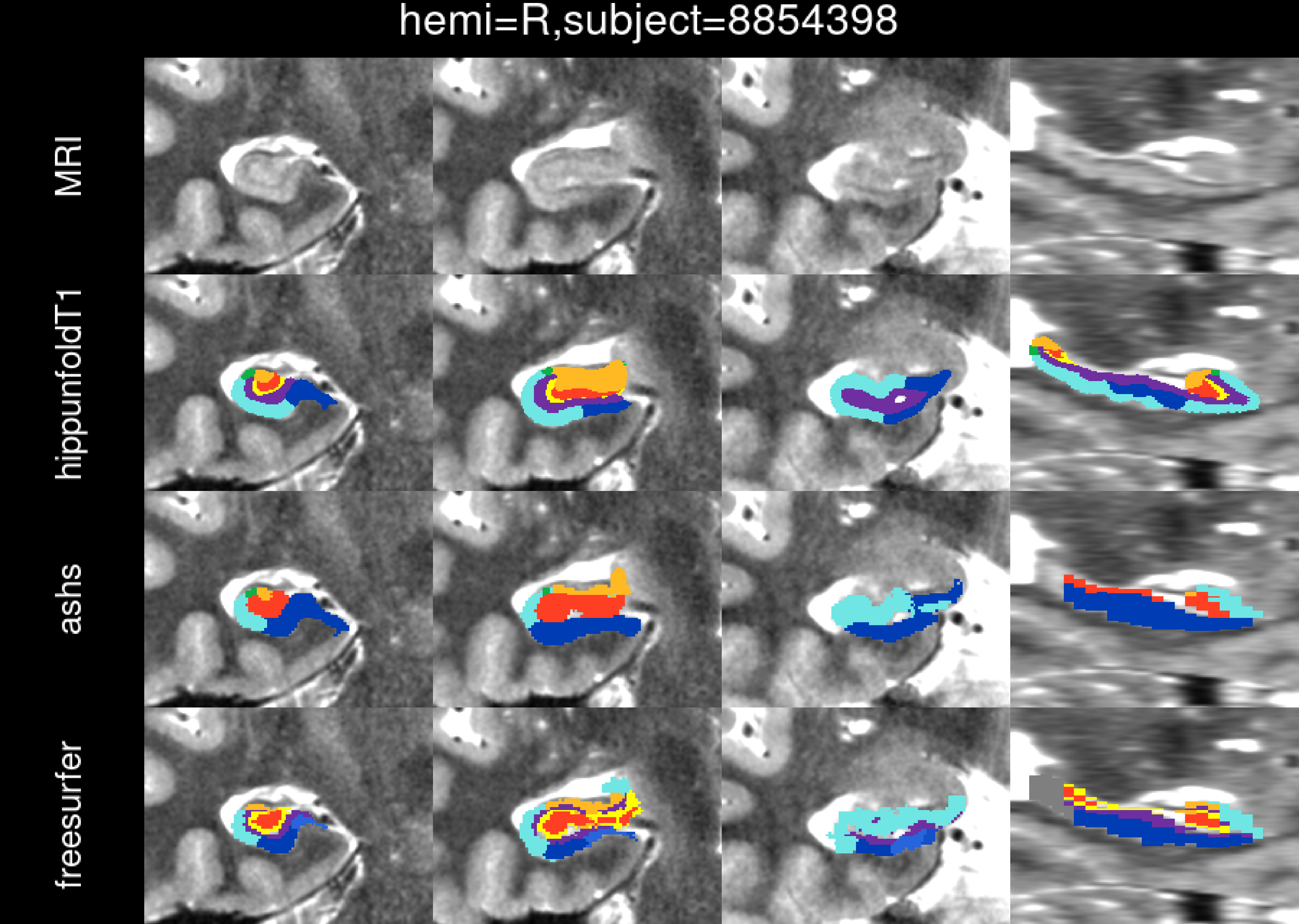
hemi=R,subject=8854398
MRI
hippunfoldT1
ashs
freesurfer

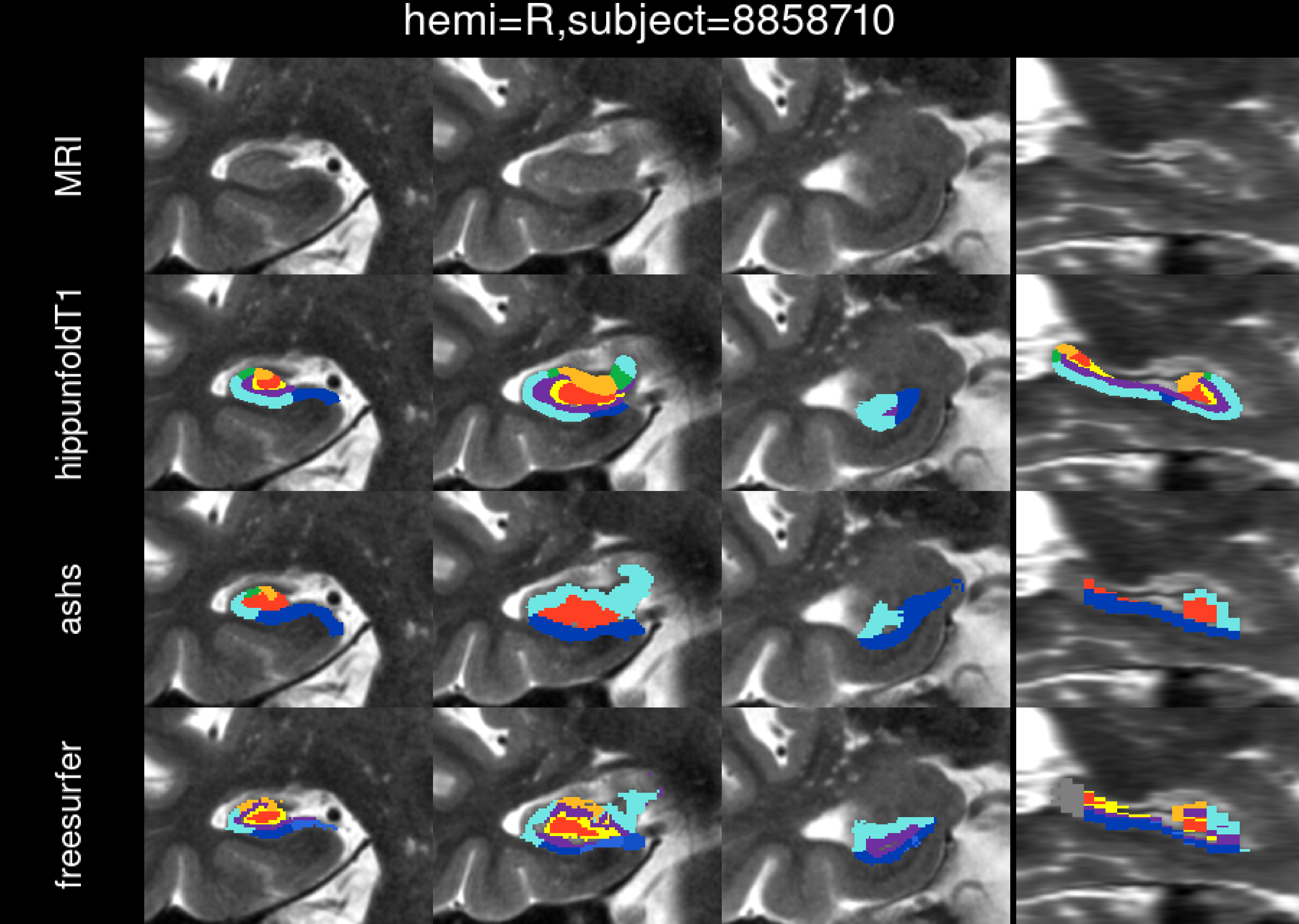
hemi=R,subject=8858710
MRI
hippunfoldT1
ashs
freesurfer

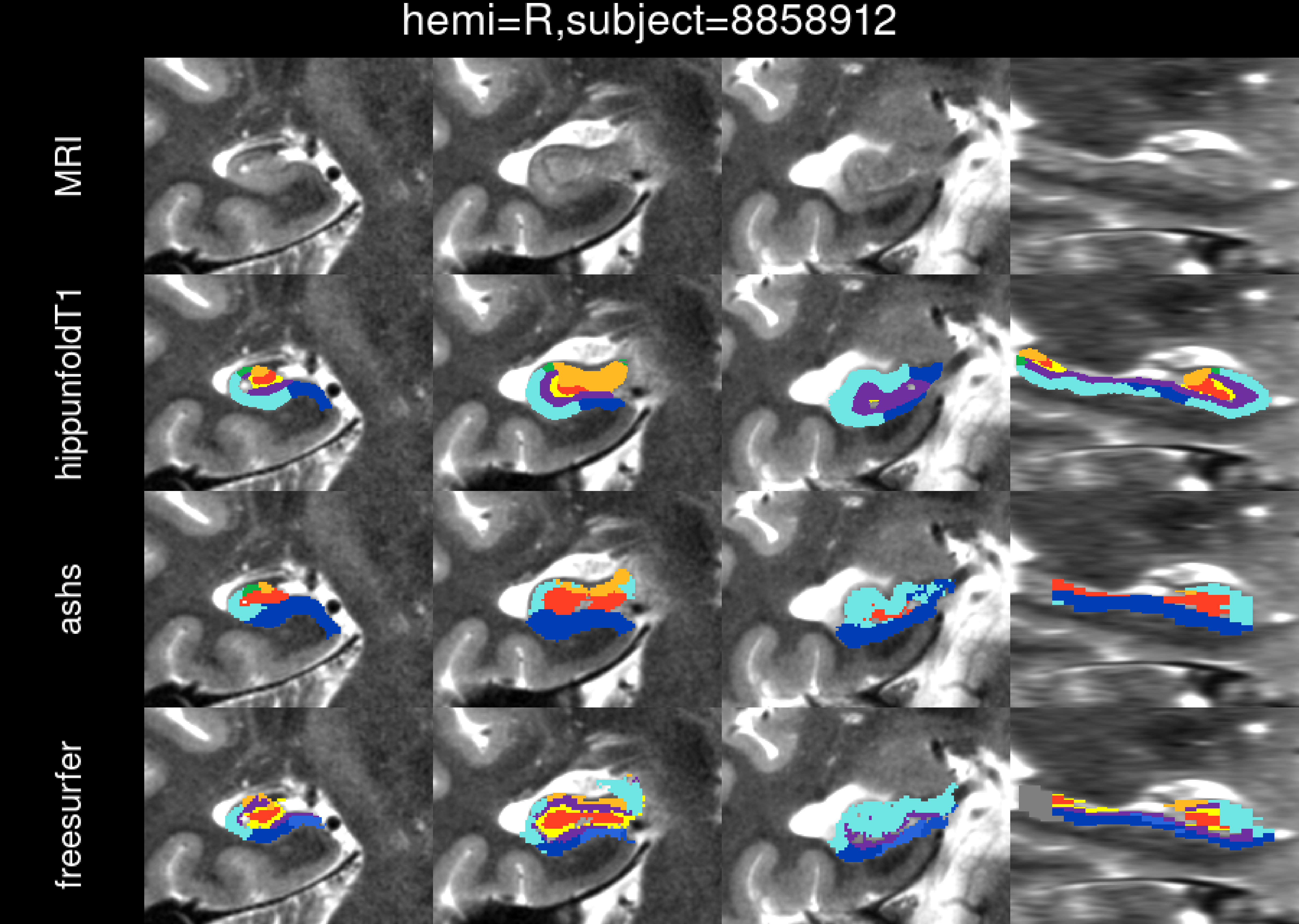
hemi=R,subject=8858912
MRI
hippunfoldT1
ashs
freesurfer

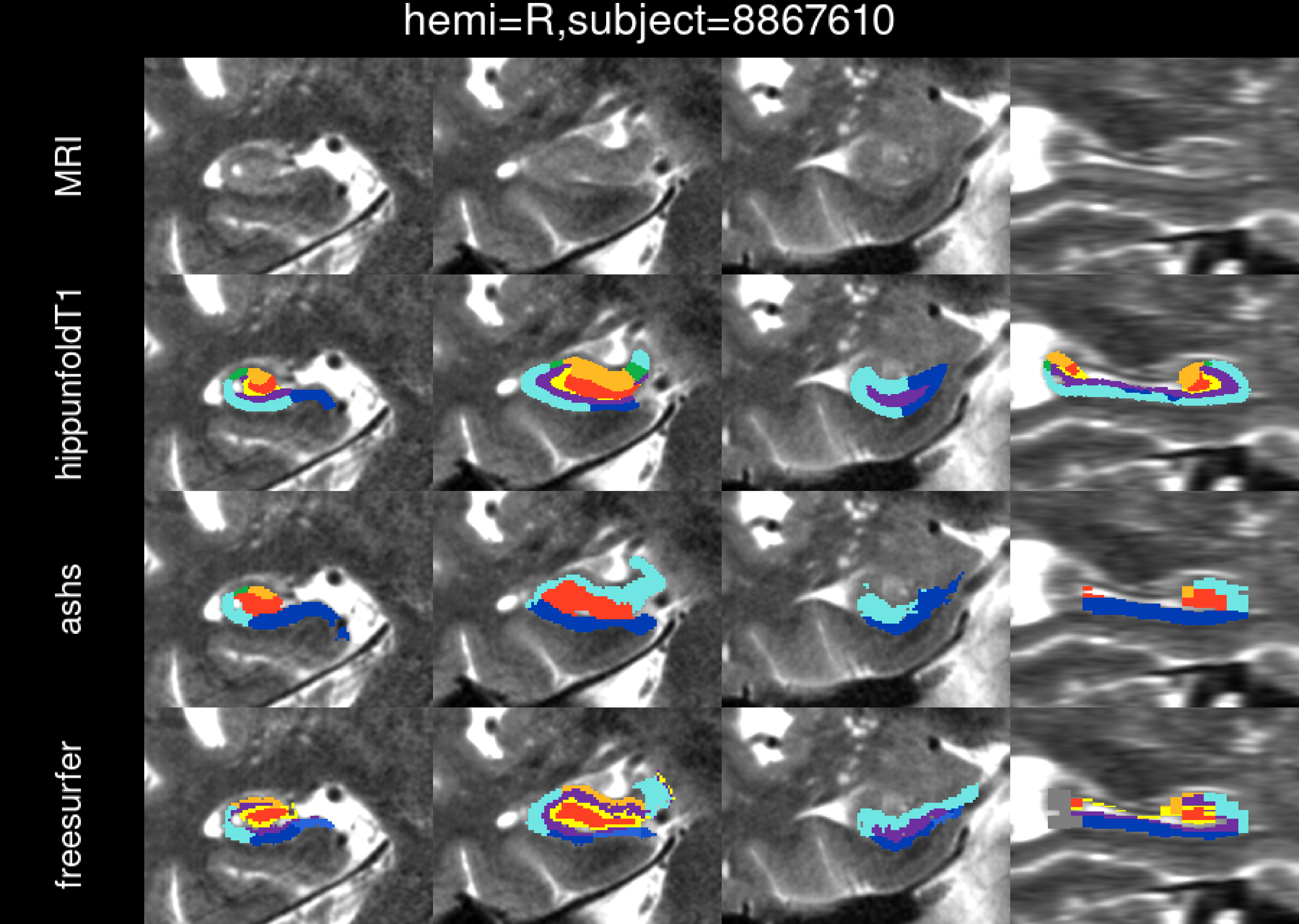
hemi=R,subject=8867610
MRI
hippunfoldT1
ashs
freesurfer

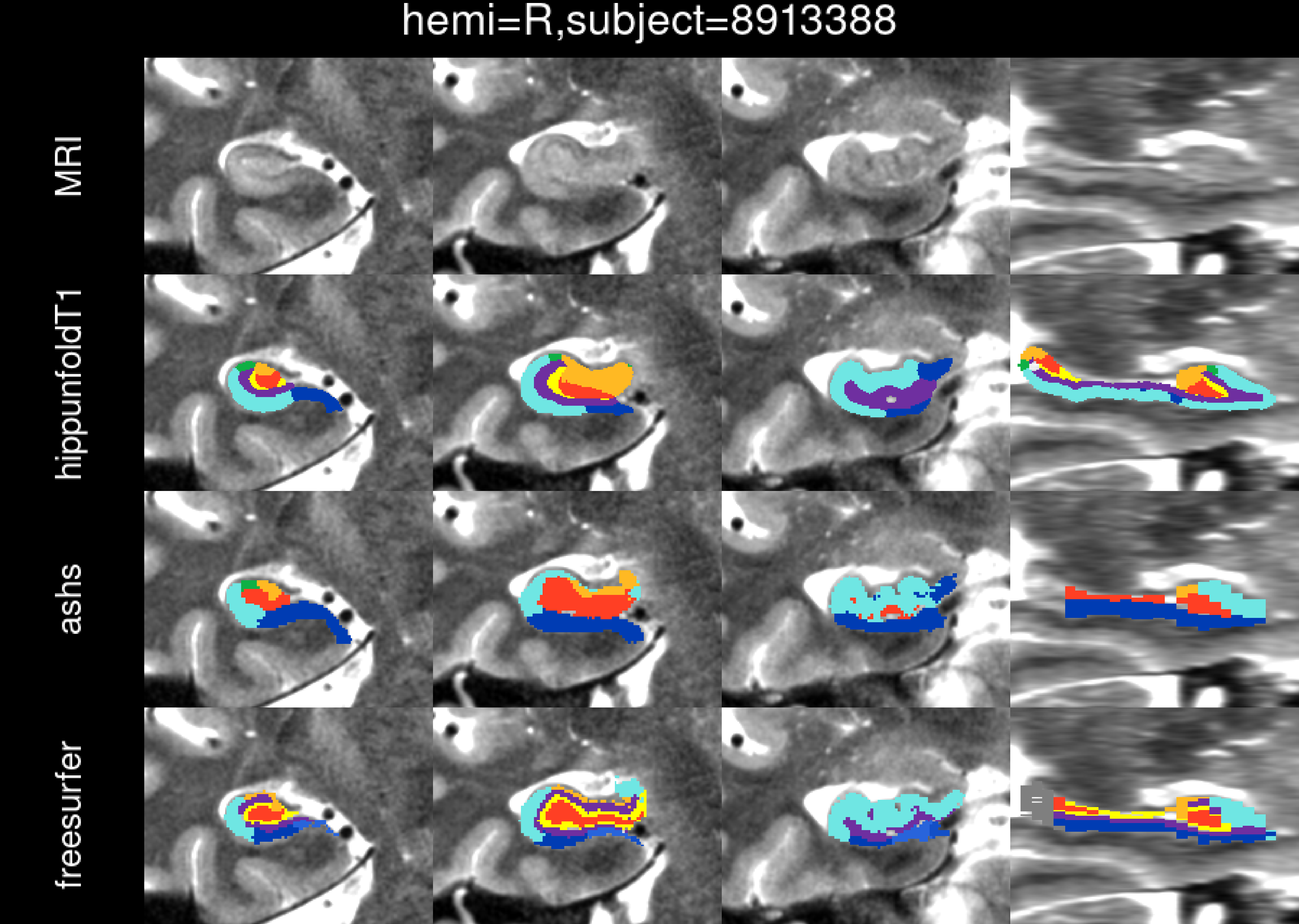
hemi=R,subject=8913388
MRI
hippunfoldT1
ashs
freesurfer

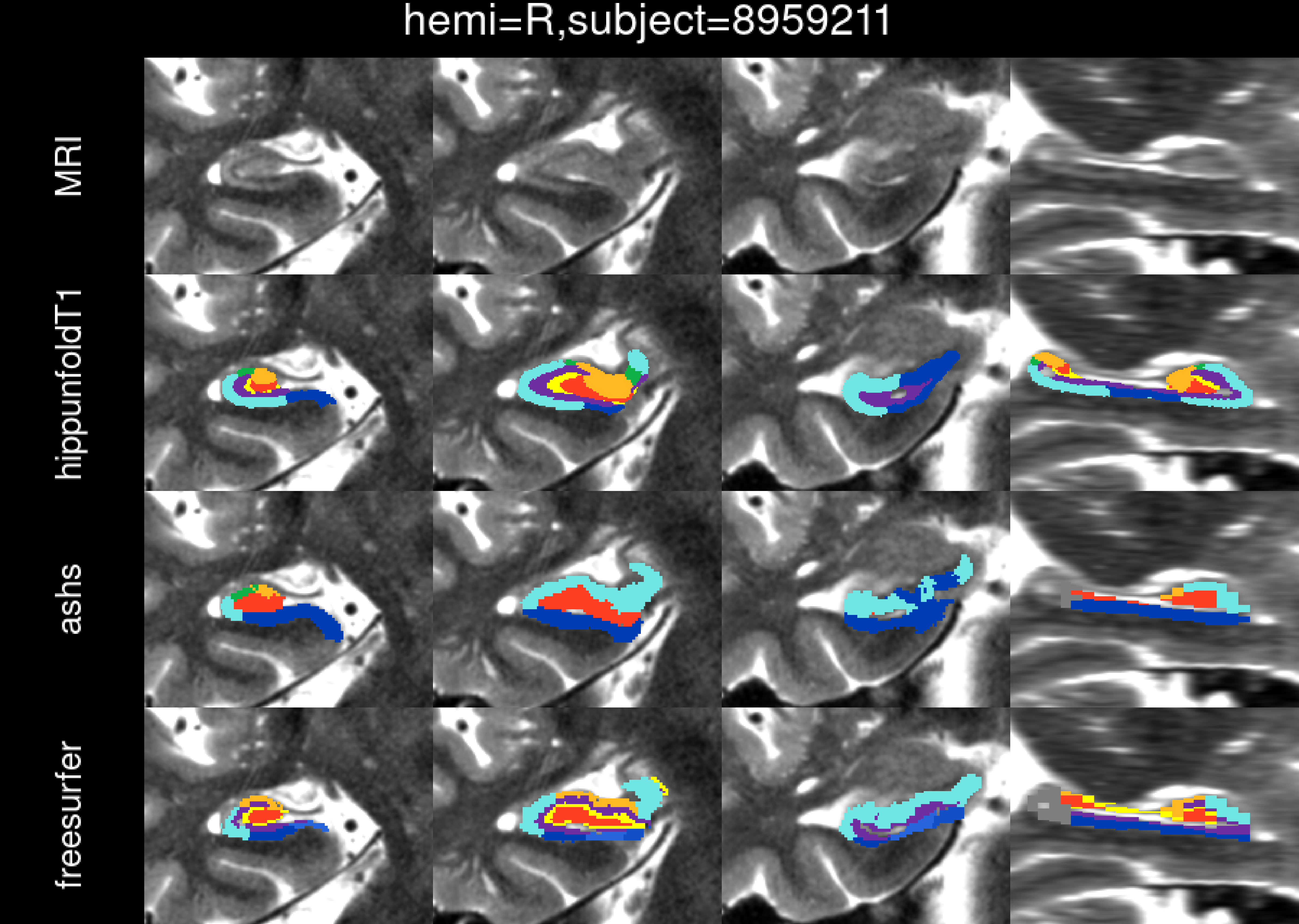
hemi=R,subject=8959211
MRI
hippunfoldT1
ashs
freesurfer

hemi=R,subject=9086990

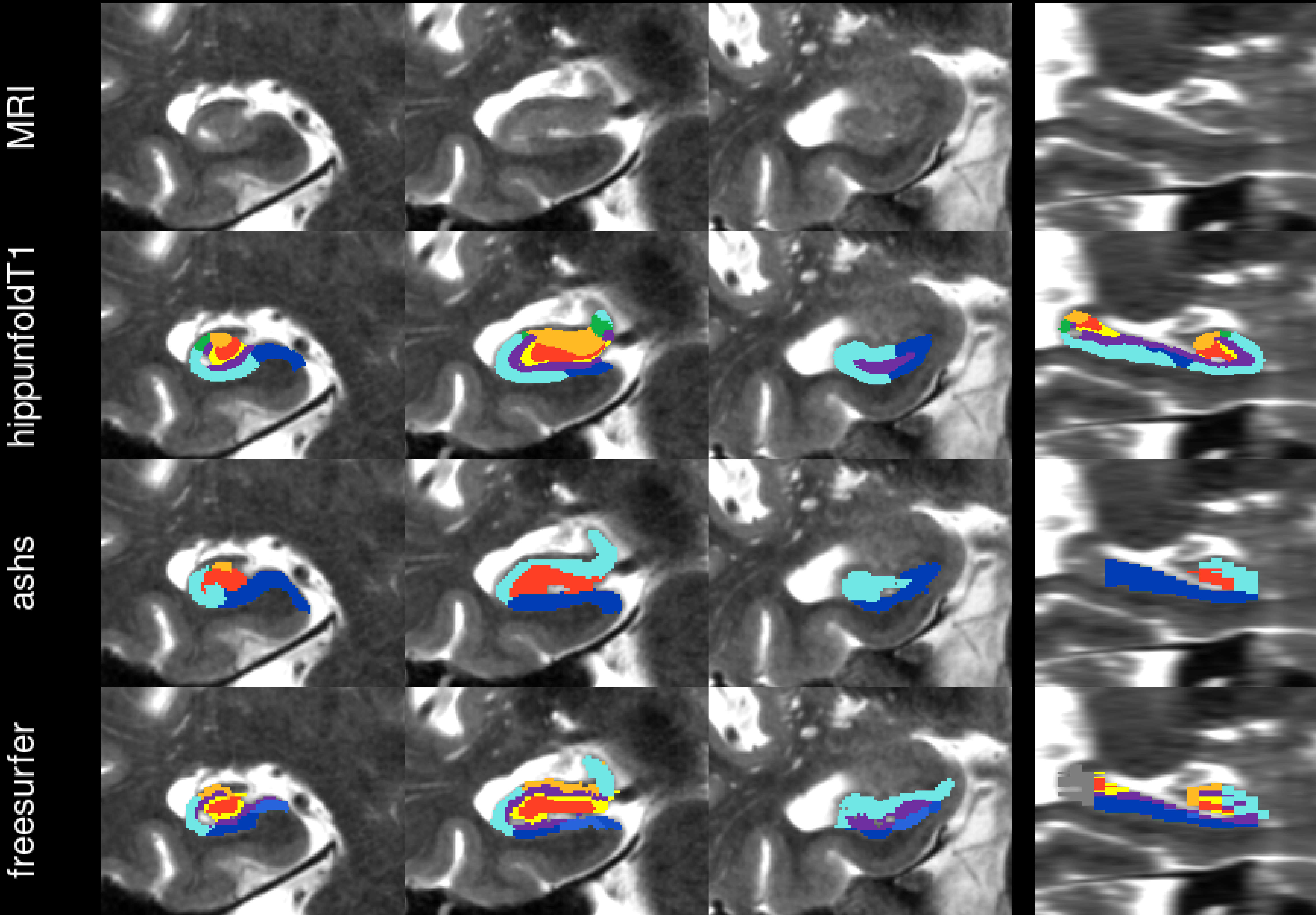

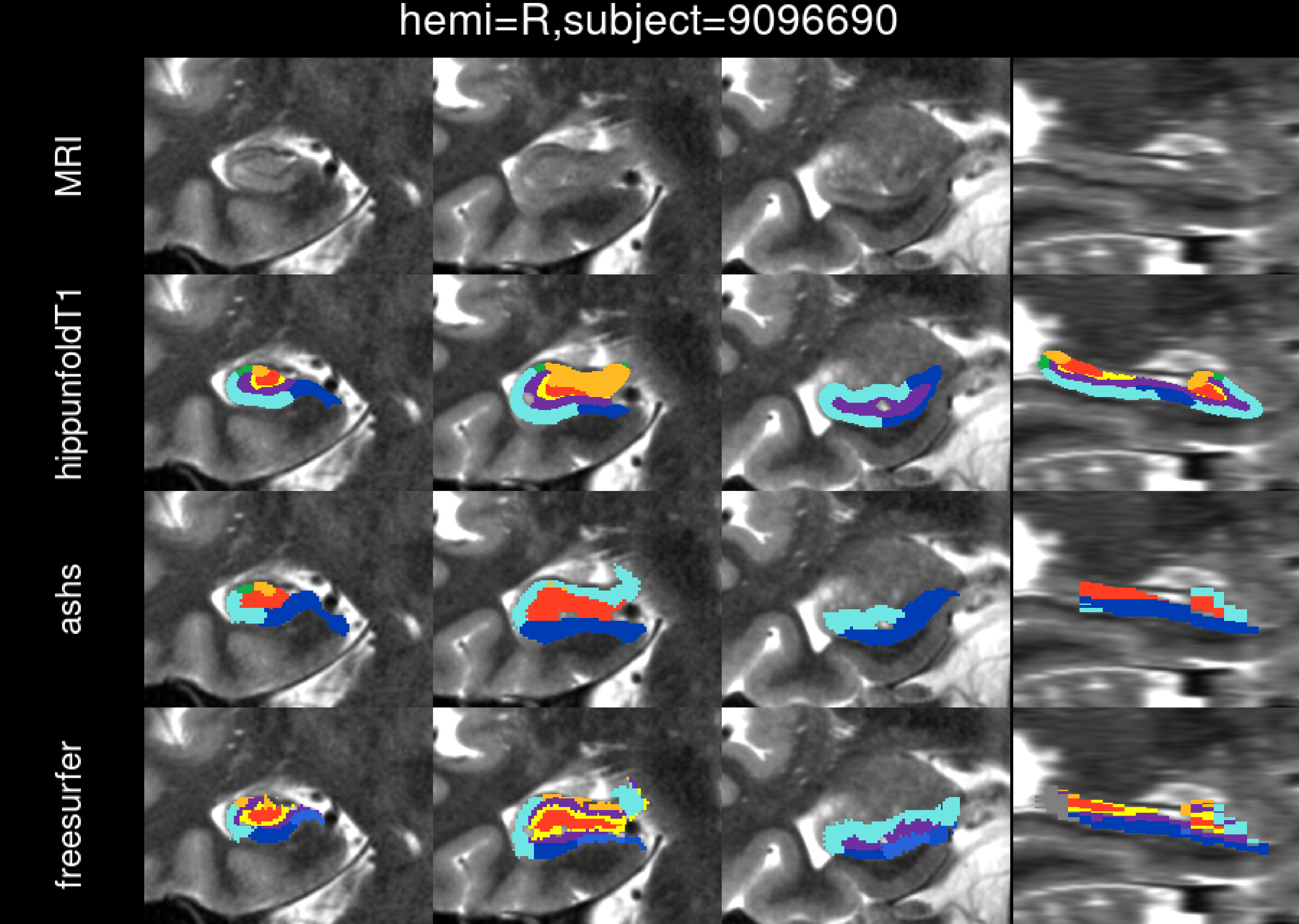
hemi=R,subject=9096690
MRI
hippunfoldT1
ashs
freesurfer

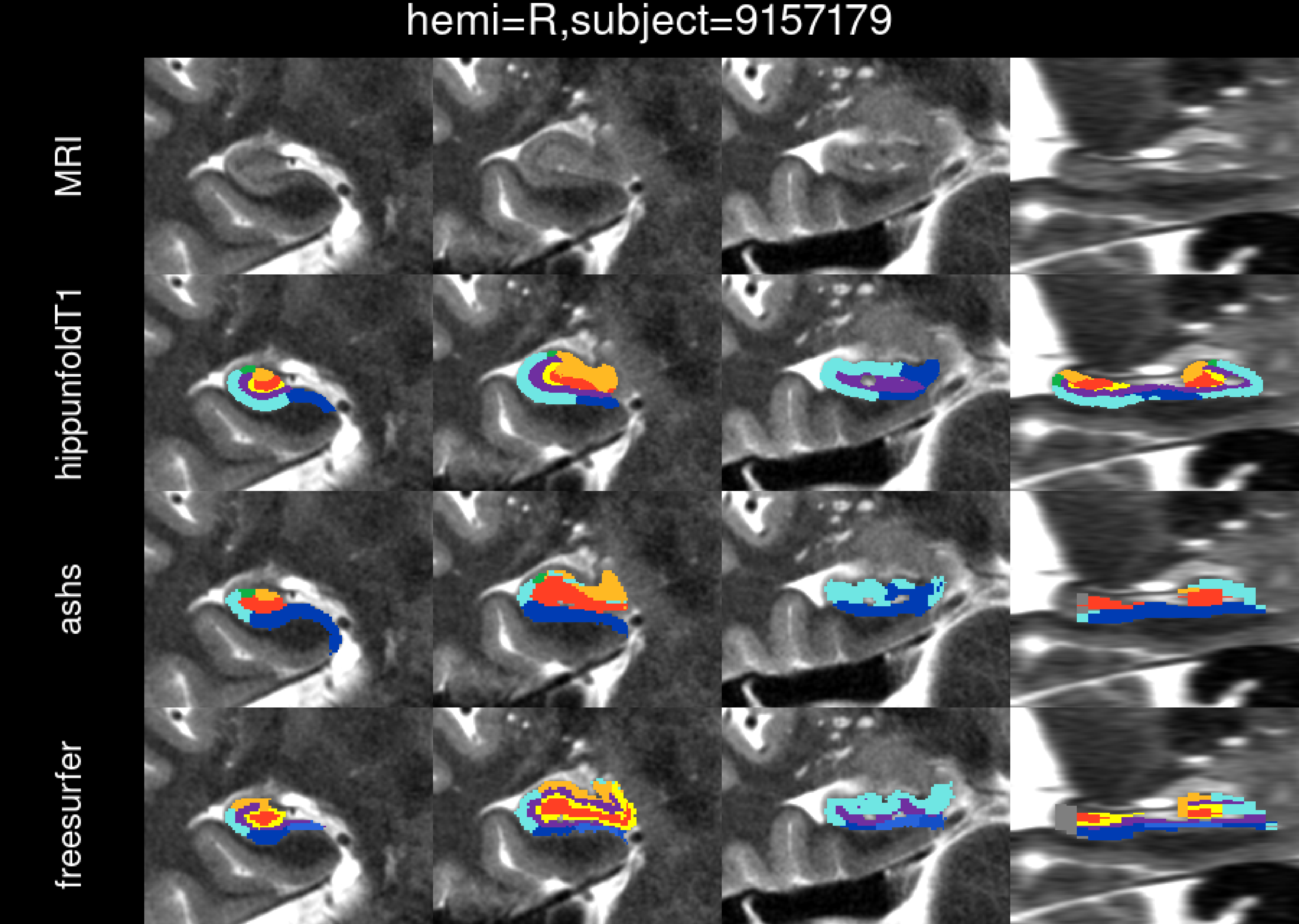

hemi=R,subject=9157179
MRI
hippunfoldT1
ashs
freesurfer

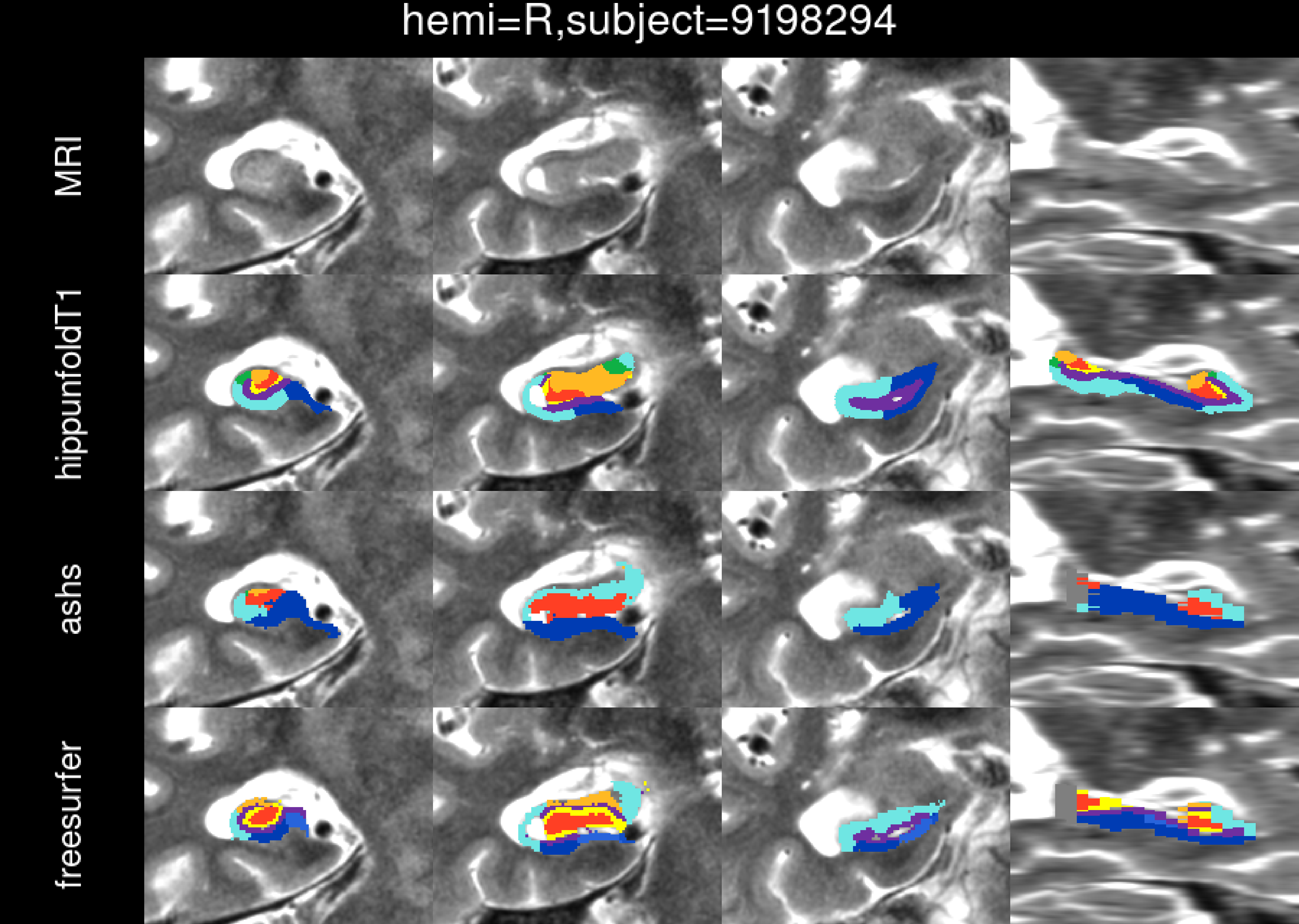

hemi=R,subject=9198294
MRI
hippunfoldT1
ashs
freesurfer

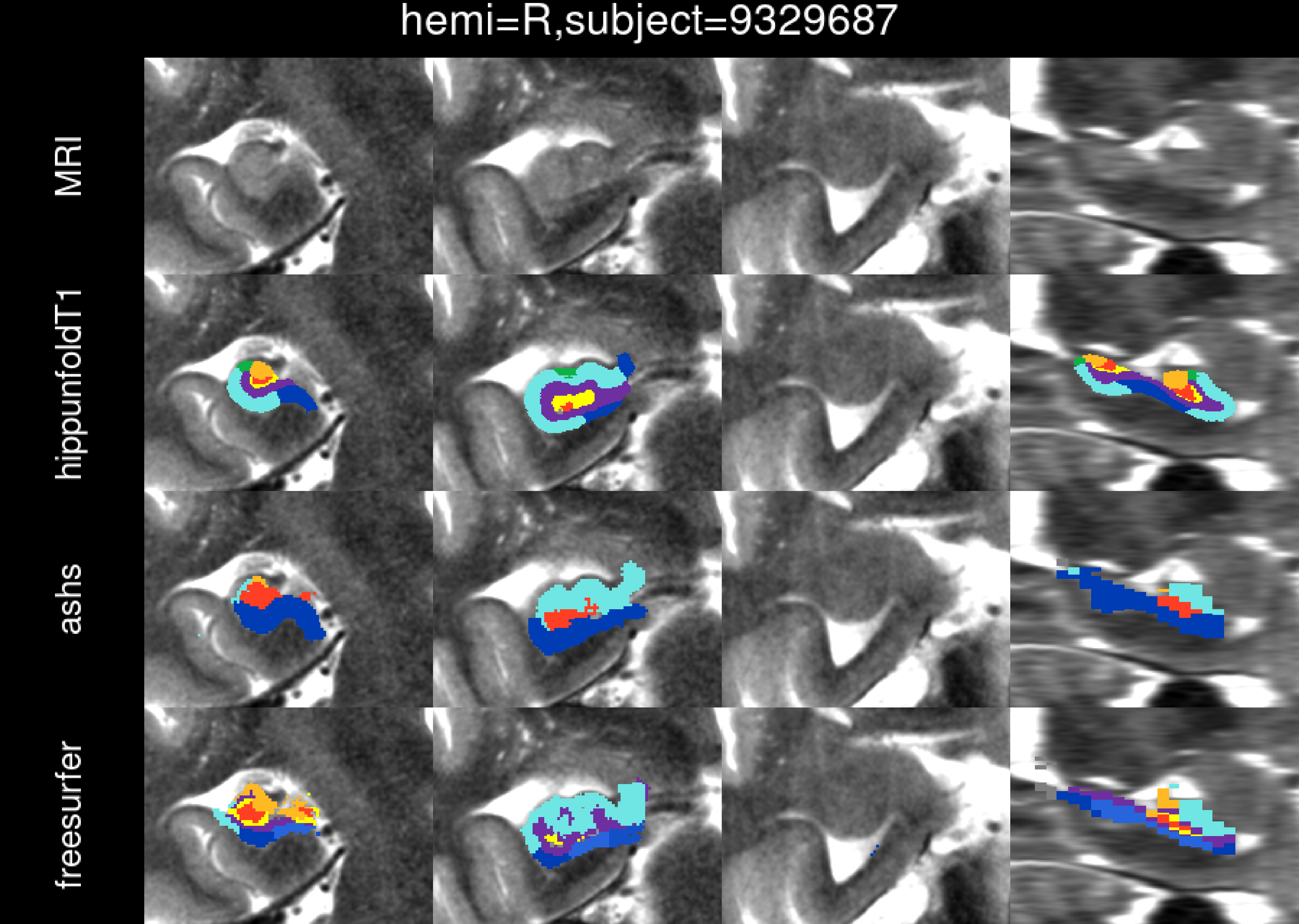
hemi=R,subject=9329687
MRI
hippunfoldT1
ashs
freesurfer

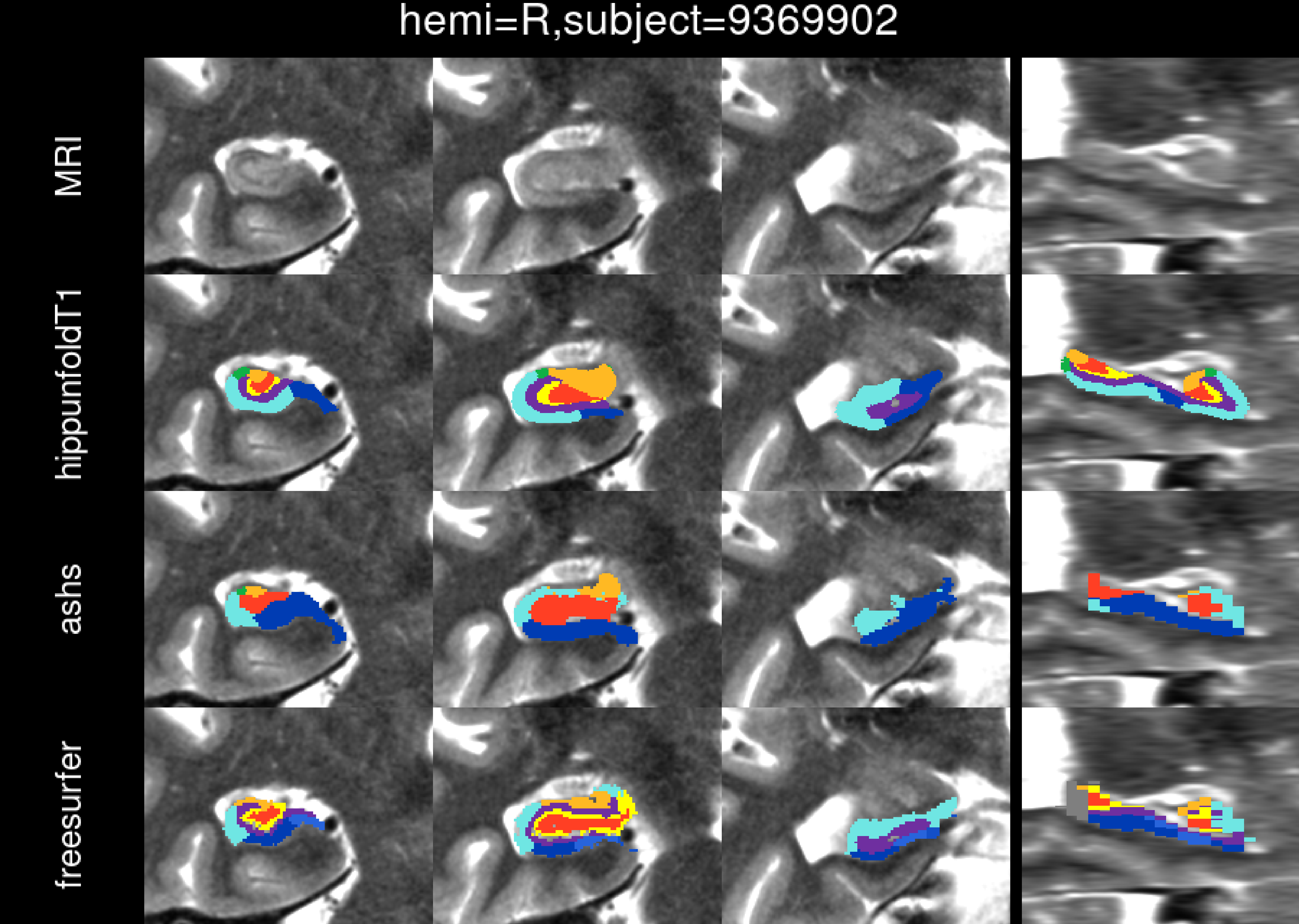

hemi=R,subject=9369902
MRI
hippunfoldT1
ashs
freesurfer

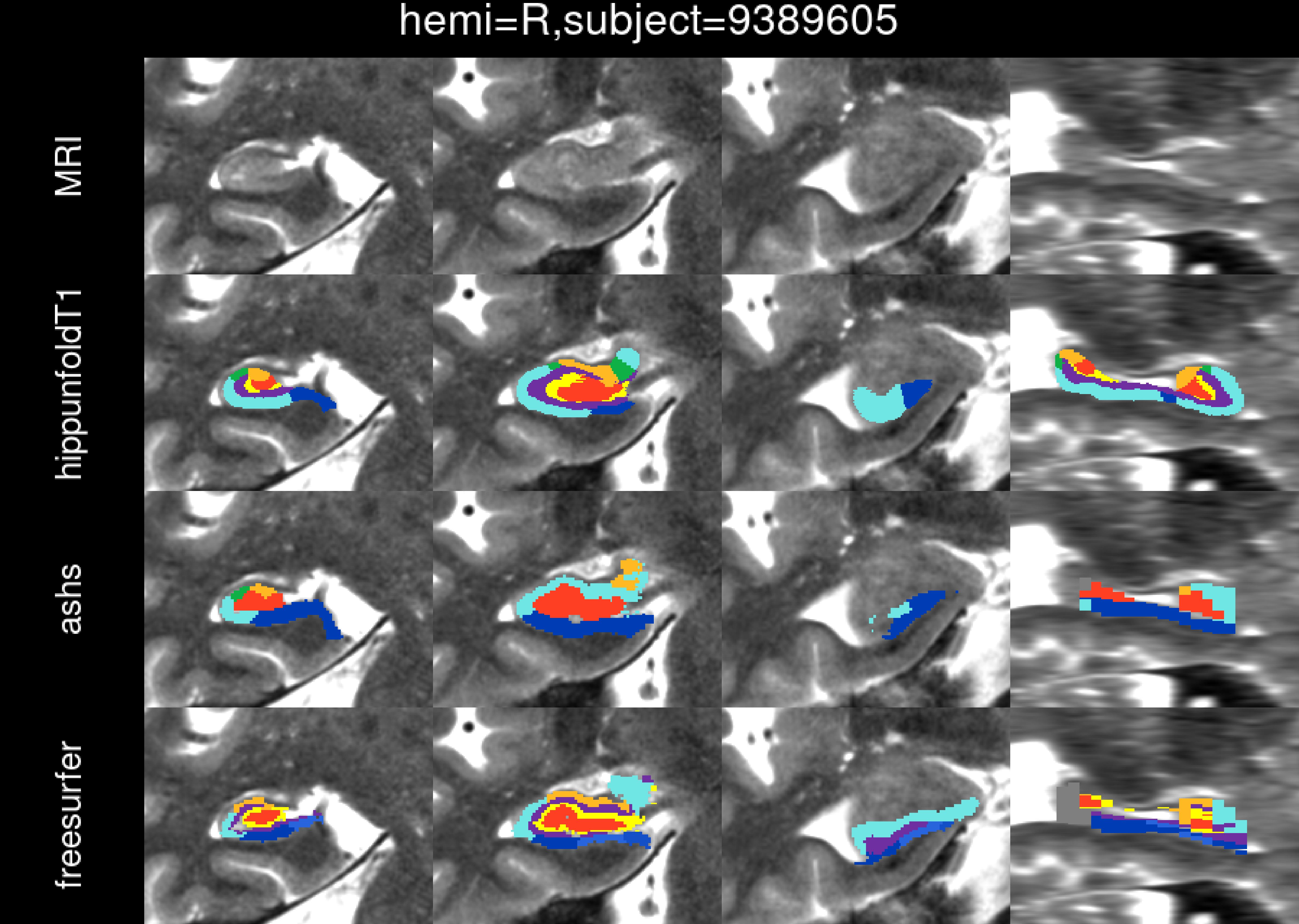
hemi=R,subject=9389605
MRI
hippunfoldT1
ashs
freesurfer

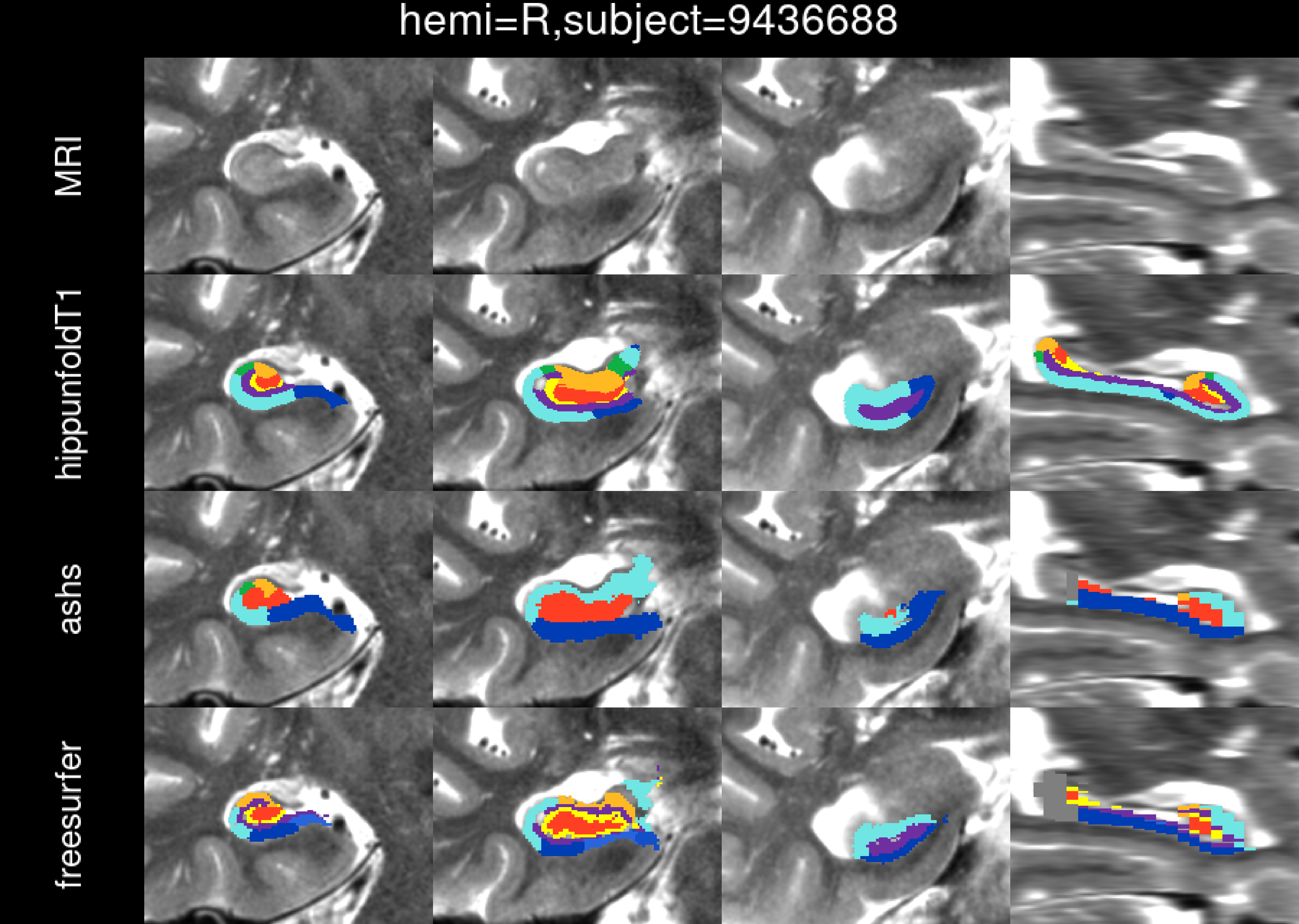

hemi=R,subject=9436688
MRI
hippunfoldT1
ashs
freesurfer

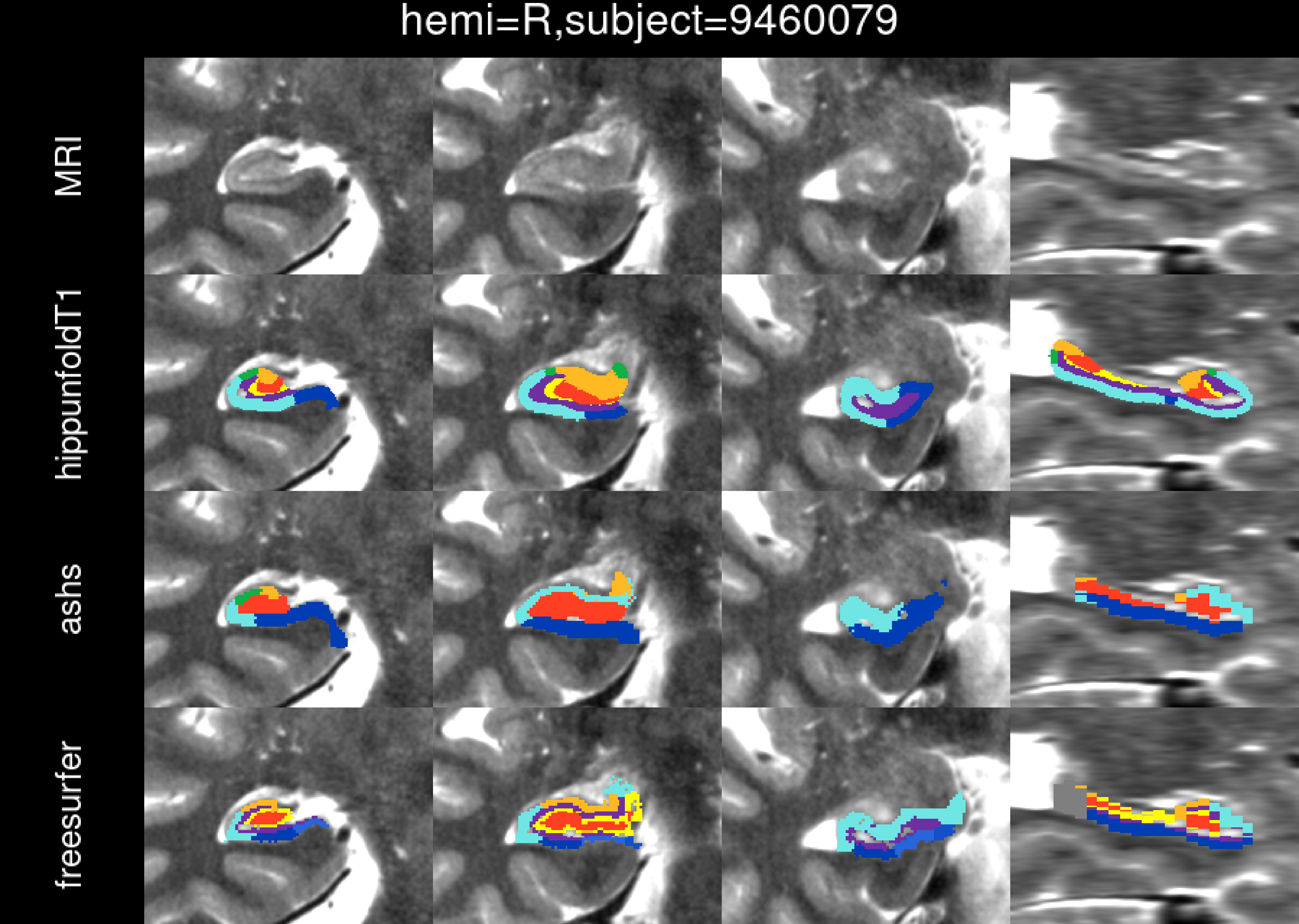

hemi=R,subject=9460079
MRI
hippunfoldT1
ashs
freesurfer

hemi=R,subject=9481794

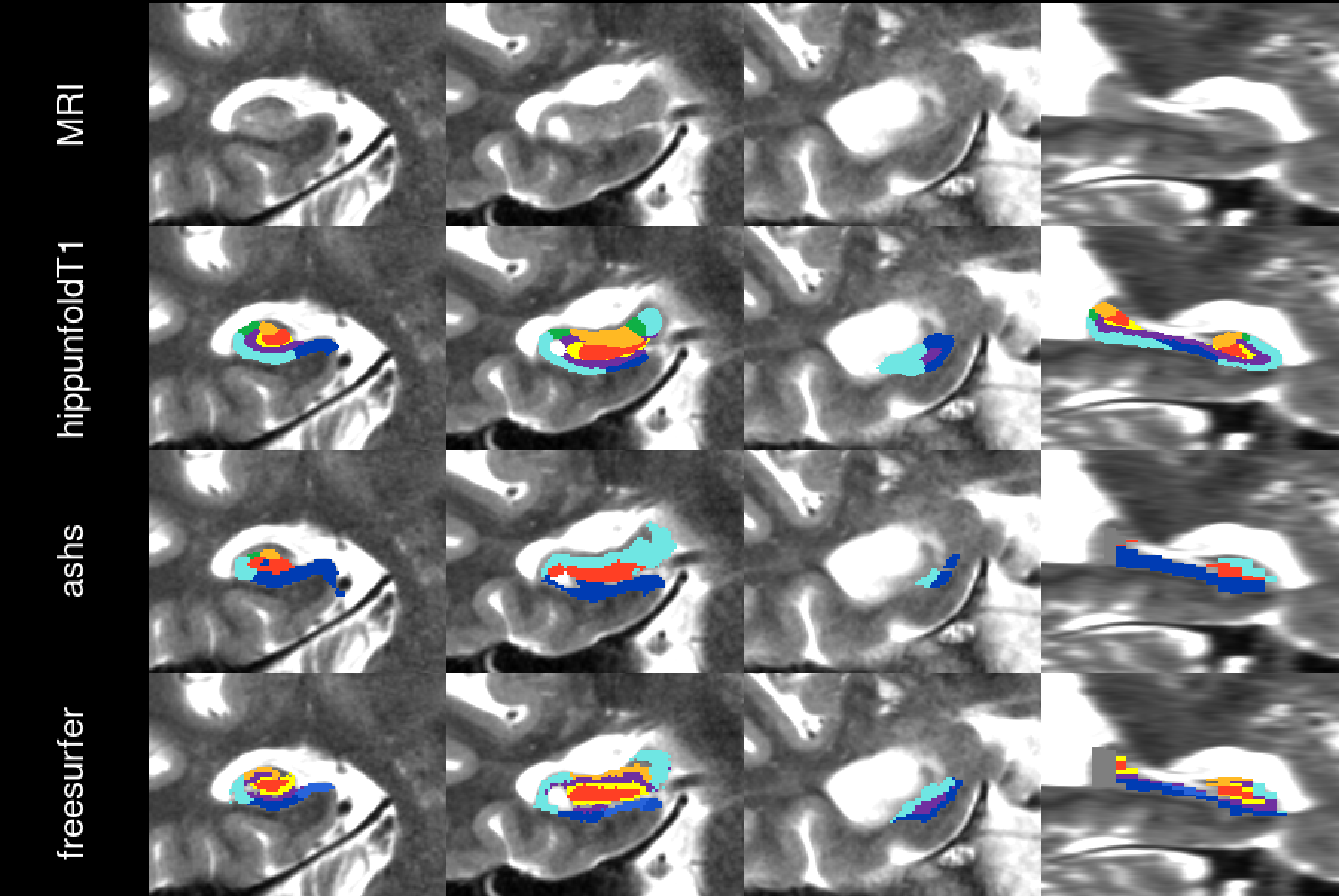

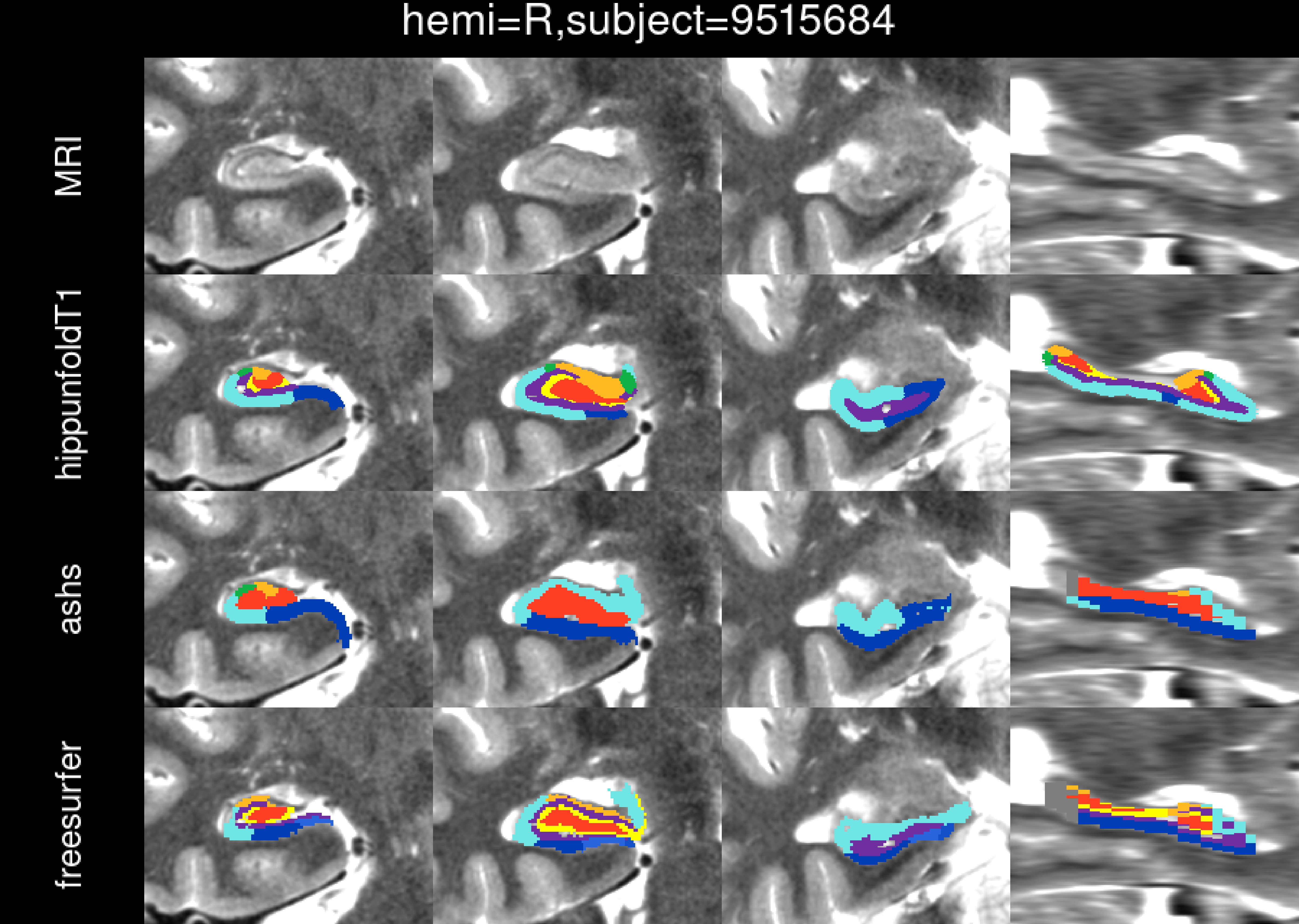
hemi=R,subject=9515684
MRI
hippunfoldT1
ashs
freesurfer

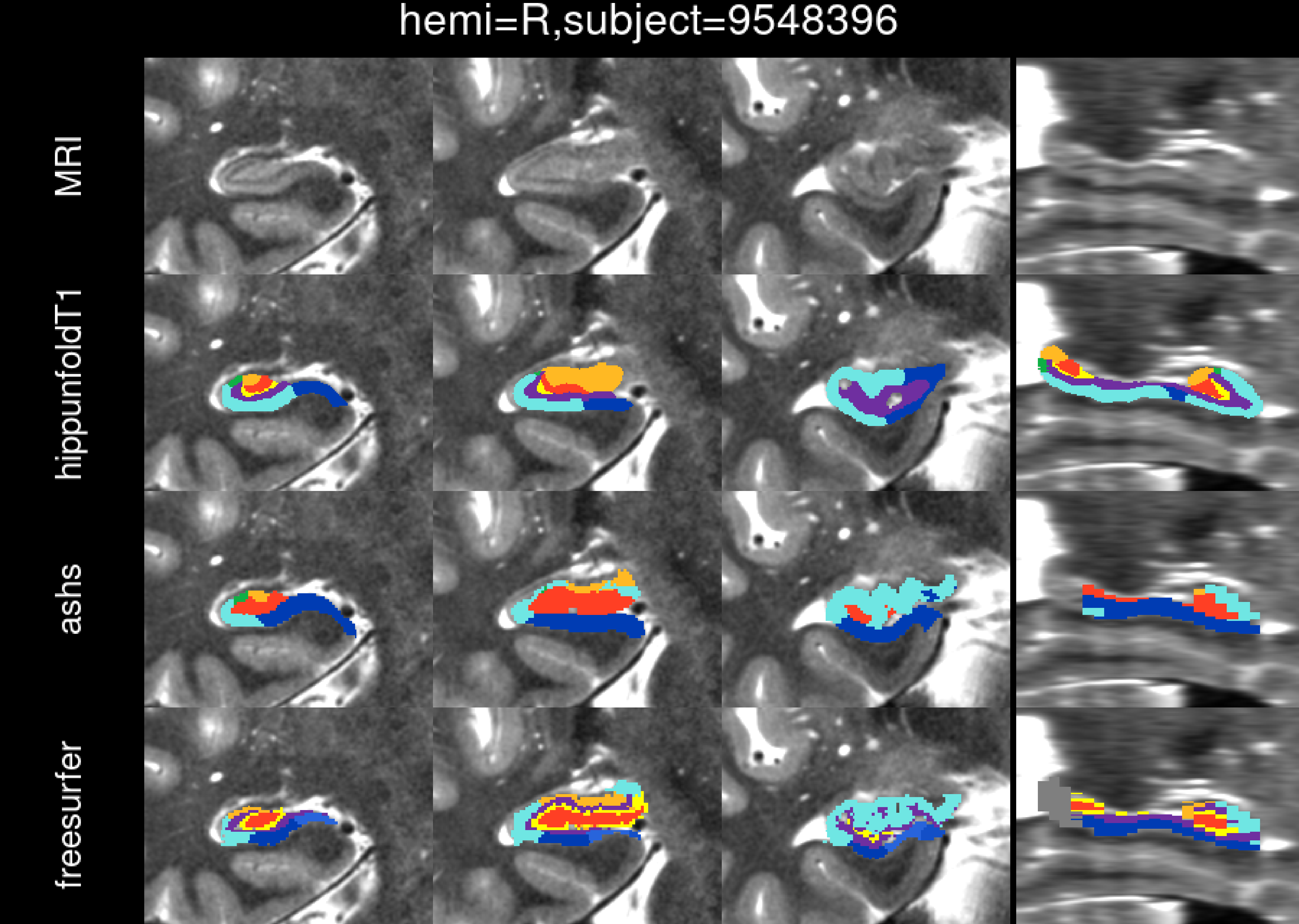
hemi=R,subject=9548396
MRI
hippunfoldT1
ashs
freesurfer

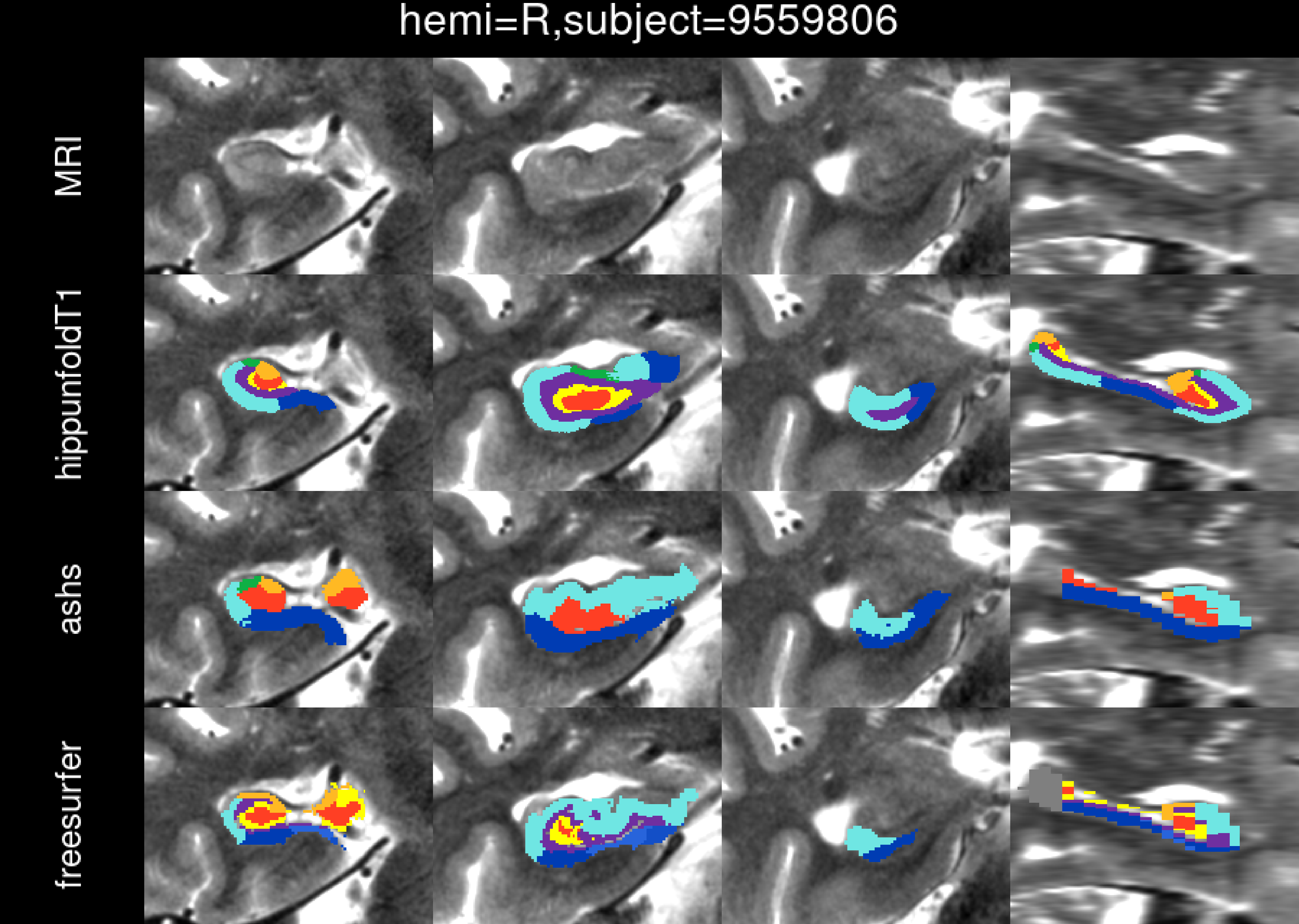
hemi=R,subject=9559806
MRI
hippunfoldT1
ashs
freesurfer

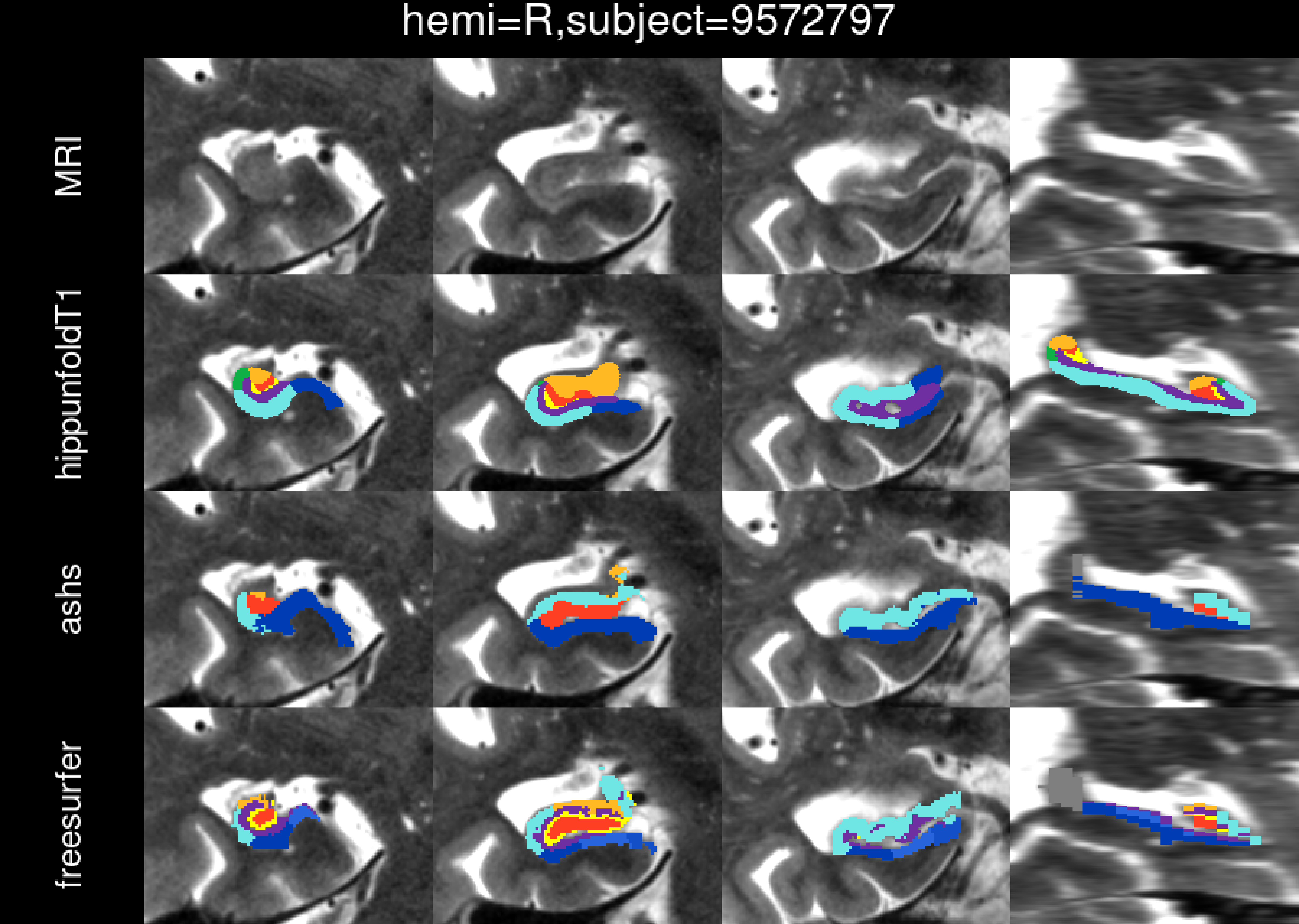
hemi=R,subject=9572797
MRI
hippunfoldT1
ashs
freesurfer

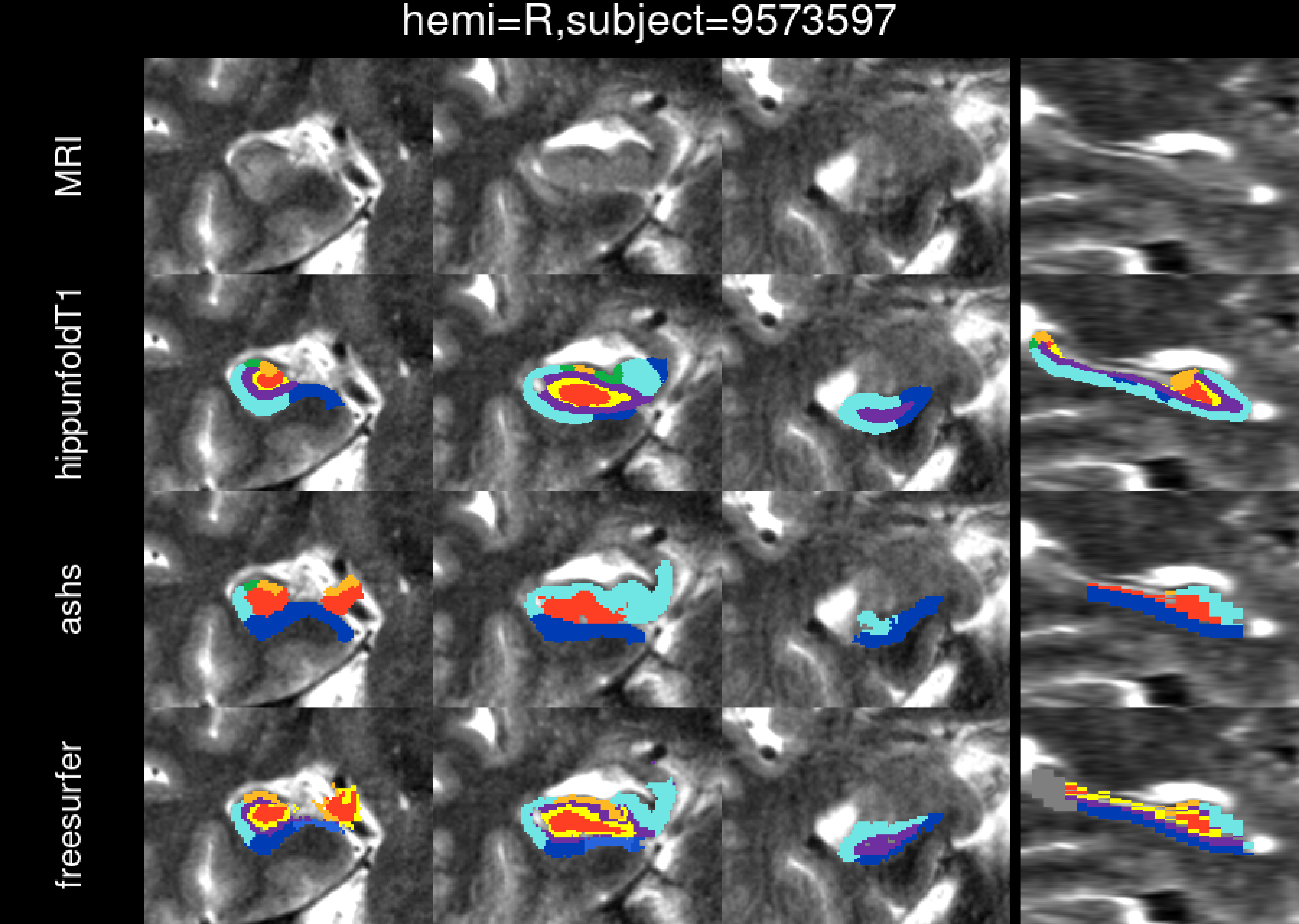
hemi=R,subject=9573597
MRI
hippunfoldT1
ashs
freesurfer

hemi=R,subject=9578406

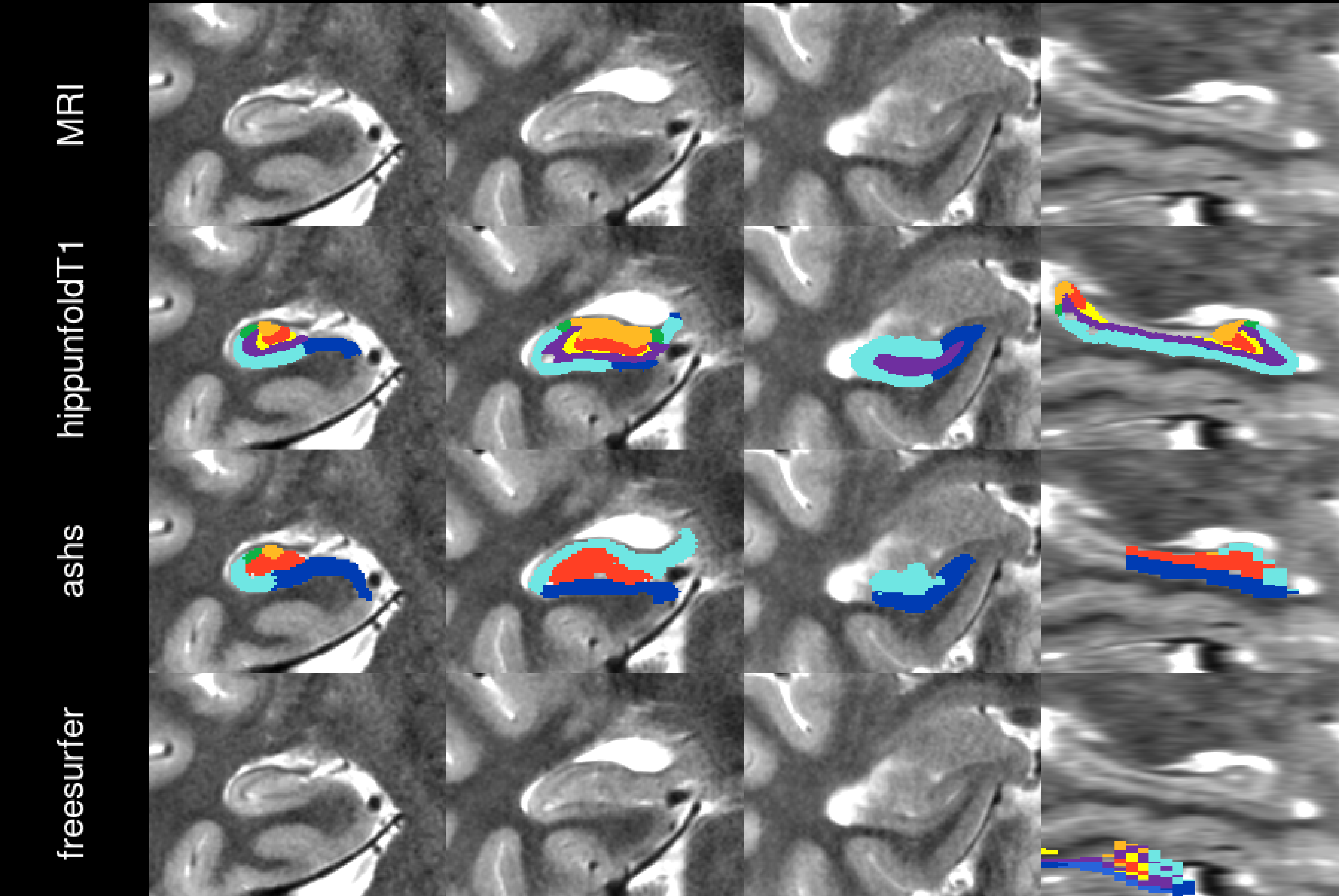

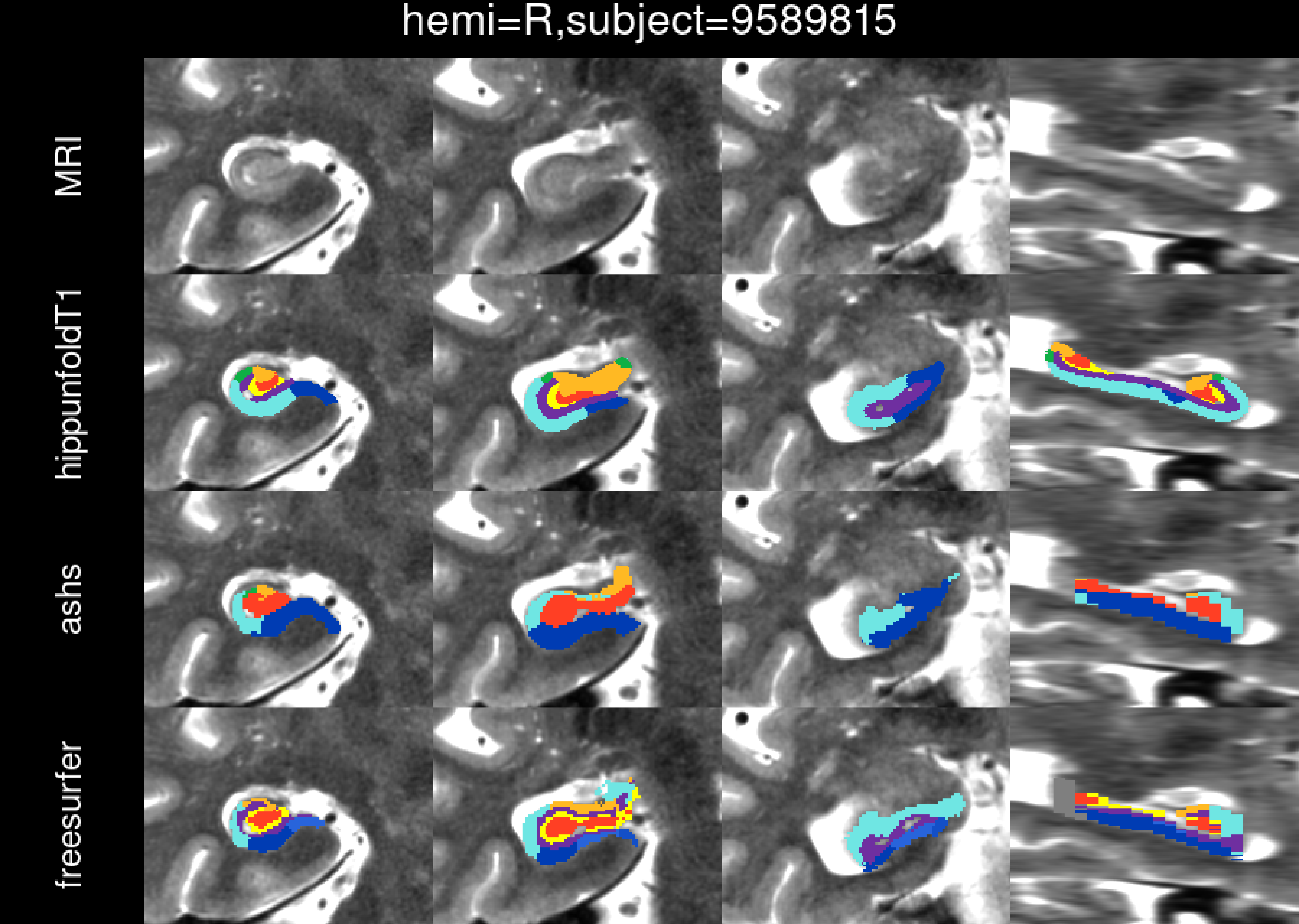
hemi=R,subject=9589815
MRI
hippunfoldT1
ashs
freesurfer

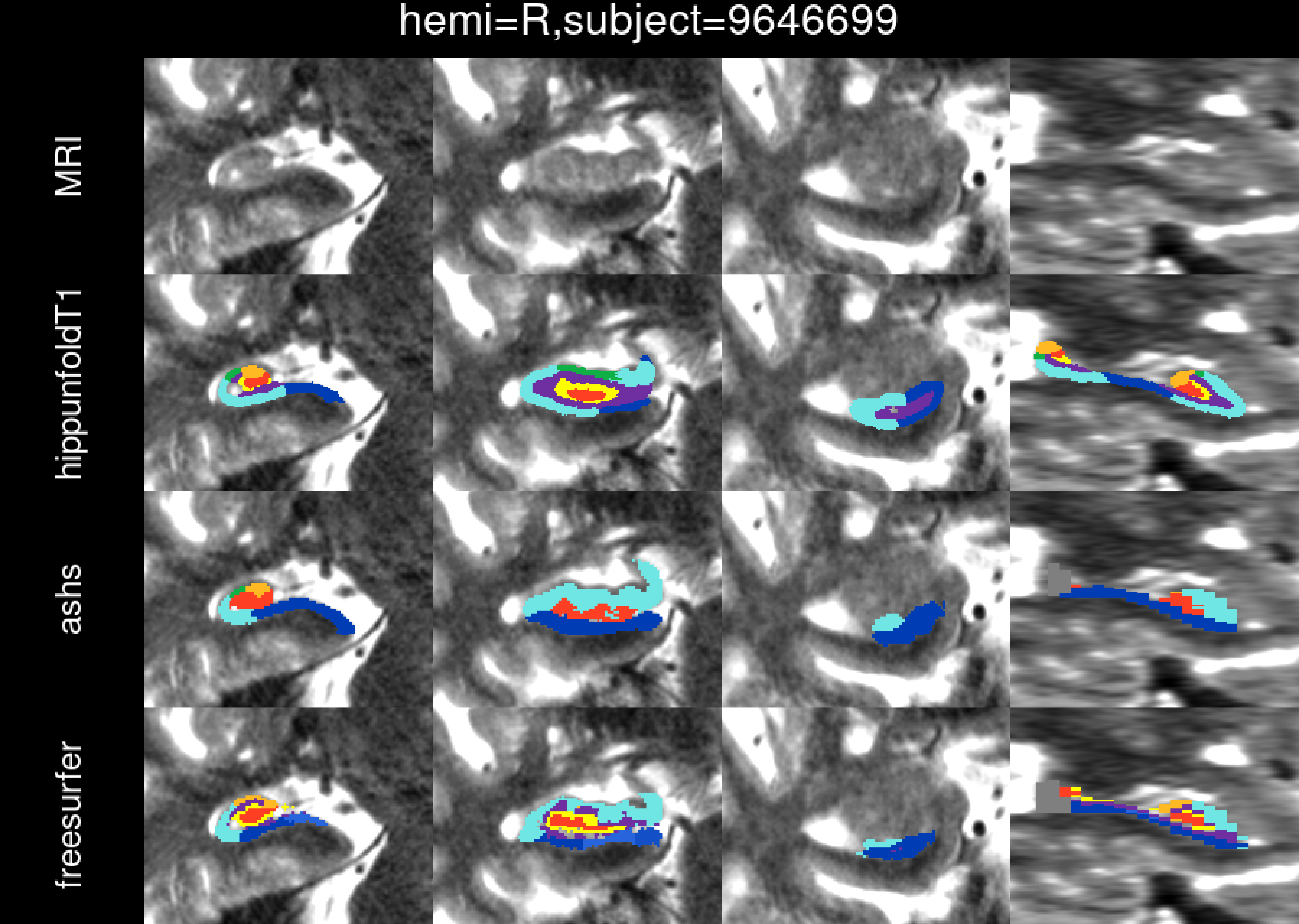
hemi=R,subject=9646699
MRI
hippunfoldT1
ashs
freesurfer

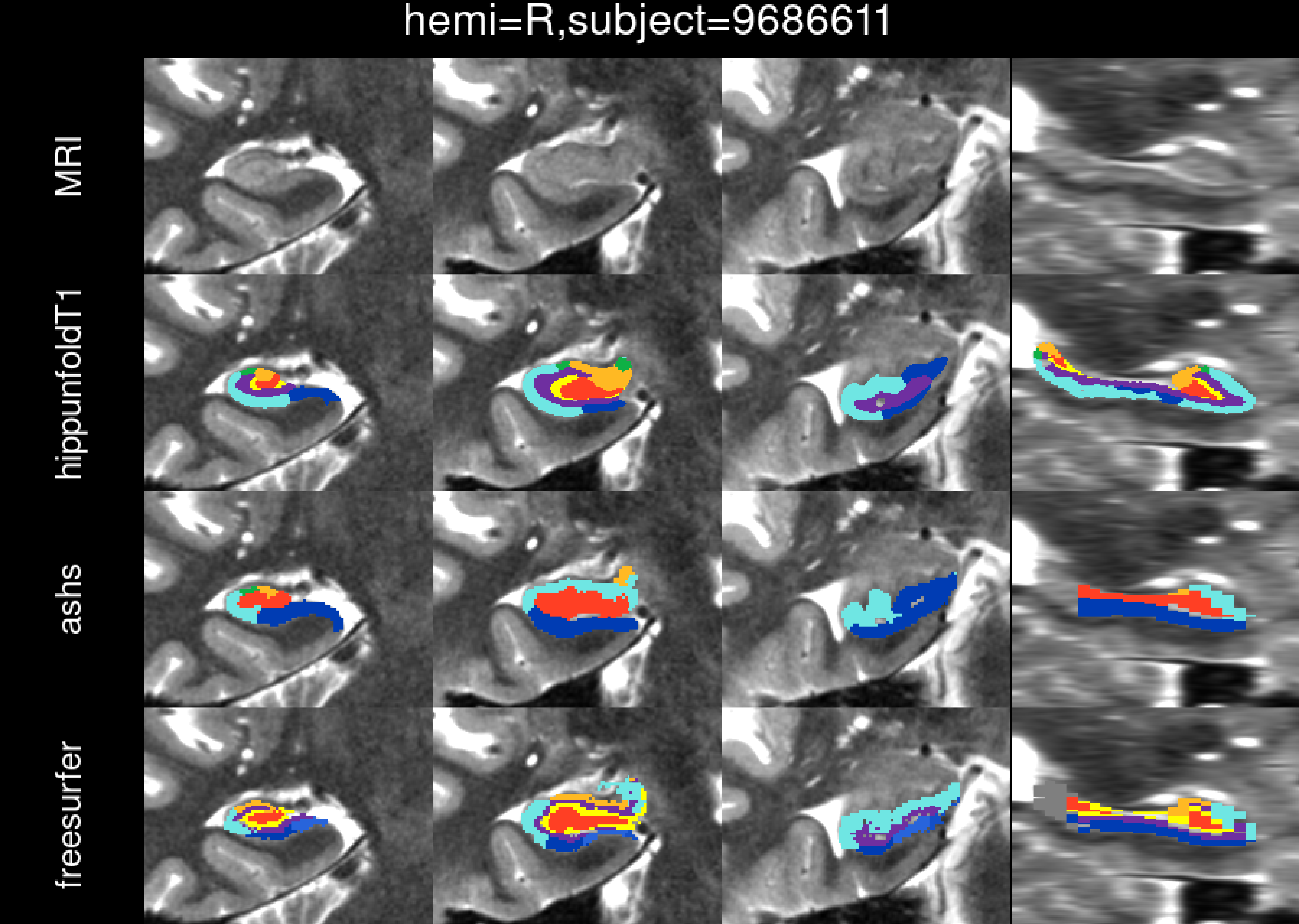
hemi=R,subject=9686611
MRI
hippunfoldT1
ashs
freesurfer

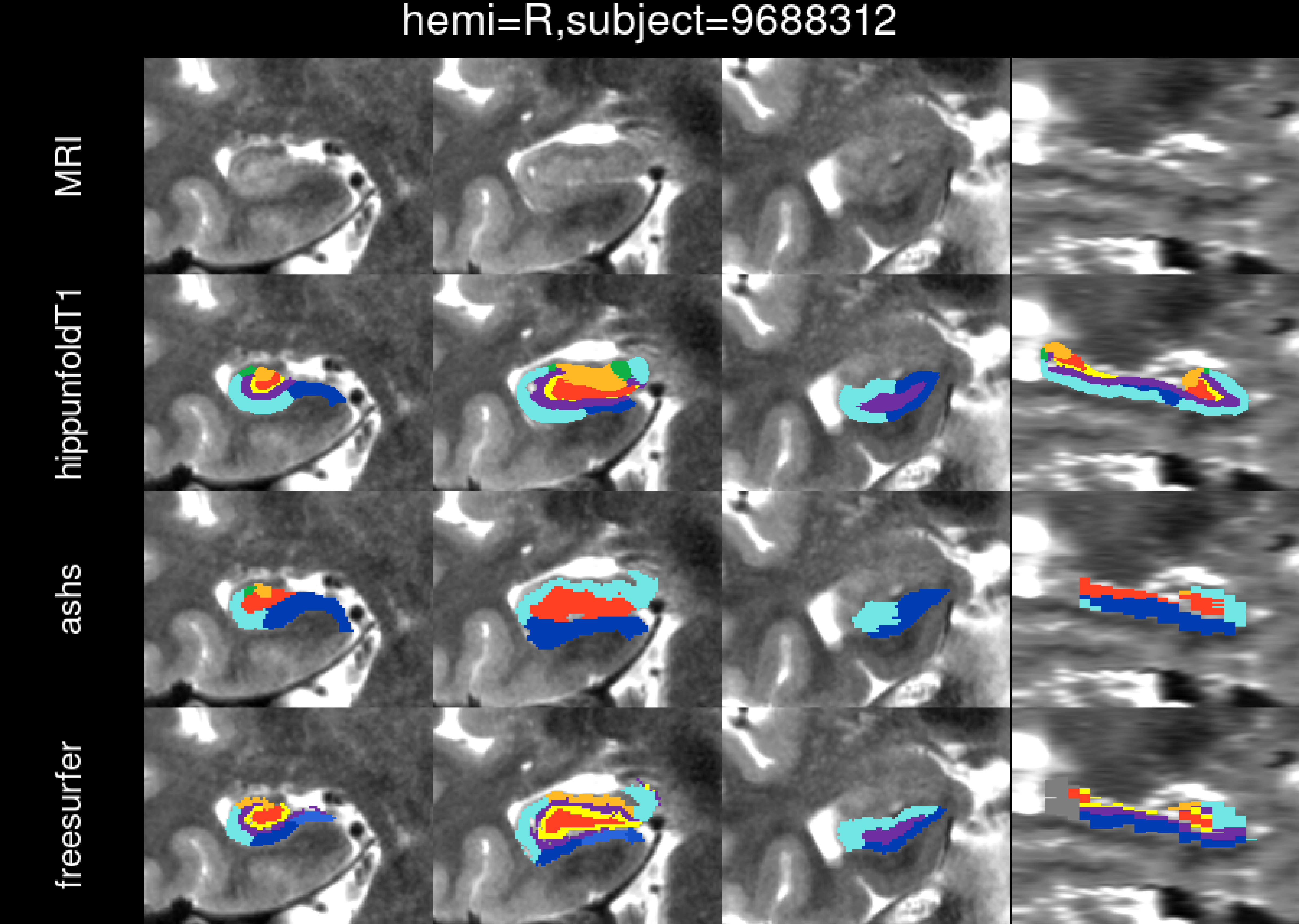
hemi=R,subject=9688312
MRI
hippunfoldT1
ashs
freesurfer

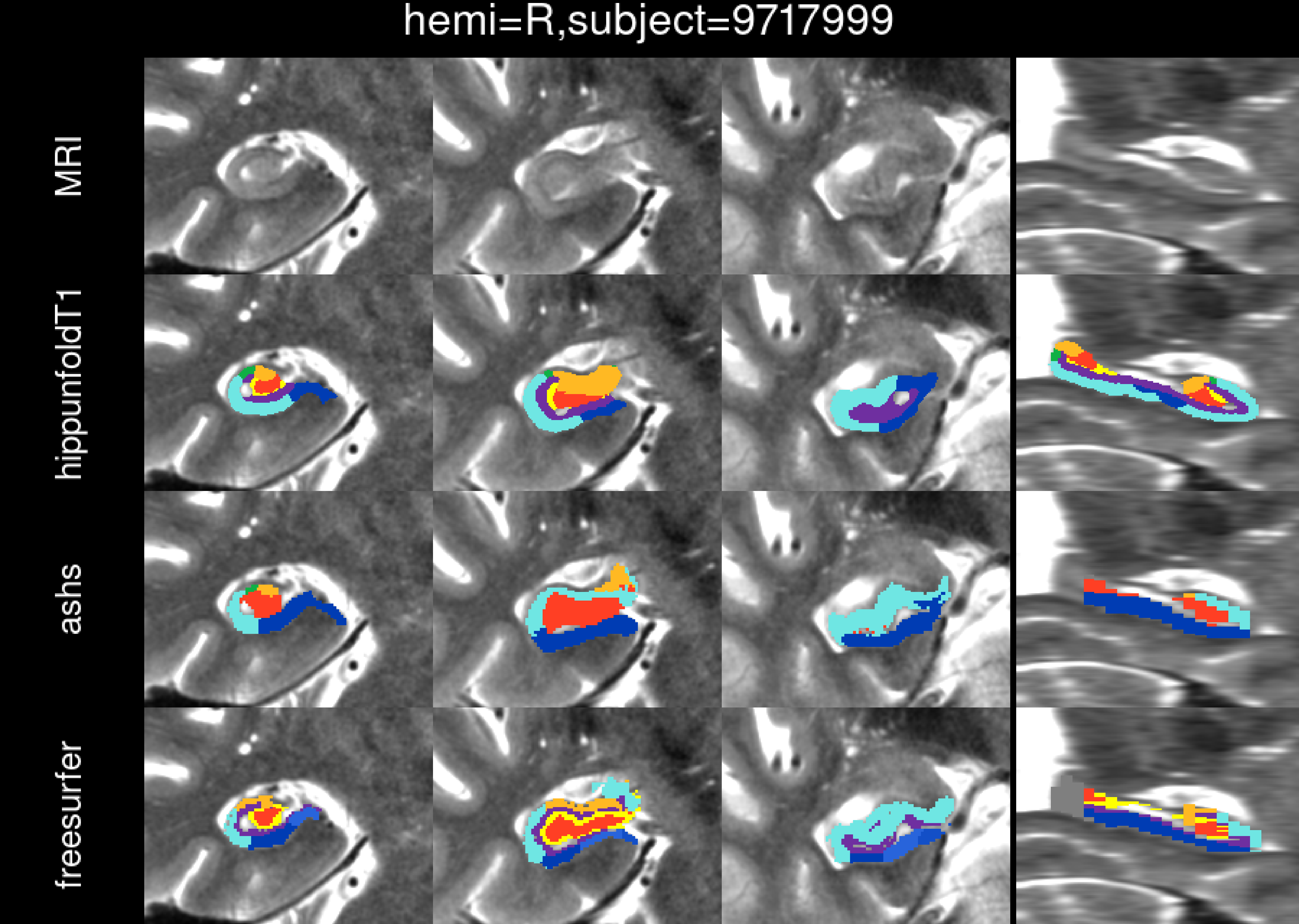

hemi=R,subject=9717999
MRI
hippunfoldT1
ashs
freesurfer

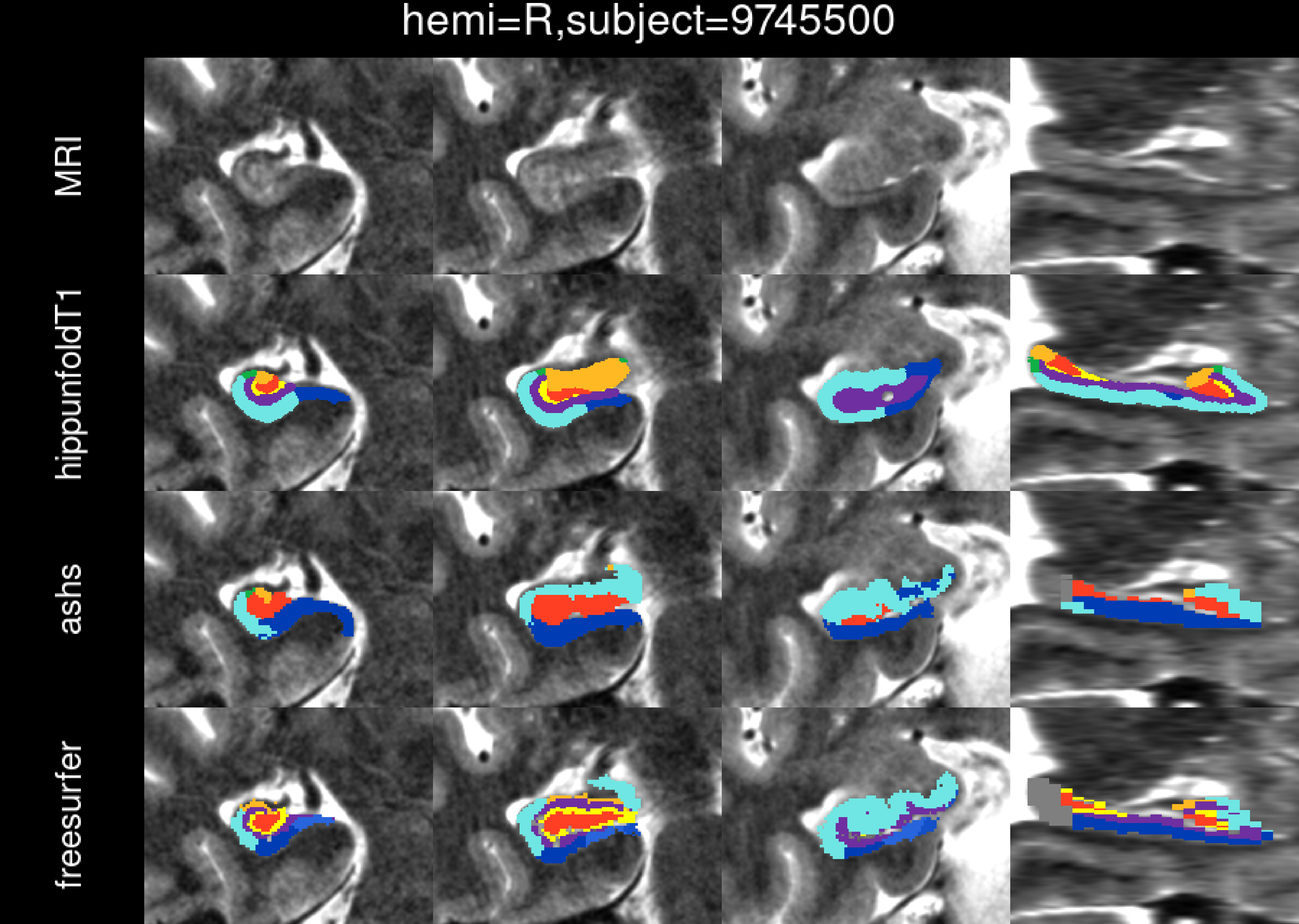
hemi=R,subject=9745500
MRI
hippunfoldT1
ashs
freesurfer

hemi=R,subject=9845403

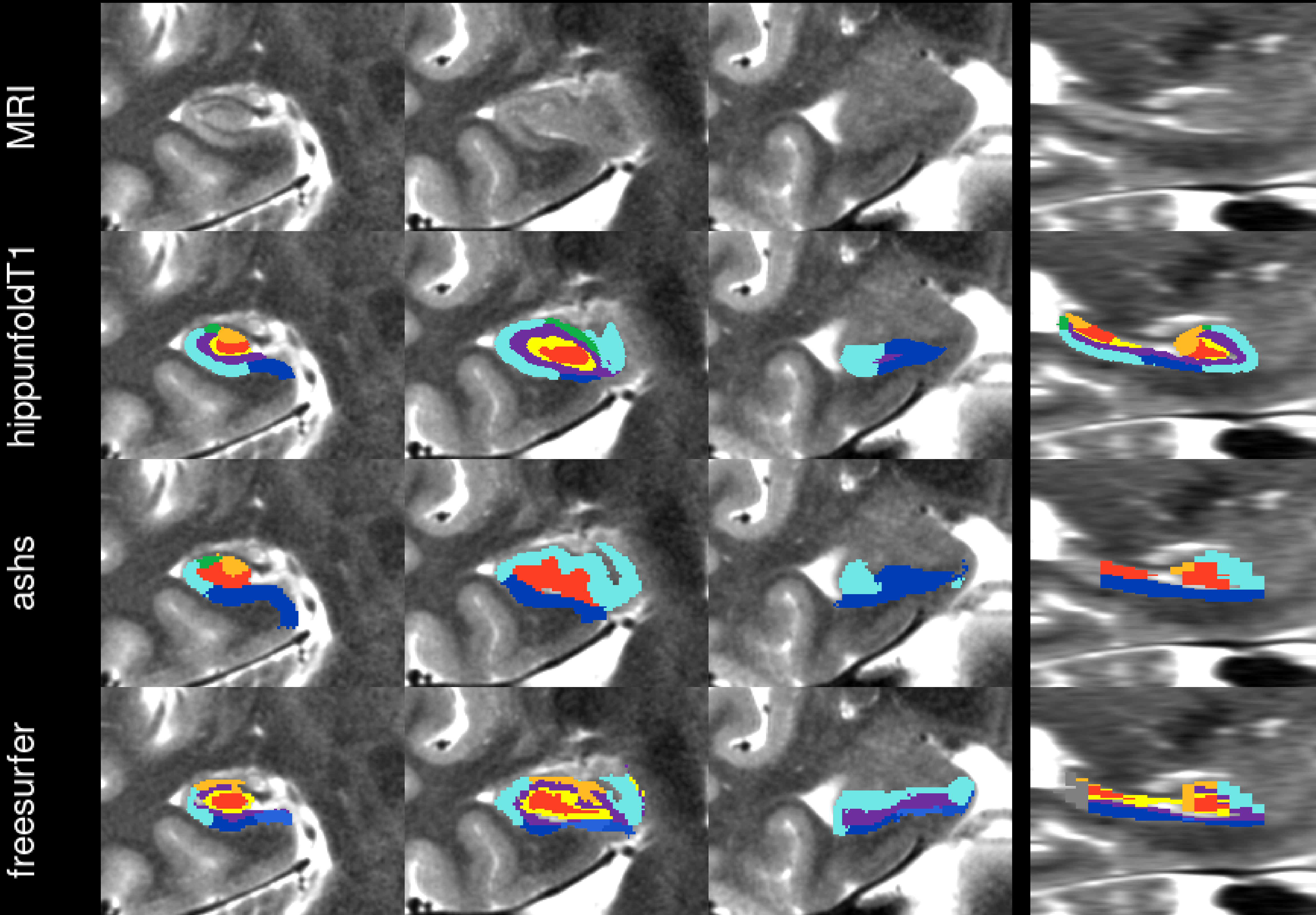

hemi=R,subject=9882308

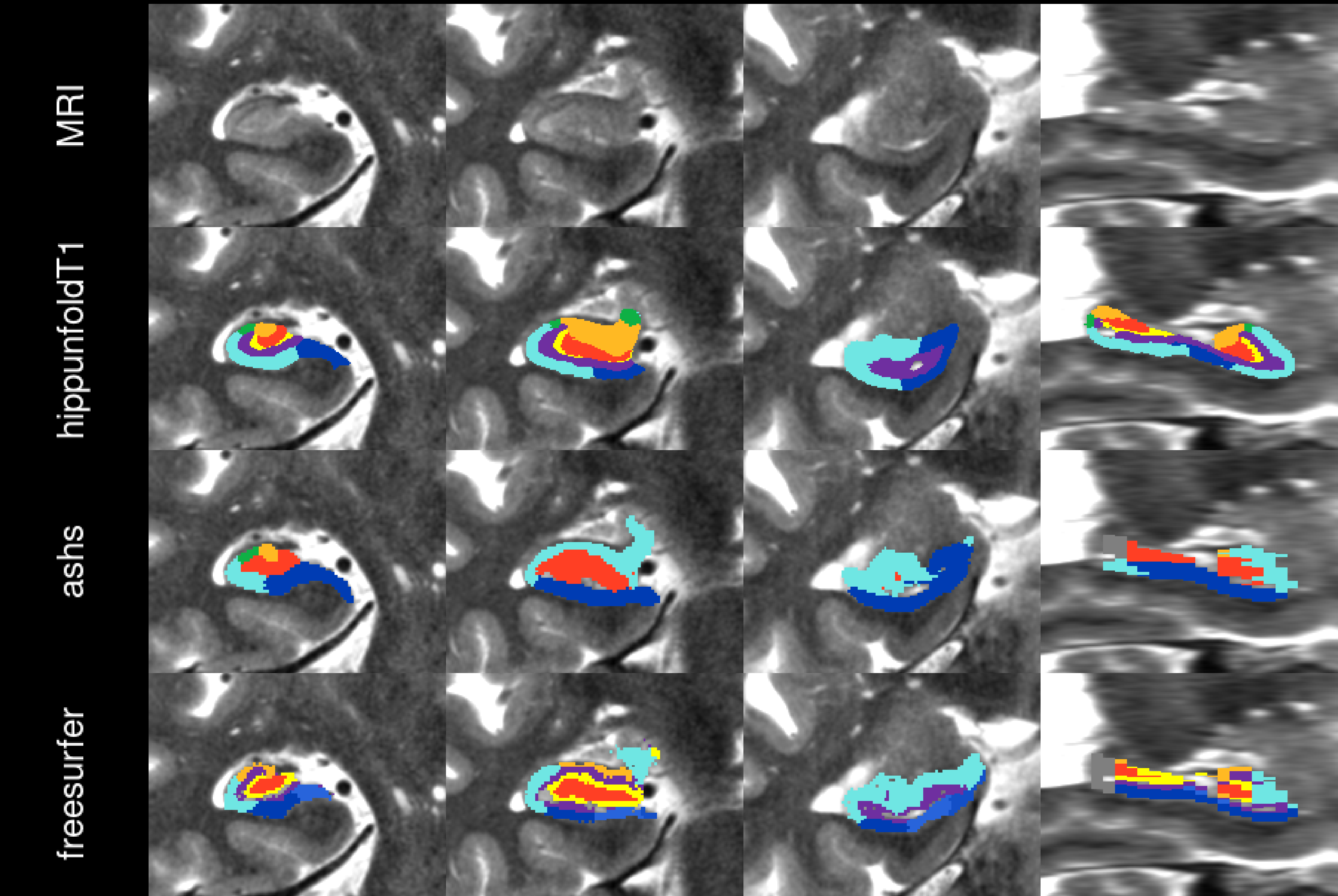

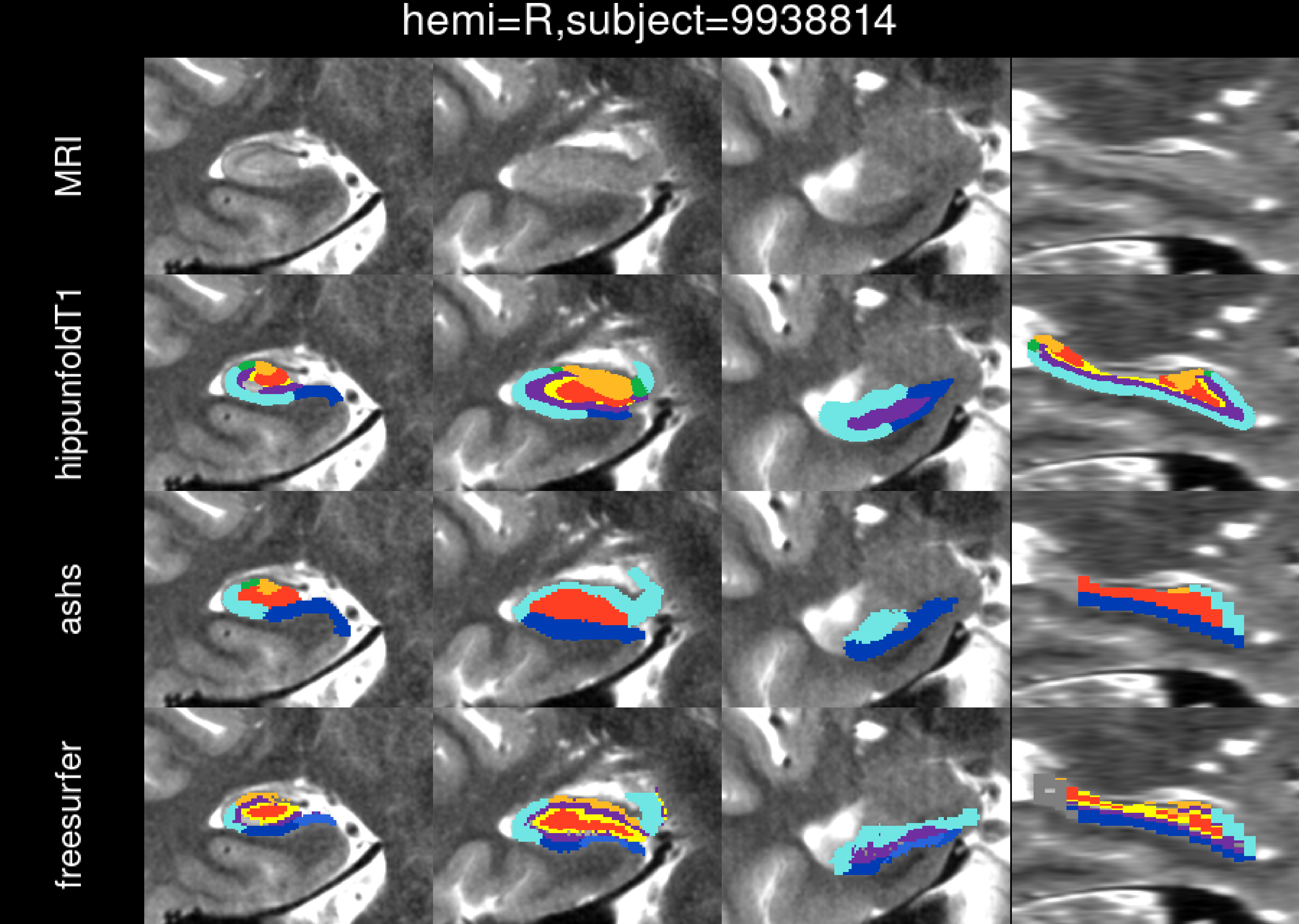
hemi=R,subject=9938814
MRI
hippunfoldT1
ashs
freesurfer

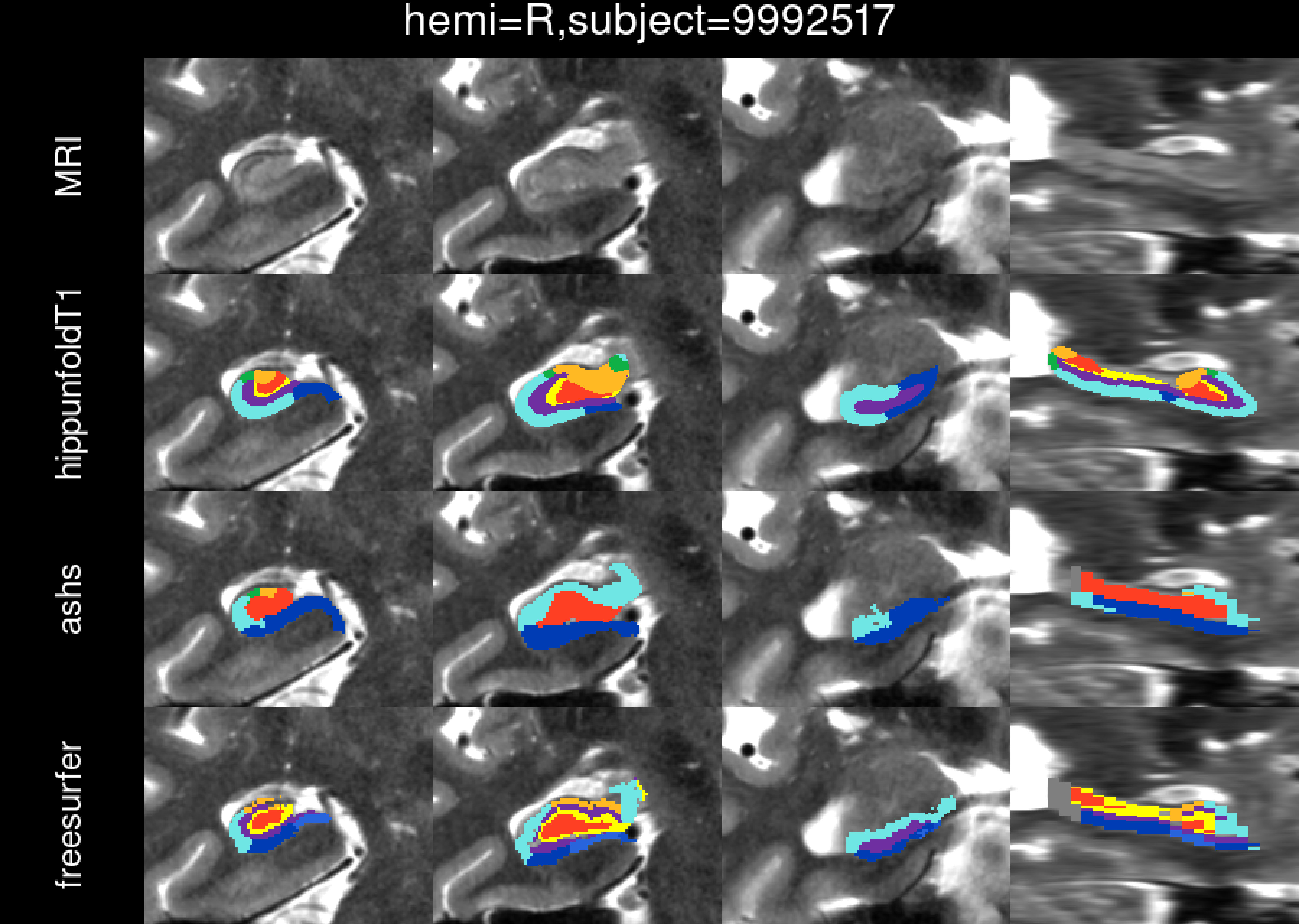
hemi=R,subject=9992517
MRI
hippunfoldT1
ashs
freesurfer
